# Biochemical communication with noisy feedback under energy constraints

**DOI:** 10.1101/2024.08.17.608427

**Authors:** Maximilian Gehri, Lukas Stelzl, Heinz Koeppl

**Affiliations:** Centre for Synthetic Biology, Technical University of Darmstadt, 64283 Darmstadt, Germany; Institute of Molecular Physiology, Johannes Gutenberg University Mainz, 55122 Mainz, Germany

## Abstract

Biochemical systems process signals through stochastic reaction dynamics that are inherently continuous in time and often exhibit memory, feedback, and nonequilibrium driving. At the same time, they are frequently modeled by effective reactions, e.g., multi-step processes such as transcription are treated as single events, while energetic bookkeeping is commonly omitted. Moreover, mesoscopic dissipation estimates are highly sensitive to whether coarse-graining and reservoir coupling are performed in a thermodynamically consistent way. Together, these features complicate the direct application of classical Shannon information theory and stochastic thermodynamics “as is” to biochemical reaction networks. This paper provides a self-contained route from first principles to a practically usable framework for studying information transmission through chemical reaction networks (CRNs) under energetic constraints. In particular, we discuss and extend the notions of classical information theory, methodically progressing to a level of generality that is necessary for the theme of causal communication through general CRNs. We then derive expressions for mutual information and directed information between bipartite CRN trajectories of disjoint sets of molecular species and show that the MI diverges without bipartiteness. These expressions account for cases in which different reactions are indistinguishable after projection to the respective subnetworks or where multiple driving mechanisms produce the same observable effect. We finally introduce a rigorous, operational Shannon-style continuous-time chemical communication model: messages are encoded by time-dependent chemostat protocols for a set of signaling molecules, the causal channel law is an immutable property of the reaction dynamics, and channel capacity is posed as an optimization over causal chemical encoders subject to thermodynamic costs of encoding and transmission. Trajectory information measures and the operational channel capacity are related by a Fano-type converse theorem. Complementary, we formulate the dual perspective of minimum-energy-per-bit necessary for reliable communication. A tractable promoter-switching example illustrates the practical application. Our work provides a formal and general framework to obtain universal energetic bounds for reliable communication in biochemical systems.

## I. INTRODUCTION

### Motivation

Energy consumption is integral to gene regulatory mechanisms and intracellular signal transduction in both prokaryotes and eukaryotes. For example, chromatin remodeling complexes in eukaryotes use ATP to maintain gene expression profiles [1, 2], while in prokaryotes, ATP conversion to cAMP provides a cofactor for transcription factors such as the catabolite-activating protein (CAP) [3]. These processes indicate that biological information processing is fundamentally constrained by energetic costs. This observation motivates a closer investigation of how upper bounds on energy dissipation shape the ability of cellular systems to process and transmit information.

Gene regulatory networks form an essential part of intracellular information processing [4], and information-theoretic tools are increasingly used to analyze their signaling capabilities and regulatory architectures [5, 6]. There is growing evidence that cells encode information in the spatiotemporal variation of populations of molecular species when responding to internal or external stimuli [7–10]. Classical information theory further suggests that in systems featuring feedback and network-mediated memory, meaningful communication can only be accurately characterized by information-theoretic quantities defined over entire trajectories [11]. Fundamental results of information theory, such as channel coding theorems, usually only apply when communication occurs over an extended period with a sustained flow of information. The coding theorems serve as asymptotic characterizations of information transmission channels, meaning they are operationally accurate only for very long, sustained transmissions. Specifically, a coding theorem connects the fundamental limit of how much information a system can process per time unit, ensuring that the probability of failure asymptotically approaches zero, with an optimization problem over trajectory information measures. Continuous-time Markovian stochastic chemical reaction networks (CRNs) offer a natural modeling framework in this context, capturing both the discrete and stochastic nature of molecular reactions. Any dynamically encoded information is thus accurately captured in the trajectories of a CRN. In complement, CRNs also allow an accurate characterization of thermodynamic costs.

### Knowledge gaps

Despite growing interest in information processing in biochemical systems, a unified communication-theoretic framework for chemical reaction networks is still missing [12]. Previous information-theoretic studies of biochemical signaling have mainly focused on quantifying information flow, statistical dependencies, and optimization principles, rather than on a general Shannon-style information-theoretic framework for communication in chemical reaction networks [13]. By contrast, the literature on information thermodynamics has primarily examined the complementary perspective, investigating how information is incorporated into non-equilibrium thermodynamic balances and the energy costs associated with measurement, erasure, and other information processing operations [14, 15]. However, these information-theoretic and stochastic-thermodynamic perspectives have not yet been unified in a general framework that addresses the central information-theoretic questions concerning communication, namely, the fundamental performance limits of information encoding for reliable transmission over a noisy communication channel. This is especially true for continuous-time biochemical systems, where information is carried by stochastic trajectories, feedback is inherently part of the dynamics, and asymptotic performance criteria, such as channel capacity, are thus rigorously defined in terms of reliable transmission over an infinite timeframe rather than static input-output relation.

Recent work has begun to connect stochastic thermodynamics and Shannon information theory in the context of driven copying channels [16, 17]. Thereby, channel capacity is understood as a symbol-wise characterization rather than a continuous-time rate. This viewpoint is valid for discrete-time memoryless channels that are derived under a timescale separation assumption for consecutively encoded symbols. While this approach provides an important complementary perspective, it does not address the specifics of biochemical information processing through chemical network structures and the associated concrete energy dissipation modeling.

More generally, the lack of a unified communication-theoretic framework for continuous-time biochemical systems is also limiting a detailed understanding of energy as a relevant resource constraint in biological signal processing. In particular, the question of energy per reliably transmitted bit has not yet been formulated in a thermodynamically faithful manner in this setting. Part of the challenge is conceptual, but part is also technical: a precise treatment requires trajectory-level information measures and dissipation estimates together with a careful measure-theoretic formulation of causality and feedback for continuous-time stochastic processes. Standard memoryless-channel models and symbol-wise mutual information alone are therefore not sufficient.

### Contributions

We develop a unified framework for biochemical communication that connects Shannon’s information theory of communication, stochastic thermodynamics, and stochastic chemical reaction networks. Its central aim is to pave the way for rigorous coding theorems for chemical communication under energetic constraints, that is, for statements identifying the maximal rate of asymptotically reliable chemical encoding and relating this operational quantity to optimized trajectory-level information measures. The dual perspective of such coding theorems under energy constraints is the minimum energy required to ensure reliable transmission of information over the same channel. We show that these perspectives are fundamentally interconnected.

On the thermodynamic side, we provide a self-contained trajectory-level formulation of stochastic thermodynamics for open chemical reaction networks in a notation aligned with the information theory for such networks. Sec. III makes the modeling choices needed for thermodynamically faithful dissipation estimates explicit. Stochastic thermodynamics is presented in a common chemical reaction network notation that later allows energetic constraints on chemical encoding and transmission to be imposed in a way that is consistent.

On the information-theoretic side, we formulate causal communication systems with general time structure in physical time, covering both discrete- and continuous-time models in Sec. IV. An important feature of this formulation is that communication is described in natural time rather than through an imposed symbol-wise indexing, which is essential for continuous-time biochemical channels with memory and feedback. A key conceptual contribution is the separation of deterministic encoding specifications for messages from a fixed stochastic feedback mechanism, that generates the actual stochastic channel encoding based on the output history and the message-encoding. This distinction clarifies the operational structure of noisy-feedback communication by separating the fixed system from the choice of encodings: the channel law and the feedback architecture are fixed, and coding theorems concern the existence of reliable codes that map from a finite set of messages to a finite codebook. Such a codebook is a subset of deterministic encoding specifications that satisfy a prescribed set of coding constraints, for example, a limited energy dissipation rate. This, in turn, allows us to formulate principled coding-theorem statements at the level of generality required for biochemical communication channels.

Independently of the operational communication framework, we next develop the information measures required to describe chemical information processing in continuous time. In Sec. V, we adapt Weissman *et al*.’s general definition of continuous-time directed information (DI) to natural physical time and establish the corresponding structural identities. In Sec. VI, we then derive mutual- and directed information expressions for trajectories of disjoint CRN subnetworks, including the effect of indistinguishable reaction channels under projection and thereby generalizing previously established expressions [18]. In particular, we show that mutual information (MI) is well defined under a bipartiteness condition, whereas without bipartiteness it generally diverges. Together, these sections provide the detailed understanding of trajectory-level information measures needed later for biochemical communication models.

Building on these ingredients, we then formulate a rigorous continuous-time chemical communication model based on stochastic chemical reaction networks in Sec. VII. In this model, which is visualized in Fig. 1, each message is encoded into time-dependent signaling molecule trajectories, which are equivalently understood as externally manipulated chemostat protocols from a thermodynamic perspective. These protocols modulate a prescribed set of reaction rates which determine the stochastic dynamics of a chemical sub-population X, whose trajectories consitute the noisy channel encoding. The trajectories of another set of molecules Y represent the channel output, given that it has no species in common with the encoding population X and the joint dynamics of X and Y is bipartite. We show how this model induces the communication-theoretic notions of (i) the channel law, (ii) the feedback encoder architecture and (iii) deterministic encoding specifications in a way that is consistent with the causal communication-system formalism developed before in Sec. IV. This, in turn, allows us to (i) identify the information measures relevant for coding theorem statements in CRNs with noisy feedback, (ii) relate the construction to Massey’s [11] fundamental notion of causal feedback-communication systems, and (iii) formulate heuristic coding-theorem statements under different design constraints, including partially variable CRN structures.

**FIG. 1.**
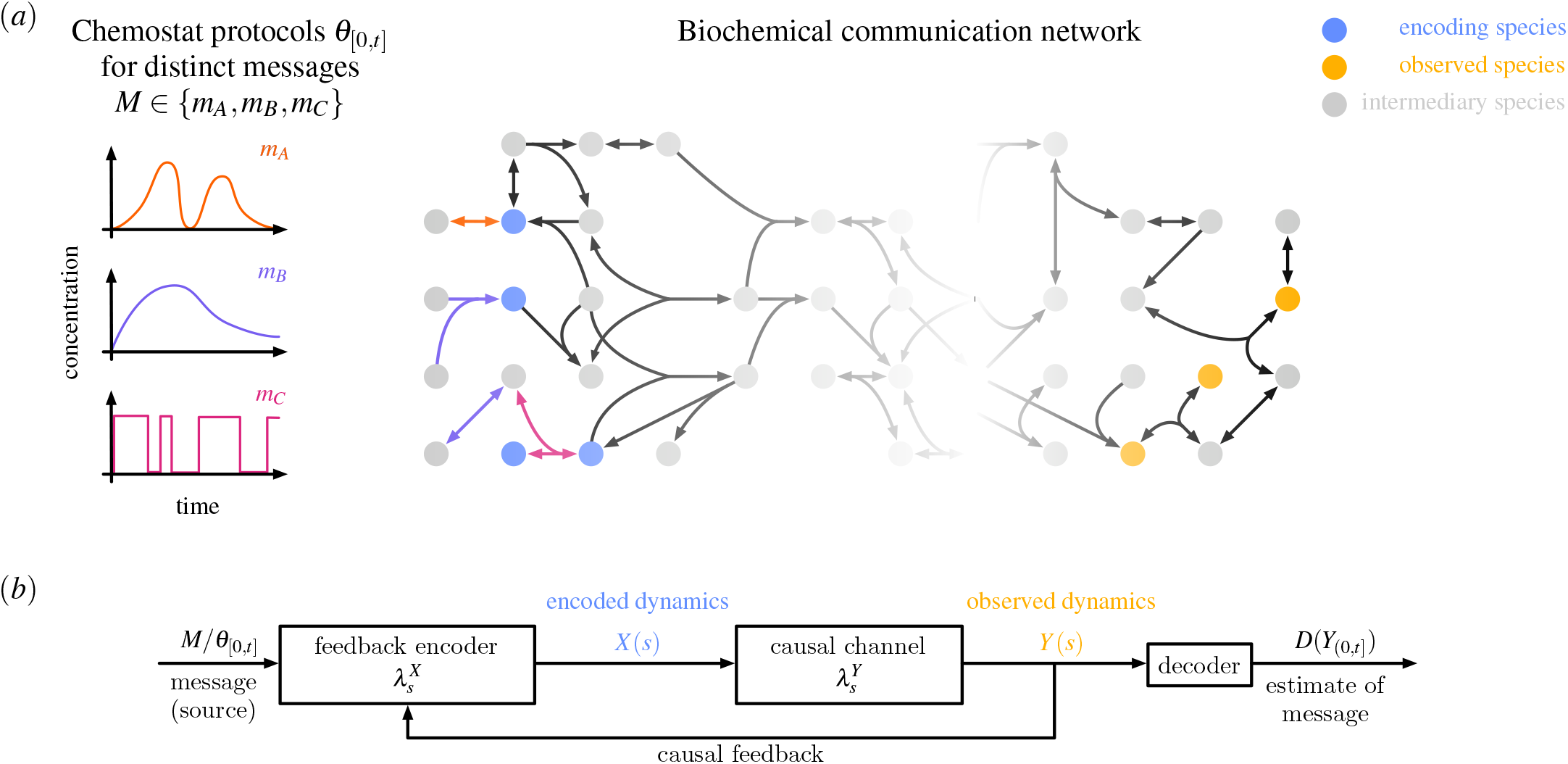
Biochemical communication through a chemical reaction network. **(a)** A random message *M* ∈ {*m*_*A*_, *m*_*B*_, *m*_*C*_} selects a sourceside signaling protocol *θ*_[0,*t*]_, represented here by deterministic, time-dependent concentration profiles of distinct signaling chemostat species (orange, purple, magenta). These chemostat protocols act on designated encoding species *S*_*X*_ (blue) within the biochemical communication network, whose dynamics then propagate through intermediary species (gray) to the observed species *S*_*Y*_ (yellow). **(b)** Corresponding communication-theoretic representation of the same system with time-wise causal information flows for every *s* ∈ [0, *t*] within the transmission interval. The source *M/θ*_[0,*t*]_ enters a feedback encoder through time-dependent modulation of channel-encoding reaction-intensities 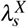, generating the stochastic encoded dynamics *X* (*s*). The channel, characterized by output reaction-intensities 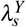, causally generates the noisy observed dynamics *Y* (*s*) from the histories *X*_[0,*s*)_ and *Y*_[0,*s*)_. The knowledge of these histories is also integrated in causal feedback encoding. A decoder *D*, acting on the full observation *Y*_(0,*t*]_, returns an estimate of the message.

Finally, in Sec. VIII we introduce thermodynamically principled energetic constraints on chemical population encoding and transmission through reaction networks. On this basis, we prove a Fano-type converse theorem for capacity-cost functions, which connects the operational coding problem under energetic constraints to maximized trajectory-level information measures. We then formulate the dual perspective of the minimum energy dissipation required per reliably transmitted information unit. Minimal promoter-switching models with analytically tractable information measures illustrate the framework in practice and expose modeling pitfalls that should be avoided in thermodynamically and information-theoretically faithful applications.

This work is accompanied by two supplementary files: Supplement S1 contains detailed technical derivations relevant to the key results presented in Sections V, VI, VII and VIII. Supplement S2 contains further background material on stochastic thermodynamics of chemical reaction networks, particularly for readers whose main field of research is different. Among other topics, it offers a more detailed account of thermodynamic coarse-graining and demonstrates how mesoscopic descriptions align with both classical microscopic Hamiltonian formulations and macroscopic thermodynamics. Additionally, it reviews the connection between entropy production and statistical time reversal, presenting it in the formal mathematical style used throughout.

### A. Connection to prior work

As both, purely thermodynamic and purely information-theoretic prior work, are well accounted for in Sections III and IV, we discuss here only prior work in the context of biochemical information transmission.

Although Barlow [19] has already hypothesized in the early 60s that biological signal processing systems may function as communication systems, applications of information theory in systems biology have only started to develop into a research field for around two decades [12]. Research in this field has addressed a range of problems, including the analysis of connectivity, sensitivity, and cross-talk in signaling pathways [20]; the identification of optimal cellular adaptation strategies and their adaptation targets [21]; and the study of gene regulatory motifs at steady state [6], at specific time points [22], or in terms of dynamic features such as amplitude and frequency [23]. In the context of biochemical signal transduction, Tostevin and ten Wolde first considered the MI between trajectories, recognizing that relevant information may be contained in the temporal dynamics of the input signal [24]. They provide the mutual information rate (MIR), i.e., the asymptotic differential information gain, under a linear-noise Gaussian approximation. However, the idea of quantifying information between dynamic biological stimuli has a longer history in the context of neural spike trains [25] and continuous-time path information measures, in general, are widely attributed to Pinsker [26]. Furthermore, the trajectory relative entropy associated with Markovian jump processes [27] (see also [28, Eq. (2)]) has been utilized in the contexts of model approximation and statistical inference. In biochemical signaling, others have considered mutual information between a static concentration variable and a time series observation [29, 30]. Lestas *et al*. used MI between trajectories to lower bound the cost of suppressing molecular fluctuations via feedback under a Langevin approximation [31], later extended by Nakahira *et al*. who identified continuous-time DI as the appropriate measure [32]; however, the stringency of these lower bounds has been questioned by Parag [33].

An expression for the mutual information between the trajectories of two species of a class of biochemical reaction networks based on filtered intensity processes was first given by Duso and Zechner [34] and later generalized by Moor and Zechner [18]. Moor and Zechner further showed that the trajectory mutual information decomposes into a pair of continuous-time transfer entropies [35]. The transfer entropy is in fact identical to the DI under certain conditions (see Appendix B for a brief comparison). In this paper, we further generalize the expression for the mutual information in [18] to encompass the information flow between disjoint subnetworks without simultaneous reactions in the two sets. Without this bipartite-type requirement for the subnetworks of interest, the MI has been argued to diverge [36]. In contrast to Spinney *et al*. [35], we equivalently derive the continuous-time DI for CRNs via the extremal-based approach of Weissman *et al*. [37] instead of explicitly constructing a Radon-Nikodym derivative. Directed information remains well defined without the bipartite assumption, whereas mutual information generally does not.

Sinzger *et al*. and Gehri *et al*. advanced techniques for the evaluation of filtered intensity processes to facilitate analytically and numerically computing the mutual information of the Poisson-type channel modulated by a telegraph process, which can be regarded as a minimal module of a CRN [38– 40]. In particular, [40] presents a custom filtering and projection procedure for semi-Markov processes to isolate sufficient statistics for the history dependence of the transition intensities of projected semi-Markov processes. This, in turn, facilitates the evaluation of MI and DI for a class of small chemical reaction models. Moor and Zechner [18] proposed several MI computation methods in the context of CRNs: a combination of stochastic simulation (SSA) [41] and moment closure approximation, a quasi-exact approach combining SSA and numerical integration, and an analytical approximation. Lastly, Reinhardt *et al*. developed an exact Monte Carlo method called path weight sampling to compute the MI between trajectories [42], which has recently been adapted for the computation of transfer entropies [43]. Analytical derivations of information channel capacities for the Poisson-type channel under peak and average-constraints on the message waveforms have been put forth by Kabanov [44], Davis [45] and Frey [46], where Frey also considered marked Poisson-type channels. For small biology-motivated systems, information channel capacities have been analytically derived by Thomas and Eckford [47–49].

The combination of thermodynamics and information theory of biological information processing has been discussed in several studies. Some early works proposed the entropy production rate (EPR) as a measure of the energetic cost of cellular information processing and decision-making [50, 51]. Later, models of diffusive transport were used to assess the thermodynamic cost of moving information-carrying molecules across cellular compartments [52]. More recently, stochastic thermodynamics has been combined with communication theory to analyze finite-state bipartite Markov systems at nonequilibrium steady state, treating each channel use as a readout of a steady-state output symbol [16]. In that framework, entropy production is not a constraint but a quantity co-optimized alongside information capacity, revealing trade-offs in the steady state. Another approach considers a discrete memoryless channel driven by an i.i.d. binary input, where the output is observed once sufficient time has passed such that input and output are significantly correlated [17].

### B. Notation and conventions

In the following let (Ω, ℱ, ℙ) be a probability space with complete filtration ℱ = {ℱ_*t*_}_*t* ≥0_. The symbol denotes the join of *σ*-algebras, i.e., the smallest *σ*-algebra containing all elements of the individual *σ*-algebras [53]. For any measurable function *f* : Ω → *X*, where (*X*, Σ_*X*_) is a measurable space, *f* ^−1^ : Σ_*X*_ → ℱ generally denotes the preimage. For basic notions of probability and measure theory, we refer the reader to the textbooks by Kallenberg [54] or Klenke [55]. For the reader’s convenience, we have added a small review of relevant measure theoretic notions in Supp. Sec. S1.1.

#### Paper-internal conventions

We use 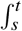 to denote integration over the half-open interval (*s, t*] (with *s < t*); this convention is important for integrals with respect to counting processes. All stochastic pure jump processes considered here (including counting processes and CRN copy-number processes) are assumed to have càdlàg trajectories, i.e, trajectories that are right-continuous and have left-limits. To distinguish between conditional probabilities and causally conditional probabilities we use ℙ(· | ·) and ℙ(· ∥ ·), respectively.

In the context of information theory of communication, we often refer to *X* and *Y* as the channel input and output, respectively; however, this identification is not strictly enforced throughout. For quantities in stochastic thermodynamics we follow the IUPAC conventions [56].

## II. PRIMER: MARKOVIAN STOCHASTIC CHEMICAL REACTION NETWORKS

A powerful formalization of the description of Markovian chemical reaction networks was provided by Kurtz and Anderson [57]. They demonstrated how counting processes relevant to these networks can be represented using Poisson-type processes, leading to a random time-change representation. Therewith, they obtain a stochastic equation for continuous-time Markov jump processes, i.e., their state-level description. Our introduction to CRNs largely follows their work, supplemented by some thermodynamically relevant concepts.

A stochastic chemical reaction network (CRN) consists of a finite set of chemical species *S* with labels U = {U_*d*_ | *d* ∈ *S*} and a finite set of reaction channels ℛ with labels ℜ_*r*_, *r* ∈ ℛ, with stoichiometric balance equations

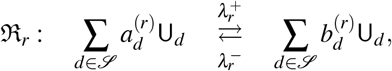

wherein 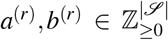 represent the number of reacting molecules of each species and the number of product molecules, respectively, for the forward reaction, and vice versa for the backward reaction. The difference of product and reactant species of ℜ_*r*_ forms the stoichiometric change vector *ν*_*r*_ = *b*^(*r*)^ − *a*^(*r*)^ of the forward reaction. For each reaction channel, the backward reaction microscopically reverses the forward reaction. We assume that all reactions are elementary [58], i.e., there are no reaction intermediates and each reaction has only a single transition state in the reaction coordinate diagram as illustrated in Fig. 2 (a).

**FIG. 2.**
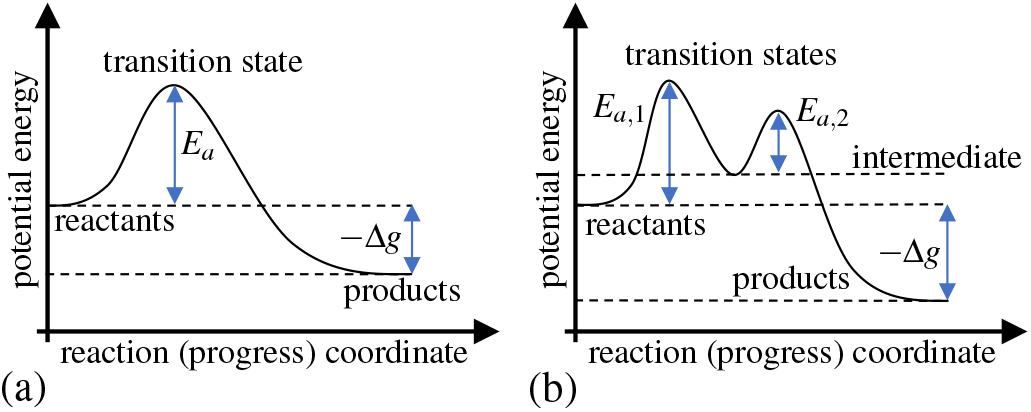
Exemplary reaction coordinate diagrams. **(a)** shows an elementary reaction, and **(b)** a non-elementary reaction with a single intermediate. *E*_*a*_ denotes the activation energy of the reaction, while −Δ*g* is amount of energy released by the reaction.

Let 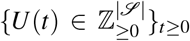 be the continuous-time Markov chain of the copy number vector of the chemical species U. Throughout, we sometimes assume that the rates of the reactions, commonly referred to as propensity functions, obey stochastic mass action kinetics [57, 59]

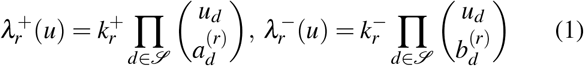

with 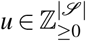 and stochastic rate constants 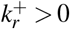 and 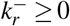. A conversion of the discrete molecule numbers to concentrations in a constant, finite solution volume V *>* 0 gives rise to another pair of rate constants 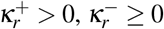, such that

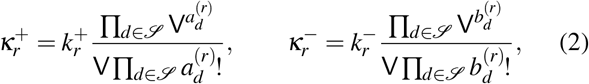

due to consistency of physical units in both the deterministic ODE model (of the concentrations) and the probability evolution equation of the stochastic model [57, 60]. 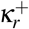 and 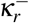 can be identified with the rate constants of the deterministic law of mass action by taking the macroscopic limit. The mass action assumption can be dropped in the following if it is not explicitly used. A reaction (channel) ℜ_*r*_ is called microscopically reversible if 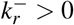, or more generally if 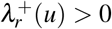, implies 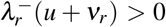 and vice versa for all *u*. A CRN is called microscopically reversible if all reactions are microscopically reversible. In the following, a CRN is assumed to be microscopically reversible unless otherwise noted.

For each reactions of channel ℜ_*r*_ we denote the counting processes of reactions in the forward and reverse direction as 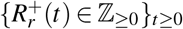 with 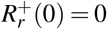 and 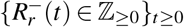 with 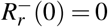, respectively. These can be represented via random time changes [57]

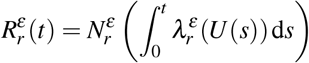

of 2|ℛ| independent unit-rate Poisson processes 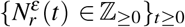 with 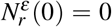 for *r* ∈ ℛ and *ε* ∈ {+, −}. Then the random state *U* (*t*) of the CRN with random initial value *U* (0) obeys

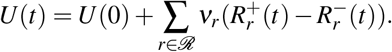

The probability mass function *p*(*u, t*) = ℙ(*U* (*t*) = *u*) follows the chemical master equation [61, 62]

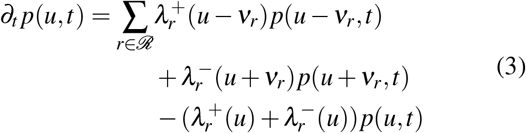

Before proceeding, we recapitulate the model assumptions made so far, as they are fundamental for stochastic thermodynamics of CRNs. Assuming a Markov process description means that the future dynamics of the mesoscopic system is sufficiently described by the current state and does not depend on the past. In particular, the future dynamics do also not depend on the particular microstate, the system is in, but just on the current mesostate *u*.

The above assumptions about the system state *U* have implications for the environment in which the reaction network is embedded. Since the propensity functions are independent of time and space, the embedding environment must be seen as thermally equilibrated and spatially homogeneous from the perspective of the chemical species, which is often referred to as well-mixedness [59]. That is, the surrounding of each molecule of *U* homogenizes spatially on a much faster time scale than the reactions occur, or in other words, diffusion is fast relative to the volume V and inter-reaction times. In addition, the temperature of the solvent is constant over time and the volume V. Similarly, fluctuations in the solvent’s molecular composition, such as changes in pH or ionic strength, are assumed to be small enough to leave the mesoscopic rate constants effectively time-invariant.

In later sections, isothermal–isobaric conditions will be adopted as a natural assumption for biological cells. Under such conditions, the stochastic rate constants 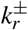 in (2) could, in principle, depend on the mesostate *u* due to volume fluctuations. However, we assume these fluctuations are negligible, allowing 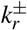 to remain effectively constant, except in scenarios involving slow, time-dependent changes such as cellular growth.

According to Gillespie, in the light of collision theory the elementary reaction assumption is plausible up to bimolecular reactions and the dynamics is shown to follow stochastic mass action kinetics under the above assumptions on the environment [59]. In other cases, in particular when the chemical species represent different conformations of polymers as is common in gene expression models, a more careful justification of the elementary reaction assumption and the Markov assumption may be required. The relevance of the elementary reaction assumption for stochastic thermodynamics of CRNs is discussed at the end of Sec. III A.

## III. PRIMER: STOCHASTIC THERMODYNAMICS FOR CHEMICAL REACTION NETWORKS

This section establishes consistent notations for heat, work, and entropy production at the trajectory level for open chemical reaction networks with multi-reservoir couplings. We proceed from closed CRNs to chemostat-coupled open CRNs and then introduce the corresponding trajectory-level thermodynamic quantities before turning to stationary dissipation rate estimates and the effects of effective model descriptions. In particular, we explain how modeling choices, such as effective reactions and parallel reaction channels, lead to effective dissipation estimates that may systematically underestimate the true dissipation. Stochastic thermodynamics for CRNs has been introduced by Schmiedl and Seifert [63]. We also refer to Rao and Esposito [64] as a more recent CRN-specific reference work and the textbooks by Peliti and Pigolotti [65] and Seifert [66]. Readers familiar with stochastic thermodynamics of open CRNs may refer directly to Secs. III E for the discussion of the effects of coarse-graining on dissipation estimates and to Supp. Sec. S2.3 for a link between common multi-reservoir entropy production rate expressions and local detailed balance equations of open CRNs.

### A. Closed stochastic chemical reaction networks

A thermodynamic system is called closed if it exchanges energy, but not matter, with its environment. In the CRN setting, we use the term more specifically: a CRN is called closed if its core species U do not exchange molecules with the surrounding solvent. We therefore partition the species in solution into the reactive core species U and solvent species S, where the latter do not appear as reactants or products in reactions involving U. Assuming that the solvent remains in thermal and chemical equilibrium at constant temperature *T*, it acts as both a heat bath and an equilibrated environment for the CRN.

The combined solution U ∪ S is treated as an isothermal closed system whose classical, non-reactive microscopic dynamics are governed by a Hamiltonian H. A mesoscopic description becomes valid under a time-scale separation in which microstates equilibrate rapidly compared with the slower transitions between mesostates. This local-equilibrium assumption is the key coarse-graining step in stochastic thermodynamics: it allows each mesostate *u* to be assigned a welldefined thermodynamic potential [65]. At constant volume this potential is the Helmholtz free energy *f* (*u*), whereas under constant pressure – as is more appropriate for biological systems – it is the Gibbs free energy *g*(*u*). Since the systems of interest here operate under isothermal–isobaric conditions, we use *g* throughout. A derivation from an underlying Hamiltonian model is reviewed in Supp. Sec. S2.1; alternatively, *g*(*u*) can be obtained from equilibrium statistical mechanics of ideal dilute solutions provided that all solute species are included [64, App. A].

The zeroth law of thermodynamics for chemical reaction networks dictates that a closed, microscopically reversible CRN that accounts for all solute species will always relax to equilibrium. The equilibrium distribution depends on the initial state *U* (0) of the system and differs between stoichiometric compatibility classes [67], i.e., the sets of all states that are connected via the reaction channels ℜ_*r*_, *r* ∈ ℛ. In the following we assume that any initial distribution assigns positive mass only to states of a single stoichiometric compatibility class, since otherwise the limiting distribution

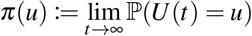

can become a mixture of the equilibrium distributions of individual compatibility classes. Equilibrium statistical mechanics further dictates that the equilibrium distribution of a closed CRN under isothermal–isobaric conditions is (i) a Gibbs-Boltzmann distribution *π*(*u*) ∝ *e*^−*βg*(*u*)^, where *β* = (*k*_B_*T*)^−1^, and (ii) it satisfies detailed balance

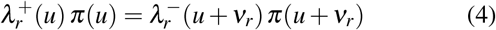

for all admissible states *u* and all reaction channels ℜ_*r*_, since we assume throughout that every modeled reaction channel is microscopically reversible.

To describe the energetics associated with each reaction channel ℜ_*r*_, we introduce forward and backward difference operators acting on functions of mesoscopic states.

#### Definition 1

*For each reaction channel* ℜ_*r*_ *we introduce the forward and backward difference operators* 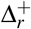 *and* 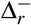, *respectively, such that for any function* 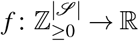 *we define*

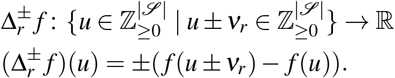

The definition implies 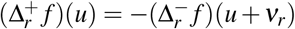. Starting from state *u* in forward direction, the change in Gibbs free energy of the state associated with reaction ℜ_*r*_ is given by

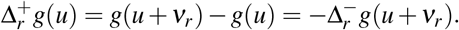

Substituting the Gibbs–Boltzmann distribution into (4) yields the local detailed balance condition, or thermodynamic consistency relation, for closed CRNs

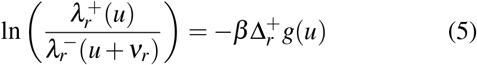

for all admissible states *u* and all microscopically reversible reactions ℜ_*r*_. Equation (5) relates the kinetic parameters of the model to the underlying thermodynamic potential. Therefore, it does not depend on the initial condition and also holds during a transient relaxation to equilibrium.

However, the consistency relation can be broken if reactions are not elementary, which violates the fundamental coarse-graining assumption that mesostates *u* define local equilibrium states [68]. Any metastable or intermediate state along the reaction progress coordinate of a non-elementary reaction (cf. 2 (b)) must be accounted for as a species of the CRN. Deviations from the well-mixed assumption caused by individual elementary reaction events are neglected under the implicit assumption of instantaneous diffusion, which ensures local equilibrium within the finite reaction volume. Consequently, the local equilibrium assumption of stochastic thermodynamics aligns with the kinetic discrete-state Markov process model.

A common modeling oversight, only recently emphasized in [69], is the misidentification of forward and backward reactions that in fact correspond to distinct microscopic channels. In such cases, the thermodynamic consistency relation is, quite naturally, not valid. This issue is particularly relevant for models of gene expression, where the production and decay of biopolymers such as mRNA and proteins typically do not represent elementary reactions and are not microscopic reverses of one another.

With a slight abuse of notation, we denote the heat dissipated into the environment due to reaction ℜ_*r*_ by 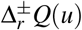. Strictly speaking, *Q*(*u*) is not always a well-defined state function, as heat is a path-dependent quantity. We incorporate the operator into the definition since it satisfies the property 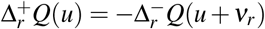.

As reviewed in Supp. Sec. S2.1, the mesoscopic Gibbs free energy *g*(*u*) of the solution can be decomposed into its mesoscopic enthalpy h(*u*) and the internal entropy *h*_i_(*u*) of the mesostate *u*, averaging over the fast equilibrated degrees of freedom

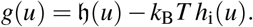

Following the mesoscopic formulation of the first law of thermodynamics under the local equilibrium assumption (cf. Supp. Sec. S2.1), the heat released to the environment during a state transition can be identified with the decrease in enthalpy:

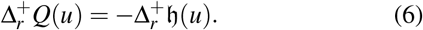

Substituting this into the free energy change yields

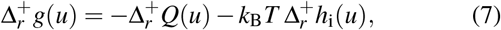

which shows that the local detailed balance condition (5) links the kinetic parameters of the model with both external (heat exchange) and internal entropy changes associated with each reaction.

### B. Open chemical reaction networks – Coupling to chemostats

An open CRN allows the reactive exchange of molecules with external reservoirs. Relative to the closed case, the only new thermodynamic ingredient is that each reaction event may now be driven by chemical work supplied by the reservoirs. Here, we assume that these reservoirs are part of the solvent, which itself must be maintained by an external environment in order to justify the steady-state assumption inherent to any reservoir description.

The core system U be coupled to *L* chemostats Z_1_, …, Z_*L*_, which are part of the solvent species S and immediately equilibrate after participation in reactions of the core system U. The energetic coupling is illustrated in Figure 3. We assume that the copy numbers of chemostat molecules in the solvent are sufficiently large that they can be represented by fixed real-valued concentrations *z* = (*z*_1_, …, *z*_*L*_). In the biological setting, this corresponds to a homeostatically maintained cellular environment. Time-dependent chemostat protocols have also been considered in the literature [63, 64, 70]. For clarity, we first treat constant chemostats and return to time-dependent protocols at the end of this section.

**FIG. 3.**
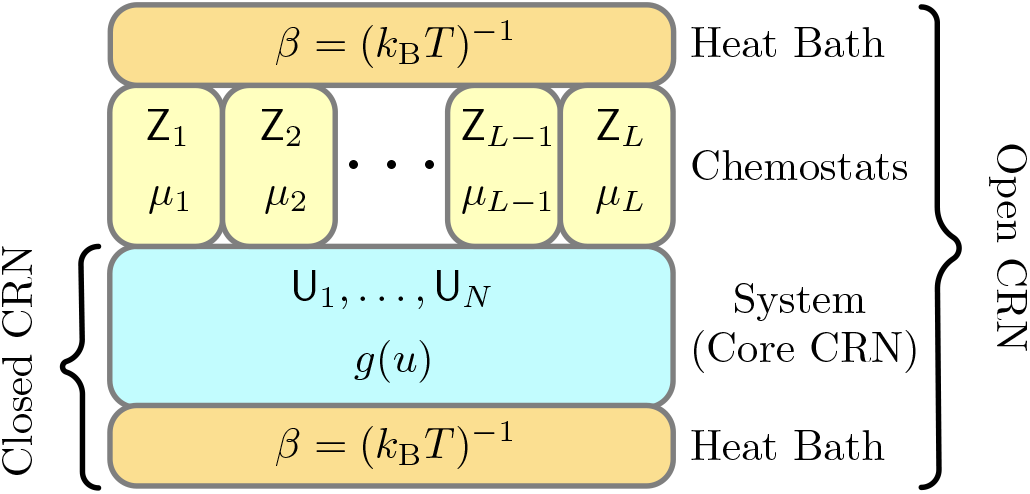
Visualization of an open CRN. The core system U is coupled to a number of chemostat species Z_*l*_ of chemical potential *µ*_*l*_ via reactions. The solvent acts as a common heat bath with temperature *T* for the core system and all chemostats. Thus, each horizontal layer in the illustration can exchange energy with its vertically adjacent layers.

The open CRN has stoichiometric balance equations

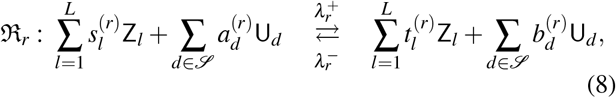

where *γ*_*r*_ = *t*^(*r*)^ − *s*^(*r*)^ is the stoichiometric change vector of the chemostat species associated with ℜ_*r*_. Immediately after each reaction, the chemostat is assumed to correct the copy number of all chemostat species. Consequently, under the assumption of stochastic mass action kinetics, the propensity functions of the coupled reactions maintain the form (1) with the adjustment that the stochastic rate constants now depend on the chemostat concentrations insofar as they incorporate a deterministic law of mass action

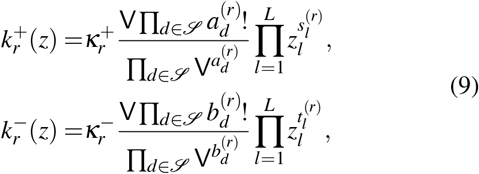

where the units of the deterministic rate constants 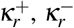, are adapted accordingly. Hence, for any set of Markovian kinetic laws of the form 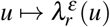 (i.e., not necessarily mass action kinetics), *z* is a fixed parameter of the propensity functions. Therefore, the probability evolution is still described by (3).

To describe the energetics of each reaction channel ℜ_*r*_, let *µ*_*l*_ denote the chemical potential of chemostat species Z_*l*_, that is, the Gibbs free-energy change associated with adding one molecule of Z_*l*_ to the reservoir. The chemical work performed on the system by the chemostats via reaction ℜ_*r*_ is then

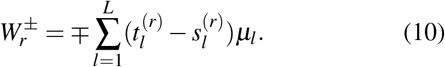

Accordingly, the heat dissipated to the environment in a reaction event is

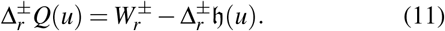

Thus, compared with the closed case, the only modification is that the dissipated heat now includes the chemical work exchanged with the chemostats. Intuitively, the dissipated heat is the chemical work done on the system but not “stored” in the systems enthalpy. Eq. (11) reduces to Eq. (6) when the chemostat species Z are incorporated into the closed system rather than treated as externally controlled part of the solvent S.

In open CRNs the thermodynamic consistency relation then becomes

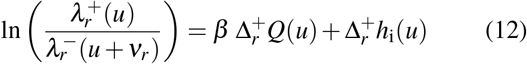

for all admissible states *u* and all reversible reactions ℜ_*r*_. The local detailed balance relation (5) together with (7) follows the same expression as (12), where however the identities of the dissipated energies 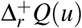 per reaction differ. Thus, (12) is more general in the sense that it potentially accounts for chemical work through (11) in addition to enthalpy changes. A detailed mesoscopic derivation of (12), including multiple reservoirs, is given in Supp. Sec. S2.2.

In Supp. Sec. S2.4 we further review the consistency of in the macroscopic limit for an ideal dilute solution, obeying mass action kinetics.

More generally, a CRN can be perturbed away from equilibrium not only by constant chemostat couplings, but also by explicitly time-dependent external control. Following the terminology of [65], these perturbations fall into two (not mutually exclusive) categories: (i) “Driving”, which refers to coupling the system to an external agent that exchanges energy with it during state transitions, as in the case of chemostats; and (ii) “Manipulation”, which involves time-dependent control of system parameters, such as changing chemostat concentrations or variations in the solution volume, for example due to cellular growth. Note that this terminology is not used consistently across the literature.

In the presence of a time-varying chemostat protocol 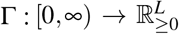, the propensity functions become explicitly time dependent through Γ(*t*), yielding a time-inhomogeneous Markov jump process with propensities 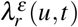. The structure of the chemical master equation (3) remains unchanged after replacing the propensity functions by their time-dependent counterparts. Likewise, thermodynamic quantities such as *g*(*u, t*) and *µ*_*l*_(*t*) acquire explicit time dependence, yielding the generalized local detailed balance condition

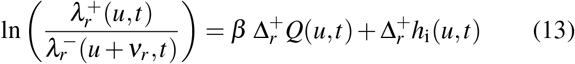

together with the reaction-wise enthalpy balance

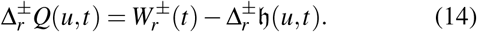

Eq. (13) is derived by introducing an auxiliary instantaneous stationary distribution *π*(*u, t*). This distribution is defined as the limiting distribution achieved by hypothetically freezing the protocol at Γ(*t*) and allowing the system to relax. Importantly, *π*(*u, t*) does not follow the CME (3) and instead satisfies the timepoint-wise algebraic relation

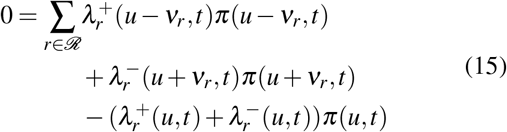

for all admissible states *u*, while ∂_*t*_*π*(*u, t*) ≠ 0 is still allowed. Hence, the true probability evolution of *p*(*u, t*) can only approximate *π*(*u, t*) timepoint-wise if the relaxation to the stationary distribution of the CRN is fast compared to changes in the protocol. Using this distribution, (13) can be derived through the method detailed in Supp. Sec. S2.2. Additionally, we review in Supp. Sec. S2.5 how the entropy production – which is introduced in the following section – decomposes into a so-called adiabatic and a non-adiabatic part.

The above relations for manipulated CRNs naturally extend to stochastic protocols, where the external control parameters are themselves governed by a stochastic process. In such cases, it is essential that the influence of the core system on the protocol, such as feedback on chemostat concentrations, remains negligible.

### C. Thermodynamics of random trajectories

A central appeal of stochastic thermodynamics is that quantities like heat dissipation, work, and entropy production can be defined along individual random trajectories. In the following, we review definitions of these quantities that are consistent with the principles of macroscopic thermodynamics (cf. Supp. Sec. S2.1). The presentation builds on Kurtz’s process-based formulation of CRNs, which naturally accounts for the instantaneous, finite changes at the mesoscopic scale caused by chemical reactions of U.

Let 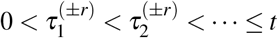 be the random times, when reaction ℜ_*r*_ happens in forward or backward direction.

Heat is dissipated during reaction events due to “excess” chemical work supplied by the chemostats. Under external manipulation, however, an additional contribution can arise even if the mesoscopic state does not change. The reason is that a time-dependent protocol can continuously modify the conditional equilibrium over the unresolved microscopic degrees of freedom associated with the currently occupied mesostate. This protocol-induced re-equilibration changes the internal entropy and, in general, is accompanied by heat exchange with the surrounding bath [66]. We therefore decompose the calorimetric heat along the random trajectory {*U*_[0,*t*]_} := *U* (*s*) _*s*∈ [0,*t*]_ into a reaction-induced part and a manipulation-induced part,

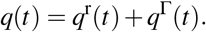

The reaction-induced contribution is

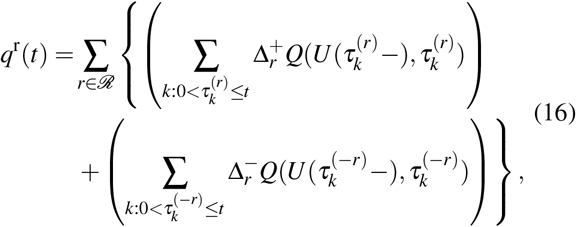

whereas the manipulation-induced contribution is defined by

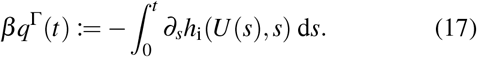

Here, *U* (*t*−) = lim_*s*↗*t*_ *U* (*s*) (or *U* (*t*^−^)) denotes the left limit in time, such that 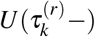 is the state of the CRN just before the *k*-th occurrence of reaction ℜ_*r*_ in forward direction [63]. The left limit in the second argument of 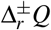 is irrelevant if Γ is a continuous function, but may be relevant for stochastic manipulation protocols. We restrict the discussion to deterministic protocols Γ that are piecewise-differentiable and hence drop the left limit in the second argument.

We rewrite (16) via stochastic integrals of the reaction counting processes:

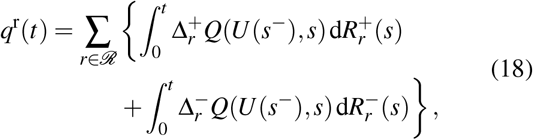

so that the total calorimetric heat becomes

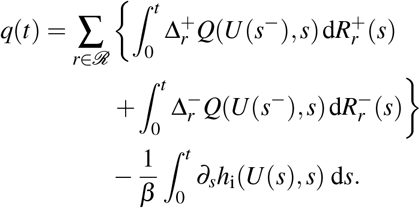

Defining the forward, backward, and net probability fluxes of reaction ℜ_*r*_ by

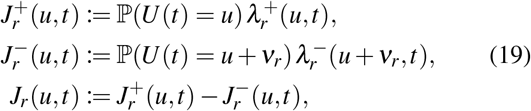

the mean heat dissipation rate, also called the thermal power [56], is

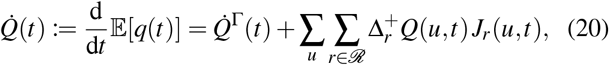

where

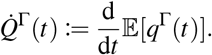

Equation (20) is rigorously derived in Supp. Sec. S2.7.

In a similar fashion, the chemical work performed on the trajectory *U*_[0,*t*]_ is

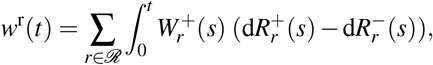

which gives rise to the mean rate of chemical work

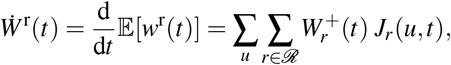

by following the steps in the proof of (20). The mean rate of chemical work 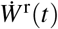 can also be understood as the amount of energy required to maintain the chemostat concentrations, i.e., the energy expenditure of the external agent. In a manipulated system, the change of the free energy of the system amounts to another work contribution along *U*_[0,*t*]_. For simplicity we assume that the Gibbs free energy is a continuous function of time [65], such that

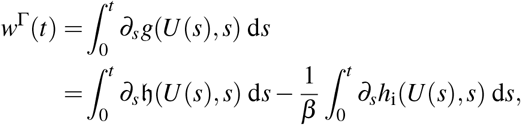

where the second line splits the manipulation work into an enthalpic and an internal entropic term. Hence, the mean rate of manipulated work is

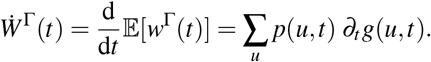

The entire work, associated with manipulation and driving, is then

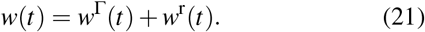

The enthalpy change along the trajectory can be decomposed into contributions from reaction events and continuous protocol-induced changes, i.e.,

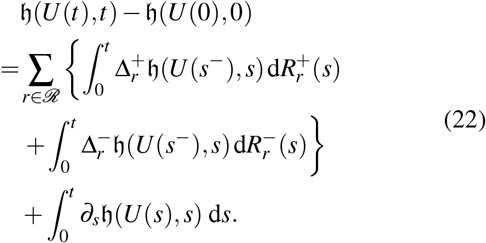

Likewise, the internal entropy change along the trajectory is

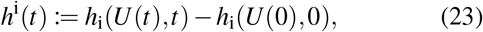

which decomposes into reaction-induced contributions and the protocol-induced contribution already identified in (17):

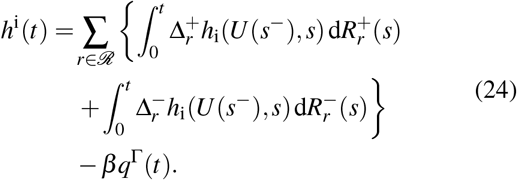

Plugging (14) into (18) and using (22) together with (21), we obtain the enthalpy balance relation

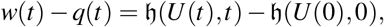

which is the trajectory-level first law of stochastic thermodynamics [64, Eq. (69)].

In the stochastic thermodynamics literature, one also encounters the alternative identity

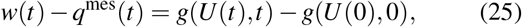

which uses the mesoscopic heat exchange

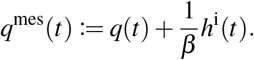

Using (24) together with the decomposition *q*(*t*) = *q*^r^(*t*) + *q*^Γ^(*t*), the mesoscopic heat exchange can be rewritten as

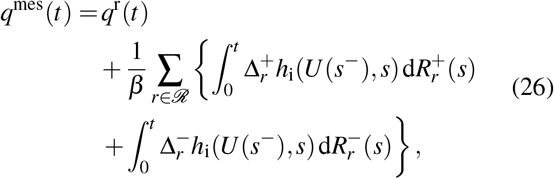

so that the protocol-induced contribution cancels explicitly. In other words, *q*^mes^(*t*) differs from the calorimetric heat *q*(*t*) by the internal-entropy contribution associated with the unresolved microscopic degrees of freedom. It is therefore the natural quantity in local-detailed-balance identities and trajectory-level fluctuation relations, i.e., (13) can be rewritten as

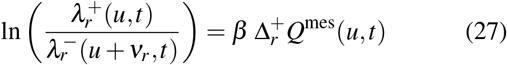

However, *q*^mes^(*t*) is in general not the actual heat released to the environment. It was first noted by Sekimoto [71] that the first-law-like equation (25) was not consistent with the macroscopic first law, since the designated “mesoscopic heat exchange” *q*^mes^(*t*) contains an additional internal entropic part such that it does not account for the actual heat flow, but only a change in the population entropy. Since then, *q*(*t*) is distinctly called measured heat [71] or calorimetric heat [65].

We now turn to a trajectory-level account of the total entropy production of the open system. The total entropy production along *U*_[0,*t*]_ comprises entropy changes of the system U ∪ S and the entropy exchange with the environment

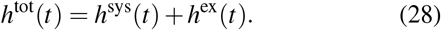

The system entropy itself contains two parts: (i) changes in the mesoscopic self-entropy

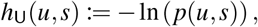

whose expectation is simply the population (Shannon) entropy of the core system *H*[*U* (*s*)] = E[*h*_U_(*U* (*s*), *s*)] = ∑_*u*_ − *p*(*U* (*s*), *s*) ln(*p*(*U* (*s*), *s*)) at time *s* ≥ 0, and (ii) changes in the internal entropy of the mesoscopic states *h*_i_. We denote the change of the population entropy along *U*_[0,*t*]_ as

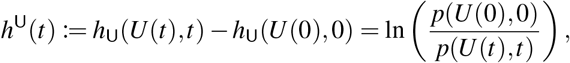

and, in turn, define the system entropy as

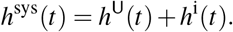

Not much can be said about the change of internal entropy, but the population entropy is a quantity, which is given in terms of the kinetic model of the open CRN. Following an analogous decomposition as (23) we obtain

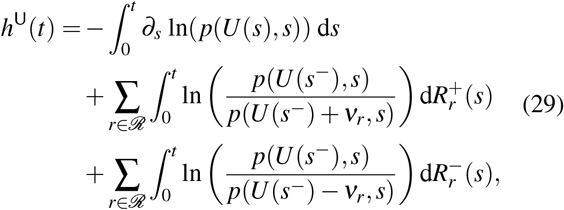

where by the use of the CME (3) we may substitute

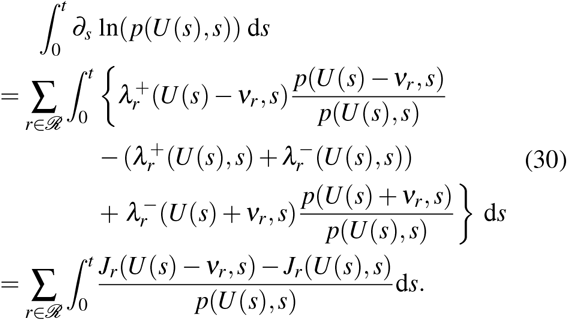

We further have 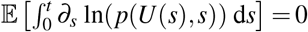, which follows_*s*_ by pulling the expectation into the integral (Fubini’s theorem) and applying a shift of summation. Using the CME (3) we can express the derivative of the mean population entropy as

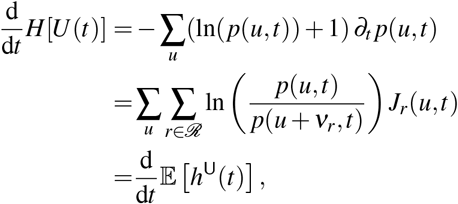

where the last equality follows from (29) and the proof method for Eq. (20) in Supp. Sec. S2.7.

According to macroscopic thermodynamic principles [68], the heat dissipated into the environment *q*(*t*) must be consistent with the increase of entropy in the environment *h*^ex^, such that

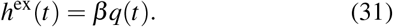

Equation (26) shows that *q*^mes^(*t*) retains only the reaction-resolved entropy flow, with the protocol-induced contribution already canceled. Applying the generalized consistency relation (13) to this representation therefore yields

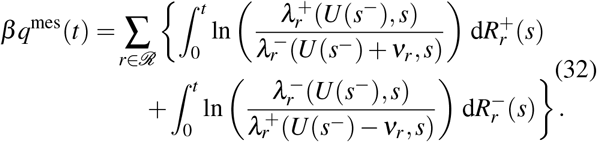

Hence *q*^mes^(*t*) is determined by the local propensity ratios and directly encodes the directional bias of each reaction channel ℜ_*r*_. In this sense, it is the natural heat-like quantity for connecting the stochastic jump dynamics to thermodynamic reversibility [72], as further discussed in Supp. Sec. S2.6; cf. Supp. Eq. (S2.17).

Again following the proof method for Eq. (20), the mean mesoscopic heat exchange rate (in units of *k*_B_*T*) satisfies

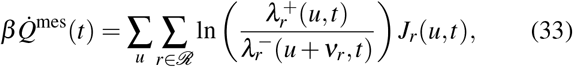

and thus depends only on the kinetic model of the open CRN.

In conclusion, the total entropy production satisfies

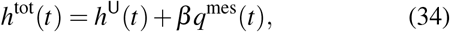

which is determined exclusively by the time-dependent population distribution and the time-dependent propensity functions. The *βh*^i^(*t*) terms, which contain encode internal entropy changes, cancel out between *h*^sys^(*t*) and *h*^ex^(*t*).

That is, regardless of internal energy changes due to external manipulation, the total entropy production of the open CRN depends solely on experimentally accessible quantities: the probability distribution over states and the reaction propensities. This key observation has been emphasized before [63, 66]. Using equations (29), (32), and (19), we obtain

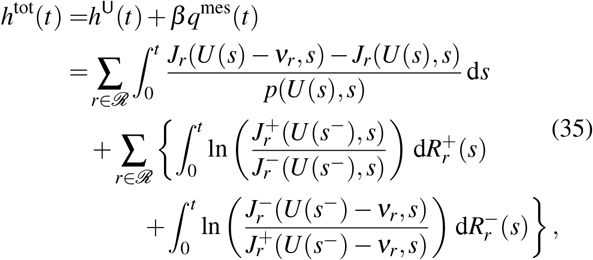

where we used (29), (32) and (19). The (total) entropy production rate (EPR) then follows with 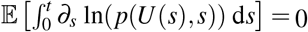 and the method of proof of (20):

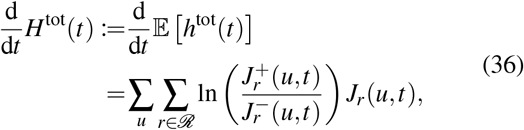

which can also be related to other mean rates

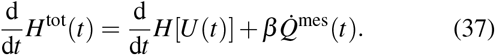

This decomposition motivated earlier inconsistent identifications of 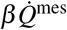 with the time-dependent entropy exchange rate [73, 74].

By virtue of ln(*x*)(*x* − 1) ≥ 0 for all *x* ≥ 0 (with convention ln(0) := − ∞), each term on the r.h.s. of (36) is non-negative and hence

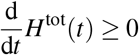

shows consistency with the second law of macroscopic thermodynamics.

### D. Energy dissipation and entropy production at stationarity

Now, we look at the entropy production rate at stationarity, which in the following will be identified with the mean energy dissipation rate in a non-equilibrium steady state (NESS).

At stationarity, we have lim_*t*→∞_ *H*[*U* (*t*)] = − ∑_*u*_ *π*(*u*) ln(*π*(*u*)), implying 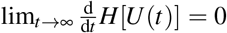. Consequently, with (37) the entropy production rate at equilibrium or in a NESS satisfies

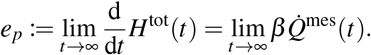

The EPR at NESS hence quantifies the entropy changes in the whole system, consisting of U ∪ S and the environment, with all entropic changes that are associated with the dynamics of the CRN and not just changes in the population distribution.

In the next step, we observe that the stationarity of *U* (*t*), combined with the local equilibrium condition Supp. Eq. (S2.5), implies that the process involving all microscopic degrees of freedom Ξ(*t*) is also stationary (cf. Supp. Sec. S2.1). Then the mean rate of change of the internal entropy (23) also vanishes:

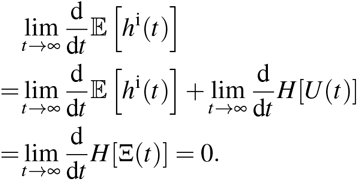

Applying (32) then establishes the identification of the EPR with the energy dissipation rate (in units of *k*_B_*T*) at stationarity:

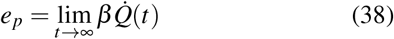

with mean heat exchange rate 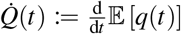. Eq. (38) plays an important role in the applications discussed in Sections VII and VIII D.

### E. Effective entropy production rate

In practice, the chemostat species of an open CRN are often omitted when formulating the stoichiometric balance equations and the corresponding propensity functions (1). As (9) shows, this simplification does not necessarily affect the definition of the stochastic dynamics of the core process *U*. However, in the context of stochastic thermodynamics, it is essential to keep track of all species involved in each reaction. In particular, using such “effective” reactions leads to a systematic underestimation of the entropy production rate.

In a properly defined open CRN it typically holds that 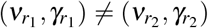 and 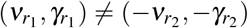 if *r*_1_ ≠ *r*_2_. However, the same changes in the core species U can be driven with different chemostat species.

#### Definition 2

*Two reaction channels* 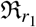 *and* 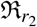 *are called* U*-*parallel, *if* 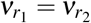 *and* 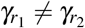, *or* 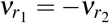 *and* 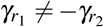.

Let *C* := {1, …, |*C*|} with |*C*| ≤ |ℛ| be the index set of the U-parallel reaction classes *C*_*α*_. Together, *C*_1_, …, *C* _|*C*|_ partitions ℛ. All reactions within such a class be directionally aligned such that 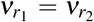 for all *r*_1_, *r*_2_ ∈ *C*_*α*_ with *r*_1_ ≠ *r*_2_. Then we define effective reaction counters 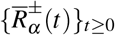 with effective propensity functions

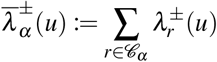

and effective probability fluxes

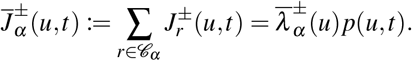

Applying (36) with these effective fluxes yields the “effective EPR” of the model with effective propensities:

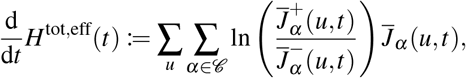

The effective EPR is a lower bound on the true EPR, i.e.,

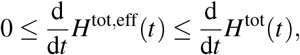

as has been shown in [65, Eq. (3.81)], using a multi-reservoir formulation of stochastic thermodynamics that we relate to the open CRN description Supp. Sec. S2.2. We exemplify this inequality in Sec. VIII D with a two-state Markov model exhibiting two reaction channels. While for this example 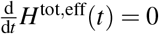 for any choice of rate constants,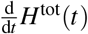 can be positive.

## IV. INFORMATION THEORY OF COMMUNICATION FOR GENERAL CAUSAL CHANNELS: PRELIMINARIES AND GENERALIZATIONS

For discrete-time systems with finite alphabets, the basic notions of information theory can often be developed without much measure-theoretic machinery. This changes in the presence of continuous-time signals or abstract, not necessarily finite alphabets, where a precise formulation requires more formal concepts from probability theory. In particular, measurable spaces, stochastic kernels, and probability measures on path spaces become indispensable.

It also brings in less familiar information measures that extend the classical notion of capacity and allow coding theorems to be formulated for arbitrary channels. Although the resulting capacity expressions are often difficult to evaluate, they offer an important conceptual advantage: memory, general time structure, and noisy feedback can be treated within a common framework, without requiring a separate and highly technical analysis of each channel class. In this way, the fundamental modeling structure of the communication system remains in the foreground.

The purpose of this section is therefore twofold. First, it collects the notation and concepts needed in the sequel. Second, it adapts and extends standard discrete-time notions to the more general time structure and to the specific modeling requirements of biochemical communication systems.

### A. General information measures: Preliminaries

Information measures, such as entropy and mutual information, were first introduced for finite spaces. General definitions have later been introduced and discussed by Kolmogorov and his colleagues Gel’fand, Yaglom, Dobrushin and Pinsker [75, Ch.7]. Most well-known information measures, such as entropy and mutual information, are instantiations of relative entropy or (KL-) divergence. For an abstract measurable space (Ω, ℱ) and any two probability measures ℙ: ℱ → [0, 1] and ℚ: ℱ → [0, 1], the relative entropy of ℙ with respect to ℚ is defined as

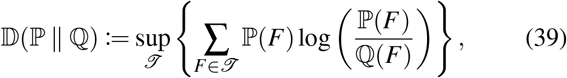

where the supremum is taken over all finite measurable partitions *T* ⊆ ℱ of Ω. For intuition, note that any such finite partition induces probability mass functions *F* ↦; ℙ(*F*) and *F* ↦; ℚ(*F*) on the finite set *T* such that the expression in the braces of (39) is simply a relative entropy on a finite alphabet.

Gel’fand, Yaglom, and Perez [75, 76] related the evaluation of the relative entropy (39) to the logarithm of a Radon–Nikodym derivative. This derivative describes a transformation law between two probability measures under the regularity requirement of absolute continuity [55]. That is, if a probability measure ℚ dominates ℙ in the sense that for any set *A* ∈ ℱ with ℚ(*A*) = 0 it also holds that ℙ(*A*) = 0, then ℙ is called absolutely continuous with respect to ℚ (denoted as ℙ ≪ ℚ). Then exists a density function *f* : Ω → [0, ∞) such that

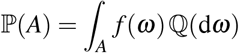

for all *A* ∈ ℱ. On a discrete space, absolute continuity simply means that supp(ℙ) ⊆ supp(ℚ). The density function *f* is called the Radon-Nikodym derivative and is often denoted as

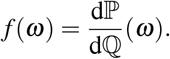

The Gel’fand-Yaglom-Perez Theorem states that the relative entropy is equivalently expressed as

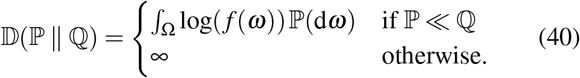

Hence, whenever ℙ ≪ ℚ, we refer to the function

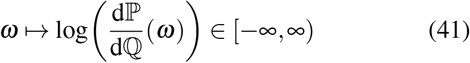

as the *divergence density* of ℙ relative to ℚ. These general divergence expressions and density notions are fundamental to this paper. Quantities of particular importance are the mutual information density it and the mutual information I.

Let *X* : Ω → *X* and *Y* : Ω → *Y* be random variables on the probability space (Ω, ℱ, ℙ) with values in abstract measurable spaces (*X*, Σ_*X*_) and (*Y*, Σ_*Y*_). The joint and the product probability on (*X* × *Y*, Σ_*X*_ ⊗ Σ_*Y*_) are given by

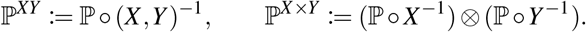

If ℙ^*XY*^ ≪ ℙ^*X* ×*Y*^, then the *information density* is defined as the divergence density

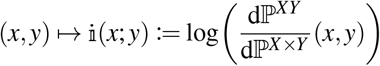

of ℙ^*XY*^ relative to ℙ^*X* ×*Y*^. In that case, the mutual information between *X* and *Y* is

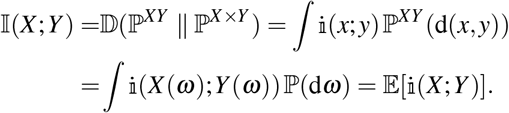

In conclusion, this subsection introduced relative entropy, information density, and mutual information in a manner that can be directly applied to path-space laws.

### B. Abstract communication channels, channel codes and constraints: Preliminaries

A central step in Shannon’s theory is the abstraction of noise corruption of a signal through a “transmission channel”, specified by a conditional distribution of the received signal given the transmitted (encoded) signal. The goal of channel coding is essentially to find good signal representations for the given noise statistics such that the transmitted message can be accurately reconstructed from the received signal. Following the measure-theoretic generalization of Shannon theory due to Kolmogorov and collaborators [76], as well as Kadota [77], Mittelbach and Jorswieck [78] and the book of Csiszár and Körner [79, Ch. 6], we formulate the noisy channel coding problem for general (not necessarily finite) alphabets. A message *M* on a finite index set ℳ ⊂ ℕ_*>*0_ is encoded into an input signal *X* on the abstract measurable space (*X*, Σ_*X*_), passed through the channel to produce an output *Y* on the abstract space (*Y*, Σ_*Y*_), and decoded into an estimate 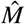 of the original message on the space ℳ ∪ {0}, where ‘0’ represents a decoding failure. These random variables are constructed to form the Markov chain

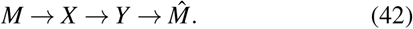

The transmission channel W is most generally defined as a stochastic/Markov kernel from *X* to *Y* [55, p. 204], i.e.,

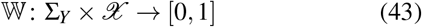

such that *B* ↦ W(*B, x*) is a probability measure for any *x* ∈ *X* and *x* ↦ W(*B, x*) is Σ_*X*_-measurable for any *B* ∈ Σ_*Y*_. For a channel W with input set *X* and output set *Y*, a channel code is a pair of mappings (f, *D*), where

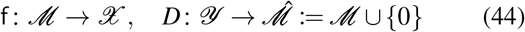

for a fixed message set ℳ. The finite image set f(ℳ) ⊆ *X* is called the codebook (or set of codewords). Although desirable, it is not always assumed that the encoding function f is injective (a one-to-one mapping), such that generally |f(ℳ) | ≤ |ℳ |. Further, assume that the message *M* is distributed according to the probability mass function *m* ↦ *p*_*M*_(*m*) such that *p*_*M*_(*m*) *>* 0 for all *m* ∈ ℳ. In the prospect of “coding into a random chemical system”, we also introduce the more general noisy channel code

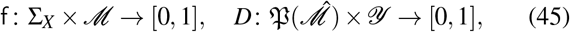

as stochastic kernels instead of the mappings in (44), where P denotes the operator generating the power set (i.e., the set of all subsets). In this case, the signal *x* ∈ *X* is randomly generated given a message *m* ∈ ℳ, and the message estimate at the decoder is also generated randomly given the received *y* ∈ *Y*. The random channel encoder f reduces to a deterministic code on a finite codebook *C* ⊆ *X* if it has the form

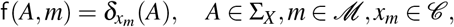

where *δ* denotes the Dirac-measure. However, the existence of such a codebook is not a requirement for general noisy channel codes. Similarly, in the case of a deterministic decoder, the noisy channel decoder kernel reduces to the form

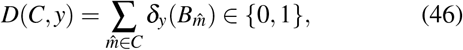

where 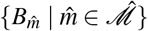 is a partition of *Y* in which empty sets are allowed and *B*_0_ := *Y \*∪_*m*∈ℳ_ *B*_*m*_.

Together, the channel W and the (noisy) channel code (f, *D*) induce yet another stochastic kernel 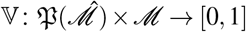, such that

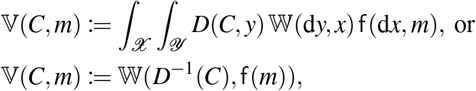

where the latter represents the case of a deterministic channel code (f, *D*). The kernel V determines the probability of any estimate 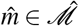, if actually *m* ∈ ℳ was transmitted over the channel W, using the code (f, *D*). As a “fidelity criterion” for the code (f, *D*), given the message distribution *p*_*M*_ and the channel W, we consider either the (mean) probability of error

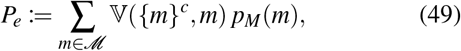

which reflects that common messages should be decodable more reliably than less common ones, or the traditionally preferred “worst-case” maximum probability of error

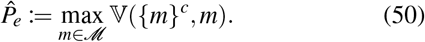

The channel coding problem, which we formalize in the next section, roughly describes the task of finding a good coding scheme for the fixed channel W, which allows to maximize the cardinality of ℳ while keeping *P*_*e*_ as low as possible.

The above construction defines encoded and received signals solely based on measurable signal spaces and the stochastic kernels defined between them. However, the defined stochastic kernels do also allow for a joint description on the observable joint space 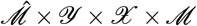, i.e.,

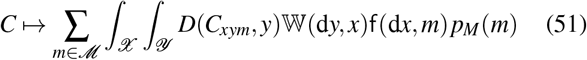

with 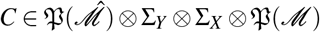 and

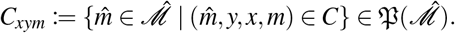

Until now, we have not used the basic probability space (Ω, ℱ, ℙ) and the random variables 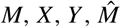 as functions on Ω. A useful way to interpret random variables in this context is that they link an underlying unobservable joint probability space, on which all randomness is defined at once, to the measurable signal and message spaces. We find the above construction of the channel and the code more instructive in the context of the channel coding problem; however, a basic probability space and random variable perspective can be consistently included, and we will use both perspectives in this paper.

#### Definition 3

(Connection via a channel). *Let X* : Ω → *X and Y* : Ω → *Y be random variables on the probability space* (Ω, ℱ, ℙ). *We say that X is connected to Y by the channel* W *if for all B* ∈ Σ_*Y*_

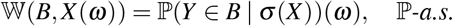

In other words, W is, by definition, a ℙ-regular conditional distribution of *Y* given *X* [54, pp. 106]. Only if the joint distribution 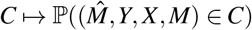 equals (51), then the random variables form the Markov chain (42) under ℙ and we will say that a subset of random variables is ℙ-consistent with the communication model.

Thus, while W specifies only the effects of the transmission medium on the signal, any basic probability measure ℙ must simultaneously govern *all* random elements of the communication system jointly and there is in general no unique a priori reference measure ℙ with respect to which W should be *defined* as a ℙ-regular conditional distribution. We introduce such a strong conceptual distinction here in prospective of coding theorem formulations, which in essence require optimization over certain parts of the communication system in two different ways, while keeping the channel law and potentially also its substructure fixed. In the context of chemical communication models such substructures can be intermediate species that forward a signal between other sets of species. This level of detail is not resolved in the classical channel formulation, but it becomes a relevant modeling choice in chemical communication systems, as discussed later in Sec. VII.

Lastly, signals and encoders used for information transmission are usually constrained by technical/system specifications and energy expenditure. There are three hierarchical levels of such constraints. First, the input signals can be constrained by a restriction of the signal space to some subset *A* ⊂ *X*. Then, the channel can be directly defined as a stochastic kernel from the measurable space (*A*, Σ_*A*_) (with some *σ*-algebra Σ_*A*_) to (*Y*, Σ_*Y*_) without requiring the kernel property w.r.t. the larger space (*X*, Σ_*X*_) [78]. Equivalently, the space of encoders and consistent basic probability measures, still defined on *X*, can be restricted to satisfy

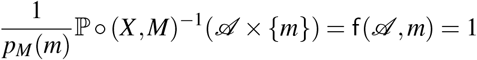

for all *m* ∈ ℳ, i.e., the support of the distributions is restricted to *A*. This is the strongest kind of constraint.

Often, such signal space constraints are formulated via a (measurable) cost functional Φ: *X* → [0, ∞] such that

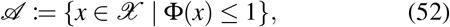

where the upper bound 1 is generic in the sense that we can always normalize Φ by any positive real number. This type of constraint has been used for channels with (continuous-) time structure, e.g., in [78, 80]. These works emphasize the technical requirement that the cost functions must be measurable, which is not always guaranteed.

Second in the hierarchy are message-wise expected cost constraints, which directly constrain the space of encoders or consistent basic probability measures such that

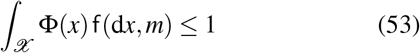

for all *m* ∈ ℳ. Given a cost functional Φ a signal space constraint Φ ≤ 1 implies (53), but generally not vice versa. Expected cost constraints just restrict the probability of high-cost signals, but they generally do not exclude them entirely. However, in the special case of deterministic codes the constraints (52) and (53) are equivalent.

Thirdly, there is the even weaker expected cost constraint, which additionally averages over the fixed message distribution, i.e.,

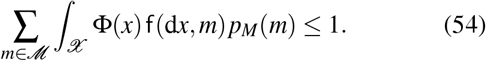

In this work, we will specifically discuss an energy constraint of the kind of (53), where the input cost functional is derived from a measurable input-output cost functional 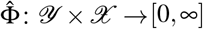 such that [81]

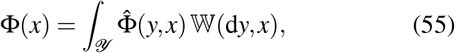

or more generally on the consistent basic probability space,

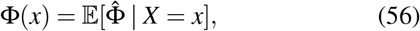

where 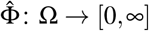 is a random variable that may include explicit “knowledge” of parts of the communication system that are not captured by *X* and *Y* alone. Such general communication system constraints encourage the use of the basic probability space perspective discussed above. In the next section we will introduce the role of time in information transmission and subsequently formalize channel coding theorems that specifies the temporally asymptotic property “channel capacity” for channels with time structure.

### C. Causal channels with time-structure: Preliminaries and Adaptations

In the previous section we have described the noisy channel coding problem as the task to make ℳ as large as possible while keeping *P*_*e*_ below a certain threshold or even arbitrarily close to zero. This problem is usually solved asymptotically for infinite transmission duration. Following Mittelbach and Jorswieck [78], we introduce a unified description of channels with time structure that applies equally to discrete-time as well as continuous-time signals.

*General time index and trajectory spaces*. Denote by T a general set of time indices, which may represent either the integers ℤ or the reals ℝ. Intervals are used in both cases, such that for *s, t* ∈ T with *s* ≤ *t*, the interval notation (*s, t*] is always understood as T ∩ (*s, t*] ⊂ T if *s < t*, and (*s, t*] := {*t*} if *s* = *t*. Let {(*X*_*t*_, Σ_*X,t*_) |*t* ∈ T} be a time-structured family of alphabets with (*X*_*t*_, Σ_*X,t*_) = (*X*, Σ_*X*_) for all *t* ∈ T. Here, we abuse notation slightly, since this alphabet is not necessarily identical to the measurable signal space introduced in the previous section. Typically, such a family is used to define the product space [55]

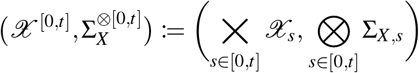

by taking the Cartesian product together with the product *σ*-algebra over all time-indices in the interval. This is the smallest *σ*-algebra that resolves differences between trajectories at countably many time points. In continuous time, where the product index set is uncountable, this *σ*-algebra is sometimes too small, and cost functions such as

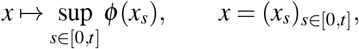

need not be measurable [78, 80]. Since in this work we assume a countable discrete state space *X* and stochastic processes with càdlàg trajectories, this issue does not arise for us: rather than working with the raw product measurable space, we always work on the càdlàg path space equipped with the Borel *σ*-algebra induced by the Skorokhod *J*_1_ topology, namely

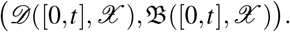

This also covers the discrete-time case, since any function on a discrete time index set is trivially càdlàg. For all *t* ≥ 0, and also for the interval [0, ∞), this is a Polish space and hence a standard Borel space, which entails a number of measure-theoretically useful structural properties [82]. For example, given a second discrete set *Y*, the spaces

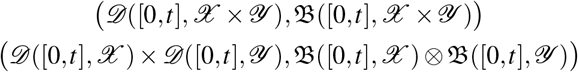

are canonically identical via the bimeasurable bijection that maps trajectories on *D* ([0, *t*], *X* × *Y*) to a pair of trajectories on *D* ([0, *t*], *X*) and *D* ([0, *t*], *Y*). As a result, any measure on the product space is uniquely identified with a measure on the joint space. We will therefore use these spaces, and also any finite product, interchangeably without further mention. In the information-theoretic literature, such spaces are sometimes referred to as standard alphabets; see, for example, [75].

#### Causal discrete time channels with initial state

To provide intuition let us first focus on the discrete-time setting with finite input and output alphabets, respectively, *X* and *Y*, and let *n* ∈ ℕ_*>*0_. The channel behavior for transmitting *n* symbols can be described by a conditional probability mass function W_*n*_ : *Y* ^*n*^ × *Y* × *X* ^*n*+1^ → [0, 1], which assigns probabilities to output sequences *y*_(0,*n*]_ = (*y*_1_, …, *y*_*n*_) given input sequences *x*_(0,*n*]_ = (*x*_1_, …, *x*_*n*_) and initial states (*y*_0_, *x*_0_). In particular, the mapping W_*n*_ can be represented as a matrix. In the case of a discrete-memoryless channel, this matrix is a product of the function W_1_ : *Y* × *X* → [0, 1] such that

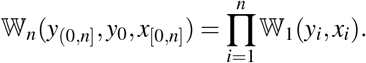

The discrete memoryless channel is independent of its initialization (*x*_0_, *y*_0_) and it is the standard case, extensively discussed in information-theoretic textbooks [79, 81, 83]. Any channel that adheres to the principle of causality satisfies a recursive product decomposition of the form

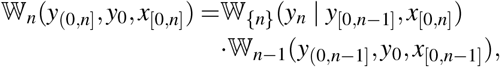

where W _{*n*}_ (*y*_*n*_ | *y*_[0,*n* −1]_, *x*_[0,*n*]_) represents the conditional probability mass function of *y*_*n*_ given the past input and output symbols *y*_[0,*n* −1]_, *x*_[0,*n* − 1]_ and the currently sent input *x*_*n*_. This implies,

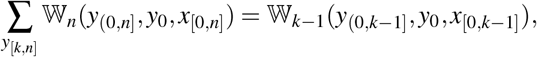

where the r.h.s. does not depend on *x*_[*k,n*]_ although we did not sum over it on the l.h.s. An equivalent channel characterization, that is often adopted in the literature, is the sequence of conditional probabilities 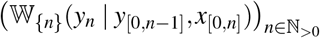. We will later observe that, for continuous-time CRNs, an analogous channel characterization is provided by intensity processes. Conceptually, each W_*n*_ is a causally conditional probability, as first introduced by Kramer [84]. After examining the role of time in indexed sequences, we will introduce a definition of causally conditional probabilities at the end of the following paragraph.

#### The arrow or time

Before we can define channels with general time structure, the role of time needs to be reevaluated. In discrete time (0, *n*], indicates the number of channel uses, i.e., the length of the encoded and the received signal sequence, respectively, *X*_(0,*n*]_ and *Y*_(0,*n*]_. Since *Y*_*i*_ is always received after *X*_*i*_ was sent for all *i* ∈ (0, *n*], the arrow of time may be represented as

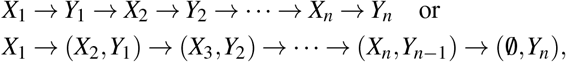

where in the first example, transmission and reception time-blocks alternate and in the second example, transmission and reception happens simultaneously, but the received signal is shifted by Δ*t* = 1 relative to the sent signal. The exact visualization of the arrow of time depends on the technical specifications of the system. It was noted by Newton [85] that this deviates from a natural time indexing, where *X*_*i*_ and *Y*_*i*_ are always simultaneous events, such that the arrow of time is visualized as

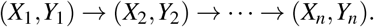

We emphasize that is arrow of time does not symbolize a Markov chain structure.

Using a symbol-wise index instead of a natural time index is certainly a useful abstraction in discrete-time communication, as it allows modeling to be decoupled from the technical system specifications that would otherwise be required. Continuous-time communication models without feedback often do not include an explicit delay either and allocate the generation and the detection of a continuous time signal in the same time block, e.g., (*k*Δ*t*, (*k* + 1)Δ*t*], *k* ∈ ℕ, Δ*t* ∈ ℝ_*>*0_ in the same natural time slot [78, 80]. In particular, a clarification of the arrow of time is always required when systems with feedback are considered, where encoding depends on the feedback information available at specific times. As the biochemical communication model developed in this paper is not restricted to consecutive block-encoding schemes, but rather views the entire transmission duration as a single block, we formulate the channel with general time structure with respect to natural time. In this case, the single-time channel characterization is replaced with

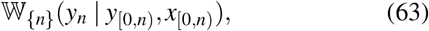

where (0, *n*] no longer represents the length of the encoded or received sequence, but the discretized transmission duration. The sequence 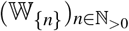 is actually equivalent to a discrete time stochastic processes with time-dependent transition matrices and history dependence on its own history and the history of the input signal. The causality requirement is then equivalently represented by

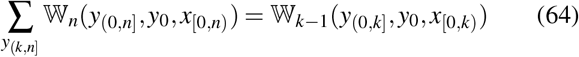

for all *k < n*. The probability law at the trajectory level for infinite output trajectories is represented by lim_*n* →∞_ W_*n*_, which encapsulates the same process information as the infinite sequence. This motivates the Def. 4 of causal channels for general time structure in the following paragraph. Before, we briefly introduce a definition of causal conditional probabilities in natural discrete time.

Assuming that both *X*_[1,*n*]_, and *Y*_[1,*n*]_ have countable discrete state spaces, then the conditional probability mass function of *Y*_[1,*n*]_ given *X*_[1,*n*]_ is defined as [86, p. 6]

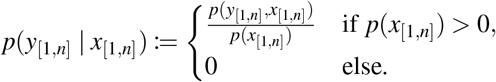

In contrast, the causally conditional probability mass function of *Y*_[1,*n*]_ given *X*_[1,*n*]_ in natural discrete time is defined as

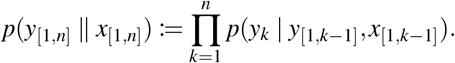

While a causal channel model, as motivated above, describes a modeling assumption, causal conditioning is just a different kind of conditional distribution for ordered sequences of random variables. While the normal conditional probability can be seen as a statistical conditional, the causally conditional probability explicitly ignores statistical dependencies between the present *y*_*k*_ and the future *x*_[*k,n*]_.

#### Causal channels with time structure

We now introduce a mathematical channel model for infinite transmission duration, for signals that are càdlàg trajectories with a general time index. This approach is formally advantageous as it encapsulates the model of the physical system for any transmission duration using a single stochastic kernel. Given that physical systems, particularly chemical reaction networks, operate causally, we directly integrate the constraint of causally functioning channels into our definition. Hence, for any countable set ℤ and any interval *I* ⊂ [0, ∞) let

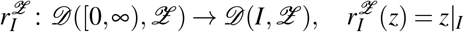

be the restriction of infinite signals to the interval *I*. Formally, this function is measurable with respect to the Borel*σ*-algebras, facilitating the following definition [78, 80].

##### Definition 4

(Causal channel with time structure). *A stochastic kernel*

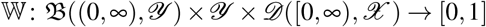

*that satisfies*

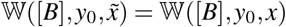

*for all t* ∈ T_*>*0_, *for all B* ∈ B((0, *t*], *Y*) *with* 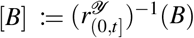 *and all* 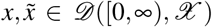 *with* 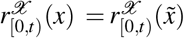 *is called a causal channel with time structure and initial state*.

Note that the causality condition in Def. 4 is equivalent to (64) in the discrete case. We increase generality compared to the Def. in [78, 80] by incorporating initial states (*y*_0_, *x*_0_) ∈ *Y* × *X* as arguments of the kernel. Hence, the channel may behave differently for distinct initializations. For a comprehensive overview of various classes of channels with state, the reader is directed to [81, Ch.7].

We emphasize once more that this channel definition is simply a way of characterizing a time-dependent, generally non-Markovian stochastic process whose evolution depends causally on the input signal. Accordingly, the operation of a physical channel over the interval (0, *t*] is described by a stochastic kernel from *Y* × *D* ([0, *t*), *X*) to *D* ((0, *t*], *Y*), namely

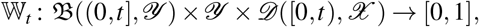

which satisfies

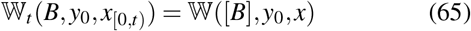

for all *t* ∈ T_*>*0_, all *B* ∈ B((0, *t*], *Y*), all *x*_[0,*t*)_ ∈ *D* ([0, *t*), *X*), and any representative 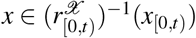. Equation (65) directly generalizes the discrete-time kernels W_*n*_. Abusing terminology, we will also refer to W_*t*_ as a causal channel.

#### Trajectory random variables

Given a basic probability space (Ω, ℱ, ℙ) with complete filtration ℱ = {ℱ_*t*_}_*t*≥0_ and trajectory-valued random variables

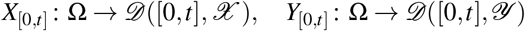

we denote the induced path-probability measures

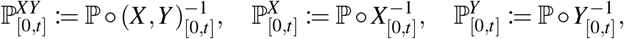

where 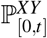 denotes the joint path measure, while 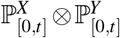 denotes the product of the marginal path measures. By ℱ^*XY*^ we denote the natural filtration of the process (*X,Y*): Ω → *D* ([0, ∞), *X* × *Y*), such that for all *t* ≥ 0 it holds

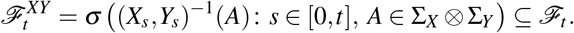

The sigma-algebra 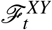 is just the projection of the trajectory-event space B([0, *t*], *X × Y*) to the basic probability space Ω.

#### Noisy initialization

As a new element we introduce a probability mass function

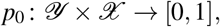

which models a potentially (joint) noisy initialization (*y*_0_, *x*_0_) of the encoded signal and the received signal. With respect to a basic probability measure ℙ the random variables (*Y*_0_, *X*_0_) and *M* must be statistically independent. The initialization distribution *p*_0_ depends on the technical specifications of the system and the specific degrees of freedom in code design. In wireless or molecular communication, *Y*_0_ could for instance arise from background noise, while *X*_0_ is deterministically set to the “null”-signal associated with no transmission. Another case are potential memory and/or noise effects of the joint encoder-channel system before the transmission is started, which is actually the case in chemical communication. The initialization *p*_0_, hence, has a peculiar status in the general formulation of the channel coding problem. Sometimes it is a fixed specifications together with W [81, Ch.7], and sometimes it can be part of the design degrees of freedom under certain constraints. For generality, we assume the latter case.

#### Channel codes with time structure

We now connect the causal channel with time structure to the abstract communication framework introduced in Sec. IV B. For a transmission of duration *t* ∈ T_*>*0_, we interpret W_*t*_ as the channel describing the conditional distribution of the output trajectory *y*_(0,*t*]_ ∈ *D* ((0, *t*], *Y*), given the initial condition together with the transmitted input trajectory, (*y*_0_, *x*_[0,*t*)_) ∈ *Y* × *D* ([0, *t*), *X*), in the sense of Eq. (43). A noisy channel code for signals with time structure and duration *t* is then defined in direct analogy with Eq. (45). At this point, we do not impose a causality constraint on the code itself, since the message set ℳ is not endowed with its own time structure.

Including the noisy initialization, the code and channel jointly induce the stochastic kernel

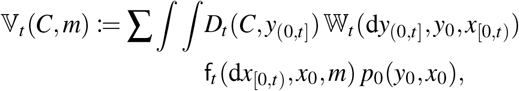

for 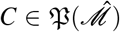 and *m* ∈ ℳ, where the sum and integrals extend over the full initialization and path spaces. Thus, V_*t*_(*C, m*) is the probability that the decoder output lies in *C*, conditioned on transmission of the message *m*, under the channel W_*t*_, the noisy code (f_*t*_, *D*_*t*_), and the initialization law *p*_0_. The definitions of error probabilities (49), (50) and the input constraints (52) (55) then carry over without change. Definition 3, together with the corresponding notion of ℙ-consistency on a basic probability space (Ω, ℱ, ℙ), also extends to this setting, provided that the message *M* is ℱ_0_measurable and statistically independent of the random initialization (*Y*_0_, *X*_0_). In this sense, the message is fixed at time zero and initiates the operation of the communication system, including the encoder f_*t*_. This formulation is sufficient for the present setting, but it will require refinement once feedback is introduced, as discussed in Sec. IV E.

### D. General channel capacity of causal channels with time structure: Preliminaries and Generalizations

#### One-shot codes and code rates

For any transmission duration *t* ∈ T_*>*0_, a channel code (f_*t*_, *D*_*t*_) on the interval (0, *t*], together with the causal channel W_*t*_, the initialization distribution *p*_0_, and the message distribution *p*_*M*_ on ℳ, determines a communication system over the time interval (0, *t*]. We refer to such a code (f_*t*_, *D*_*t*_) as a (|ℳ |, *t*)-code.

The standard code rate of a (|ℳ |, *t*)-code is denoted as

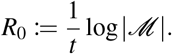

If the message *M* is uniformly distributed, then 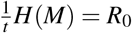. In classical channel coding, one often assumes a uniform message distribution, since for deterministic codes the coding problem can be reduced to that case by a suitable randomization argument [81, p. 73]. Moreover, bounds such as Feinstein’s lemma [77] are typically formulated independently of *p*_*M*_ when the fidelity criterion is the maximum probability of error. In the present biochemical setting, however, the semantic meaning of a message may be tied to its physical encoding, so that this reduction to a uniform prior need not be appropriate. We therefore keep the message distribution explicit and define the mean code rate under *p*_*M*_ by [87]

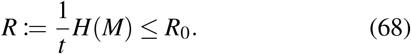

Later we will also use the equivalent notation *H*(*p*_*M*_) in order to make the dependence on the message distribution explicit.

#### Limitations of classical block concatenation coding for channels with significant memory

In classical channel coding, longer block codes are often constructed by concatenating shorter codewords, or “letters,” in time [37, 78, 80, 88]. Given a (|ℳ_*a*_|, *t*_*a*_)-code and a (| ℳ_*b*_ |, *t*_*b*_)-code, one transmits a message pair (*m*_*a*_, *m*_*b*_) ∈ ℳ_*a*_ × ℳ_*b*_ by first sending the codeword associated with *m*_*a*_ and then, after resetting the encoder, the codeword associated with *m*_*b*_.

Under additional assumptions, this temporal concatenation satisfies a discrete block-memoryless property, which is a key ingredient in the coding theorem for channels with general time structure derived in [78]. This encoder-side memoryless property does not, however, eliminate memory effects at the channel output. If the channel itself is not block-memoryless, then consecutive blocks may still interfere through the channel memory and thus exhibit intersymbol interference across blocks [78]. Enforcing approximately block-memoryless behavior at the receiver may therefore lead to very low practical code rates for channels with memory, because the guard or relaxation intervals required to return the system close to a prescribed initialization *p*_0_ can be substantial and may dominate the overall transmission time [88]. We assume that the scenario of long relaxation times is common in biochemical communication systems, leading us to move away from the concept of consecutive block coding.

#### Monotone families of message sets

For channels with significantly long memory it is natural to treat the entire transmission interval as a single coding block and to use one larger codebook over that interval rather than a sequence of shorter blocks decoded separately. This leads to families of message sets and code families indexed by the transmission duration itself. To exclude degenerate parametrizations, we impose the following regularity condition.

##### Definition 5

(Monotonicity of messages and codes). *A collection* {ℳ_*t*_ | *t* ∈ T_*>*0_} *of message sets (and message variables M*_*t*_ : Ω → ℳ_*t*_*), together with a collection of probability mass functions* {*p*_*M,t*_ : ℳ_*t*_ → [0, 1] | *t* ∈ T_*>*0_} *is called* monotone *if*

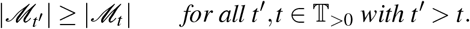

*Any associated collection of codes* 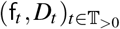 *for* 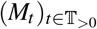 *is also called* monotone.

This definition does not imply that the code rate 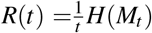 is monotone in *t*. Rather, it expresses a minimal consistency requirement: increasing the available transmission time should not force a reduction of the message alphabet. In this sense, monotonicity is compatible with the usual temporal concatenation idea, but formulated here at the level of a single channel use with varying duration.

#### Example and scope of the monotonicity requirement

As an example, suppose one concatenates identical (|ℳ_0_|, Δ*t*)slots with |ℳ_0_| ≥ 2. Then

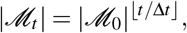

and the corresponding standard code rate is

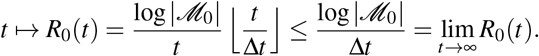

Thus, standard block concatenation produces a monotone family in the above sense.

By contrast, continuous-time signal models do not in general impose a canonical minimal slot length. Without additional regularity or cost constraints, one may therefore obtain artificially large per-unit-time rates by allowing increasingly rich message sets on arbitrarily short transmission intervals. The monotonicity requirement is a first structural restriction that rules out families in which the message alphabet decreases as the transmission time increases. At the same time, it is deliberately weak: by itself, it does not exclude pathologically rapid, exponentially increasing rate behavior.

#### Reliable finite-duration codes and admissible classes

Having introduced the basic structural restriction on message families, we now turn to the notions of reliability and admissibility needed for asymptotic coding statements.

##### Definition 6

*Let ε* ∈ [0, 1) *and t* ∈ T_*>*0_. *A code* (f_*t*_, *D*_*t*_) *with initialization p*_0_ *is called an* (|ℳ|, *t, ε*)*-code for the channel* W_*t*_ *and message distribution p*_*M*_ *if*

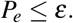

In the following, let

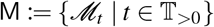

denote a monotone collection of message sets, together with a family of probability mass functions

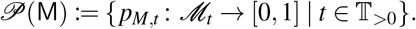

We allow the class of admissible pairs (M, *P* (M)) to be restricted to a prescribed subclass *C*_M_, which we leave unspecified. Further, let *P*_0_ denote the class of admissible initialization distributions *p*_0_. Any element of *C*_M_ or *P*_0_ is called *admissible*. By a slight abuse of terminology, we will also refer to a pair (W, *C*_M_) or a triple (W, *P*_0_, *C*_M_) as a channel.

#### Achievable rates and channel capacity

We now introduce the asymptotic notions of achievability and capacity required for coding theorems. Informally, the question is the following: as the utilized transmission time becomes large, how much information per unit time can be conveyed reliably through the channel? In the present setting, this question is slightly broader than in the classical formulation, because not only the code itself but also the admissible message families and the initialization distribution may be part of the design space. The next definition formalizes the requirement that, for all sufficiently large transmission durations, one can find an admissible communication scheme whose mean code rate is arbitrarily close to a prescribed value *R*^∗^ while the decoding error remains below a prescribed threshold. The formulation is generalized from Csiszár and Körner [79].

##### Definition 7

(*ε*-achievable rates and *ε*-capacity). *Let ε* ∈ [0, 1), *let* W *be a causal channel with time structure, let P*_0_ *be a class of admissible initialization distributions, and let C*_M_ *be a class of admissible monotone message collections*.

*A non-negative number R*^∗^ *is called an ε*-achievable rate *for* (W, ℙ_0_, *C*_M_) *if, for every δ >* 0, *there exists t*_0_ ∈ T_*>*0_ *such that for all t* ∈ T_*>*0_ *with t* ≥ *t*_0_, *there exists an admissible pair* (M, *P*(M)) ∈ *C*_M_ *and an admissible initialization p*_0_ ∈ *P*_0_ *such that*

i. 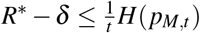 *and*
ii. *there exists an* (|ℳ_*t*_|, *t, ε*)*-code for* (W_*t*_, *p*_*M,t*_).

*A number R*^∗^ *is called an* achievable rate *if it is ε-achievable for every ε* ∈ (0, 1).

*The supremum of all ε-achievable rates is called the ε*capacity *and is denoted by C*_*ε*_. *The supremum of all achievable rates is called the* capacity *and is denoted by C*.

In words, an *ε*-achievable rate refers to a rate that can be closely approximated for all sufficiently large transmission durations through an appropriate selection of message family, initialization, and code, ensuring the error probability does not exceed *ε*. Consequently, the *ε*-capacity is therefore the best asymptotic rate if a fixed nonzero error level *ε* is tolerated. The capacity *C* is the corresponding zero-error-threshold notion in the usual Shannon sense, requiring that the same asymptotic rate be attainable regardless of how small the target error probability is set.

Since *C*_*ε*_ is non-decreasing in *ε* ∈ (0, 1), it follows that

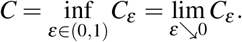

#### Message distributions as part of the design space

In contrast to the classical formulation, we regard the message sets together with their associated distributions as part of the admissible design space. This generalization is relevant for biochemical communication, where semantic restrictions may limit which message distributions are meaningful or where ℳ itself may represent a quantized signal space of an underlying physical phenomenon. As a consequence, even uniform message distributions need not be admissible elements of *C*_M_. We hypothesize that the resulting admissible class can be important for the information-theoretic characterization of a given biochemical system, but we do not assess this hypothesis further in the present work. Importantly, while capacity is usually formulated via the standard code rate *R*_0_ and the maximum probability of error 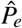, related prior-aware perspectives do appear in the literature [89, 90]. However, these results do not cover the present setting, where admissible message distributions themselves are part of the design space. Developing a corresponding coding theorem is beyond the scope of this work.

#### Equiprobable block messages as a classical special case

For comparison with other works, we may consider the family obtained by concatenating identical (|ℳ_0_|, Δ*t*)-slots with |ℳ_0_| ≥ 2 and Δ*t* ∈ T_*>*0_:

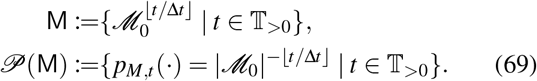

Thus, for every *t* ∈ T_*>*0_ all available messages are equiprobable. Previous works such as [78, 80] have implicitly considered the class *C*_M_ generated by such message families for all |ℳ_0_| ∈ ℕ_*>*1_ and Δ*t* ∈ T_*>*0_, and referred to Δ*t* as the block transmission duration.

#### Finite-duration information measures induced by an input law

The capacity definition in Def. 7 is operational. To formulate coding theorems, we now connect it to the general information measures introduced in Sec. IV A, which provide computable upper and lower bounds and, under suitable conditions, exact characterizations of that operational quantity. For the moment, let the channel and the encoded signal be independent from the initialization state (*y*_0_, *x*_0_) ∈ *Y* × *X*. Then the channel over the interval (0, *t*] reduces to a stochastic kernel of the form

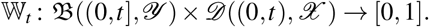

Let *µ*^*X*^ be an arbitrary probability measure on *D* ((0, ∞), *X*), i.e.,

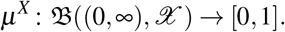

For each *t* ∈ T_*>*0_, define its restriction to (0, *t*) by

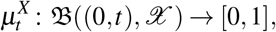

where

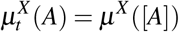

for all *A* ∈ B((0, *t*), *X*), with 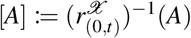. The class of admissible probability measures *µ*^*X*^ is assumed to satisfy any constraint imposed on the admissible encoder family f_*t*_ for all *t* ∈ T_*>*0_.

From 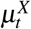 and W_*t*_ we define three further probability measures:

i. the joint measure 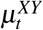 on B((0, *t*), *X*)⊗B((0, *t*], *Y*) by

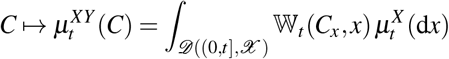

with *Cx* := {*y* | (*x,y*) ∈ *C*} for all *x* ∈ *D*((0,t),*X*),
ii. the output measure 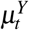 on B((0, *t*], *Y*) by

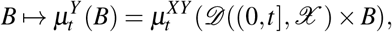
iii. and the product measure 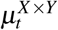 on B((0, *t*), *X*) ⊗ B((0, *t*], *Y*) determined by

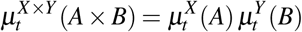

for all *A* ∈ B((0, *t*), *X*) and *B* ∈ B((0, *t*], *Y*).

This motivates the definition of the *mutual information density* [91, 92] for the pair (*µ*^*X*^, W) restricted to (0, *t*]. If 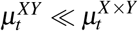, define

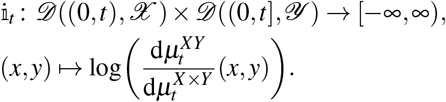

In that case, the associated mutual information is

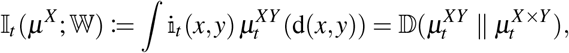

where the last equality follows from (40).

Equivalent formulations in terms of the Radon-Nikodym derivative 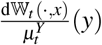 are often used in the literature [91, 93]. We employ the construction via the joint and product measures because it fits directly into the general divergence framework introduced in Sec. IV A and is technically more direct on abstract spaces [94].

#### Information rates

To pass from finite-duration information measures to asymptotic information rates, we use the liminf and also the liminf in probability of a sequence of real-valued random variables (*A*_*n*_)_*n*∈ℕ_ on (Ω, ℱ, ℙ), defined by [91]

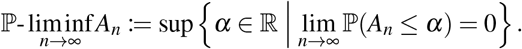

For a fixed channel W, there are two equivalent ways to construct the joint path measure *µ*^*XY*^. One may start from an arbitrary input path measure *µ*^*X*^ and combine it with W to obtain *µ*^*XY*^. Alternatively, one may start from a basic probability measure ℙ together with random variables (*X,Y*) such that W connects *X* to *Y* under ℙ; in that case, the induced input law may specify *µ*^*X*^ = ℙ^*X*^, such that the resulting joint law satisfies *µ*^*XY*^ = ℙ^*XY*^. Hence, the information density and mutual information associated with (*µ*^*X*^, W) agree with those associated with (*X,Y*) under any ℙ-consistent realization.

For chemical communication, however, the formulation in terms of a basic probability measure ℙ is often more informative. It allows one to impose additional structural consistency requirements on the admissible realizations beyond the sole requirement that *X* and *Y* be connected by the abstract channel law. In the case of communication through a CRN process *U*, such additional structure may include, for example, the presence of specific reactions, prescribed propensity functions, whole-network energy constraints, or consistency with intermediate chemical species that are not resolved at the level of the abstract channel from *X* to *Y*. Once the abstract channel description is refined in this way, the corresponding code-design problem and the capacity problem can also be formulated more specifically.

By construction, *µ*^*X*^ is a probability measure on the input trajectory space, not a stochastic kernel on that space. In particular, it should not be confused with a noisy encoder, which is a kernel from the message space to the input signal space.

##### Definition 8

(Information rates). *Let X* : Ω → *D* ((0, ∞), *X*) *and Y* : Ω → *D* ((0, ∞), *Y*) *be random variables on the probability space* (Ω, ℱ, ℙ). *Suppose that X is connected to Y by the causal channel with time structure* W, *and let* i_*t*_ *be the information density of the pair* (ℙ^*X*^, W) *restricted to D* ((0, *t*), *X*) × *D* ((0, *t*], *Y*).

*Then*

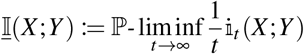

*is called the* inf-information rate *between X and Y*. *Further*,

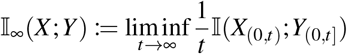

*is called the* mutual information rate *between X and Y*.

For discrete time it is known that [92], [93, Thm. 8]

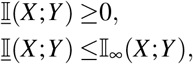

but the extension to continuous-time is straightforward.

#### General coding theorems and information stability

For a discrete-time channel W, deterministic channel codes, and the class of equiprobable message families from (69), Verdú and Han [93] proved the general coding theorem for the capacity

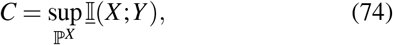

where the supremum is taken over all basic probability measures ℙ such that *X* is connected to *Y* by the channel W. This theorem applies to arbitrary discrete-time channels without further structural assumptions.

Under an additional *information stability* condition, originally attributed to [26, 95], the general formula reduces to the more familiar Shannon-type expression

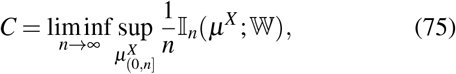

provided there exists a probability measure ℙ (equivalently *µ*^*X*^) such that

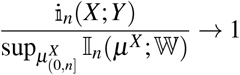

in probability as *n* →∞ [89]. This resembles a weak law of large numbers for the information density. In practice, however, information stability is often difficult to verify, so many works instead prove (75) directly for specific channel classes [78, 93]. A common sufficient condition is that the joint process (*X,Y*) is stationary ergodic together with suitable restrictions on the memory of W and/or *X*.

#### Constraints and noisy initialization

For noiseless channel codes, input restrictions can be imposed directly through admissible signal sets, as in the hard constraint (52). The general coding theorem (74) remains valid under such constraints [92, Thm. 3.6.1]. In that case, the supremum over *µ*^*X*^ is taken only over those distributions that satisfy 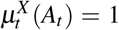 for all *t* ∈ T_*>*0_ on prescribed constraint sets *A*_*t*_ ⊂ *D* ((0, *t*), *X*).

A fixed initialization can be incorporated into this framework as an additional hard constraint. More precisely, if the encoder is initialized at a prescribed state *x*_0_ ∈ *X*, then one may instead work on the path space *D* ([0, *t*), *X*) and restrict attention to admissible sets *A*_*t*_(*x*_0_) consisting only of càdlàg trajectories with initial value *x*_0_. Thus, all admissible encoding trajectories start from the same prescribed state. Analogously, a fixed receiver initialization *y*_0_ ∈ *Y* may be incorporated into the channel law by restricting the output to trajectories in *D* ((0, *t*], *Y*) that are generated from that prescribed initial condition.

A genuinely noisy initialization is different. If the initialization law *p*_0_ is itself part of the admissible design space, then the input object is no longer just a probability measure on the future input trajectories, but more naturally a stochastic kernel

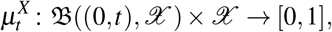

which specifies the conditional law of the future input trajectory *x*_(0,*t*)_ given the initial state *x*_0_. In that case, the joint law is constructed by averaging over the initialization distribution such that

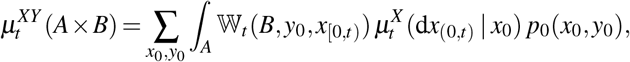

for all *A* ∈ B((0, *t*), *X*) and *B* ∈ B((0, *t*], *Y*), where *x*_[0,*t*)_ denotes the trajectory obtained by adjoining the initial value *x*_0_ to *x*_(0,*t*)_.

Accordingly, if *p*_0_ is allowed to vary within an admissible class, then the associated information-rate optimization becomes an iterated optimization: first over admissible input kernels *µ*^*X*^ and then over admissible initialization laws *p*_0_.

#### Interpretation of coding theorems

In general, the mathematical significance of channel coding theorems lies in the fact that they render channel capacity numerically accessible through information measures [79]. Channel coding theorems are usually (non-constructive) existence results, just as the capacity in Def. 7 is defined as an “existence” property. The iterative construction of code families with low *P*_*e*_ is typically considered intractable, in contrast to the optimization of information measures over *µ*^*X*^, at least for certain classes of channels. Note that while the capacity formalizes a supremum over channel codes, the r.h.s. of (74) and (75) takes a supremum over the probability measures *µ*^*X*^, where the two optimization sets are related only by the constraints [96]. The practical significance is that they identify the theoretically achievable optimum for a given channel and thereby provide a benchmark against which actual codes can be evaluated [79]. The proofs for coding theorems always consist of two parts, a *direct part* stating the existence of a “good” (low error) code whose code rate is upper bounded by the supremum of the information measure, and a *converse part* stating the nonexistence of “good” codes with higher rate.

#### Context for biochemical communication

In the context of chemical communication, we note that coding theorems are usually formulated for deterministic channel codes and under equiprobable messages. Chemical populations, however, typically exhibit intrinsic fluctuations, so that signals cannot in general be encoded directly into population trajectories by a deterministic map. This motivates the two-step encoding construction via chemostats in Sec. VII A. The separation between optimization over code families and optimization over input laws will also help place the case studies in Sec. VIII D into a coherent information-theoretic framework. In the following sections, the concepts of channels with time structure and channel capacity are extended to the case where noisy encoders may also use noisy feedback.

### E. Causal channels with time structure and noisy causal feedback: Preliminaries and Generalizations

#### Historic context

It was first noted by Marko [97] that (human) communication, unlike mere information transmission, is intrinsically bidirectional. Building on his ideas, Massey initiated a consistent extension of the channel coding problem to incorporate the use of feedback [11]. Importantly, he emphasized the distinction between statistical and causal dependence in the formulation of proper information measures. Feedback is not an inherent feature of the channel itself; rather, it is integrated into the encoding of the signals transmitted through the channel. In particular, Massey introduced an information measure called directed information and demonstrated that, when used in place of mutual information, it might sometimes accurately extend channel coding theorems for causal communication systems in discrete time to transmission with the use of feedback. For channels that are used without feedback, directed information from *X* to *Y* reduces to the mutual information between them. Figure 4 shows such a causal discrete time communication system. According to Massey, the defining property of a causal communication system is that the causal channel W _{*n*}_ at every transmission instance is conditionally independent of the sent message *m*, given *y*_[0,*n*)_, *x*_[0,*n*)_. This reflects the understanding that the encoding process happens before transmission, with the encoding device sending only the signal into the channel.

**FIG. 4.**
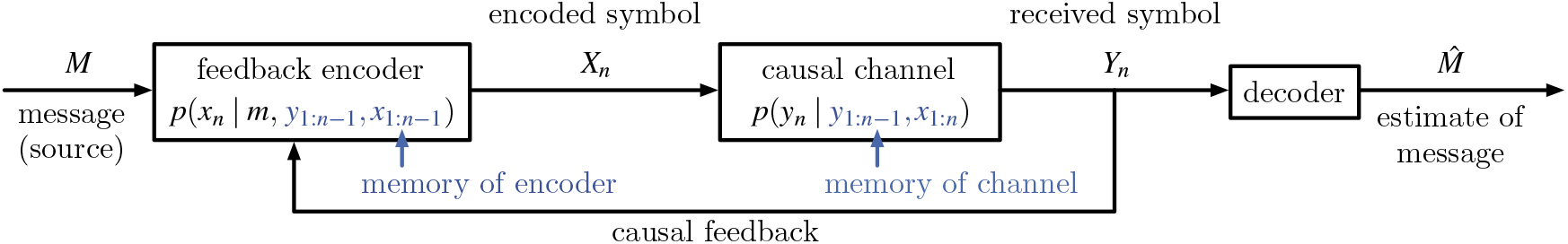
General discrete-time causal communication system. In a general discrete-time causal communication system [11], a random message *M* with finite state space is encoded in a discrete sequence {*X*_*n*_}_*n*∈{1,…,*N*}_, such that each symbol *X*_*n*_ is generated by the conditional probability function *p*(*x*_*n*_ | *m, y*_1:*n*−1_, *x*_1:*n*−1_) based on its own history *X*_1:*n*−1_, the history of received symbols *Y*_1:*n*−1_ (causal feedback), and the message *M*. The received sequence {*Y*_*n*_}_*n*∈{1,…,*N*}_ is transmitted through a channel, such that each *Y*_*n*_ is generated by the probability *p*(*y*_*n*_ | *y*_1: −*n* 1_, *x*_1:*n*_), independently of the message. The receiver then implements a decoding function to estimate *M* based on the observation of the entire trajectory *Y*_1:*N*_. In particular, the running index of the process stands for subsequently transmitted and received symbols and does not represent natural time.

We believe differentiating between constructing the communication model via kernels and modeling it on a basic probability space adds clarity. Therefore, we will first outline the kernel-based approach and then relate it to a probability measure ℙ.

#### Causal noisy feedback encoders with time structure

In essence, the construction of a causal noisy feedback encoder with time structure is analogous to that of the causal channel with time structure in Sec. IV C. In discrete time, a sequence

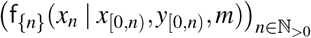

is equivalent to a discrete time stochastic processes with time-dependent transition matrices and history dependence on its own history and the history of the received signal. However, for a noisy encoder this does not mean that the history *y*_[0,*n*)_ is fed directly into the “encoding device” without prior noise corruption. This stochastic process model does not provide an explicit specification of this level of detail. The causality requirement is equivalently expressed at the kernel level for finite encoder operation times as

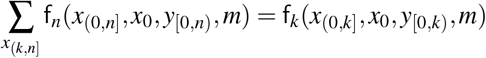

for all *k < n*. In discrete time, f_*n*_ is exactly analogue to a causally conditional probability, as first defined by Kramer [84]. The definition of a noisy encoder as a kernel describing the statistics of infinitely long input-signal trajectories for general time structure is as follows.

##### Definition 9

(Causal noisy encoder with time structure). *A stochastic kernel*

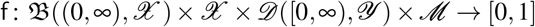

*that satisfies*

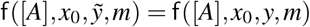

*for all m* ∈ ℳ, *for all t* ∈ T_*>*0_, *for all A* ∈ B((0, *t*], *X*) *with* 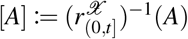 *and all* 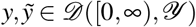 *with* 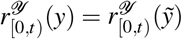 *is called a causal noisy encoder with time structure and initial state*.

Accordingly, operating the encoder over the interval (0, *t*] is described by a stochastic kernel from *X* × *D* ([0, *t*), *Y*) × ℳ to *D* ((0, *t*], *X*), namely

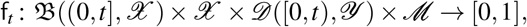

which satisfies

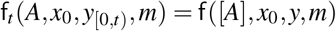

for all *t* ∈ T_*>*0_, all *A* ∈ B((0, *t*], *X*), all *y*_[0,*t*)]_ ∈ *D* ([0, *t*), *Y*) and any representative 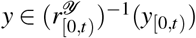.

#### Joint law of encoder and channel

Thus far, we have defined a causal channel and a causal encoder as stochastic kernels that satisfy a causality condition. Once feedback is present, however, the input and output trajectories no longer form a Markov chain of the form (42). Hence, unlike in Eq. (51), the joint law cannot be obtained by a single integration of the channel against the encoder. For intuition, we briefly return to symbol indexing and consider the conditional factorization of the joint law of two discrete sequences [11, Eq. (3)]:

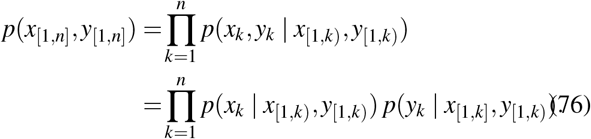

For discrete sequences with symbol index, where the arrow of time is *x*_*k*_ → *y*_*k*_ → *x*_*k*+1_ rather than (*x*_*k*_, *y*_*k*_) → (*x*_*k*+1_, *y*_*k*+1_), Eq. (76) is exactly the causality-preserving rule for constructing the joint law from an encoder kernel and a channel kernel. In natural time, by contrast, such an ordering within a single time step is no longer built into the indexing itself. Therefore, once encoder and channel are modeled as separate causal devices, an additional modeling choice is required to specify how they are coupled at equal times. In the present framework, we adopt the convention that the coupling acts only through the accumulated past. Conditionally on the history up to time *k* − 1, the encoder generates the new input update and the channel generates the new output update, but neither update depends on the other’s simultaneously generated new value.

Adapting the above factorization to natural discrete time and to our original encoder and channel kernels, we therefore define

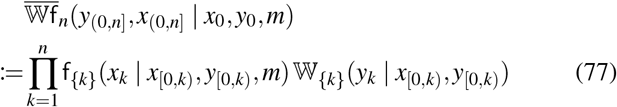

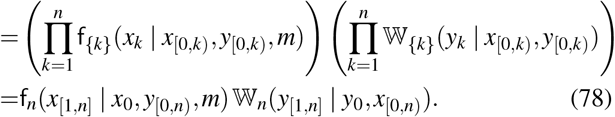

Line (77) exhibits the factorization at the level of the one-step conditional transition law of the joint process, while the last line is the corresponding factorization of the induced path densities. The product in Eq. (78) is therefore understood at the density level, not as a product of measures. In this way, the construction defines a generally non-Markovian joint process whose conditional transition law at each step *k* has the product form

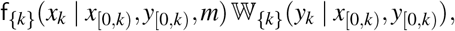

while the induced path density factorizes accordingly over the full time interval. Thus, the encoder and the channel may still be “strongly” coupled through their common dependence on the past history, but the construction excludes same-time instantaneous causal dependence between the encoder update and the channel output update.

Defining the joint law of encoder and channel for a general time structure at the level of generality used so far requires a more elaborate measure-theoretic construction, which we do not pursue here. A general construction principle is outlined in [35, App. A] and is briefly revisited in Appendix Sec. B. The discrete-time factorization above should therefore be understood as fixing the type of joint law we wish to realize also in general time: conditionally on the past history, the encoder update and the channel update are generated separately rather than jointly. In natural discrete time this still allows both variables to change within the same coarse time slot. For continuous-time jump processes, however, the analogous construction excludes simultaneous jumps of *X* and *Y* almost surely. This matches our interpretation of encoder and channel as distinct causal devices. If one started instead from a causally coupled joint law in which the new values of *X* and *Y* were generated jointly given the past, then causally conditional probabilities of *Y* given *X* and of *X* given *Y* and *M* could still be derived from that joint law. However, the channel kernel and encoder kernel alone would then no longer determine the full joint law, because they would not encode the instantaneous coupling itself.

For the continuous-time jump processes associated with the CRN models considered in this paper, one can work with explicit trajectory-level likelihoods, namely local Janossy densities [98, Ch. 7.3]. In this setting, the discrete-time causal factorization in Eq. (78) is replaced by a corresponding factorization at the level of trajectory likelihoods, as discussed at the end of Sec. VI. More specifically, Kramer’s notion of causally conditional probabilities in discrete time was extended to general time by Spinney *et al*. [35], who also derived the corresponding Janossy-density expressions for continuous-time jump processes. These likelihoods can be used to represent stochastic kernels such as W_*t*_ and f_*t*_, provided they arise from the construction method of Spinney *et al*. on an ambient probability space ℙ. In this case, we will also call the kernels ℙregular, as detailed in the following (cf. Def. 10). Since we introduce these local Janossy densities later in the CRN setting, in Sec. VI, together with the bipartiteness condition that excludes simultaneous *X*- and *Y*-jumps, we do not formalize the construction here.

For the present discussion, we simply assume that the joint law for infinite channel operation

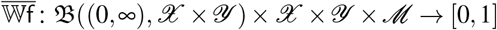

is available and determines the corresponding joint law for channel operation over the finite interval (0, *t*], namely

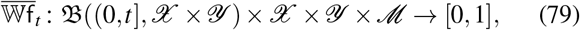

which satisfies

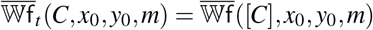

for all *t* ∈ T_*>*0_, all *C* ∈ B((0, *t*], *X* × *Y*), and all (*x*_0_, *y*_0_, *m*) ∈ *X* × *Y* × ℳ.

#### Induced channel, error probability and ℙ-consistency

The joint law allows us to specify the induced channel from ℳ to 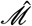 for the operation over (0, *t*] as

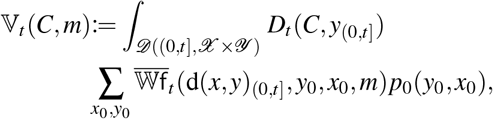

for 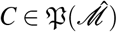 and *m* ∈ ℳ. This induced channel V_*t*_ allows us to specify the probabilities of a decoding error as in Eqs. (49), (50).

As already mentioned, Def. 3 for connection via a channel is no longer viable once feedback is used and the joint law of encoder a standard joint distribution. Causality for both the channel and the encoder needs to be respected by any consistent ambient probability measure ℙ.

##### Definition 10

(Causal connection via channel and encoder). *Let* (Ω, ℱ, ℙ) *be a probability space with filtration* ℱ = {ℱ_*t*_}_*t*≥0_ *and*

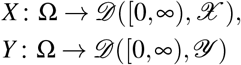

*be adapted stochastic processes (i*.*e*., *X*_[0,*t*]_ *and Y*_[0,*t*]_ *be* ℱ_*t*_*measurable for all t* ∈ T_≥0_*). We say that X is causally connected to Y by the channel* W *if for all B* ∈ B((0, ∞), *Y*)

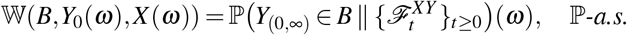

*Similarly, if M* : Ω → ℳ *is a* ℱ_0_*-measurable random variable, we say that Y, given M, is causally connected to X by the encoder* f *if for all A* ∈ B((0, ∞), *X*)

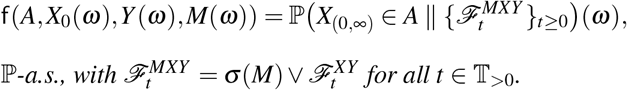

In analogy to “regular conditional distributions” we introduce a similar notion for causally conditional probabilities in Appendix B. Then, satisfying the above definition is equivalent to saying that W is a ℙ-regular causally conditional distribution of *Y* given *X* and that f is a ℙ-regular causally conditional distribution of *X* given *Y* and *M*. In the following we will use the short notation in Eq. (B1) for a ℙ-regular causally conditional kernel and write W = ℙ^*Y* ∥*X*^ or f = ℙ^*X* ∥*Y,M*^. This should be distinguished from the ordinary conditional law ℙ^*X* |*Y*^, which may serve as a posterior distribution and is applicable, e.g., in decoding methods like maximum-a-posteriori estimation [99]. However, it does not play the role of the causal channel law in the coding problem.

Again, we also call a communication system ℙ-consistent if the joint distribution induced by 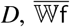, *p*_0_ and *p*_*M*_ equals the joint distribution of 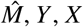 and *M* on (Ω, ℱ, ℙ). We emphasize that a probability ℙ can satisfy W = ℙ^*Y* ∥*X*^ without being ℙ-consistent for a specified communication system, which will be needed in the following for the formulation of coding theorems for causal channels with (noisy) causal feedback.

### F. General channel capacity under noisy causal feedback: Preliminaries and Generalizations

In channel code design, one usually optimizes over deterministic objects that specify the encoder, rather than over the stochastic signal realizations generated during transmission. In the classical setting without feedback, these objects are codewords. With feedback, they are replaced by code functions, namely sequences of functions of the feedback information available at the encoder. This representation was introduced to convert the feedback coding problem into an equivalent problem without feedback [99]. How these functions are defined depends on the type of feedback signal that the encoder receives from a feedback link. To connect with the classical feedback-capacity literature, we therefore briefly return to discrete time and deterministic code-function representations.

#### Ideal feedback link in classical communication

A prominent case in the literature is the ideal feedback link [37, 99, 100]. In this case, the received signal is sent back to the encoder without noise corruption, but possibly with a delay. The corresponding code functions are then deterministic special cases of the causal feedback encoders f discussed above. In the classical discrete-time notation with symbol index, one considers sequences of the form

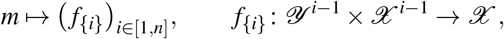

where the function value represents the symbol *x*_*i*_. Thus, for a fixed code-function sequence *f*_[1,*n*]_, the input symbols are generated recursively according to

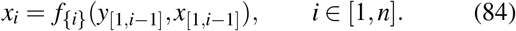

This recursive structure satisfies the causality requirement. Substituting it into the causal channel law yields the effective causally conditioned law from code functions to output trajectories,

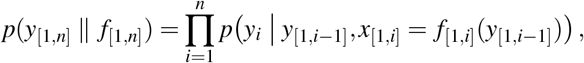

where *x*_[1,*i*]_ = *f*_[1,*i*]_(*y*_[1,*i*−1]_) means that the sequence *x*_1_, …, *x*_*i*_ is generated recursively by Eq. (84). Hence, once *f*_[1,*n*]_ is fixed, the channel output law is fully determined by the causal channel together with that recursive input construction.

Accordingly, the feedback coding problem may be reformulated as an equivalent problem without feedback, in which the input alphabet is the space of admissible code-function sequences and the corresponding optimization variable is a probability measure 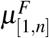 on that space. The associated information quantity is I(*F*_[1,*n*]_;*Y*_[1,*n*]_). The same measure 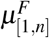 induces a causally conditional input kernel,

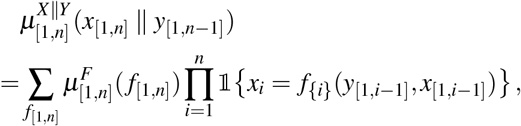

whose evaluation at a fixed feedback realization *y*_[1,*n*−1]_ is precisely the induced probability law of the input trajectory *x*_[1,*n*]_. Tatikonda and Mitter [99] show that the capacity problem can therefore be rewritten directly as an optimization over the induced causally conditional kernels *µ*^*X* ∥*Y*^, rather than over the auxiliary measures 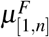 on code-function space. Under the assumption of information stability, the coding theorem then reads

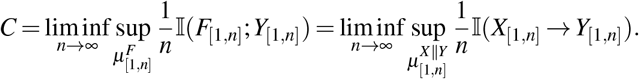

The second equality reflects the passage from the auxiliary code-function representation to the induced physical channel-input process. A probability measure 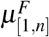 on admissible code-function sequences induces, through the causal recursion of the encoder, a causally conditional input kernel *µ*^*X*∥*Y*^. The feedback-capacity optimization can therefore be reformulated on the space of induced causal input laws. At that level, the appropriate information measure is directed information from *X*_[1,*n*]_ to *Y*_[1,*n*]_, not because *F*_[1,*n*]_ and *X*_[1,*n*]_ are identical random objects, but because the resulting information quantity depends only on the causal law of the physical input process [99].

#### Noisy feedback link in classical communication

A second important case in the literature is a noisy feedback link [101, 102]. In this setting, the encoder does not observe the channel output history directly, but only through a noisy causal feedback channel with fixed law

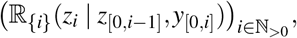

where ℤ is a finite alphabet. The corresponding deterministic code functions are then of the form

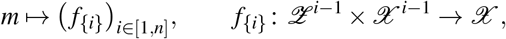

so that, for a fixed code-function sequence *f*_[1,*n*]_, the input symbols are generated recursively by

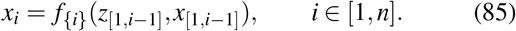

Given the time order *X*_1_ → *Y*_1_ → *Z*_1_ → *X*_2_ → *Y*_2_ → *Z*_2_ · · ·, the forward channel law together with the feedback channel law induces the effective causally conditioned joint law

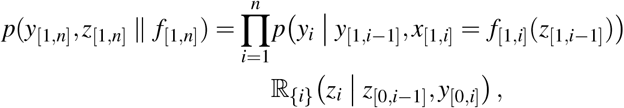

and hence the effective causally conditioned law from code functions to channel outputs

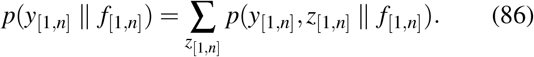

Thus, once *f*_[1,*n*]_ is fixed, the output law is fully determined by the causal forward channel, the noisy feedback channel, and the recursive input construction in Eq. (85).

As in the ideal-feedback case, the coding problem may therefore be reformulated as an equivalent problem without feedback, in which the input alphabet is the space of admissible code-function sequences and the corresponding design variable is an auxiliary probability measure 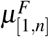 on that space. The same measure induces a causally conditional input kernel

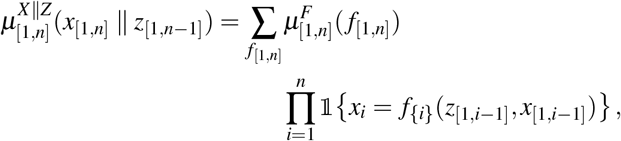

whose evaluation at a fixed noisy-feedback realization *z*_[1,*n*−1]_ is the induced probability law of the input trajectory *x*_[1,*n*]_.

In contrast to ideal feedback, however, the corresponding information measure is no longer ordinary directed information from *X*_[1,*n*]_ to *Y*_[1,*n*]_, but residual directed information, i.e., directed information with the contribution of the noisy feedback process removed; see Sec. V, Eq. (100) for the formal definition. Li and Elia [101, 102] show that the noisy-feedback capacity problem can therefore be reformulated as an optimization over the induced kernels 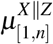, rather than over the auxiliary measures 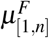 on code-function space. Under the assumption of information stability, the resulting coding theorem reads

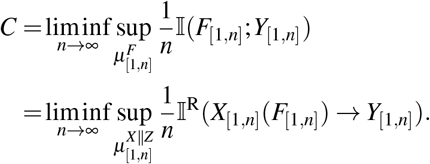

The second equality reflects the same reduction principle as in the ideal-feedback case: a measure on admissible code-function sequences induces a causally conditional kernel on the physical channel inputs, and the corresponding residual directed information depends only on that induced kernel.

In the biochemical communication models proposed in this work, the role closest to code functions is played by deterministic signaling molecule chemostat protocols. The reduction of the optimization problem to *µ*^*X* ∥*Z*^, however, relies on two premises that do not hold in our setting: first, the remaining part of the encoder law is assumed fixed given the specified noisy feedback link; second, the channel input symbols are generated deterministically from the noisy feedback history. In our biochemical communication model, by contrast, the channel-encoding trajectory is itself intrinsically stochastic, so that the above reduction does not apply directly.

#### Encoding specifications and finite-duration information measures

To formulate general coding theorems under feedback, we begin with an auxiliary prior on a space of deterministic encoding specifications. This is directly analogous to the classical coding-theorem constructions: without feedback one uses auxiliary priors *µ*^*X*^ on deterministic codewords, whereas with feedback one uses auxiliary priors *µ*^*F*^ on deterministic code-function sequences.

Let ℰ be a measurable state space, and let the admissible deterministic encoding specifications form a measurable sub-space of

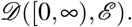

In analogy with the classical feedback setting, we denote by *µ*^*F*^ an arbitrary probability measure on this admissible specification space. In our biochemical communication model, such a deterministic encoding specification is given by an admissible signaling-molecule chemostat protocol trajectory. In contrast to the classical setting, the specification state space ℰ need not coincide with the channel-input state space *X*.

Once a deterministic encoding specification *f* ∈ *D* ([0, ∞), ℰ) is fixed, the feedback encoder architecture and the causal channel law W jointly determine the output statistics through the causality-preserving construction introduced above for 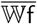. That is, conditionally on the accumulated past, the encoder update and the channel update at each time *s* ∈ (0, *t*] during operation are generated separately, and neither depends on the other’s simultaneously generated new value. In the classical discrete-time noisy-feedback setting, this construction reduces precisely to the effective causally conditioned law *p*(*y*_[1,*n*]_ ∥ *f*_[1,*n*]_) in Eq. (86). We now introduce its general-time analogue as a causal stochastic kernel

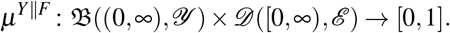

Thus, for each fixed specification trajectory (*f*_0_, *f*), the map *B* ↦ *µ*^*Y* ∥*F*^ (*B, f*) gives the probability law of the resulting output trajectory on the event set *B*.

For each *t* ∈ T_*>*0_, we denote the corresponding finite-duration kernel on (0, *t*] by

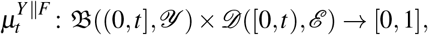

defined through restriction of the infinite-time kernel as

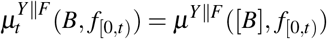

for all *B* ∈ B((0, *t*], *Y*), where 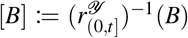, and for any representative 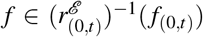. In this way, 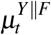 plays exactly the same role for general time structure that *p*(*y*_[1,*n*]_ ∥ *f*_[1,*n*]_) plays in the discrete-time construction. The corresponding marginal output measure is

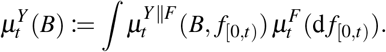

Together, 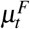 and 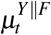 define a joint measure

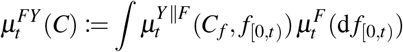

for all

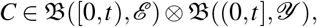

where

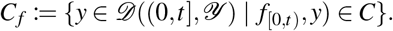

We also define the corresponding product measure

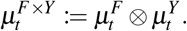

Whenever 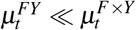, we define the mutual information density by

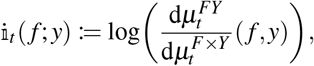

and the associated mutual information by

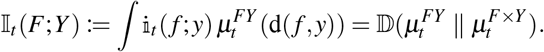

Thus, I_*t*_(*F*;*Y*) is the amount of information that the output trajectory over (0, *t*] carries about the deterministic causal encoding specification selected at time zero. For channels with noisy feedback, these are the natural finite-duration operational quantities.

#### Fixed encoder architecture and causal connection

To make the role of the deterministic encoding specification more explicit, we introduce a fixed causal stochastic kernel ℍ from *X* × *D* ([0, ∞), *Y*) × *D* ([0, ∞), ℰ) to *D* ((0, ∞), *X*), which represents the encoder architecture together with all parts of the feedback side of the communication system that do not vary depending on the sent message *m*. The point of this parametrization is that all message-encoding parts of the encoder are absorbed into the specification process *F*. Thus, once the design space of *F* is fixed, the kernel ℍ determines a fixed feedback-encoder architecture which processes message-dependent information under consideration of feedback at the encoder side. This structure is visualized in Fig. 5.

**FIG. 5.**
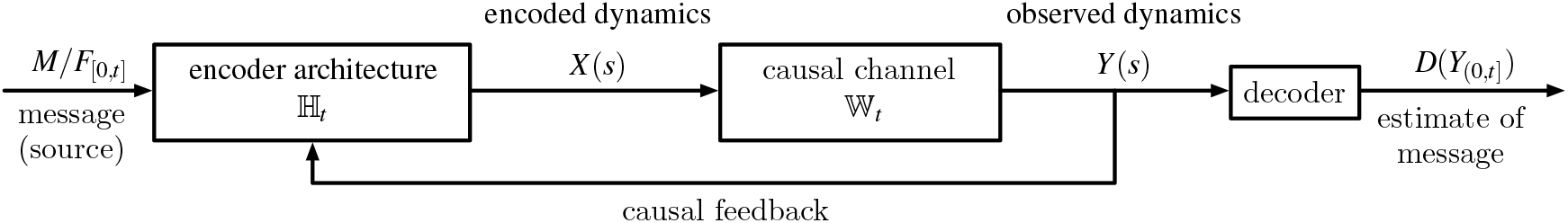
Schematic of a general communication model with noisy feedback. A message *M*, or equivalently its deterministic source-side encoding specification *F*_[0,*t*]_, initiates transmission. Through the fixed encoder architecture ℍ_*t*_, this specification generates the stochastic channel-encoding dynamics *X*, while the observed dynamics *Y* are fed back causally to the encoder. The fixed causal channel W_*t*_ maps the encoded dynamics *X* to the observed dynamics *Y*. A deterministic decoder *D*, acting on the full observation *Y*_(0,*t*]_, produces an estimate of the message. The figure emphasizes the separation between deterministic source-side design, stochastic encoder-side dynamics, and the fixed channel law from *X* to *Y*.

We emphasize again that all message-dependent encoding design degrees of freedom are represented by the state space of *F*. In contrast, the kernel ℍ fixes the feedback encoder architecture for all possible message encodings. The codingtheorem optimization is then performed over auxiliary priors on admissible specification trajectories *F* for a fixed kernel ℍ.

A classical example of a fixed encoder architecture ℍ, for deterministic encoding specifications of the form (85), is given by a discrete-time noisy feedback link. In this case,

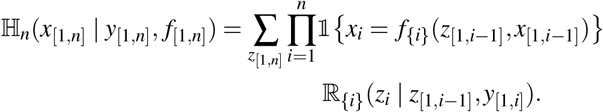

The previously introduced output kernel *µ*^*Y* ∥*F*^ should therefore be understood as the marginalization of the joint causal kernel 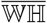, to the output trajectory space by integration over *D* ((0, ∞), *X*).

##### Definition 11

(Causal connection via a fixed encoder architecture). *Let* (Ω, ℱ, ℙ) *be a probability space with filtration* {*Ft*}*t≥0, and let*

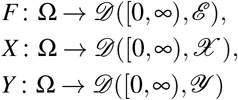

*be stochastic processes, where F is* ℱ_0_*-measurable and X and Y are adapted. Let* ℍ *be the fixed causal kernel of the encoder architecture. We say that X, given F, is causally connected to Y by the fixed encoder architecture* ℍ *if for all*

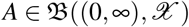

*it holds that*

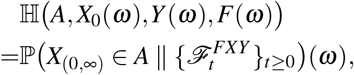

ℙ*-a*.*s*., *where*

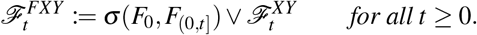

In other words, ℍ is the ℙ-regular causally conditional distribution of the channel-input trajectory *X*_(0,∞)_ given the observed output trajectory *Y*, the initial channel-input state *X*_0_, and the deterministic encoding specification *F*.

#### Reduction to admissible kernel classes

Although the finite-duration information quantities introduced above are naturally expressed in terms of the auxiliary prior *µ*^*F*^ on deterministic encoding specifications, the corresponding optimization problem is often more conveniently rewritten in terms of the induced causal feedback law on the channel input process. More precisely, given the fixed encoder architecture ℍ and an auxiliary prior *µ*^*F*^, one obtains a causally conditional kernel on the physical channel-input process by averaging ℍ over the specification space. On the infinite time interval, this induced kernel is

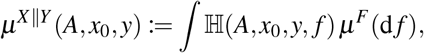

for all *A* ∈ B((0, ∞), *X*), and with the standard restriction for finite duration *t* ∈ T_*>*0_. Thus, *µ*^*X* ∥*Y*^ is the causal law of the channel-input trajectory induced by the fixed encoder architecture under the auxiliary prior *µ*^*F*^. Note that this construction did not include the initialization *p*_0_ in the encoding specifications *F* because the initialization is here not considered as a message-encoding design freedom, but rather a parameter of the whole communication system. Consequently, it needs to be treated separately.

In the classical ideal-feedback setting, this construction is exactly the familiar kernel *µ*^*X* ∥*Y*^ induced by a prior on deterministic code-function sequences [99]. In the classical noisy-feedback setting, by contrast, one first separates the fixed noisy feedback link from the remaining encoder-side design freedom. The latter is then represented by a kernel *µ*^*X* ∥*Z*^, which describes the causal law of the channel input given the noisy feedback signal *Z* and should therefore be regarded as the remaining optimization variable rather than as the full induced law *µ*^*X* ∥*Y*^ itself [101].

In general, the relevant induced kernel class *µ*^*X* ∥*Y*^ depends on which parts of the encoder architecture are treated as fixed and which parts remain designable. We may therefore define a class of causal input kernels by declaring a kernel to belong to that class whenever it is induced by some allowed auxiliary prior on deterministic encoding specifications through an architecture ℍ. The explicit form of that auxiliary prior is then irrelevant for the subsequent coding theorem; what matters is only the induced causal law *µ*^*X* ∥*Y*^ of the channel input process (or the part of it that varies with *µ*^*F*^). In this way, all restrictions on admissible encoders – for example those imposed by a fixed noisy feedback link or by further mechanistic constraints – are absorbed into the resulting class of causal input kernels *µ*^*X* ∥*Y*^.

Whenever the relevant information quantity depends only on that induced law – as in the ideal-feedback case, and in the noisy-feedback case after passage to residual directed information – the original optimization over auxiliary priors *µ*^*F*^ on deterministic encoding specifications can be replaced by an equivalent optimization over the induced kernel class *µ*^*X* ∥*Y*^. However, this reformulation does not in general imply a reduction in the complexity of the optimization problem [103].

#### Information rates

We now pass to the corresponding asymptotic rates. For simplicity, we again first assume that W and ℍ do not depend on the initialization state *x*_0_, *y*_0_.

##### Definition 12

(Information rates from encoding specifications to outputs). *Let*

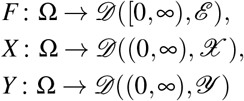

*be random variables on the probability space* (Ω, ℱ, ℙ). *Suppose that F is* ℱ_0_*-measurable, that X, given F, is causally connected to Y by the fixed encoder architecture*

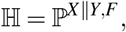

*and that X is causally connected to Y by the channel kernel*

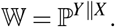

*If* 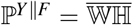 *is the induced causally conditional law of Y given F, and* : i_*t*_ (*F*;*Y*) *denotes the mutual information density associated with the pair* 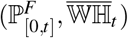 *on the finite interval* [0, *t*], *as introduced above*.

*Then*

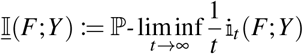

*is called the* inf-information rate *from the encoding specification F to the output Y*. *Further*,

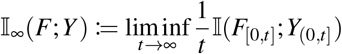

*is called the* information rate *from the encoding specification F to the output Y*.

Since the encoding specification *F* is selected at time zero and is not causally influenced by the channel output, the asymptotic quantities above are defined in terms of ordinary mutual information rather than directed information. The channel coding problem, even under noisy feedback, is hence reduced to the non-feedback case.

To include the effect of a designable or non-designable initialization *p*_0_, we need the adaptation

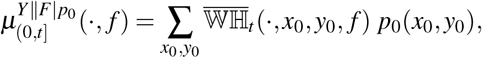

since the actual initialization state (*x*_0_, *y*_0_) will be unknown to both encoder and decoder. The mutual information measures are then taken with respect to the adapted pair

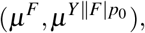

where *µ*^*F*^ is as before, *p*_0_ is any admissible initialization probability mass functions that merely acts as a parameter and not as a kernel input argument. Importantly, *p*_0_ does not belong to the deterministic encoding specifications in the sense that its choice is not part of finding a good channel code. Instead, it is a parameter under which any choice of channel code may perform better or worse.

#### General coding theorems and information stability

Because the encoding specification *F* is selected at time zero, the feedback coding problem is reduced to an effective non-feedback problem from *F* to *Y*. Consequently, the natural analogue of the general coding theorem of Verdú and Han [93] is

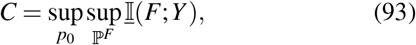

where the supremum is taken over all basic probability measures ℙ such that *F* is ℱ_0_-measurable, *p*_0_ = ℙ((*X,Y*)(0) = ·), *X* given *F*, is causally connected to *Y* by the fixed encoder architecture ℍ = ℙ^*X* ∥*Y,F*^, and *X* is connected to *Y* by the channel W = ℙ^*Y* ∥*X*^.

As we do not prove this generalized channel coding theorem with noisy feedback, we state it here as a conjecture. This conjecture is however, strongly supported by the generalized Feinstein-lemma [77], which in adaptation to our case states that for every message set ℳ, any *p*_0_, *µ*^*F*^ and any *γ, t >* 0 exists an (|ℳ|, *t*)-code with standard code rate 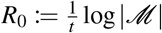 such that the maximum probability of error is bounded as

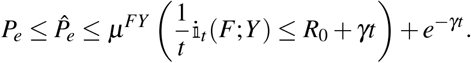

When the r.h.s. is less than one, it ensures the existence of an injective code. This Feinstein-lemma is a fundamental bounds that is used to prove general coding theorems [93].

Under the additional assumption of information stability, this general formula reduces to the Shannon-type expression

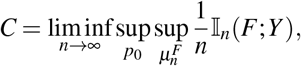

provided there exists an auxiliary prior *µ*^*F*^ (equivalently a consistent probability measure ℙ) such that

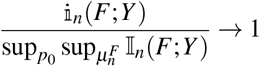

in probability as *n* → ∞. Thus, after reduction to deterministic encoding specifications, the formal structure of the coding theorem is the same as in the non-feedback case, with *F* replacing the usual channel input variable.

Lastly, we add a note of caution w.r.t. these conjectured coding theorems. While they are well motivated when considering the standard code rate *R*_0_ and the maximum probability of error 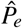 as performance criteria, there is still a theoretical gap for the formulation of coding theorems, where the message distribution is constrained and one therefore instead considers the mean code rate *R* and the message-weighted probability of error *P*_*e*_. In particular, the auxiliary prior *µ*^*F*^ was never required to contain any information about *p*_*M*_ such that the r.h.s. of the coding theorems is always independent of the message distribution.

### G. Adaptation to biochemical communication

In contrast to classical noisy-feedback coding, where the design object is typically a deterministic code function that causally maps an available feedback signal to the channel input, the biochemical communication model developed here is organized around deterministic encoding specifications that act through the CRN dynamics. In the present setting, each message selects a deterministic signaling protocol, represented by non-negative trajectories of selected signaling molecules. These signaling molecules form a subset of the chemostatted species introduced in Sec. III B. The protocol is therefore not itself the channel-input trajectory *X*, but the deterministic source-side specification that governs how the stochastic channel encoding is generated. The resulting dynamics of *X*, together with the propagation of feedback through the network, are then determined by the encoder-side architecture ℍ.

This follows the same structural principle as in the classical coding-theorem framework: the design variables are deterministic specifications of how encoding is carried out, while the transmitted trajectory may still be stochastic. In the classical setting, these specifications are codewords or code-function sequences. In the biochemical setting, they are signaling-molecule chemostat protocols and, in more general formulations, families of such protocols associated with admissible encoder architectures. This allows the biochemical communication problem to be formulated, as before, as an optimization over auxiliary priors on deterministic encoding specifications *F*, while the stochastic part of the encoder side is absorbed into the kernel ℍ.

What carries over directly is that the physical channel remains represented by a fixed causal law W = ℙ^*Y* ∥*X*^. Accordingly, all basic probability measures over which we optimize must be consistent with the channel law W and with the chosen encoder-side architecture ℍ = ℙ^*X* ∥*Y,F*^. The initialization may likewise be part of the design problem. In that case, the preparation of the initial distribution can be viewed as a separate pre-communication control stage, for instance by holding chemostats at prescribed levels until the population distribution approaches the desired initialization law *p*_0_.

Within this formulation, the fundamental asymptotic quantity is the inf-information rate I(*F*;*Y*) from the deterministic encoding specification to the output trajectory. When the induced pair (*F,Y*) is information stable, this reduces to the ordinary information rate 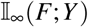. In the biochemical setting, this means that the admissible signaling-molecule protocol trajectories, and hence the auxiliary prior *µ*^*F*^, cannot be chosen arbitrarily if one wants a Shannon-type capacity expression in terms of I_∞_(*F*;*Y*). In general, a reduction to simpler induced kernel classes, as in the classical examples, is no longer guaranteed and must instead be verified from the structure of the specific biochemical communication model under consideration.

## V. DIRECTED INFORMATION IN CONTINUOUS TIME WITH NATURAL TIME STRUCTURE

Here we discuss directed information measures for general continuous time stochastic processes. As the proofs are often technical, we only present results and refer the reader to the Appendix or Supplement for further details.

### Definitions

For discrete sequences, the directed information (DI) was originally defined by Massey [11] as

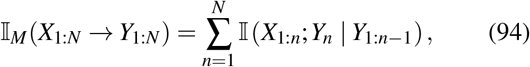

where *X*_1:*N*_ = {*X* (*n*)} _*n* ∈{ 1,…*N*_} and *Y*_1:*N*_ = {*Y* (*n*)} _*n* ∈{ 1,…*N*_} are random sequences for all *N* ∈ ℕ. As noted by Newton [85], this definition was developed in the context of communication channels, where *Y*_*n*_ = *Y* (*n*) is received after *X*_*n*_ = *X* (*n*) is transmitted. Thus, the enumeration index *n* does not represent physical time for the joint process {(*X* (*n*),*Y* (*n*))} _*n* ∈{1,…*N*}_. If the index *n* does denote physical time for the joint sequence, then (94) includes undirected “instantaneous information exchange” terms [104] of the form I(*X*_*n*_;*Y*_*n*_ | (*X,Y*)_1:*n*−1_), which should be excluded when aiming to quantify the causal influence of *X*_1:*N*_ on *Y*_1:*N*_.

To address this, Newton proposed a causally faithful definition of DI for discrete sequences, where the index represents physical time:

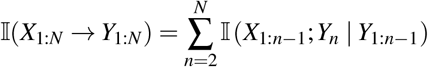

To generalize this definition to continuous-time trajectories, we follow the extremal-based approach of Weissman *et al*. [37], who defined the directed information for continuous processes by analogy with Massey’s formulation as

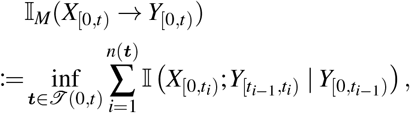

where *T* (*t*_0_, *t*) = {***t*** = (*t*_*i*_)_*i* _∈{_1,…,*n*}_ : *t*_0_ *< t*_1_ *<* · · · *< t*_*n*_ = *t*} is the set of all finite partitions of an interval [*t*_0_, *t*), *t*_0_ *< t*, and *n*(***t***) denotes the length of the partition. However, we modify the definition to follow the causal structure of Newton’s proposal instead. Thus, for continuous-time processes on the right-open interval [0, *t*), we define

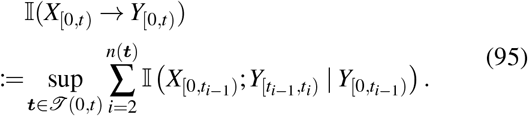

The use of the supremum, in contrast to the infimum, is justified in Supp. Lemma S1.1.

Directed information on a closed interval [0, *t*] is given by the right-limit

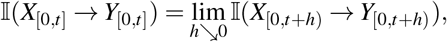

and the directed information rate (DIR) is

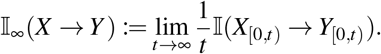

In other works, such as [36, 85], directed information and transfer entropy have been extended to continuous time by taking the continuity limit of their discrete-time forms to construct a time-local rate, which is then integrated. The limitation of this approach for general continuous-time processes is that boundary terms or discontinuous jumps of the information measure as a function of time may be overlooked (although such discontinuities do not occur for the well-behaved jump processes considered here). In contrast, the approach of Weissman *et al*. provides a structurally sound alternative, as it is grounded in measure theory through the definition of generalized divergence (39) and thus defined directly in terms of continuous-time path space probability measures, equivalent to Radon-Nikodym derivative-based definitions (cf. Appendix B or [35, App. A]).

### Integral representations

In Supp. Sec. S1.4 we show that, under certain continuity conditions, the DI satisfies

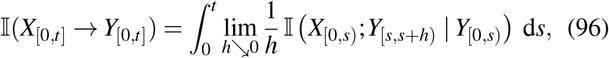

which is easier to evaluate in practice than definition (95). Similarly, for the continuous generalization of Massey’s DI, we obtain

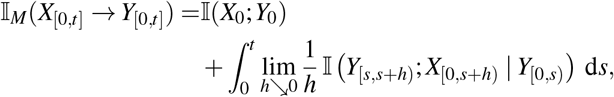

under the same continuity conditions. A similar expression was previously proposed by Weissman *et al*. [37], though without accounting for potential discontinuities or including the MI of the initial values, and without a formal proof.

### Conservation laws

If I(*X*_[0,*t*)_;*Y*_[0,*t*)_) *<* ∞, then mutual and directed information are related through the information conservation law

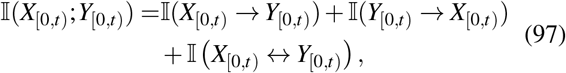

where the continuous-time instantaneous information exchange is defined as

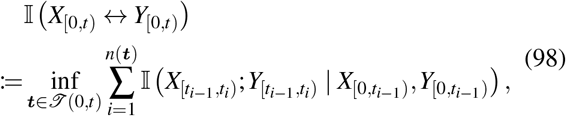

generalizing the corresponding discrete-time definition [104]. This conservation law has already been addressed in [36, Eq. (38)] and follows from its discrete counterpart in [85], as we show in Supp. Proposition S1.1.

In contrast, the Massey DI obeys the more asymmetric conservation law [37]

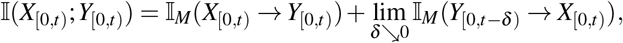

where the delayed-input Massey DI is defined by

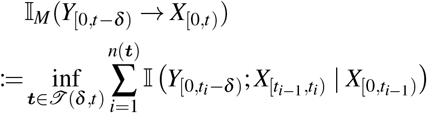

for any delay *δ* ∈ (0, *t*].

### Conditional and causally conditional directed information

The conditional DI given a random variable *Z*, and the causally conditioned DI given an ℱ-adapted stochastic process *Z*_[0,*t*)_ = {*Z*(*s*)}_*s*∈[0,*t*)_, are defined respectively as

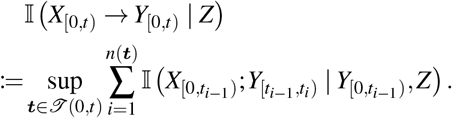

and

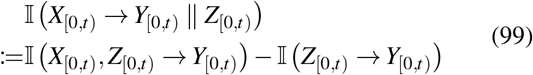

if I *(X*_[0,*t*)_, *Z*_[0,*t*)_ → *Y* _[0,*t*)_)*<* ∞ and I (*Z*_[0,*t*)_ →*Y*_[0,*t*)_ *<* ∞. Definition (99) is motivated by the chain rules for the discrete-time Newton DI, as detailed in Appendix A. An extremal-based definition is not viable since

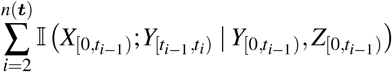

is not necessarily monotonous under refinement of the partition ***t*** ∈ *T* (0, *t*). However, there exists a sequence of partitions that simultaneously approximates I (*X*_[0,*t*)_, *Z*_[0,*t*)_ → *Y*_[0,*t*)_ and I (*Z*_[0,*t*)_ →*Y*_[0,*t*)_, and hence also I (*X*_[0,*t*)_ → *Y*_[0,*t*)_ ∥*Z*_[0,*t*)_). This can be shown in analogy to the proof of Supp. Lemma S1.2. Similarly, we define

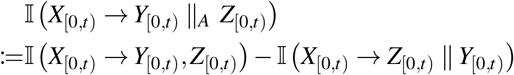

if the r.h.s. exists. Here, ∥_*A*_ denotes “anticipatory” causal conditioning. This definition is again motivated in Appendix A, and it can be shown that there exists a sequence of partitions ***t*** ∈ *T* (0, *t*) such that I *X*_[0,*t*)_ → *Y*_[0,*t*)_ ∥_*A*_ *Z*_[0,*t*)_ is approximated by

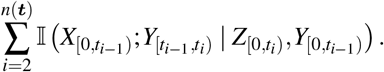

#### Lemma 1

*It holds that*

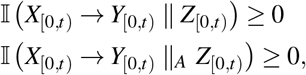

*if the quantity on the l*.*h*.*s exists*.

This lemma follows directly from the non-negativity of the approximating sequences.

*Residual directed information*. Another quantity of interest is the residual directed information, defined by Li and Elia [102] in discrete time as the difference of a directed information and a causally conditional directed information. With the above derivations, their definition can be directly adopted for continuous time, such that

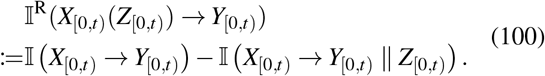

This relatively new information measure sometimes replaces mutual information and even directed information for channels with noisy (non-ideal) feedback due to the following proposition.

#### Proposition 1

*Let M be a* ℱ_0_*-measurable random variable. If* I(*M* → *Y*_(0,*t*)_ ∥ *X*_(0,*t*)_) = 0, *then*

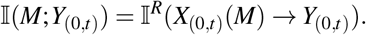

The conceptual add-on compared to the mutual information is that it is defined in terms of information measures that measure the information flow from *X*_(0,*t*)_ to *Y*_(0,*t*)_. Proposition 1 suggests the interpretation that the amount of information exchanged between *M* and *Y*_(0,*t*)_ equals the part of the causal information flow from *X*_(0,*t*)_ to *Y*_(0,*t*)_ that remains attributable to the message *M*, rather than to additional stochasticity of the channel encoding itself. The condition I(*M* → *Y*_(0,*t*)_ ∥ *X*_(0,*t*)_) = 0 thereby encapsulates the notion that the evolution of *Y*_(0,*t*)_, causally given the channel encoding *X*_(0,*t*)_, is (causally) independent of a message variable *M* that initiates information transmission. This is an essential requirement for a communication system to behave causally [11]. An analogous discrete time, finite state proposition has been provided in [101], relying on the entropy decomposition of directed information. As such entropies are not defined on the process level, we provide a short proof, based on chain rules for directed information.

*Proof*.

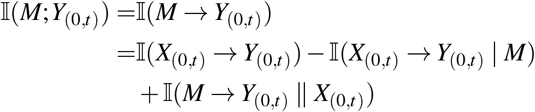

The last term vanishes by the assumption.

## VI. INFORMATION MEASURES FOR CHEMICAL POPULATION TRAJECTORIES

In this section, we derive mutual- and directed-information measures for trajectories of chemical population subnetworks, explicitly accounting for indistinguishable reaction channels under projection, and show that mutual information is well defined only under a bipartiteness condition. The section further introduces directed-information densities, formulated via log-likelihood-ratios, that support the later discussion of biochemical communication systems.

To simplify notation compared to Sec. II, we define the set of directed reaction 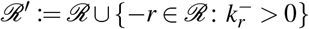, which has cardinality |ℛ^′^|. The directed reaction channels 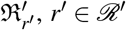 have stoichiometric balanced equations

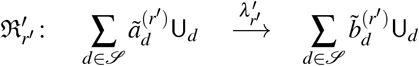

and change vectors 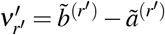. For clarity, we will omit the prime notation in the remainder of this section.

Let *S*_*X*_, *S*_*Y*_ ⊆ *S* be disjoint subsets of chemical species. They specify the two subnetworks between which we want to characterize information exchange. We therefore define the projected processes

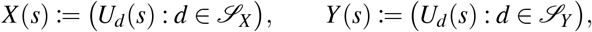

for all *s* ≥ 0 with respective state spaces 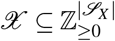 and 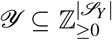, and write *X* = {*X* (*s*)}_*s*≥0_, *Y* = {*Y* (*s*)}_*s*≥0_.

### Subnetwork-indistinguishable reactions

Different reactions of the full network may induce the same net stoichiometric change on the observed subnetwork. To formalize this projection from the full CRN to the *X*- and *Y*-subsystems, we introduce the projected change vectors. Let

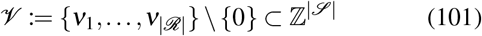

be the set of nonzero change vectors on the full network. Since we only need to project change vectors that actually arise in the CRN, we define

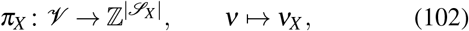

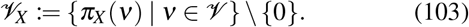

Analogously, we define *π*_*Y*_ and *V*_*Y*_.

#### Definition 13

*The directed reaction channels* ℜ_*r*_ *and* 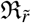 *are called X-indistinguishable if*

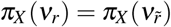

*and analogously for Y*.

In other words, two distinct reaction channels are *X*-indistinguishable if they produce exactly the same observable jump in the *X*-subnetwork. This distinction is essential because the trajectory of *X* resolves only the projected stoichiometric change, not the underlying reaction label in the full network. As an example, consider the species X, Y, and A together with the reactions

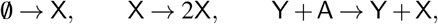

which are all *X*-indistinguishable since each increases the amount of X by one. Definition 13 therefore captures the fact that changes in the marginal subnetwork need not identify the underlying reaction event uniquely. This is important for the definition of the reaction counting processes that generate the trajectories associated with the joint and marginal probability measures 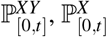 and 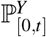.

### Projection to subnetwork CRN models

Let *V* : ℛ → *V* ∪ {0}, *r* ↦ *ν*_*r*_ denote the mapping from reaction indices to their corresponding change vectors. This function is not necessarily injective, as we will see in Sec. III B.

To describe the jump structure of the projected processes, we now merge all reactions that are indistinguishable at the level of the observed subnetwork. We now distinguish between reaction counters of changes in *X* or *Y*. Set

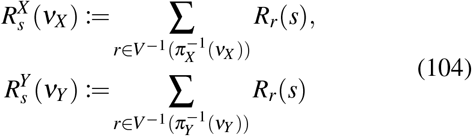

for all *ν*_*X*_ ∈ *V*_*X*_ and *ν*_*Y*_ ∈ *V*_*Y*_. Thus, 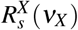 counts how often the *X*-subnetwork has undergone the observable jump *ν*_*X*_ up to time *s*, irrespective of which full-network reaction produced that jump. The analogous interpretation holds for 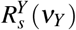.

In this notation the time dependence is shifted to the lower index and the functional arguments are the respective marginal change vectors. The corresponding propensity functions of subsystem changes are

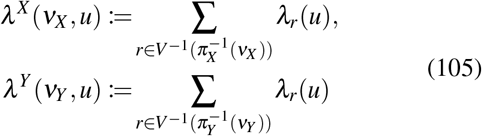

These merged counting processes provide a jump representation of *X* and *Y* in which reactions that are indistinguishable at the subsystem level have been combined.

With this notation, we obtain for all *s* ≥ 0 the process equations

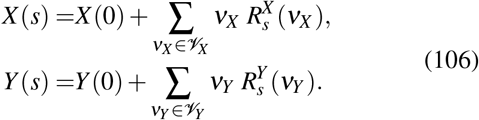

To compute the mutual information, it is not enough to describe the subsystem dynamics through the current state of the full CRN. Instead, we need the predictable jump intensities of the merged counting processes relative to the filtrations generated by the observed histories. This leads from the state-dependent propensity functions to history-dependent stochastic intensity processes. For intuition, one may read the dependence on the filtration 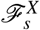 as dependence on the observed trajectory segment *X*_[0,*s*]_ up to time *s*, even though the conditional expectations themselves are still defined as 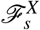 measurable functions on the basic probability space Ω.

To compute the MI, we must represent the evolution of *X* and *Y* relative to the histories of the marginalized and joint subsystems, specifically 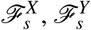, and 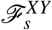, rather than relying on a representation that depends on the current state of *U*. This is done by changing the propensity functions of the reaction counters 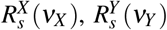 to stochastic processes, that depend on the history of the respective marginal system.

Accordingly, the merged counting process 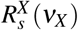 has an 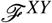-intensity process *λ* ^*X*^ (*ν*_*X*_) and an ℱ^*X*^-intensity process 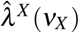 given by [53]

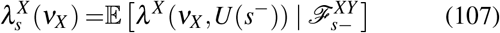

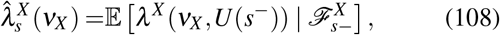

where *U* (*t*−) = lim_*s*↗*t*_ *U* (*s*) (or *U* (*t*^−^)) denotes the left limit in time. Thus, 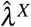 is the intensity seen from the history of *X* alone, whereas *λ* ^*X*^ is the intensity seen from the joint history of (*X,Y*). Analogously, 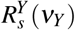 has the ℱ^*XY*^-intensity *λ*^*Y*^ (*ν*_*Y*_) and the ℱ^*Y*^-intensity 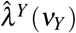 with

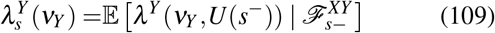

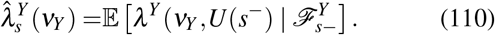

By definition, all jump processes are non-explosive and all intensities are left-continuous processes and therefore satisfy the predictability property [105] w.r.t. their particular filtration.

### Weak and strong bipartiteness

For the path-space information measures below, it is important to distinguish between a structural property of the CRN and a pathwise property of the projected processes. This motivates the following two notions of bipartiteness.

#### Definition 14

*Let X and Y be the subnetwork processes as described above. We say that X and Y are*

a. *weakly bipartite if the probability of a simultaneous jump vanishes, i*.*e*.,

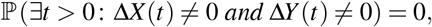

*where* Δ*X* (*s*) := *X* (*s*) − *X* (*s*^−^) *for all s* ≥ 0 *and analogously for Y*.
b. *strongly bipartite if* ℛ *does not contain reactions that simultaneously change X and Y, i*.*e*.,

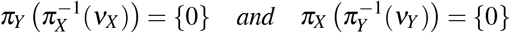

for all *ν*_*X*_ ∈ *V*_*X*_, *ν*_*Y*_ ∈ *V*_*Y*._

Strong bipartiteness is a structural condition on the underlying reaction network: no reaction channel is allowed to change both subnetworks simultaneously. Weak bipartiteness is instead a property of the induced trajectories and only requires that simultaneous jumps of *X* and *Y* occur with probability zero. The weakly bipartite case agrees with the usual notion of bipartiteness for Markovian and non-Markovian jump processes [36, cf. p. 7]. Heuristically, for all *s* ≥ 0 and small *h >* 0,

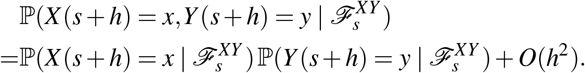

That weak bipartiteness does not imply strong bipartiteness can be seen by considering particular initial conditions. Assume that *X* and *Y* are not strongly bipartite. There may now be certain initial states for which simulatneous jumps in *X* and *Y* do not occurr with probability one, while other initial states do allow for simultaneous jumps with positive probability. For example, consider an isomer that may transition between conformation X and conformation Y. Then *X* and *Y* populations are not strongly bipartite, but they are trivially weakly bipartite given the initial condition ℙ(*X* (0) = 0,*Y* (0) = 0) = 1. On the other hand, that strong bipartiteness implies weak bipartiteness can be formally seen by the discussion in Supp. Sec. S1.2. The reaction counters (104) comprise a multivariate point process in the strongly bipartite case, which directly implies weak bipartiteness as the different coordinates of a multivariate point process are not allowed to jump simultaneously by definition.

### Expressions for mutual information and directed information

Both *X* and *Y*, individually, and (*X,Y*) are “fundamental processes” of the kind discussed in § 2.13 of [105], even without bipartiteness. The following theorem expresses the path-space mutual information in terms of the subsystem intensity processes. The bipartiteness assumption excludes singular simultaneous-jump contributions, while the logarithmic integrability condition ensures that the corresponding likelihood-ratio terms are integrable. The result is similar in structure to that in [18], but we additionally prove that the non-bipartite cases are singular.

#### Theorem 1

*If the conditions*

i. I(*X* (0);*Y* (0)) *<* ∞,
ii. *X and Y are strongly bipartite*,
iii. *the intensities meet the requirement*

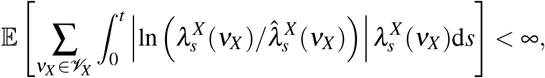 *and analogously for Y*, *are satisfied, then the MI is finite and satisfies*

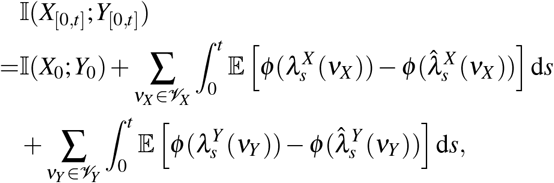

*where φ* (*a*) := ln(*a*) 1_(0,∞)_(*a*) *with a* ≥ 0 *and* 1 *denoting the indicator function of the measurable set in its lower index*.

*If X and Y are not weakly bipartite and exists r* ∈ ℛ *with π*_*X*_ (*ν*_*r*_), *π*_*Y*_ (*ν*_*r*_)≠ 0 *such that* ℰ[*R*_*r*_(*s*)] *>* 0 *for all s >* 0, *then* 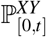 *is not absolutely continuous with respect to* 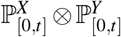 *for all t >* 0.

If absolute continuity fails, then the Radon-Nikodym derivative of the joint path measure with respect to the product of the marginals does not exist, and the mutual information is therefore infinite (cf. Eq. (40)). A proof of the Theorem is provided in Supp. Sec. S1.2.

In the strongly bipartite case, the mutual information splits naturally into contributions carried by jumps in *Y*, jumps in *X*, and the information already present in the initial condition. Two of the terms are identified with directed-informations.

#### Theorem 2

*Under the conditions of Theorem 1 (strongly bipartite case), it holds*

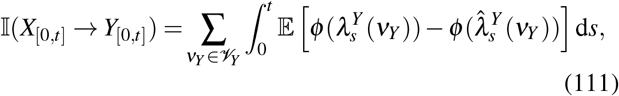

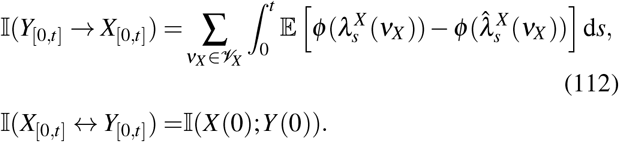

*and for the Massey DI*

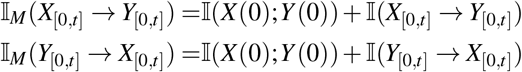

A formal proof is given in Supp. Sec. S1.5. The expressions for Newton’s DI are well-defined even in the nonbipartite case, but require a different method of proof, which we do not provide here. The instantaneous information exchange and the Massey DI both contain simultaneous jump contributions, which cause them to diverge in this case.

### Causally conditional directed information

The same intensity-based construction extends directly to causally conditional directed information. To this end, let *S*_*Z*_ ⊆ *S* be an additional set of species and define the projected process

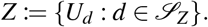

The corresponding coordinate maps and projected nonzero change vectors are defined analogously to (102) and (103).

#### Corollary 1

*Let* (*S*_*X*_ ∪ *S*_*Z*_) ∩ *S*_*Y*_ = ∅. *Then*

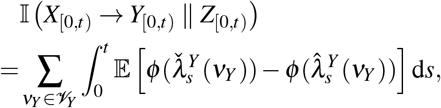

*where* 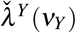 *is the* ℱ^*XZY*^ *-intensity of* 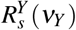 *and*

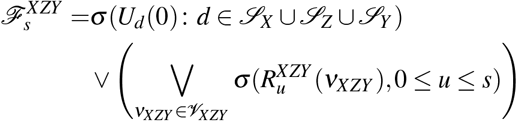

*for all s* ≥ 0.

### Why merging indistinguishable reactions is important

Finally, we return to the role of indistinguishable reactions. The merged counters introduced above are not merely a notational simplification; they are required to correctly represent the histories (filtrations) generated by the observed subsystem. Instead of following the above projection procedure, define the index set

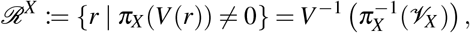

i.e., the set of all reactions that modify *X*. While

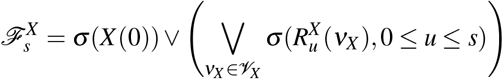

the newly constructed set of reaction counters only satisfies

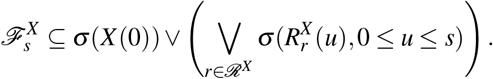

Analogous definitions can be made for *Y* and (*X,Y*).

To build intuition, consider a given trajectory *x*_[0,*s*]_. From *x*_[0,*s*]_, one can uniquely construct the corresponding counting process realizations 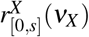 for all *ν*_*X*_ ∈ *V*_*X*_. Conversely, given these counting process realizations together with the initial condition *x*(0), the trajectory *x*_[0,*s*]_ can be uniquely reconstructed. Similarly, *x*_[0,*s*]_ can also be reconstructed from *x*(0) and the realizations of the reaction counters *R*_*r*_, *r* ∈ ℛ^*X*^, on the interval [0, *s*]. However, the reverse is not generally true: if the system includes *X*-indistinguishable reactions, the trajectory *x*_[0,*s*]_ does not contain enough information to determine which specific parallel reaction *r* caused each observed stoichiometric change. A more formal discussion of this equivalence of jump process representations has been provided in [40, Appendix A]. In conclusion, not merging *X*-indistinguishable reactions can lead to misrepresentation of ℱ^*X*^ and thus of the intensity processes 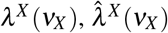. For instance, it must be taken into account in the stochastic filtering equations used to evaluate the various intensity processes [106]. The corresponding filtering equations are provided in Supp. Sec. S1.3.

It is also useful to compare the merged intensity description with the finer description that keeps the original reaction labels. This comparison clarifies how merging indistinguishable reactions affects the directed information expressions. So we consider the ℱ^*XY*^ and ℱ^*X*^-intensity-like processes defined for all *r* ∈ ℛ^*X*^ by

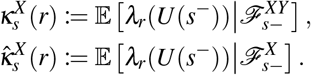

The processes are not always proper intensity processes in the sense of [53, p. 27] since *R*_*r*_, *r* ∈ ℛ^*X*^, are not necessarily ℱ^*XY*^ or ℱ^*X*^-adapted processes.

Similarly, for all *r* ∈ ℛ^*Y*^, define the ℱ^*XY*^- and ℱ^*Y*^-intensity-like processes as

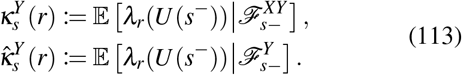

Note that

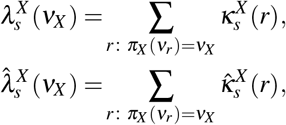

and analogous relations hold for *Y*. By the log-sum inequality,

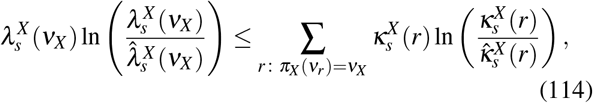

with equality if the sum consists of a single term. Thus, passing from reaction-labeled intensities to merged subsystem intensities can only decrease the corresponding directed-information expression. Equality holds precisely when no nontrivial merging occurs. It follows that the directed information satisfies

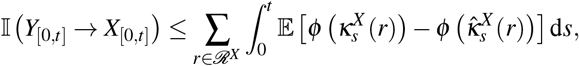

with equality holding when |*V*_*X*_| = |ℛ^*X*^|. Thereby the tower property of the conditional expectation was used to recast both sides of (114) in the familiar representation, that includes *φ*. An analogous inequality holds for I (*X*_[0,*t*]_ → *Y*_[0,*t*]_).

We provide a more detailed discussion of the mismatch of not accounting for indistinguishable reactions in Appendix C, including a precise identification of terms that constitute the mismatch.

### Point process entropies and local Janossy densities

For completeness, we also introduce point process entropies and causally conditioned point process entropies, which extend the corresponding discrete-time concepts of Kramer [84] to the present class of continuous-time processes. Kramer introduced causally conditioned probabilities in discrete time, and Spinney *et al*. [35] generalized this notion to continuous time. These causally conditioned path probabilities will be important in Sec. VII, where they define the chemical communication channel. We denote by 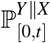 the ℙ-regular causally conditional path probability of 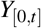 given *X*_[0,*t*]_; see Appendix B for a definition.

The corresponding trajectory likelihoods for jump processes are expressed in terms of local Janossy densities [98, Ch. 7.3]. For the reaction counters 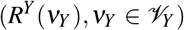, these likelihoods are defined on the space 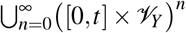, conconditionally on the initial value *Y*(0). The likelihood associated with 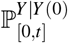 is

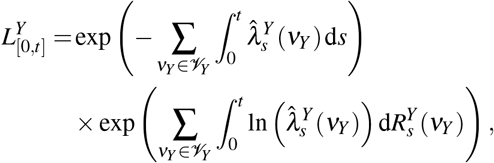

and the likelihood associated with 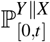 is

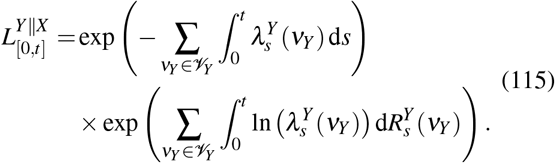

The related notion of point process entropy goes back to McFadden [107], who introduced this particular notion of entropy through the likelihood of a counting-process trajectory. The marginal point process entropy is then defined as

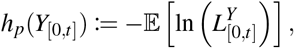

and the causally conditioned point process entropy as

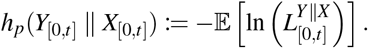

Consequently, the directed information satisfies the familiar relation

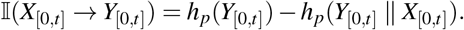

### Directed information density

We now introduce the directed information density, for later use in Sec. VII. Since the encoded signal starts only after initialization, we formulate the density on the post-initialization interval (0, *t*), and the Janossy densities below are understood as the corresponding trajectory likelihoods after averaging over the initialization distribution. Using these Janossy densities, we define

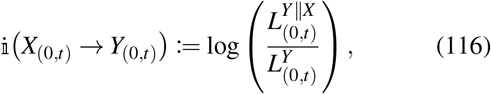

where, as above, the Janossy densities are evaluated along the realized jump trajectory on (0, *t*). Thus, the directed information density is the trajectory-level log-likelihood ratio between the output law causally conditioned on the channel-encoding trajectory *X* and the marginal output law. If the expectation of the directed information density is finite, then it yields the directed information,

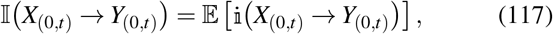

which is equivalently given by the point process entropy difference introduced above.

## VII. CHEMICAL REACTION NETWORKS AS COMMUNICATION CHANNELS

This section develops the communication channel description of chemical reaction networks, depicted in Fig. 1. In this setting, continuous-time path mutual information and directed information are combined with stochastic thermodynamic constraints by interpreting deterministically manipulated CRNs as communication systems. Thereby, information is encoded into controlled chemical signaling dynamics, transmitted through the network, and read out from selected output species, while the corresponding encoding and transmission steps incur thermodynamic costs in the form of heat dissipation, entropy production, or work.

In both information theory and thermodynamics, however, one should be careful with continuous approximations of CRNs, such as the chemical Langevin equation. While heat dissipation may still be well approximated by diffusion models, the property of stochastic consistency (cf. Supp. Sec. S2.6) is typically lost [72]. In a similar way, path-information measures are not consistent under continuous approximations because the continuum limit eliminates “discrete reaction information” [18]. For this reason, the discussion below is formulated directly at the level of the jump-process description.

### A. A causal chemical communication model

This subsection introduces the stochastic CRN model of chemical communication together with the conditional intensity processes and the resulting causal path probabilities. In doing so, it connects the communication-theoretic viewpoint to the CRN primer of Sec. II and to the thermodynamic primer of Sec. III, in particular to the coupling of open CRNs to chemostats discussed in Sec. III B. At this stage, however, our goal is only to specify the underlying stochastic model and its causal structure. This model is later cast into a full noisy channel coding problem in Sec. VII D.

We introduce the communication model of interest, loosely following the marked Poisson-type channel described by Frey [46]. As long as not mentioned otherwise, all random variables defined in this subsection are defined on the probability space (Ω, ℱ, ℙ). Let *t >* 0 be the transmission duration and *M* : Ω → ℳ be a random variable, ℱ_0_-measurable, on a finite index set ℳ. We call *M* the message index (or simply message) and ℳ the message set of a (|ℳ|, *t*)-code. The symbol *R*, without any index, is reserved for mean code rates (cf. Eq. (68)) in this section. Properties of codes and code rates have been introduced in Sec. IV D.

For notational simplicity we assume that all U-parallel reactions *r*_1_, *r*_2_ ∈ ℛ, *r*_1_ ≠ *r*_2_, satisfy 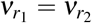, i.e., the change vectors have the same orientation for *ε* = +. For a CRN *U* with |ℛ| potentially reversible reaction channels we redefine the set of change vectors *V* (cf. (101)) to contain only forward change vectors, i.e.,

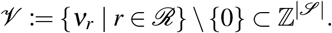

As before in Sec. VI, assume *S*_*X*_, *S*_*Y*_ are disjoint subsets of species. The marginal reaction counters of *X* are

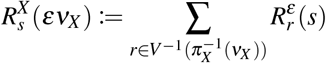

for all *ν*_*X*_ ∈ *V*_*X*_ and *ε* ∈ {+, −*}*, where *V* denotes the mapping from reaction indices to their corresponding change vectors in *V*. Reaction counters for *Y* are defined analogously. Consequently, we have

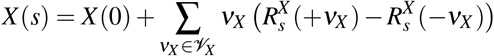

for all *s* ∈ [0, *t*] and analogously for *Y*. At time zero, the system *U* be always prepared with a well defined initial distribution *p*(*u*, 0) for every independent use of the communication channel (and hence independent of *M*).

#### Specification of the source encoder

Assigned to each message index *m* ∈ ℳ is a deterministic manipulation protocol 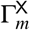 of the open CRN *U*, represented by bounded, piecewise-differentiable càdlàg chemostat trajectories on [0, *t*]. Consequently, the model incorporates source-level control through the deterministic, time-dependent chemostat manipulation previously discussed in Sec. III B. We assume that this protocol is turned on at time 0, and that the map 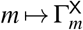 is bijective. The restriction to piecewise-differentiable trajectories serves as a minimal requirement, allowing the protocol to be expressed as an ordinary differential equation with jumps and ruling out potentially pathological cases. Next, we define the ℱ_0_-measurable message process *θ*, such that for all *s* ∈ [0, *t*]

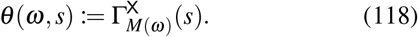

The manipulation protocols 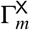 have the superscript X as the effect of the manipulation is assumed to solely affect certain reaction channels 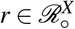 that exclusively change species X:

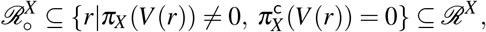

where 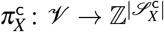 is the coordinate map to coordinates of 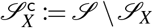, complementary to *S*_*X*_. This restriction ensures that the source protocol enters the network only through the designated channel encoding *X*. Since the manipulated reactions do not directly change any species outside *S*_*X*_, any influence of *θ* on the rest of the CRN, and in particular on the output *Y*, must be mediated by the dynamics of *X*. This keeps the interpretation of *X* as the channel input unambiguous. The restriction to reactions in 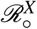 can be alleviated slightly to encompass reactions that are “upstream” of *X* in the sense that they influence the dynamics of *Y* only via the dynamics of *X*. This wider notion of encoding reactions was already used in Fig. 1. However, a formal disambiguation of such reactions for arbitrary CRNs requires further mathematical tools and is, therefore, beyond the scope of this work.

The propensity functions for reaction channels 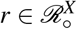 on [0, *t*] depend on *m* ∈ ℳ through the protocol 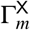, which we denote as 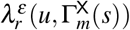. If the protocols are non-constant, then the propensities are explicitly time-dependent, given *m*. All other reactions 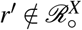 are assumed to have no explicit time-dependence for simplicity.

While *θ* can be regarded a (noiseless) source encoding, we refer to *X* as the (noisy) channel encoding with channel encoding propensities

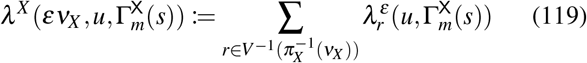

analogous to (105), where we do not distinguish between *m*-dependent and *m*-independent reactions on the r.h.s. for brevity. We call *Y* the channel output with output propensities

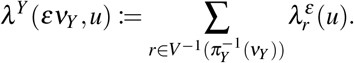

#### Specification of the causal communication system

To connect the CRN model to the abstract feedback-channel framework of Sec. IV E and IV F, we now introduce the intensity processes that depend only on the observed histories of the communication variables *M, θ, X*, and *Y*. The ℱ^*θXY*^ – intensities

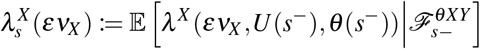

describe the actual stochastic channel encoding architecture and will be called *feedback channel encoding intensities*. The ℱ^*MXY*^-intensities

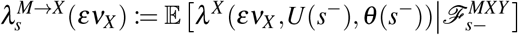

describe the same encoding dynamics, but viewed directly from the message level, and will be called *feedback source-channel encoding intensities*. Further, the ℱ^*XY*^-intensities

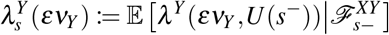

will be called the *causal channel output intensities*. As we discuss below, the intensities {*λ*^*Y*^ (*εν*_*Y*_)|*ε* ∈ {+,−*}, ν*_*Y*_ ∈ *V*_*Y*_*}* represent realizations of the causal communication channel itself. Finally, for message-to-output information measures of a channel with feedback and memory, we also require the ℱ^*MY*^ – intensities

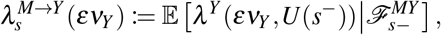

which we call *source-output intensities*.

The intensity processes *λ*^*Y*^ and *λ* ^*M*→*X*^ are the CRN counterpart of, respectively, the causal channel and the causal encoder introduced in Sec. IV E. As already discussed in Sec. VI, each of the above intensity processes determines a local Janossy density (a causally conditional path likelihood; cf. Eq. (115)), and hence a causally conditional path probability. In particular, the feedback source-channel encoding intensities induce the Janossy density 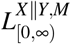 and therefore the causally conditional law ℙ^*X* ∥*Y,M*^ of *X* given *Y* and *M*. In this sense they specify a ℙ-regular version of a specific causal noisy encoder f from Def. 9. Likewise, the causal channel output intensities induce the causally conditional law ℙ^*Y* ∥*X*^ and thereby a ℙ-regular version of a specific causal channel W from Def. 4. [108]

The above encoding intensities also distinguish two encoding levels. The message *M* selects a deterministic protocol, represented by the noiseless source encoding process *θ*, whereas the actual channel encoding is the generally noisy process *X* generated by the reaction dynamics under that protocol. In the present setup, however, the source level can be equally well represented by *M* or *θ*, because *θ* is a deterministic function of *M*. Hence, conditioning on *M* or on *θ* yields the same predictive information for the encoding dynamics. Consequently, for the present noiseless source encoding via chemostat trajectories, the feedback source-channel encoding intensities coincide with the feedback channel encoding intensities as functions on (0, *t*] × Ω. For the same reason, the source-output intensities may equivalently be written by conditioning on *θ* instead of *M*. The formal statement is given next, while the precise measure-theoretic formulation and proof are deferred to Supp. Sec. S1.6 A (cf. Supp. Prop. S1.3).

##### Proposition 2

*If θ is a noiseless source encoding, defined as above by the deterministic protocol family* 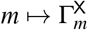, *then conditioning on M is equivalent to conditioning on θ for the corresponding intensity processes. In particular, for all ε* ∈ {+,−*} and ν*_*X*_ ∈ *V*_*X*_, *the feedback channel encoding intensities and the feedback source-channel encoding intensities agree almost everywhere on* (0, ∞) ×Ω, *i*.*e*.,

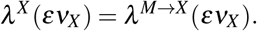

*Moreover, for all ε* ∈ {+, −} *and ν*_*Y*_ ∈ *V*_*Y*_,

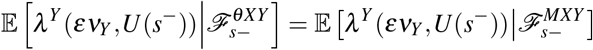

*and the source-output intensities satisfy*

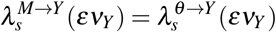

*almost everywhere on on* (0, ∞) ×Ω, *where*

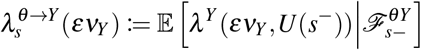

Lastly, let

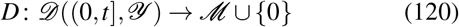

be a deterministic decoding function (analogous to (44)), which maps every output trajectory back to either of the messages in ℳ or declares an erasure, by mapping to zero. We may associate two distinct conceptions with this decoding function: (i) the *extrinsic decoder* is a (computationally) learned function by an external observer, based on experimental observations of *Y*_(0,*t*]_ given *M*; (ii) the *intrinsic decoder* refers to the conception that each *m* ∈ ℳ stands for a certain action that is noisily implemented by a biological cell, where 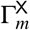 is the input that triggers this action and the output is a exactly the associated noisy downstream action. However, in the following we consider an *abstract decoder* which just determines the function space required for information theoretic analyses of chemical communication models. The (mean) probability of a decoding error be defined as

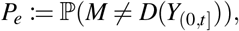

and is equivalent to Eq. (49) and reflects that common messages should be decodable more reliably than less common ones.

Another, more traditional “worst case” performance measure, is the maximum probability of error

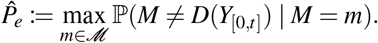

### B. Chemical channel and chemical channel codes

In the previous subsection we have assumed the perspective of a single chemical communication model defined on a fixed ambient probability space (Ω, ℱ, ℙ). While this is the natural way to introduce a physical model, we have motivated several times in Sec. IV that a stochastic kernel based description of a communication system adds clarity and is advantageous for the formulation of the capacity problem. In this subsection we hence link the physical model description of a chemical communication system to the stochastic kernel definitions of the causal channel W and the causal noisy feedback encoder f, respectively, Definition 4 and 9. This will also clarify, why we have introduced the notion of a noisy channel code in the first place and not just used the classical deterministic description, in Sec. IV B; because a general chemical feedback encoder is actually noisy. However, as in Eq. (120) the decoder will always assumed to be deterministic in our discussion.

#### Kernel-intensity representations of channel and encoder

As noted above, the channel law ℙ^*Y* ∥*X*^ is equivalently represented by the collection of predictable intensities *λ*^*Y*^ (*εν*_*Y*_)|*ε* ∈ {+,−*}, ν*_*Y*_ ∈ *V*_*Y*_ *}*, which is unique up to stochastic equivalence [53] (p. 31). This equivalence is readily apparent from the path likelihood (115). Whereas the mathematical literature on local Janossy densities formulates such likelihoods on the marked-jump space ∪_*n*=0_ ([0, *t*] × (*V*_*Y*_ ∪ − *V*_*Y*_))^*n*^ [98, Ch. 7.3], we work with an equivalent, and more intuitive, representation on the space of càdlàg trajectories *D* ([0, *t*], *X* × *Y*). To this end, we introduce a collection of non-negative functionals

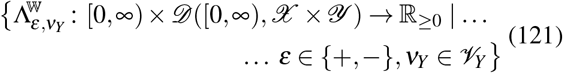

that adhere to a set of causality requirements, introduced below. With this collection we define the kernel-likelihoods

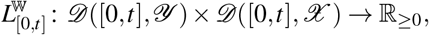

for all *t >* 0 such that

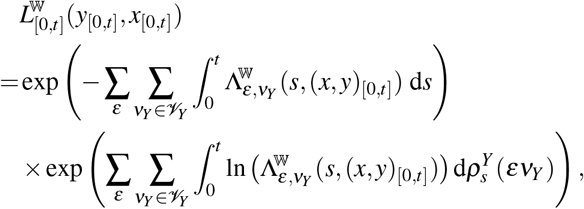

where the counting trajectories *ρ*^*Y*^ (*εν*_*Y*_) are fully determined by *y*_[0,*t*]_, and the integral is analogous to counting process integrals. As any density, a local Janossy density just represents a measure as an integral over a particular reference measure. Not wanting to go into detail, we just denote this path measure on *D* ((0, *t*], *Y*), B((0, *t*], *Y*)) by 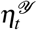, which essentially integrates and sums over all possible jump times and state changes up to time *t*. Consequently, we may define a stochastic kernel W that satisfies Definition 4 such that for all *t >* 0, all *x*_[0,*t*]_ ∈ *D* ([0, *t*], *X*), *y*_0_ ∈ *Y*, and for all *B* ∈ B((0, *t*], *Y*)

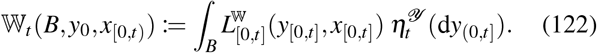

Note that the l.h.s. does not depend on *x*_*t*_ itself. Now, conversely, any causal channel W can be represented by a collection of intensities of the form (121). Under the conditions in Def. 15 below, which can be satisfied for the class of chemical communication models considered in this work, the identification is unique. This definition actually just lists a set of regularity conditions that any valid causal intensity process must satisfy. In preparation, we define the set of jump times J(*z*) := {*s* ∈ (0, ∞) : *z*(*s*)≠ *z*(*s*−)} for any càdlàg trajectory *z*.

**Definition 15**. *A collection of the form* (121) *(and equivalently its induced kernel* W*) is called* causal chemical channel *with initial state, if for all ε* ∈ {+,−*}, ν*_*Y*_ ∈ *V*_*Y*_ *exist a sequence of functions*

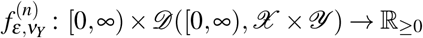

*such that*

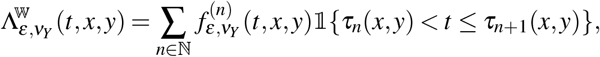

*Where*

i. *{τ*_*n*_*(x, y)* | *n* ∈ *N} = J(x, y) with τ*_*n*_(*x, y*) *< τ*_*n*+1_(*x, y*) *for all n* ∈ N *and all* (*x, y*),
ii. 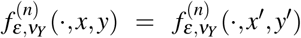 *for all* (*x*^′^, *y*^′^) *with* 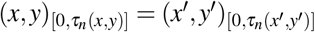, *and*
iii. 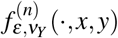 *is continuous on the interval* (*τ*_*n*_(*x, y*), ∞).

Condition (ii) just formalizes that the functions 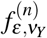 depend on the trajectories only via the past before the *n*-th jump. The uniqueness is justified by [53, p. 309, Thm. T34], which we adapted to the predictability conditions (i) and (ii), and the regenerative form of intensity processes for marked point processes whose interoccurrence times are continuous random variables with a probability density [53, pp. 59], which imply the continuity condition (iii). Actually, the functions 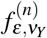 have the unique form

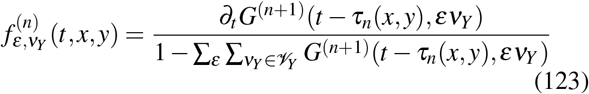

for *t* ∈ (*τ*_*n*_(*x, y*), ∞), where *t* ↦ *G*^(*n*+1)^(*t, εν*_*Y*_) is the joint distribution of the event that the (*n* +1)-th reaction will be of type *εν*_*Y*_ and satisfy *τ*_*n*+1_(*x, y*) − *τ*_*n*_(*x, y*) ≤ *t*, given *τ*_*n*_(*x, y*) and the process history 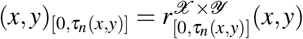; see [53, p. 60, Eq. (2.1)] for comparison.

In conclusion, we have three equivalent representations of a causal chemical channel: the stochastic kernel W, the kernel-likelihood *L*^W^, and the kernel-intensity collection, shortly denoted as Λ^W^. Comparing the kernel intensities with our natural discrete-time motivation of a causal channel with time structure, we identify them as the exact continuous-time jump-process analogue of Eq. (63), i.e., the single-instance output law W _*{n}*_ (*y*_*n*_ |*y*_[0,*n*)_, *x*_[0,*n*)_).

Analogously, we introduce a collection of non-negative functionals

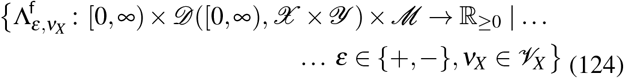

that should satisfy the same regularity conditions as in Def. 15, given any *m* ∈ ℳ. The collection (124) (and equivalently its induced kernel f) is then called *causal chemical encoder* with initial state. Here again, we have three equivalent representations of a causal chemical encoder: the stochastic kernel f, the kernel-likelihood *L*^f^, and the kernel-intensity collection, shortly denoted as Λ^f^.

#### Joint law of encoder and channel

Denoting 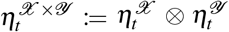 we may now obtain the finite-duration joint encoder-channel law (79) by integration of the product of local Janossy densities:

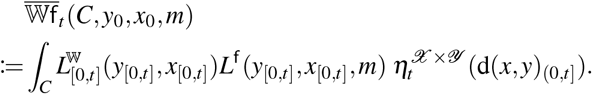

Note that the integrand is exactly analogous to the factorization of densities in natural discrete-time, presented in Eq. (78).

#### Induced channel and probability of error

The induced channel from message to message estimate is subsequently given by Eq. (81). This channel allows a purely kernel-based expression of the probability of error of an (|ℳ|, *t*) chemical code via Eq. (49).

#### Causal connection and ℙ-consistency

To connect these kernel-intensities with the intensity processes defined in Sec. VII A we now specify the notion of causal connection in Def. 10 to chemical communication systems.

##### Definition 16

(Causal connection via channel and encoder). *Let* (Ω, ℱ, ℙ) *be a probability space with filtration* ℱ = {ℱ_*t*_}_*t*≥0_ *and*

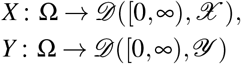

*be adapted stochastic processes (i*.*e*., *X*_[0,*t*]_ *and Y*_[0,*t*]_ *be* ℱ_*t*_*-measurable for all t* ∈ T_≥0_*). We say that X is causally connected to Y by the chemical channel* Λ^W^ *if for all ε* ∈{+,−} *and ν*_*Y*_ ∈ *V*_*Y*_, *the causal channel output intensities and the channel kernel-intensities agree almost everywhere on* (0, ∞) ×Ω, *i*.*e*.,

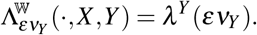

*Similarly, if M* : Ω → ℳ *is a* ℱ_0_*-measurable random variable, we say that Y, given M, is causally connected to X by the encoder* f_*t*_ *if for all ε* ∈ *{*+,*™} and ν*_*X*_ ∈ *V*_*X*_, *the source-channel encoding intensities and the encoder kernel-intensities agree almost everywhere on on* (0, *t*] ×Ω, *i*.*e*.,

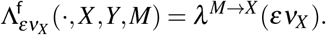

*A chemical communication system, given by p*_*M*_, *p*_0_, 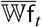, *and D is called* ℙ*-consistent if its joint distribution equals the joint distribution of* (*M, X,Y*, 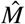) *on* (Ω, ℱ, ℙ).

By construction, chemical causal connection implies that the equivalent ℙ-regular causally conditional kernel satisfy ℙ^*Y* ∥*X*^ = W and ℙ^*X* ∥*Y,M*^ = f. Applying the noisy decoder description (46) for notational simplification, the joint distribution of the communication system for finite operation time *t >* 0 is given by

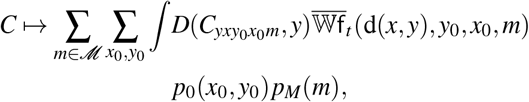

with 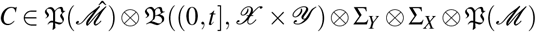,

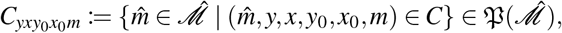

and integration over *D* ((0, *t*], *X* × *Y*).

#### Heuristic derivation of a channel kernel from a given CRN

So far, the discussion of intensity kernels has been rather abstract and one may wonder how it can be related to an actual CRN. The key advantage of stochastic kernels in the capacity problem is that they are defined for all possibly admissible càdlàg input trajectories *D* ([0, ∞), *X*) that can be realized by a chemical population process. What this facilitates is that the encoder is then also allowed assign positive likelihood arbitrarily to such admissible trajectories without risking that some of these trajectories cannot be interpreted by the channel law derived from a CRN that assigned the positive likelihood differently. In other words, the code design problem is then simply ill-defined on that subset of the design space that belonged to the ℙ^*X*^-null sets of the reference CRN. The stochastic kernel on the other hand is well-defined on all admissible input trajectories and thereby avoids the described problems.

So let us consider the model from Sec. VII A. Note again, that the stochastic kernel representing a causal channel does not depend on the probabilities with which input trajectories are generated. Further, it does also not depend on the initialization distribution *p*_0_. With this in mind, we conjecture that the following derivation of W or Λ^W^ is viable.

From the original chemical reaction network (ℛ, *S*) we construct an alternative intervened model, described on the probability space (Ω, ℱ, 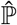), such that 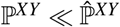. As a first step, we replace all propensities in ℛ^*X*^, i.e., those that only change species in *S*_*X*_ with unit propensities. That is, for all *s* ≥ 0 and *u* ∈ *U* set

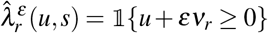

for all 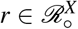. This transforms the reaction counting processes 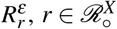, into conditional standard (unit-rate) Poisson processes. In fact, any Poisson-type counting process can be obtained from a standard Poisson process via thinning or a rescaling of time [53, 57, 105]. An alternative notion of local Janossy densities also uses the standard Poisson process as a reference measure to derive the respective densities [98]. Thus, the conditional standard Poisson process assigns positive likelihood to all possible trajectories that could be realized by a Poisson-type counting process 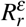, where *u* + *εν*_*r*_ ≥ 0 acts as a regularity condition ensuring *U* (*t*) ≥ 0 for all *t >* 0. In this way, we have extended the attainable dynamics of the reactions 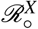 as much as possible, and the intervened model no longer depends on the source encoding protocols. However, the intervened model of *X* is still not able to realize all trajectories within *D* ([0, ∞), *X*). Changing any of the reactions 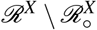 could potentially change the causally conditional law of *Y* given *X*, which is why we cannot modify them directly. However, we may instead introduce auxiliary reactions 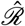 with change vectors satisfying 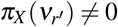 and 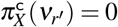 as well as 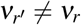 for all 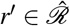 and all 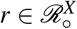. Assigned propensities are again of the form

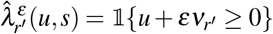

for all 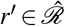. To account for all possible changes in the càdlàg space, we could require that

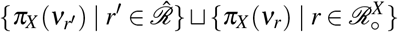

represent all possible forward reactions in 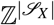. Indeed, since *X* is countably infinite for a general population model, 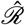 would then also be countably infinite. So, while we have formulated kernels above on the full space *D* ([0, ∞), *X*), we observe that this full space will never be practically realizable by any chemical reaction network. Which reactions should actually be included depends on the question, which dynamics the noisy encoders f over which the channel coding problem optimizes, should be able to realize. However, this question is only of practical interest when the stochastic kernel of interest is derived; it can naturally also be included as a restriction on the class of encoders f, which is why we proceed without further specification.

Finally, the intervened model also needs to assign positive probability to any state (*x*_0_, *y*_0_) if *p*_0_ is part of the design degrees of freedom. In fact, *λ*^*Y*^ generally has a hidden dependence on the initial distribution *p*(*u*, 0) of the full CRN. Consequently, we define the initial distribution 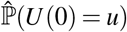 of the intervened model as a product distribution of

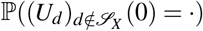

and a product Poisson distribution on *X* × *Y*. The kernel intensities are then derived, e.g., in the regenerative form with (123) for the new input process with state equation

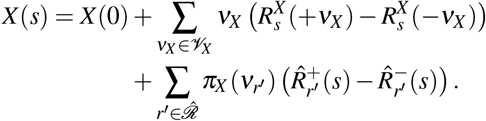

What such a kernel derivation should accomplish is first, to facilitate the formulation of a chemical channel for an existing network structure, and second, to guarantee that there indeed exists a chemical reaction network associated with a specific chemical channel. Without the latter, the channel coding problem was ill-defined.

#### Kernel-intensity representation of the feedback channel encoder

Now, let us reintroduce the noiseless chemical source encoding protocols. Assume the source encoding is performed with *N* ∈ N chemostat species. Then let 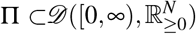 be a measurable subset that contains only bounded and piecewise-differentiable trajectories. We refer to Π as the *admissible protocol space*. Π_*t*_ denotes the restriction to 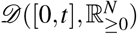. In addition the the boundedness and differentiability, this space Π may be defined via a a family cost functionals 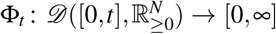 with *t >* 0 such that all Γ_*t*_ ∈ Π_*t*_ must satisfy Φ_*t*_ (Γ_*t*_) ≤ 1, analogous to (52).

##### Definition 17

*Let* Π_*t*_ *be an admissible protocol space. A noiseless chemical* (|ℳ|, *t*)*-source code is an (injective) mapping*

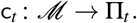

The protocol-valued random variable of such a source code is given by *θ*_[0,*t*]_ = c_*t*_(*M*), where we typically require c_*t*_ to be injective such that a bijection between the two random variables exists.

Again, we introduce a collection of non-negative functionals

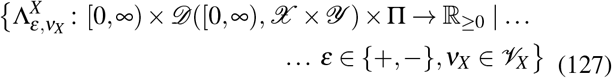

that should satisfy the similar regularity conditions as in Def. 15. Only condition *(ii)* is replaced with *(ii’)* For all *t >* 0 it holds 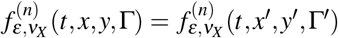 for all (*x*^′^, *y*^′^) with 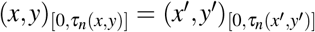 and all Γ′ with 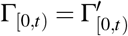.

We refer to this collection as *causal chemical feedback channel encoder* with initial state. This notion of the encoder achieves a separation between source encoding through signaling chemostat protocols and feedback channel encoding via the CRN architecture. It is again equivalent to a local Janossy density and a stochastic kernel.

##### Definition 18

(Causal connection via channel-encoder). *Let* (Ω, ℱ, ℙ) *be a probability space with filtration* ℱ = {ℱ_*t*_}_≥ *t* 0_ *and X,Y be* ℱ*-adapted processes*.

*If θ* : Ω → Γ *is a* ℱ_0_*-measurable random variable, we say that Y, given θ, is causally connected to X by the feedback-channel encoder* Λ^*X*^ *if for all ε* ∈ *{*+,−} *and ν*_*X*_ ∈ *V*_*X*_, *the channel encoding intensities and the channel-encoder kernel-intensities agree almost everywhere on on* (0, ∞) ×Ω, *i*.*e*.,

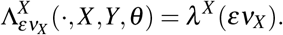

With all these kernel notions at hand, we need only one more ingredient to formulate a channel coding theorem: the right information measures and their relation to intensity processes and kernel-intensities.

### C. Information measures for chemical communication

In general, the relevant mutual information for the channel coding theorem of a continuous-time chemical communication channel with memory and use of feedback is I(*M*;*Y*_(0,*t*]_). To account for the initialization, note that

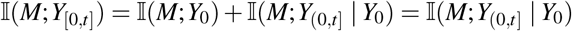

by the chain rule of information and 0 ≤ I(*M*;*Y*_0_) ≤ I(*M*;*U* (0)) = 0 by construction. With the intensity processes defined thus far, we have

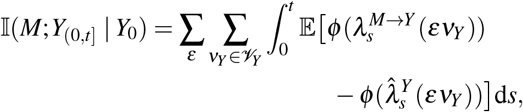

which corresponds to a use of the information channel with decoder side information of the initial state *y*_0_. However, since for any *t >* 0 the realizations of *Y*_[0,*t*]_ are càdlàg trajectories, the realized state *y*_0_ can be reconstructed from *y*_(0,*δ*]_ for any *δ >* 0 by taking the limit from the right. This implies that both 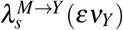 and 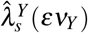 are equal to the intensity processes not conditioned on *y*_0_ for all *s >* 0, which makes the entire intensity processes stochastically equivalent. Hence, the side information is actually irrelevant for the decoder and we have

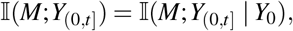

which by the above discussion directly carries over to the mutual information density, such that

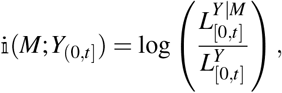

where 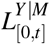 is the Janossy density associated with the source-output intensities *λ* ^*M*→*Y*^ and 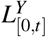 is the marginal Janossy density associated with the output intensities 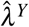.

Consistently with Proposition 2 we have

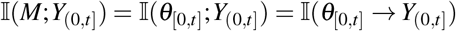

for a noiseless source-encoding *θ*_[0,*t*]_.

At the density level, the directed information from the noiseless source encoding *θ* to the channel output *Y* on (0, *t*] is represented by the log-likelihood ratio

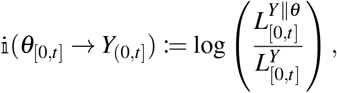

where 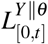 is the Janossy density associated with the source output intensities *λ*^*θ* →*Y*^. Writing out the likelihoods, we obtain

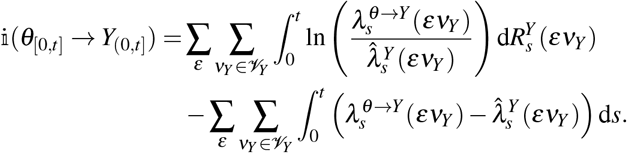

If the directed information is finite, then it holds

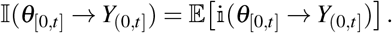

#### Causal communication systems in the sense of Massey

A useful classification of feedback communication systems goes back to Massey [11]. In the present setting, the relevant question is whether knowledge of the entire source encoding over the transmission interval provides any additional predictive information about the channel output once the channel input and output histories are already known. Since the source encoding is fixed before transmission starts, this criterion is naturally formulated by adjoining the full *σ*-field 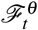 to the past history 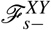. This motivates the following definition.

##### Definition 19

*For a* (|ℳ|, *t*)*-source code with message M, let θ*_[0,*t*]_ *be a (potentially noisy) source encoding and let Y*_(0,*t*]_ *be the channel output. A chemical communication model* (*M, θ,U, S*_*X*_, *S*_*Y*_) *with designated channel encoding X*_[0,*t*]_ *is called* causal *if, for all ε* ∈ {+, −} *and all ν*_*Y*_ ∈ *V*_*Y*_, *the process*

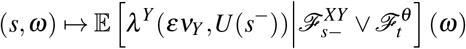

*is an* 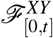 *-intensity*.

In other words, a causal communication system is defined by the property that, even when the complete source encoding over [0, *t*] is known, it does not alter the channel output intensities once the history of (*X,Y*) is given. Consequently, the output behavior is entirely governed by the specified channel encoding *X*, analogous to Massey’s definition in discrete time. We emphasize that this definition encompasses possible extensions to noisy source encodings *θ*.

Definition 19 directly implies that the causally conditioned output law given (*X, θ*) agrees with the output law given *X* alone, since both are generated by the same output intensities. Equivalently, the corresponding causally conditioned Janossy densities coincide. As a consequence, the log-likelihood ratio defining the causally conditioned directed information vanishes, and therefore

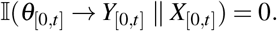

If the source encoding is noiseless, then the protocol process *θ* is a deterministic function of the message *M*, so that 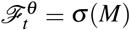. In that case,

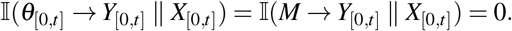

In words, the output process, causally given the channel encoding, does not causally depend on the message or the source encoding.

#### Relations between information measures for causal communication systems

In analogy with Massey’s causal communication framework, we obtain the following relations between information measures for chemical communication systems in which the source encoding *θ* may itself be noisy. We assume that the source-encoding process satisfies the regularity conditions

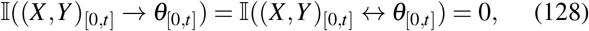

that is, the source encoder is causally independent of the rest of the system and there is no instantaneous information exchange between them. Although we do not prove these conditions here in full generality, they arise naturally from our construction, where *θ* only enters the CRN *U* as a modulation of propensity functions. For the noiseless source encoder constructed above, they hold automatically because *θ*_[0,*t*]_ is in bijection with *M*. Moreover, since

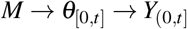

forms a Markov chain also for potentially noisy source encodings, the data processing inequality yields

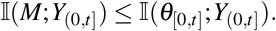

Further, recall that under the two natural assumptions on the causal communication model

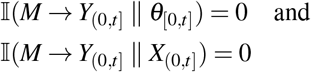

we have also shown in Lemma 1 that

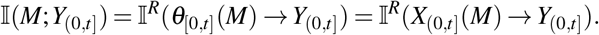

##### Proposition 3

*Let* (*θ*_[0,*t*]_, *X*_[0,*t*]_,*Y*_[0,*t*]_) *be a causal communication system for a (potentially noisy) source code θ*_[0,*t*]_ *satisfying* (128). *Then*

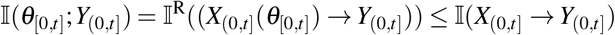

*with equality if and only if* I(*X*_[0,*t*]_ → *Y*_[0,*t*]_ ∥ *θ*_[0,*t*]_) = 0.

*Proof*. The regularity conditions and the information conservation law (97) imply

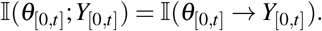

Since trajectories are càdlàg, we may always ignore the initial states the processes *X* and *Y*. Then the chain rule (99) yields

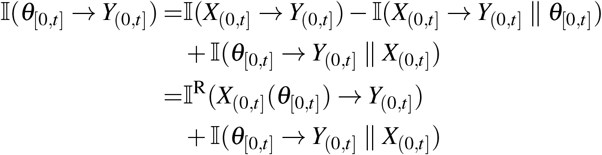

The equality is the established since Definition 19 implies

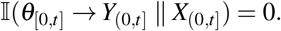

The inequality follows with Lemma 1.

In that, I(*X*_(0,*t*]_ → *Y*_(0,*t*]_ ∥ *θ*_[0,*t*]_) quantifies the surplus of causal information provided by *X* to *Y* beyond the causal information already provided by *θ*. We can think of this as the amount of channel encoding noise that is causally transmitted to *Y*. Since the ℱ^*θY*^-intensities *λ*^*θ* →*Y*^ usually differ from the ℱ^*θXY*^-intensities

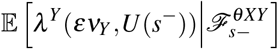

for any *ν*_*Y*_ ∈ *V*_*Y*_, *ε* ∈ {+,−*}*, this surplus information typically has a positive value. Such positive surplus information can, for example, be verified for the small models in Sec. VIII D under the additional assumption of two distinct constant chemostat concentration values as 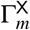-trajectories, although we do not carry out that verification explicitly here.

### D. The noisy channel coding problem in chemical communication

#### Chemical communication channel and source-side design variables

We define a *chemical communication channel* as a triple

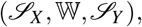

where *S*_*X*_ and *S*_*Y*_ are designated disjoint input and output species, and W is a fixed causal channel law from input trajectories to output trajectories. More precisely, we require that there exists at least one CRN *U* with *S*_*X*_, *S*_*Y*_ ⊆ *S* and a probability space (Ω, ℱ, ⊆ ℙ) such that the corresponding processes *X* and *Y* are causally connected by W in the sense of Def. 10. Implicitly, this fixes the population state spaces *X* and *Y* of *S*_*X*_ and *S*_*Y*_, since W is defined on trajectories over these spaces.

At this level, the chemical realization of W is neither unique nor fixed. The only requirement is chemical realizability: at least one CRN must exist that implements the channel law between the designated input and output trajectory spaces. To integrate source-side encoding protocols into this fixed-channel description in an unambiguous way, we impose two additional assumptions.

##### Assumption 1

*The species labels in S*_*X*_ *and S*_*Y*_ *have fixed molecular identities in the physical sense, that is, fixed atomic structure*.

This assumption fixes the intrinsic reaction-level structure of the open subnetworks associated with *S*_*X*_ and *S*_*Y*_, independent of the rest of the network. In particular, it fixes the stoichiometric balance equations (cf. Eq. (8)) of all elementary reactions 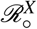 that exclusively change species in *S*_*X*_. Recall that the message-dependent source protocols 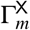 are assumed to influence the full CRN only through reactions in 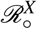.

In turn, this also fixes a finite set of possible physical identities of chemostat species involved in 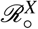. Some of these chemostats may correspond to ubiquitous chemical fuels such as ATP, and may also participate in reactions outside 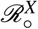 in a full CRN realization. We therefore single out a subset of chemostat species that play the role of signaling molecules. This leads to the following second assumption.

##### Assumption 2

*Under Assumption 1, there exists a subset* ℒ *of chemostat species involved in the reactions* 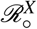 *such that there is at least one CRN U implementing the channel* (*S*_*X*_, W, *S*_*Y*_) *for all allowed time-dependent concentration variations of the chemostats in* ℒ.

Assumptions 1 and 2 together use both the species *S*_*X*_, *S*_*Y*_ and the channel law W to fix the molecular identities of signaling chemostats ℒ and, at the same time, to restrict the class of CRNs that may realize the channel. In particular, only those CRNs *U* are admitted in which these signaling chemostats act through the designated reaction family 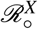.

Under these assumptions, the number |ℒ | ∈ N_*>*0_ is fixed, and every source protocol 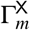 of duration *t >* 0 is represented by a bounded, piecewise-differentiable càdlàg trajectory in 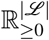. Without imposing further restrictions, we denote the resulting joint encoder-channel system by

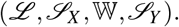

The assumptions above are not unique, and their main role is to isolate a mathematically manageable and physically reasonable notion of encoder architectures. Before formalizing this further, we distinguish two natural coding perspectives.

#### Distinct encoder design perspectives

The channel capacity is a property of a causal chemical channel (*S*_*X*_, W, *S*_*Y*_) with fixed input and output species identities and a fixed channel law. Thus, the coding problem is formulated for a prescribed causal law from *X* trajectories to *Y* trajectories, while the design freedom lies in how the channel input *X* is generated and how the output is decoded. In biochemical systems, this does not necessarily mean that the surrounding CRN realization is fixed in every detail. In particular, one may allow modifications of the dynamics that generate *X*, or more generally of parts of the surrounding network, provided that the induced forward channel law W from *X* to *Y* remains unchanged. What is excluded, however, is any intervention that results in W ≠ ℙ^*Y* ∥*X*^. Allowing such changes would turn the problem from channel coding for a fixed channel into a joint code–channel co-design problem. We deliberately discuss the former, since the assumption of a fixed channel law is the defining structural feature of the traditional noisy channel coding problem and therefore provides the clearest reference point for extending that framework to biochemical information processing in general.

Within the fixed-channel setting, two natural coding perspectives may still be distinguished. In the first, one fixes not only the channel law W, but also a specific open-CRN realization (*S*, ℛ), including stoichiometry and propensity functions. Then the only design freedom during channel operation lies in the choice of signaling chemostat trajectories of ℒ. In the second, one fixes only the specification (ℒ, *S*_*X*_, W, *S*_*Y*_), while allowing different surrounding network realizations that implement the same effective channel law. In that case, interin mediary species, reactions, and feedback pathways may vary, and with them the encoder architecture and specifications that generate the dynamics of *X*, provided that the induced forward channel W from *X* to *Y* remains unchanged. Depending on the modeling scenario, code design may therefore also include variations in reaction propensity functions and other CRN-level modifications, that leave the effective channel law W unchanged while altering how feedback from the receiving species influences the encoding species.

In this work, we focus on the first perspective. The second perspective will only briefly be discussed below without going into detail. What we do not attempt here is an explicit treatment of CRN modifications at the level of concrete reaction-network changes. Doing so would require additional mathematical tools to determine which changes of species, reactions, feedback pathways, or propensity functions leave the induced channel law invariant and which do not. That level of structural analysis is therefore left outside the scope of the present work.

#### Encoder architecture of the fixed-CRN perspective

In this scenario, the architecture of the feedback encoder responsible for producing the stochastic channel encoding is determined by the designated CRN. The designated encoding species *X* primarily structure the flow of information through the prescribed network (*S*, ℛ): the message acts first through the deterministic signaling protocols, these protocols drive the dynamics of *X*, and only through *X* is information transmitted to the output species *Y*. Hence, the stochastic part of the channel encoding is absorbed into the fixed encoder architecture, while the remaining design problem is reduced to choosing the signaling chemostat trajectories and the decoder of the form (46).

In the notation of Sec. IV F, the fixed CRN realization determines the encoder architecture ℍ. Accordingly, we identify the corresponding space of deterministic encoding specification as the set of admissible signaling chemostat trajectories (cf. Def. 17)

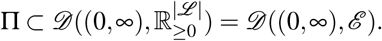

The deterministic encoding specification is a random variable

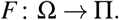

Thus, in the present setting, *F* represents a generalized version of the protocol process *θ* which is not restricted to a finite set of protocols, and the auxiliary priors *µ*^*F*^ are probability measures on Π.

The encoder architecture ℍ is then directly induced by the chemical feedback-channel encoder kernel-intensity collection Λ^*X*^ in (127), equivalently through its associated local Janossy density *L*^ℍ^. Thus, for every *t >* 0, *x*_0_ ∈ *X, y*_[0,*t*)_ ∈ *D* ([0, *t*), *Y*), Γ ∈ Π, and *A* ∈ B((0, *t*], *X*), we define

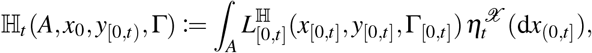

in direct analogy with the passage from Λ^W^ to W in Eq. (122). Hence, in the fixed-CRN perspective, ℍ is not an additional object beyond the chemical model itself, but simply the kernel-level representation of the fixed CRN encoder dynamics. In particular, for any ℙ-consistent realization,

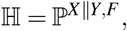

where in the present noiseless source-encoding setting *F* is represented by the protocol process *θ*.

In conclusion, the general coding theorem has exactly the form of Eq. (93), i.e.,

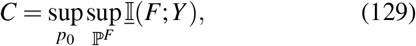

and the mutual information density for finite transmission duration *t >* 0 is given by

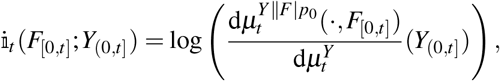

where the initialization *p*_0_ remains as an additional parameter.

#### Channel coding with variable-CRN structure

In the broader fixed-channel perspective, one no longer fixes a single CRN realization and hence not a single encoder architecture ℍ. Instead, one allows an admissible class of encoder architectures

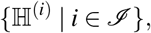

where the index *i* labels distinct CRN realizations that all induce the same forward channel law W from *X* to *Y*. Then, the reaction structure encoded by the index *i* ∈ *I* fixes a set of stoichiometric balance equations and the functional form of the propensity functions, which then do not depend on the message but rather describe a design choice of the overall communication system.

For any fixed architecture ℍ^(*i*)^, however, the deterministic encoding specification from Sec. IV F may still have a non-trivial time-independent component. Concretely, any source code for the architecture *i* ∈ *I* may assign to each message an element

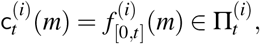

where 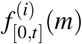 is the signaling protocol trajectory.

Accordingly, in the variable-CRN perspective the abstract pair (*F*, ℍ) is generalized in two ways: first, one works with an admissible class of architectures {ℍ(*i*)}_*i*∈*ℐ*_ rather than a single fixed ℍ; second, for each chosen architecture ℍ^(*i*)^, the deterministic encoding specification takes values in a possibly different space Π^(*i*)^. The corresponding auxiliary priors *µ*^*F*|*i*^ are then probability measures on these architecture-dependent specification spaces. The general coding theorem is then of the form

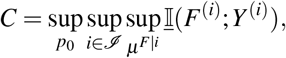

where *Y* ^(*i*)^ is also carries an encoder architecture index since any 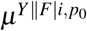 is constructed with a specific encoder kernel ℍ^(*i*)^.

## VIII. BIOCHEMICAL COMMUNICATION UNDER ENERGY CONSTRAINTS

### A. Energy constraints from stochstic thermodynamics

Let us now introduce a physically principled constraint on the search space of *X*_[0,*t*]_ and *F*_[0,*t*]_ (or, equivalently, *θ*_[0,*t*]_). In Sec. IV B we have discussed several possible types of constraints. Here, we will discuss constraints of the form of Eq. (56). We will thereby have a look at three different random cost functions 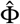: (i) total entropy production *h*^tot^, (ii) dissipated heat *q*(*t*), and (iii) work *w*(*t*) performed on the system.

#### Total entropy production rate constraint

For each *f* ∈ Π_*t*_ we consider the total entropy 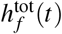 produced by the trajectory *U*_[0,*t*]_, given the external manipulation *f* represented by a signaling chemostat protocol. Let *ε*_0_ ≥ 0 be the upper bound on the time-averaged total entropy production, and define the cost function (56) as

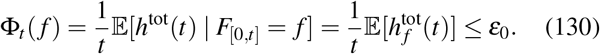

Such a constraint is also called input-output cost function [81], because it considers the dissipative cost of the whole CRN *U* and not only a cost that is directly formulated on the generation of *f*, or the cost of channel encoding into *X*_(0,*t*]_.

As discussed in Supp. Sec. S2.6, the cost function (130) can be identified with the KL divergence between the forward path of the reaction-resolved CRN and the time-reversed path of this CRN under a time-reversed protocol; cf. Supp. Eq. S2.18. It can be expressed by Eq. (36) as

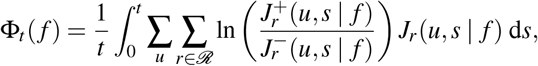

where the time-dependence is assumed to be generated by the manipulation protocol *f*. Using the decomposition *h*^tot^(*t*) = *h*^U^(*t*) + *q*^mes^(*t*), another equivalent expression is

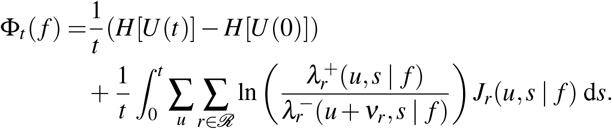

We remark that the total entropy production rate constraint requires a full-reaction resolved CRN model. In contrast to the relevant information measures of the coding theorems, which require filtered, history dependent intensity processes that can be obtained from a full network through marginalization, the EPR directly uses propensity functions an probabilities *p*(*u, s*) of the full CRN model.

#### Total EPR constraint and stationarity

Assume that Π ⊂ *D* ([0, ∞), ℰ) is a protocol space that is closed under time shifts, i.e.,

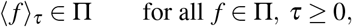

where ⟨*f* ⟩_*τ*_ denotes the trajectory shifted by *τ* in time such that ⟨*f*⟩ _*τ*_ (*t*) = *f* (*t* + *τ*) [78]. An auxiliary prior *µ*^*F*^ on Π is called *time-stationary* if

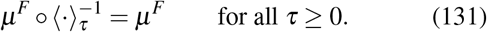

That is, the initial segment [0, *τ*) can be removed for arbitrary *τ* ≥ 0 without changing either the admissible protocol space or the auxiliary prior. Then, for each *t >* 0, the finite-duration law

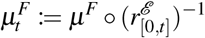

is obtained by restriction through the known time-restriction map 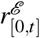, and the resulting family 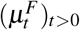 is shift-consistent in the sense that the law of every length-*t* observation window is independent of its starting time.

The appeal of such time-stationary priors can be understood heuristically from both the thermodynamic and the coding-theoretic side. If the protocol process *F* is viewed as part of a larger stochastic description of the CRN that evolves under a stationary distribution, then finite observation windows have shift-invariant statistics. On the thermodynamic side, this suggests that the time-averaged cost Φ_*t*_ from (130) may approach a stable long-time rate [109, Thm. 2.2.4.]. A related idea appears on the coding-theoretic side: the general capacity formula is expressed in terms of the inf-information rate I(*F*;*Y*), whereas the more familiar Shannon-type expression involves the ordinary information rate I_∞_(*F*;*Y*). Passing from the former to the latter requires information stability, meaning that the information density per unit time must admit a well-behaved long-time limit. Stationarity alone does not ensure this, but it provides a natural starting point under which such a simplification may be expected once additional regularity assumptions are imposed.

With the finite-time cost functional Φ_*t*_ already defined in (130), one may now impose thermodynamic constraints of different strength, in direct analogy with Sec. IV B. The strongest option is the protocol-space analogue of the hard signal constraint (52), namely the restriction of the admissible protocol set to

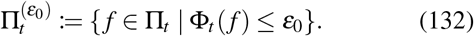

If instead one starts from a deterministic source code c_*t*_ : ℳ → Π_*t*_, *m* ↦ *f*_*m*_, then the direct protocol-level analogue of the message-wise expected cost constraint (53) is

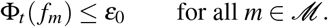

Finally, if 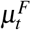 is an auxiliary prior on protocol space, one may impose the weaker expected constraint

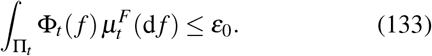

This constraint is not merely a technical relaxation. By the definition of Φ_*t*_ in (130), it is exactly the expected time-averaged total entropy production of the full communication system,

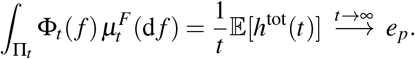

Thus, (133) directly bounds the mean dissipative cost of communication under the chosen protocol prior. In this sense, it is the natural thermodynamic counterpart of (54).

At the same time, (133) is weaker than the hard restriction (132). It controls only the mean entropy production across the ensemble of protocol realizations selected by 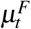, and therefore still allows individual protocols with Φ_*t*_(*f*) *> ε*_0_ to occur with positive probability. For this reason, imposing an expected thermodynamic constraint is generally not the same as restricting the admissible protocol space itself; the two formulations coincide only under additional assumptions on the communication model such as a memoryless channel; cf. [92, Remark 3.6.1].

#### Work and heat dissipation constraints

Similar to (130) we would like to formulate a work and heat constraint of the form

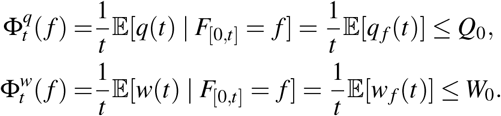

However, such constraints are not directly accessible from the kinetic model. Recall that *q*(*t*) = *q*^mes^(*t*) − *h*^i^(*t*), where *q*^mes^(*t*) is accessible for the open CRN model via Eq. (32), but *h*^i^(*t*) accounts for the entropic changes in internal degrees of freedom which are not accessible from the population model. What would be required to derive *h*^i^(*t*) is a microscopic Hamiltonian H. Similarly, recall that *w*(*t*) = *w*^Γ^(*t*) + *w*^c^(*t*), where

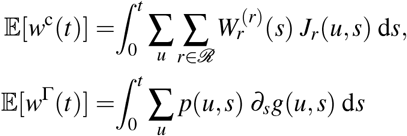

with 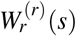 given in terms of time-dependent chemical potentials of the chemostats (10). What is needed to derive *w*(*t*) are therefore the time-dependent chemical potentials *µ*_*l*_(*s*) as well as the knowledge of the derivative of the mesoscopic Gibbs free energy under the given protocol.

Consequently, both the total entropy production *h*^tot^ and mesoscopic heat *q*^mes^(*t*) are the only directly accessible options for thermodynamically motivated energy constraints.

#### Remarks thermodynamic cost constraints

Note that the constraints discussed thus far are joint channel-feedback-encoding and transmission cost constraints. They measure the energetic cost of stochastically encoding and transmitting any *f* via the CRN. Hence, they do not assign a direct cost for the generation of *f* itself, as in the classical notions of cost constraints, where the transmission does not carry any energetic cost. Classically, transmission rather leads to a decay in signal strength and clarity, and the energetic cost to generate a strong physical signal *f* is typically expended at the encoder. Of course, the given cost functions can always be complemented by further cost functions on the protocols, such as a peak constraint of the form *f* ≤ *φ*_0_ with *φ*_0_ ≥ 0.

### B. Channel capacity and Fano-type converse theorem

#### Capacity-cost functions

As we have motivated for the fixed-CRN channel coding paradigm, the general channel coding theorem has the form (129), i.e.,

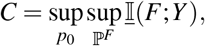

which, under the additional assumption of information stability, reduces to the Shannon-type expression

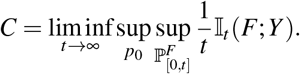

Further, irrespective of information stability, it hold that [93, Thm. 8)]

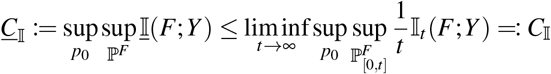

with equality if and only if information stability is satisfied. Here, we have introduced a separate notation for the infmutual information rate capacity *C*_I_ and the Shannon-type information capacity *C*_I_, to distinguish these different information measure-based quantities from the operational capacity *C*, which follows Definition 7.

Evaluating these different capacity notions under the discussed thermodynamic constraints, i.e., considering only those probability measures with 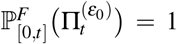, yields capacity-cost functions

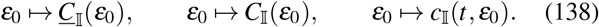

where we exemplarily use the total entropy production rate constraint (130) with upper bound *ε*_0_. Note that only the Shannon-type capacity has a time-wise notion

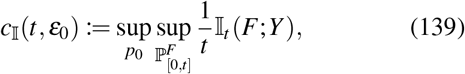

which has, however, no relevance in any coding theorem, since these are always asymptotic statements.

#### Fano-type converse theorem

For the given communication channel we can also provide a converse (channel coding) theorem [46, 110], which is valid for any of the discussed energy constraints. Importantly, it is also valid for information unstable channels, since it only provides an upper bound on the operational capacity that is not necessarily tight.

##### Lemma 2

(Fano-type inequality). *Consider an* (|ℳ|, *t*)*-code for the channel* (ℒ, *S*_*X*_, W, *S*_*Y*_) *specified in Sec. VII D. Let M* : Ω → ℳ *be the message with probability mass function p*_*M*_. *If the code exhibits the code rate R, then*

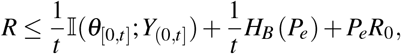

*where H*_*B*_(*p*) = −*p* log(*p*) − (1 −*p*) log(1 −*p*) *denotes the binary entropy function (for an arbitrary log-base)*.

We remark that in this Lemma, *θ*_[0,*t*]_ represents the message process (118), which can assume |ℳ| different protocol trajectories. Hence it is not the auxiliary encoding specification process *F*_[0,*t*]_. For the following theorem we exploit that any *θ*_[0,*t*]_, that does not exeed the energy dissipation bound, must still satisfy

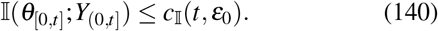

##### Theorem 3

*Let R be an achievable rate of the channel* (ℒ, *S*_*X*_, W, *S*_*Y*_) *under a given set of energy constraints on admissible codes. For simplicity, we exemplarily use the total entropy production rate constraint with upper bound ε*_0_. *Then it holds*

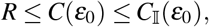

*where C*(*ε*_0_) *denotes the operational capacity under the dissipation constraint*.

Proofs for both, the Lemma and the Theorem, is provided in Supp. Sec. S1.6 A. Since the proof does actually only rely on the inequality (140), it is also naturally valid for other capacity-cost functions that do not need to have a thermodynamic motivation.

Lastly, although we formulate these optimization problems under thermodynamic constraints, they should often be complemented by kinetic constraints. For instance, at thermodynamic equilibrium, where the rate constants satisfy (5), one could in principle let all rate constants diverge while maintaining fixed propensity ratios. This, however, is clearly unphysical. Additional assumptions, such as lower bounds on mean sojourn times [38] or explicit models of the reaction constants [59], might be necessary to ensure physical plausibility. In stochastic thermodynamics, the interplay between thermodynamic and kinetic constraints has been explored in the context of thermodynamic and kinetic uncertainty relations [111].

### C. Energy-per-bit perspective

A dual perspective to information capacity is the energy-per-bit-rate function [81]. Define the *average energy per information unit* (given the code rate *R* in an arbitrary log-base) as

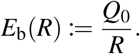

Here, we use the heat dissipation constraint with bound *Q*_0_ instead of the total entropy production constraint; otherwise we would have defined the *average entropy production per information unit*.

Since the capacity-cost functions (138) are monotone increasing, the *minimum-energy-per-bit-rate function* can be defined as their generalized inverse:

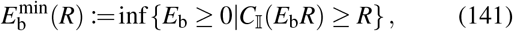

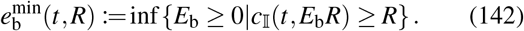

Given Theorem 3, 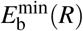 is actually a lower bound on the minimum amount of heat that must be dissipated per unit of information sent to reliably communicate at code rate *R* over a given channel. This bound is conjectured to be tight for information stable channels. The generalized inverse (141) can also be applied directly to *C*(*Q*_0_) and *C*_I_. The following Lemma makes the asymptotic version numerically more accessible

#### Lemma 3

*It holds that*

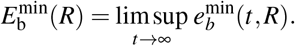

A proof is provided in Supp. Sec. S1.6 C.

Similarly, we can define the minimum average heat dissipation required for a minimum information rate *R*:

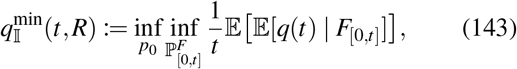

subject to the constraint I(*F*_[0,*t*]_;*Y*_(0,*t*]_) ≥ *Rt*. It’s asymptotic version is

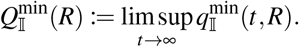

The two presented notions of minimum energy dissipation per information unit are shown to be equivalent, proving that the energy minimization per reliably sent information unit and the capacity cost-function under a maximum energy dissipation constraint are in fact dual problems.

#### Proposition 4

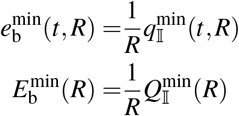

As before, the definitions and equivalence results work analogously with total entropy production, etc. A proof is provided in Supp. Sec. S1.6 C.

### D. Case studies: Accurate energy-accounting and information-dissipation trade-off in small promoter models

We now discuss thermodynamic and information-theoretic modeling for two partly analytically tractable promoter-transcription models. We use these models primarily to illustrate the thermodynamic and information-theoretic modeling pitfalls mentioned above. We and others [40] have derived tractable expressions for the mutual information of the considered model class. Promoter models also serve as prominent examples in biochemical information-theoretic studies [5, 6, 112], further enhancing the appeal of this choice. In particular, we realize that several properties of the described chemical communication model must be abandoned in this minimal model. Nonetheless, the following Shannon-theoretic discussion can be regarded as a conceptual link between the previously discussed biochemical information theoretic studies and our suggested biochemical communication framework.

In each model the promoter state trajectory acts as the channel input *F* and the mRNA transcription record acts as the output *Y*. The promoter state thereby directly modulates the propensity function of the transcription process, without a stochastic channel encoding population X in between. Hence, although we will model the promoter as a stationary Markov jump process, it is not a chemical encoding in the sense discussed so far. Instead the promoter process is understood in the sense of a stationary auxiliary prior distribution, as discussed in Eq. (131). Our task is to maximize the mutual information rate 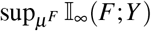 subject to an upper bound on the promoter’s intrinsic entropy-production rate *e*_*p*_. Transcription events are excluded from the thermodynamic budget because the transcription events are not modeled thermodynamically consistent for simplicity and tractability. Hence, in contrast to the EPR constraint in Eq. (130), we formulate the cost function directly as

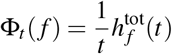

for all *t >* 0, where 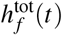 is the entropy production of *F*_[0,*t*]_ and not *U*_[0,*t*]_ | *F*_[0,*t*]_. Since *µ*^*F*^ is chosen stationary we apply the weaker expected constraint of the form (133), i.e.,

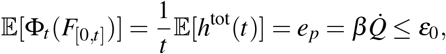

where the r.h.s is independent of *t* and equal to the heat dissipation rate by the stationarity assumption on *F*, [109, Thm. 2.2.4.] and relation (38). The resulting constrained problems are solved with fmincon (MATLAB R2022a), providing quantitative examples of channel capacity for small promoter-transcription models under thermodynamic constraints.

Consider a two-state Markov jump process with states OFF and ON, which modulate transcriptional activity by switching between the transcription rates 0 and *k*_tx_. For simplicity, we set *k*_tx_ = 1 in our analysis, thereby fixing the system’s time scale. The state transition diagram of this model, along with transition rates and thermodynamic assumptions, is shown in Fig. 6. The model features a pair of parallel reaction channels: the first channel involves ATP hydrolysis with 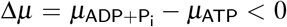 and is coupled to corresponding chemostats, while the second reaction channel is thermally “driven” and not coupled to a chemostat. We denote the Gibbs free energy difference between the states by Δ*g* = *g*(ON) − *g*(OFF). The entropy production rate at the NESS is

**FIG. 6.**
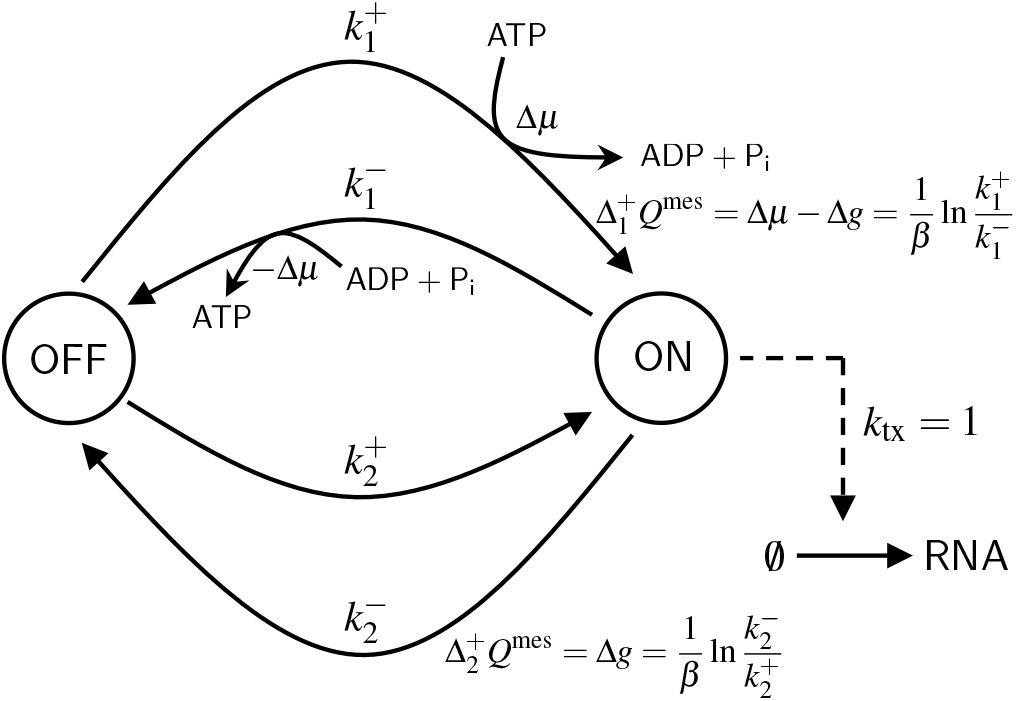
State transition diagram of a two-state promoter model with two microscopically reversible reaction channels. The first reaction channel involves ATP hydrolysis and is therefore coupled to the chemostats of ATP, ADP and P_i_. The equations next to the transition edges are thermodynamic consistency relations (27) of the respective reaction channels, while 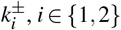 denotes the forward and backward stochastic rate constants of the reactions.

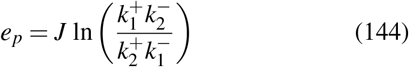

with net probability flux 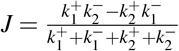, as we show in appendix E. This model provides a minimal example of the underestimation of the true entropy production by the effective entropy production (cf. Section III E), as the effective twostate model always satisfies detailed balance at stationarity, such that its effective entropy production rate vanishes.

The three-state promoter model in Figure 7 extends the twostate model by adding an intermediate state, OFF^∗^, which represents a step in a non-elementary reaction involving ATP hydrolysis. In this model, the first reaction channel handles ATP binding and unbinding between OFF and OFF^∗^, while the second channel accounts for the actual ATP hydrolysis, driving the transition from OFF^∗^ to ON. The third reaction channel allows direct transitions between OFF and ON and is not coupled to any chemostat species. Distinguishing ATP binding/unbinding and hydrolysis may thereby be a more accurate account of the elementary reaction assumption than the single driven channel in the two-state model. Thermodynamically, the three-state promoter is characterized by

**FIG. 7.**
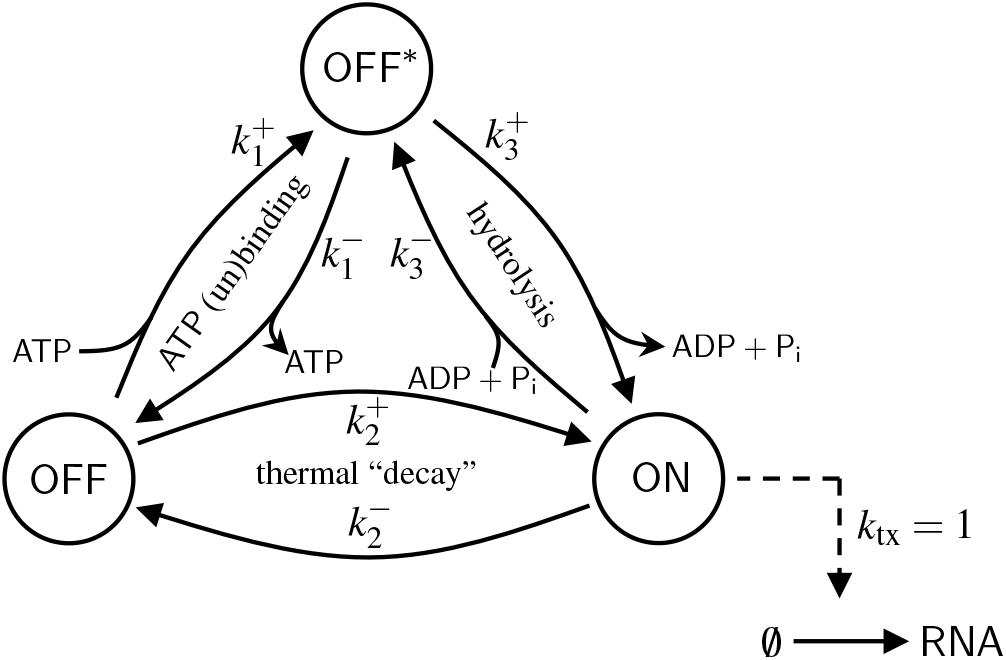
State transition diagram of a three-state promoter model with three microscopically reversible reaction channels. The upper path from OFF to ON involves ATP hydrolysis. Hence, OFF^∗^ is the intermediate of a non-elementary hydrolysis reaction, that was treated as an elementary reaction in the two-state model in Fig. 6. 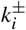, *i* ∈ {1 2 3} denotes the forward and backward stochastic rate constants of the reactions.

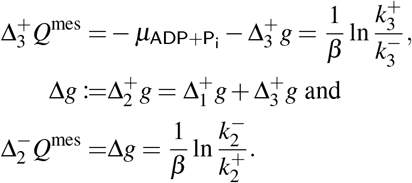

We remark that the difference between work and Gibbs free energy is the mesoscopic heat, while the difference between work an enthalpy is the calorimetric heat, in line with the consistency relation (12). The EPR satisfies

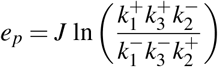

with 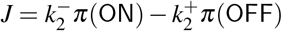, which is related to the general form of the EPR for circular state transition diagrams [113].

We now formulate the maximization problem and use the Shannon-type information capacity notion, i.e., maximum mutual information rate. For the two-state promoter we compute

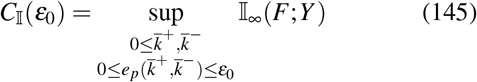

where 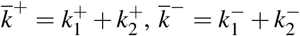 are the effective rates of the transitions between OFF and ON.

The physical boundary conditions 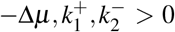 constrain the admissible region of the effective rates:

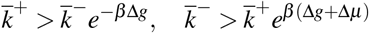

For this admissible region the EPR is thus expressed in terms of the effective rates via

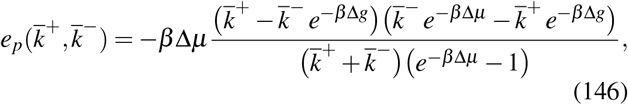

as we show in Appendix E.

The MIR for the model with a two-state promoter has been shown to satisfy [40]

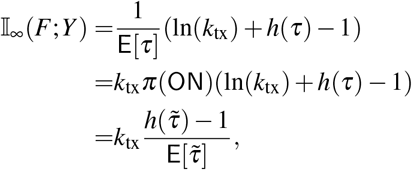

where *h*(*τ*) is the differential entropy of the random time *τ* between transcription events in the dynamics of 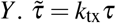 is its dimensionless version. The probability density of *τ* is

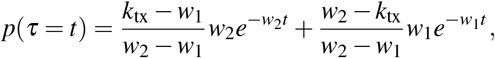

which is an affine combination of exponentials and where *w*_1_, *w*_2_ *>* 0 are the roots of 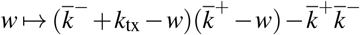 with *w*_1_ *< w*_2_.

Figure 8 presents the numerical solution of (145) for a fixed value of −*β* Δ*µ* = 16, corresponding to the chemical work done on the system via ATP hydrolysis [114], and a fixed ratio of −Δ*g/*Δ*µ* = 0.5. Thereby, fixing Δ*µ* corresponds to fixing the ratio of concentrations 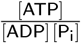 and fixing Δ*g* essentially fixes the identities of core species that participate in the promoter switching. The only degree of freedom left for practical parameter tuning is then the absolute concentration [ATP], keeping the ratio fixed. In Figure 8a the information capacity is evaluated for different *ε*_0_ *>* 0, given that −Δ*g/*Δ*µ* = 0.5. Complementarily, Figure 8b displays a density plot of the EPR *e*_*p*_ in the 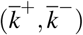-plane together with (i) its hyperbolic level sets and (ii) the curve of MIR maximizing effective rates for varying *ε*_0_ *>* 0. The density plot illustrates that upper bounding the EPR does not upper bound the kinetic rates.

**FIG. 8.**
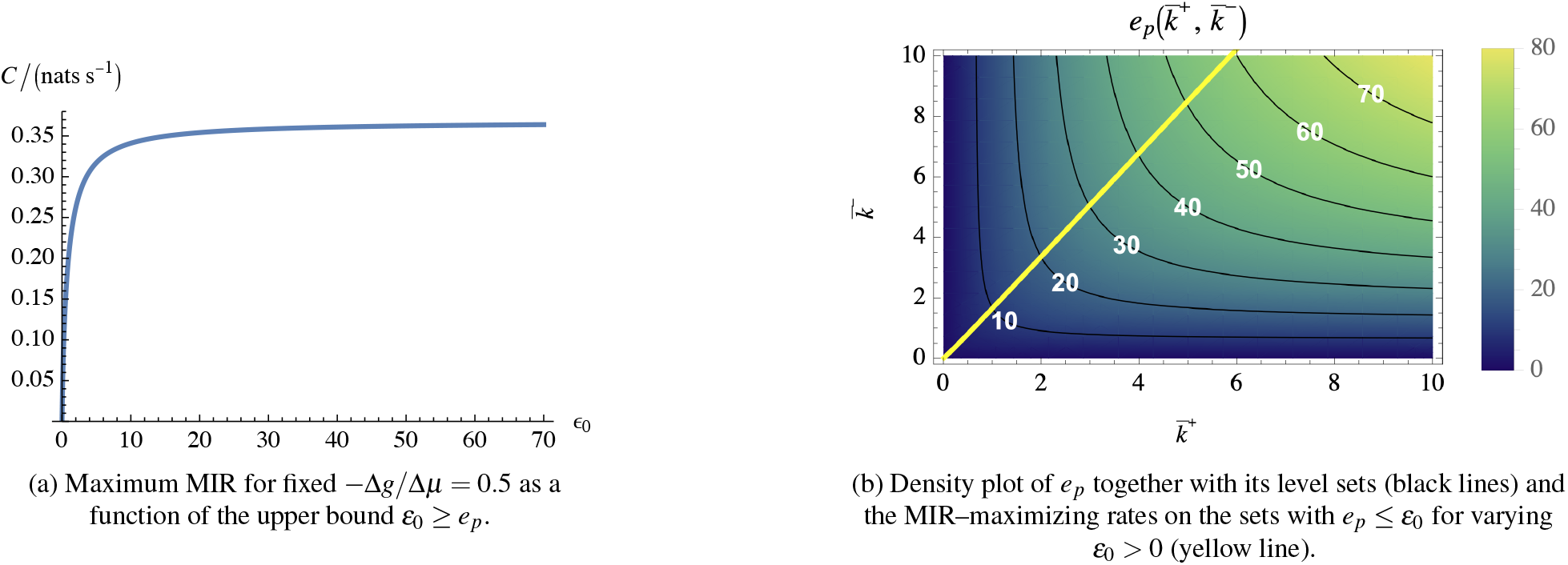
Maximum MIR of the two-state promoter model as a function of the entropy production rate bound. Both MIR and EPR are evaluated for fixed −*β* Δ*µ* = 16 and −Δ*g/*Δ*µ* = 0.5.

In contrast, Figure 9 shows the information capacity for varying −Δ*g/*Δ*µ* and different level sets of *e*_*p*_. Hence, we do not only vary the absolute ATP concentration, but also explore the maximum for a continuous range of possible molecular promoter identities. In the range −Δ*g/*Δ*µ* ∈ [0, 1] the maximum MIR increases monotonously with increasing *e*_*p*_.

**FIG. 9.**
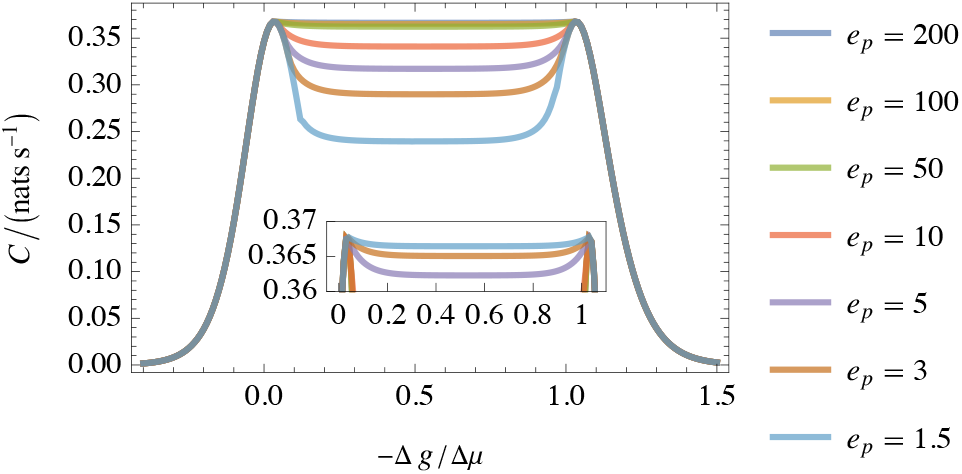
Maximum MIR of the two-state promoter model. The MIR is represented as a function of −Δ*g/*Δ*µ* with fixed −*β* Δ*µ* = 16 for different level sets *e*_*p*_ = *ε*_0_.

For −Δ*g/*Δ*µ /*∈ [0, 1] the level set *e*_*p*_ = 0 in the 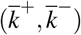 plane always intersects with the MIR maximizing ratio 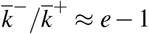 [38]. However, −Δ*g/*Δ*µ <* 0 can be regarded as implausible since ON would then have less Gibbs free energy than OFF. Δ*g/*−Δ*µ >* 1 on the other hand corresponds to 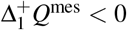, i.e., a chemically driven reaction, which additionally absorbs heat from the environment. Both scenarios are biophysically implausible.

For the three-state promoter let *F* represent the subnetwork {ON}, i.e., the switching dynamics between a transcriptionally active and any inactive promoter state {OFF, OFF^∗^}. We define the capacity problem as

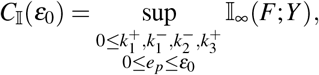

where Δ*g*, Δ*µ* and *k*_tx_ are fixed. The MIR of this model can be shown to have the form [40]

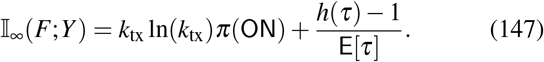

The corresponding information capacity curves for *k*_tx_ = 1 in Figure 10 are analogous to the curves of the two-state promoter.

**FIG. 10.**
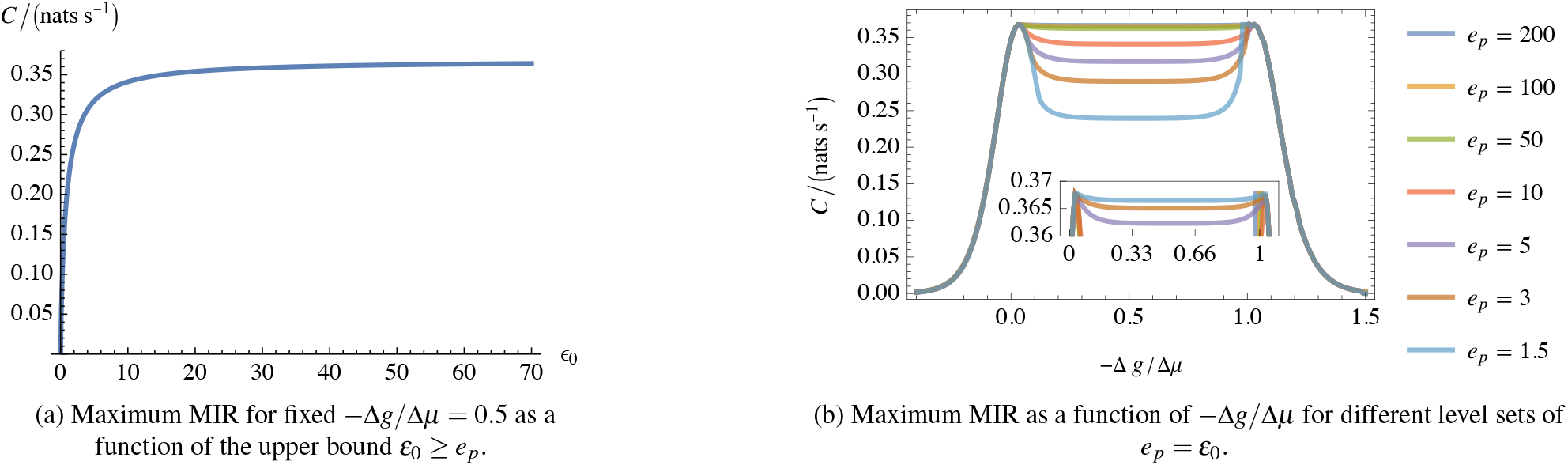
Maximum MIR of the three-state promoter model. Both MIR and EPR are evaluted for fixed −*β* Δ*µ* = 16.

In conclusion, the two distinct input topologies do not change the information capacity of the transcription process channel with respect to kinetic rate optimization under an entropy production rate constraint.

In the light of channel coding, the numerical results suggest that the maximum reliable coding rate satisfies *R*_0_ ⪅ 0.37 natss^−1^ ≈ 0.53 bitss^−1^ in the limit of *e*_*p*_ → ∞, given that *k*_tx_ = 1. Hence, under these conditions, the duration required to reliably distinguish between two different different trajectories is 1.87 s. This must be understood in the asymptotic sense that for large *n* ∈ N a transmission duration of 1.87*n* s is at minimum required to distinguish *n* different promoter trajectories.

## IX. DISCUSSION AND CONCLUSION

### Discussion of related research traditions

In this work, our primary aim was to establish a rigorous chemical communication system together with coding-theorem statements under thermodynamic energy constraints, and thereby a dual minimum-energy-per-bit perspective. In the physical sciences, however, information-theoretic studies of biochemical systems have so far focused more strongly on information-flow analyses, optimization principles and the connection of information handling and thermodynamic work. We briefly outline the research objectives of these traditions.

The field of information dynamics has focused on distributed information processing in complex systems, decomposing dynamics into computational primitives such as information storage and information transfer, with transfer entropy serving as a measure of predictive information transfer between coupled variables [115]. This line of work has also been connected explicitly to stochastic thermodynamics by relating such computational primitives to irreversibility and entropy-production balances [116]. In contrast, the principle of maximum information preservation has been used to test whether biological circuits approach information-optimal designs [6, 13, 117].

Information thermodynamics has more generally examined how information extraction or erasure from a source object via a measurement device enters nonequilibrium thermodynamic balances of work and dissipation [14, 15, 36]. Thereby, information thermodynamics uses system models that differ substantially from the communication model discussed in this work. For example, in [17] information transmission is discretized into cycles that involve a noisy copy operation of a message to the output variable and a subsequent reset of the input variable. Such cycles of information overwriting and copying are much closer to the traditional information thermodynamic engines and allow lower bounding the mismatch cost of communication by the symbol-wise mutual information between the input and output variable under the assumption of an i.i.d. information source and a discrete memoryless channel. By contrast, our communication model seems only scarcely related to previous results of information thermodynamics. In App. D we discuss that under several simplifying assumption, the integrated learning rate [36]

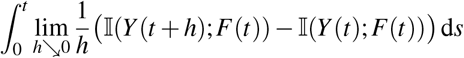

constitutes a common lower bound on both the mean entropy production *H*^tot^(*t*) and the communication-relevant mutual information I(*F*_[0,*t*]_;*Y*_(0,*t*]_). However, the entropy production rate is not necessarily and upper bound on information rate, as is the case for the discrete-time memoryless channel in [17].

From the perspective of the discussed neighboring traditions, the main utility of the present work lies in three specific advances. First, the trajectory information measures discussed here extend previous CRN expressions by considering mutual and directed information between disjoint subnetworks and by addressing indistinguishable reaction channels under projection. Second, the causal information-theoretic formulation in terms of causal kernels, feedback-encoder architectures, and admissible encoding specifications provides a more precise way to formulate optimization questions regarding biochemical information processing in general. Third, the integration of thermodynamically faithful dissipation estimates makes it possible to study information flow and related optimization principles under energetic constraints derived directly from the reaction dynamics rather than from phenomenological cost functions [118, 119]. This is relevant to the hypothesis that information flow acts as an emergent optimization principle in biological evolution [6, 13, 117], as well as to synthetic biology, where the same tools can inform the principled design of chemical reaction networks for energy-efficient information processing.

### Conclusion

This work argues that, rather than focusing primarily on static input-output relationships, biochemical communication should be examined at the trajectory level over an infinite timeframe and take thermodynamic constraints into account. In biochemical reaction networks, both energy dissipation and signaling unfolds through sustained reaction dynamics, often with network-mediated memory and feedback, so that information propagation is naturally carried by stochastic histories of population states rather than by isolated channel uses with deterministically encoded symbols. Symbol-wise descriptions can still be appropriate, but only under additional assumptions. For instance, when a decomposition into successive history-aware encoding intervals is enforced, or when a sufficient time-scale separation generates effective relaxation between symbol-encoding intervals, implying an approximately memoryless channel. Such assumptions are useful in certain reduced models, but they are not generic for biochemical networks. In particular, the resulting notions of capacity in symbol-wise communication are then not always directly related to information rate notions in physical time. Often, biological information processing models rather describe one-shot decision tasks [112, 120] than the sustained encoding and decoding procedures that coding theorems are designed to characterize. These two schemes should be properly distinguished, as the notion of channel capacity does not rigorously apply to such one-shot tasks. Performance limits and energetic costs of sustained biochemical information processing are therefore most accurately formulated at the trajectory level in physical time.

From this viewpoint, the main significance of the present work is not only that it introduces a chemical communication model, but that it makes explicit which structural distinctions are required to formulate communication-theoretic questions meaningfully for biochemical systems. Most importantly, the stochastic encoder architecture, which mediates a stochastic population encoding under the integration of feedback, must be distinguished from the deterministic encoding trajectories that represent the finite set of distinct messages. Only under this separation does it become clear what is fixed by the communication system and what belongs to the code, and only then can notions such as channel capacity, energetic cost functions, and reliable transmission be formulated in an operationally meaningful way. Consequently, the described information modeling is not a technical refinement, but a pre-requisite for posing Shannon information-theoretic questions rigorously in biochemical communication.

The framework also highlights a broader lesson for energy-aware biochemical information processing. A faithful joint treatment of information transmission and energetic cost requires a thermodynamically faithful mesoscopic model of the underlying reaction dynamics. Such a model must apply coarse-graining and represent effective multi-step reactions, and reservoir couplings in a way that preserves the dissipative properties relevant to the communication process. This matters directly for quantities such as the minimum energy required per reliably transmitted bit: if relevant reservoirs are omitted, or if multi-step mechanisms are compressed into effective reactions without including thermodynamic correction terms, dissipation is systematically underestimated. More generally, the present work suggests that a joint analysis of information transmission and energetic cost requires the level of description of the underlying model to be determined by thermodynamic consistency rather than by information-theoretic convenience alone.

Even so, the present framework is best understood as a foundation from which a more complete theory of energy-aware biochemical information processing can develop.

### Limitations and future directions

The most immediate limitation is that a thermodynamically consistent model is formulated at the level of elementary reaction networks, whereas biologically relevant models are often available only in effective, non-elementary form. In practice, this is a substantial restriction: even the transcription of a single gene may involve a very large number of elementary reactions, the corresponding mechanistic information is often only partially known, and the number of molecular states generated by such polymerization processes can quickly become computationally intractable. A central challenge for future work is therefore to develop controlled approximation schemes that connect thermodynamically faithful elementary descriptions to effective network models without losing information-theoretically relevant memory- and feedback-mediating network structures. Moreover, the thermodynamic constraints that are presently most accessible are usually those on total entropy production or mesoscopic heat dissipation, whereas calorimetric heat constraints require microscopic Hamiltonian information and work constraints require explicit thermodynamic potentials.

On the information-theoretic side, the coding-theorem statements developed here still require full validation for continuous-time models, and several relevant biochemical features remain only partially incorporated into the formalism. In particular, biologically plausible restrictions to non-uniform message priors could not yet be fully integrated into the coding-theorem framework, and naturally occurring noisy source encoding and potentially intrinsic biochemical decoding mechanisms, such as sustained decision making, have not yet been accounted for. Also, the possibility that parts of the CRN itself may vary as part of channel coding has only been touched upon here; a particularly interesting open problem is to characterize which changes in kinetics or network structure can be used as encoder architecture designs without changing the causal chemical communication channel. More generally, the present framework should be understood as a foundation rather than a finished theory. It sets up the Shannon’s classical fixed-channel communication setting as a first step toward a variety biochemical information processing problems. By making explicit what is required for a rigorous information-theoretic analysis, it also provides a basis for systematic extensions to other information-theoretic paradigms like, e.g., rate distortion problems. The extent to which particular biological architectures interpretationally approximate the biochemical communication channels described here is still to be understood, and it is likely that they will be found only in special cases. In light of the evolutionary optimization hypothesis and synthetic biology, the theme of joint code–channel co-design subject to additional goal-oriented performance criteria may be explored further in the future.

We believe that the presented theory helps sharpen questions of energy-dependent biochemical information processing by placing Shannon information theory, thermodynamic cost functions, and continuous-time stochastic chemical reaction dynamics within a single rigorous formulation.

## ACKNOWLEDGMENTS

We thank Nicolai Engelmann for helpful discussions.

M.G. developed the methodology, led the formal analysis, contributed to conceptualization, and wrote the original draft; M.G., L.S., and H.K. reviewed and edited the manuscript. L.S. contributed to conceptualization. H.K. supervised the research and led the conceptualization.

The authors declare no competing financial interest.

## AI DISCLOSURE

During preparation of this manuscript, the authors used ChatGPT as a drafting and language-polishing aid to help structure, revise, and improve portions of the text. All arguments, statements, citations, and interpretations were independently verified, revised, and approved by the authors, who take full responsibility for the final content.

## DATA AVAILABILITY

The code used to perform the optimizations is available from the corresponding author upon reasonable request.

## Appendix A Chain rules for directed information

The following chain rules for the Newton DI are derived similarly to the original proofs given by Kramer [84] for the Massey DI.

### Lemma 4

*For the random discrete sequences X*_1:*N*_, *Y*_1:*N*_ *and Z*_1:*N*_ *the DI satisfies the chain rules*

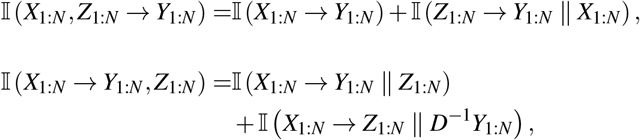

*where D*^−1^ *denotes an anticipatory shift such that*

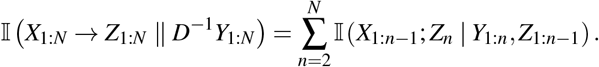

*Proof*.

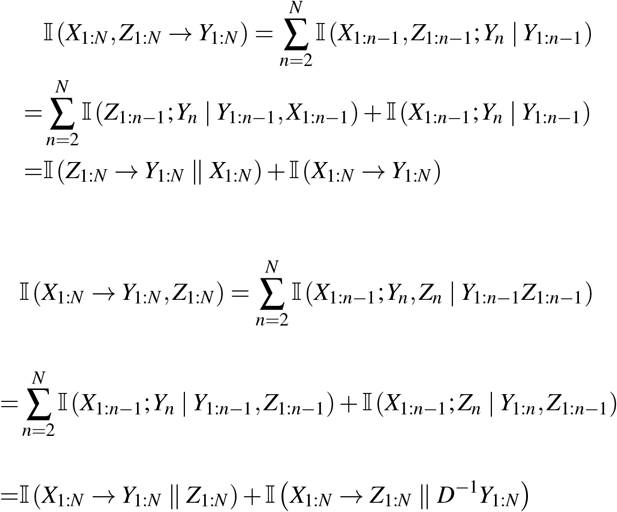

## Appendix B Transfer entropy between subnetworks and causally conditional probabilities

While the primary focus of this paper is on mutual and directed information, we also provide an expression for the transfer entropy between subnetworks to facilitate comparison for an interdisciplinary readership. Following conventional notation, we refer to *X* as the target and *Y* as the source. To accommodate the definition of transfer entropy, we extend the definition of stochastic processes from the finite interval [0, *t*] to the real line ℝ, such that the CRN is assumed to have started at time −∞ rather than at time zero.

Consider the time-ordered families of sigma-algebras 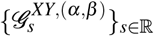 and 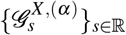, defined by

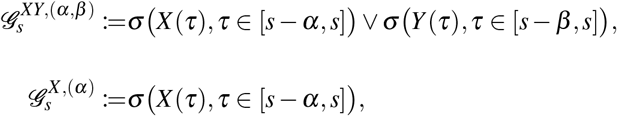

which represent the sliding-window partial histories of the joint process (*X,Y*) and the marginal process *X*, respectively. The parameters *α, β >* 0 specify the history lengths for *X* and *Y*. We emphasize that the time-ordered families are *not* to be confused with the join of *σ*-algebras or filtrations. Often a filtration even cannot even be constructed from such a family, since the fixed-length window definitions imply 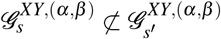 and 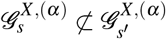 for *s* < *s* ′ in general. The time-ordered sets become equivalent to filtrations only in the limit *α, β* → ∞.

Spinney *et al*. [35] generalized the discrete-time definition of transfer entropy to arbitrary stochastic processes indexed by a strictly ordered and uncountable set T. For continuous-time processes, they assume T ⊇ [*t*_0_ − max(*α, β*), *t*) with *t*_0_ *< t*. Their approach constructs what they refer to as “natural measures”. In the formal notions of measure-theory, they construct (i) a stochastic kernel

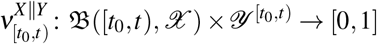

and (ii) a probability measure

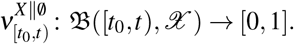

In general, 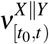 is *not* a regular conditional probability of 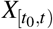 given 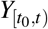, and 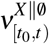 is not simply the marginal probability of 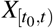, even if the construction involves a family of ℙ-regular conditional probabilities. A rigorous measure-theoretic construction involves the use of Carathéodory’s and Kolmogorov’s extension theorems and is beyond the scope of this work. Nevertheless, if the constructions are based on ℙ-regular conditional probabilities, then we refer to 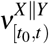 as ℙ*regular transfer probability* of *X* given *Y*, as well as to 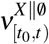 (marginal) ℙ-*regular transfer probability* of *X*, and we denote

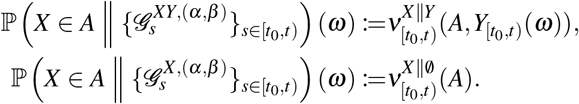

for all *A* ∈ B([*t*_0_, *t*), *X*). This notation combines the timeordered families from [35] and the “causally conditional probability” ℙ(· ∥ ·) as originally defined by Kramer [121]. We emphasize again that ℙ-regular transfer probabilities are in general distinct from ℙ-regular conditional probabilities and the time-ordered families do not represent a single *σ*-algebra or filtration.

Note that the construction of time-ordered families of *σ* algebras is not restricted to sliding windows and can also be applied to arbitrary families of filtrations. Consequently, a direct generalization of Kramer’s discrete-time definition of causally conditional probability to continuous-time (in the sense of Newton) is given by 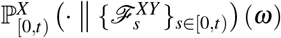 with short notations 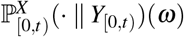 and

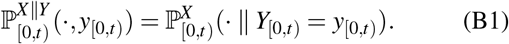

We refer to it as the P-*regular causally conditional probability* ty of *X* given *Y* on [0,*t*). Note that in this case

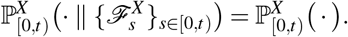

for any non-anticipatory (i.e., ℱ-adapted) stochastic process.

Following Spinney *et al*. [35], the transfer entropy is defined measure-theoretically via the Radon-Nikodym derivative of ℙ-regular transfer probability measures, i.e.,

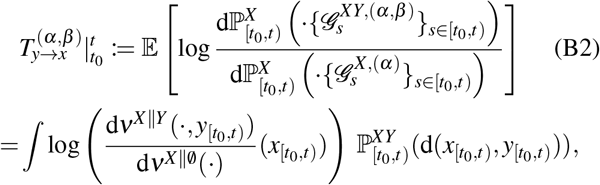

This definition looks similar to the Gel’fand-Yaglom-Perez expression (40), but it is actually neither a relative entropy nor a conditional relative entropy. Similar to (41), the loglikelihood ratio in (B2) can be used as a pathwise, random-variable version of transfer entropy, or “transfer entropy density”, as is common in stochastic thermodynamics.

For comparison, we provide a related generalization of Schreiber’s discrete-time transfer entropy [122] based on the structural, extremal-based approach of Weissman *et al*. [37]

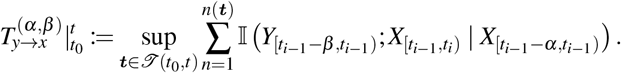

Here, we simply conjecture that the two definitions are equivalent for standard path spaces. Following the approach outlined in Supp. Sec. S1.5, it can be demonstrated that the latter definition yields the same expression for the transfer entropy of jump processes as that derived by Spinney *et al*. [35].

Using the subnetwork propensities (105) we define the (conditional) transfer intensity processes

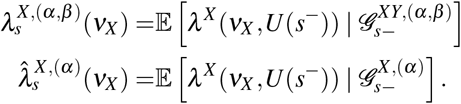

It follows that

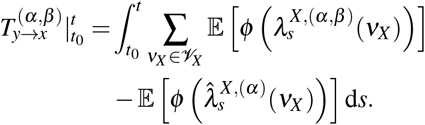

In the limit of full history this yields the relation

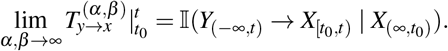

There are two ways to make the transfer entropy equal to the DI. Either artificially set the path of *U* to a constant for all *s <* 0 and randomly select *U* (0). Then

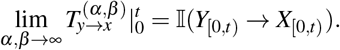

Alternatively, the parameters *α, β* can be interpreted as time-dependent functions *α*(*s*), *β* (*s*). Setting *α*(*s*) = *β* (*s*) = *s* we obtain 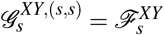 for all *s* ≥ 0, such that

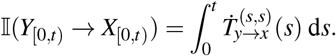

## Appendix C Properties of directed information and reaction-indistinguishability mismatch decomposition

In this section, we expand on the discussion in Sec. VI of the discrepancy that arises when indistinguishable reactions are either resolved or aggregated into a marginal counting process. Rather than using the extremal representation introduced in Sec. V, we express the directed information I(*X*_[0,*t*]_ → *Y*_[0,*t*]_) between subpopulation processes as a relative entropy between path measures, i.e., as the expectation of the logarithm of a Radon–Nikodym derivative; cf. Eqs. (116) and (117).

*Directed information as a relative entropy*. Directed information and causally conditional directed information can be represented as relative entropies between joint path measures. To see this, we define a reference measure such that for every *C* ∈ B([0, *t*], *X* × *Y*)

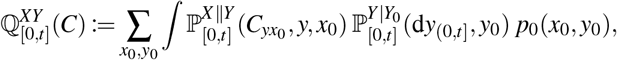

where 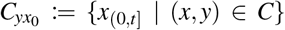. Under the absolutecontinuity assumption 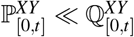, the factorization of the joint Janossy densities gives, 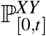-almost surely,

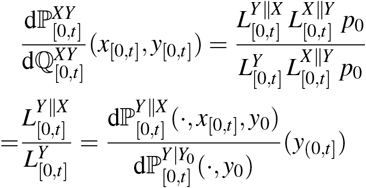

and consequently—as in the discrete-time case [99]—

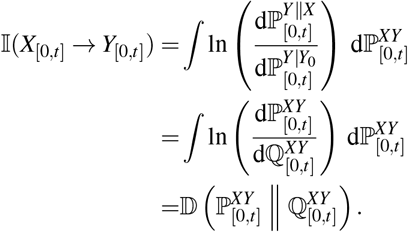

Thus, although directed information is naturally introduced through a conditional log-likelihood ratio, it is equivalently the relative entropy between the joint path law and a reference path law in which the causal influence of *X* on *Y* has been removed.

The same construction applies to causally conditional directed information I(*X*_[0,*t*]_ → *Y*_[0,*t*]_ ∥ *Z*_[0,*t*]_), where the factorization of Janossy densities gives, 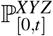-almost surely,

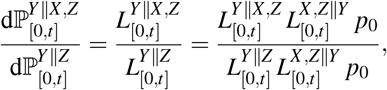

where the enumerator of the r.h.s. is a density of 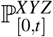.

Hence, both directed information and causally conditional directed information inherit the standard properties of relative entropy, including non-negativity. This agrees with the nonnegativity result in Lemma 1, which we obtained directly from the extremal-based definition.

### Local directed information between counting process characterizations

With this in mind, we now introduce a divergence-type notation 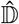, which we refer to as local directed information and which encompasses directed information and causally conditional directed information for counting and population processes. The attribute “local” will be motivated in the subsequent paragraph.

Since the local Janossy densities of counting and population processes are equivalent to a set of intensity processes, we introduce for an arbitrary subset 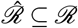 the notation

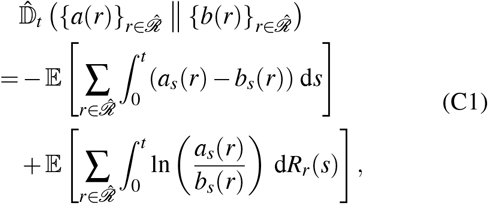

where *a*(*r*) and *b*(*r*), respectively, be *G* ^(*a*)^and *G* ^(*b*)^-predictable non-negative processes such that 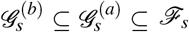 for all *s* ≥ 0 and

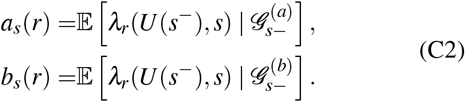

We intentionally do not call these processes intensity processes, since proper intensity processes are required to satisfy the formal adaptedness-condition (cf. [53, p. 27] or [123, Sec. 18.2.1.]), i.e., for all *t* ≥ 0

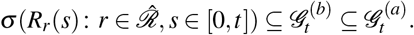

That is, the internal filtration of the counting processes must be a subfiltration of the defining filtrations *G* ^(*b*)^ and *G* ^(*a*)^. Since we have to drop this assumption for the subsequent discussion, we just refer to such processes *a*(*r*) and *b*(*r*) as intensity-type processes. Definition (C1) does analogously apply to merged counting processes, and merged reaction intensities like *λ*^*Y*^ (*ν*_*Y*_), *ν*_*Y*_ ∈ *V*_*Y*_. For instance, we identify

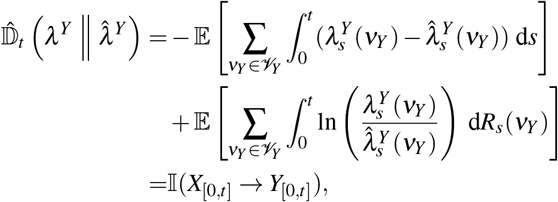

where we omitted the index set in the arguments of 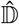 brevity, as it is clear from the context.

Structurally, these local directed information exhibits the chain rule

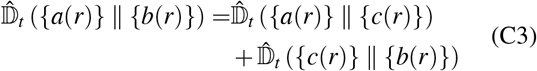

if *c*(*r*) are *G* ^(*b*)^-predictable non-negative processes defined analogously to *a*(*r*) and *b*(*r*), such that *G* ^(*b*)^ ⊆ *G* ^(*c*)^ ⊆ *G* ^(*a*)^ ⊆ ℱ. We call (C3) a chain rule since under a particular choice of intensity processes {*a*(*r*)}, {*b*(*r*)}, {*c*(*r*)}, it is equivalent to Eq. (99).

Further, by non-explosiveness, Fubini’s theorem, [53, Eq. (3.2)] and the tower property of the conditional expectation, (C1) always simplifies to

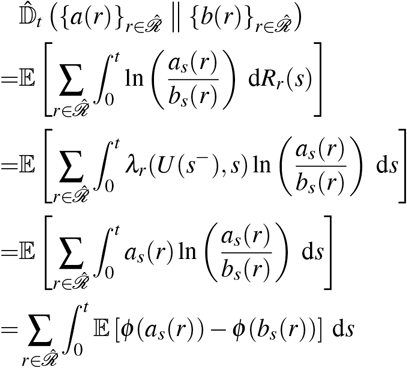

with *φ* (*x*) := *x* ln(*x*)1_(0,∞)_(*x*) as in Sec. VI. Further, by Jensen’s inequality for conditional expectations [55, Thm. 8.20] and the convexity of *φ* on [0, ∞), we have

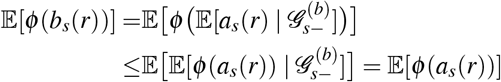

such that

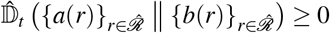

is always satisfied. So, even if one of the intensity-type processes is not a proper intensity process, the non-negativity property of relative entropy is still preserved.

### Decomposition of the indistinguishablity mismatch

The intensity processes 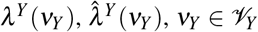, can be decomposed into the intensity-type processes 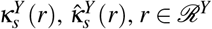; cf. Eq. (113). Then,

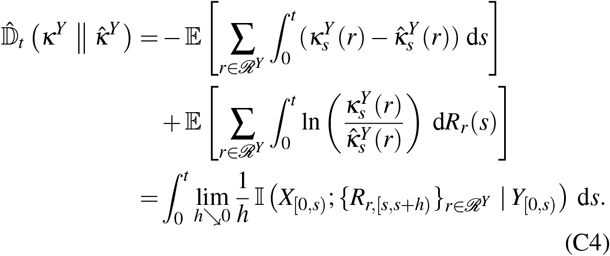

Note that 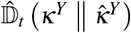 is not a directed information and also not a causally conditional directed information since the processes 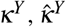 depend on the history of *Y* instead of the history of *R*_*r*_, *r* ∈ ℛ^*Y*^. Exactly for this reason, they are not proper intensity processes of the respective counting processes. Equation (C4) motivates that we refer to 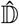 as local directed information, since the local characterization along the interval [0, *t*], i.e. the Lebesgue-density, is structurallly similar the that of a proper directed information (see Eq. (96)).

We can now identify the mismatch between these two quantities, i.e., the mismatch between the non-merged local directed information 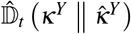 and the directed information, while the history dependence is still on the population process *Y* and not on the counting processes indexed with ℛ^*Y*^. Define

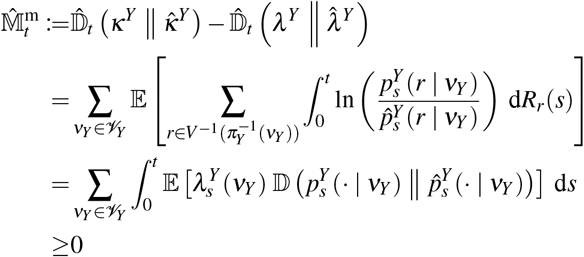

with the time-dependent conditional probability mass functions

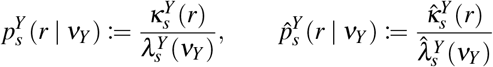

for all 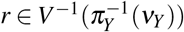.

Now denote for all *t* ≥ 0

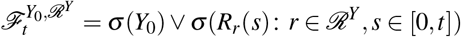

and 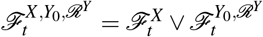.

Then, we can define for all *r* ∈ ℛ^*Y*^ the 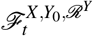 and 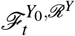-intensity processes of *R, r* ∈ ℛ^*Y*^, as

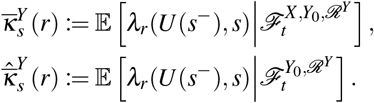

With these intensity processes, we identify

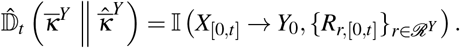

and further define the *Y*-side filtration-mismatch

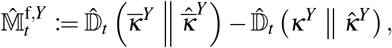

which—unlike the merging mismatch 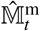—is not non-negative in general. Applying the chain rule (C3) to 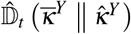 in two different ways yields an alternative expression for the *Y*-side filtration mismatch

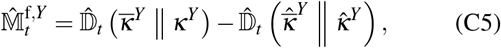

since both

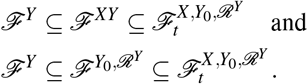

The respective local directed informations in (C5) have the integral-versions

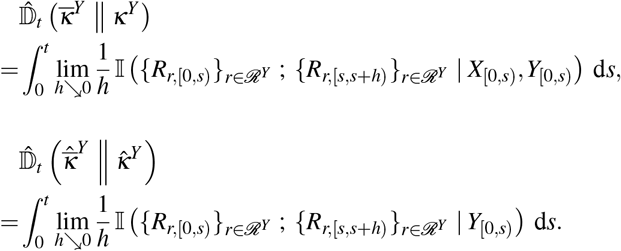

We finally define the *X*-side filtration mismatch as

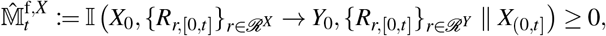

since by the chain rule (99) the directed information between reaction processes decomposes as

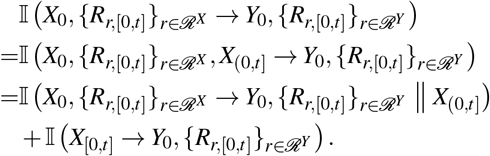

In conclusion, we have derived

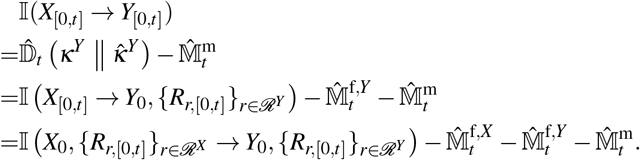

Hence, we can identify

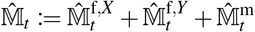

as the total mismatch between the directed information between the population processes and the directed information between the reaction processes, such that

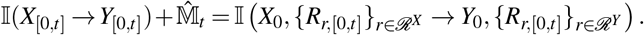

If |*V*_*Y*_ | = |ℛ^*Y*^ |, then 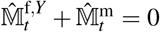. Similarly, if |*V*_*X*_ | = |ℛ^*X*^|, then 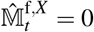. If both conditions are verified, then the total mismatch 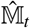 vanishes.

A representation in terms exclusively non-negative quantities is

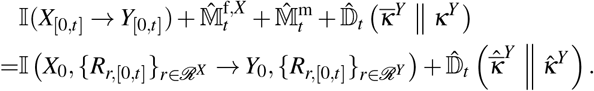

### Mismatch decomposition at density-level

Finally, we note that the mismatch decomposition can also be established at the density-level. We denote

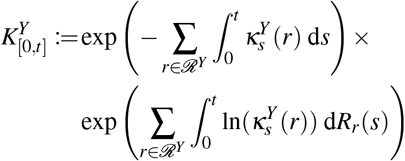

analogously to 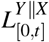 defined in terms of 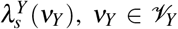. Similarly, we denote 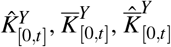 for the respective densities induced by 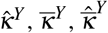. Then we have

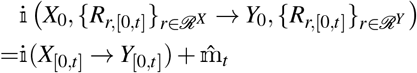

with the mismatch density decomposition

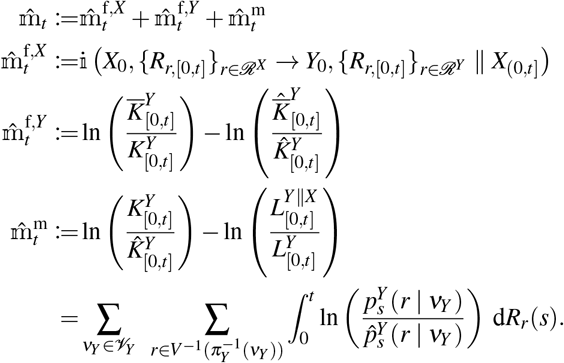

## Appendix D Communication-relevant mutual information, EPR and learning rate

A mismatch decomposition similar as in App. C can also be derived to relate i (*F*_[0,*t*]_;*Y*_(0,*t*]_) and the ℛ^*Y*^-associated part of *h*^tot^(*t*). Throughout this sections, we denote with ℛ the reversible reaction channel indices, and with *ε* ∈ {+,−*}* the direction of the reaction.

### Total entropy production of a subset of reactions

For an arbitrary subset 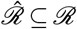 we denote the entropy production mediated by these reactions as

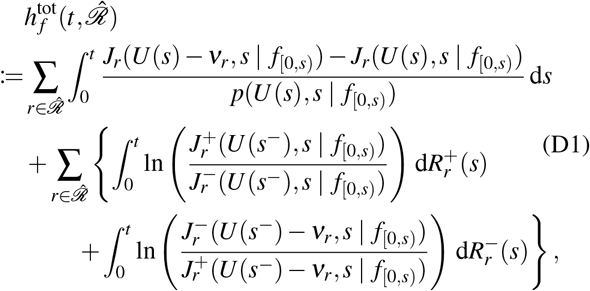

where the dependence on the deterministic chemostat encodings *f* is made explicit. Then, by Eq. (35), we have

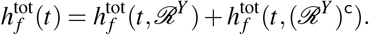

Similarly, we may define the mean entropy production mediated by these reactions as

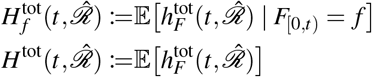

such that

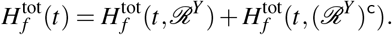

Next, for all *r* ∈ ℛ and *ε* ∈ {+,−}*}*, introduce the *F*^*FU*^ intensity process

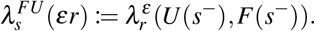

Further, define the forward-time intensity process associated with the time-reversed backward process by

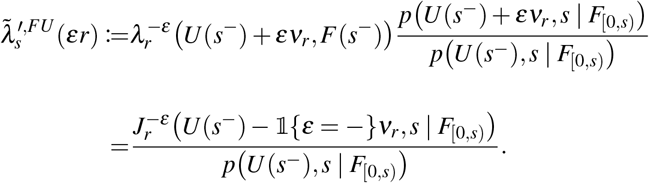

Here 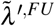 denotes the intensity processes of the time-reversed backward process *Ũ* ′ with propensity functions given in Supp. Eq. (S2.13). The backward process *U* ^′^ evolves under the reversed protocol 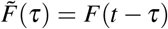, and *Ũ* ′ is its time reversal. In general, *Ũ* ′ is only a reference process and its path law need not coincide with the forward law of *U*.

With these intensity processes we may rewrite (D1) as

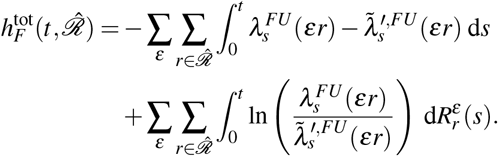

While this expression has the form of a log-likelihood ratio of local Janossy densities, and its expectation has the form of the r.h.s. of Eq. (C1), the two processes *λ* ^*FU*^ and 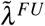 do not jointly statisfy (C2) such that

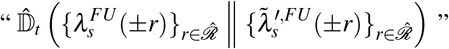

is ill-defined. We have, however, the Lebesgue-density representation

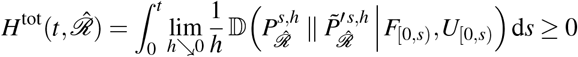

via a conditional relative entropy [75, Ch. 7.2], where

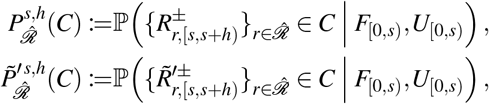

for any 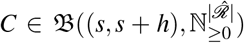, denote the path-laws of the time-forward reaction counters and the time-reversed backward-protocol reaction counters, conditioned on the full history of the CRN and the random protocol. Notably, this density representation implies that the contribution *H*^tot^(*t, {r}*) of any reaction *r* ∈ ℛ to the total entropy production is non-negative.

### Information-entropy mismatch

In the following and denote the Janossy densities of *λ* ^*FU*^ and 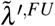 of the set ℛ^*Y*^ as, respectively, 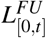 and 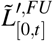. Then,

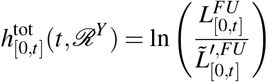

is the partial entropy production of ℛ^*Y*^. It’s mean

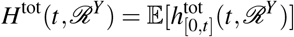

equals the partial entropy production of the population process *Y*, as discussed by Chétrite *et al*. [36, p. 11], if ℛ = ℛ^*Y*^, *S* = *S*_*Y*_ and |*V*_*Y*_ | = |ℛ^*Y*^| together with the assumption that (*F,Y*) is a Markov process. In this case, *F* does not encode information into the population dynamics of a designated encoding species set *S*_*X*_, but instead directy into the population dynamics of the output species *S*_*Y*_. In other words, the propensities of *Y*-changing reactions ℛ^*Y*^ depend directly on the chemostat protocol. Under these simplifying assumptions, the partial entropy production was shown to satisfy [36, Eq. (25)]

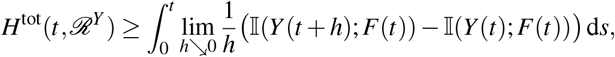

where the lower bound on the r.h.s. is known as the *integrated learning rate*.

Next, we consider the intensity(-like) processes 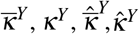, which are assumed to be predictable with respect to the respective filtrations 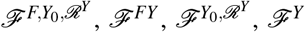. We may directly adapt all relations for these processes from App. C by substitutung *X* with *F*. For instance, it holds that

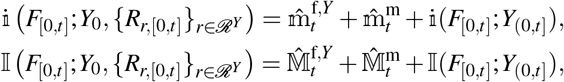

where 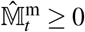, and 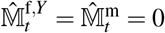 if |*V*_*Y*_ | = |ℛ^*Y*^ |.

We then define the information-entropy mismatch density

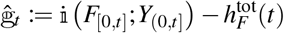

and the mean information-entropy mismatch

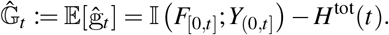

If we assume ℛ = ℛ^*Y*^, and *S* = *S*_*Y*_, then this mismatch density simplifies to

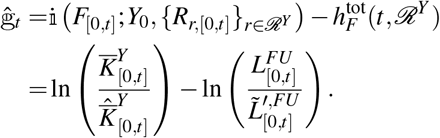

If additionally |*V*_*Y*_ | = |ℛ^*Y*^ |, then

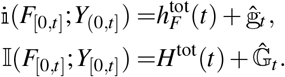

The mean mismatch 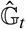 is, however, not non-negative in general and may switch sign depending on 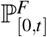. In the special case that additionally (*F,Y*) is a stationary Markov process, it hold that [36, Eq. (18)]

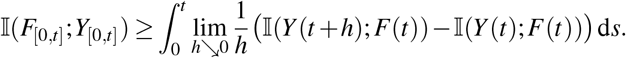

Hence, *H*^tot^(*t*) and I(*F*_[0,*t*]_;*Y*_[0,*t*]_) share the integrated learning rate as a common lower bound in this case, which still leaves the sign of 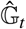 undetermined.

## Appendix E EPR of a non-equilibrium two-state model

To derive the EPR for the two-state model we use Equation (33) for the entropy exchange rate, which is equal to the EPR at the NESS:

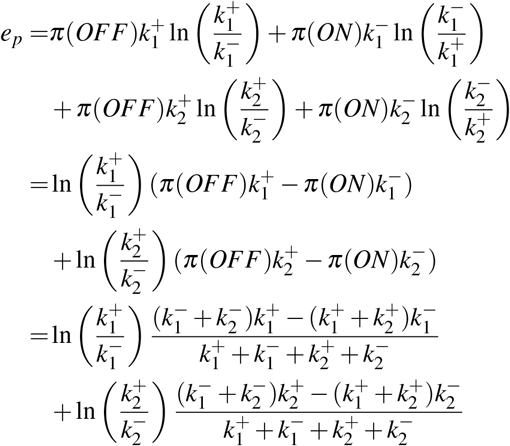

with 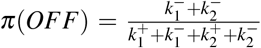 and 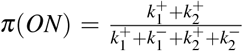. The stationary distribution is the kernel of the generator matrix corresponding to the model. Simplifying the above expression yields (144).

To obtain the EPR as a function of the effective rates 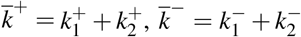, we we express the rates in the state transition diagram via the effective rates and the potential differences Δ*g*, Δ*µ*. We have

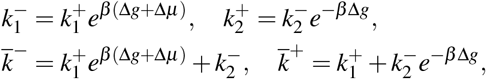

which implies

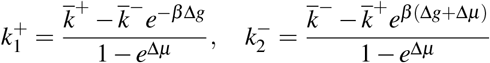

and therefore also

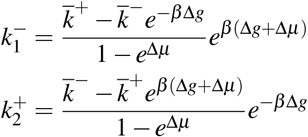

The physical boundary conditions 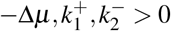 constrain the admissible region of the effective rates to

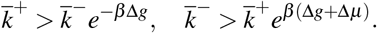

For this region we obtain (146) by substituting the original rates with the above expressions and simplifying.

**Appendix F: Interarrival time between transcription events for the circular three-state promoter**

The DI between the promoter dynamics and the RNA copy number has was linked to the differential entropy of the interarrival time in (147). Here, we apply Anderson’s filtering theorem for semi Markov processes similarly as described in [40]. Define the state space *E* := {J, ON, OFF^∗^, OFF}, where J is the jump state, which mimics ON in terms of leaving transitions. The Laplace version of the semi-Markov kernel density is

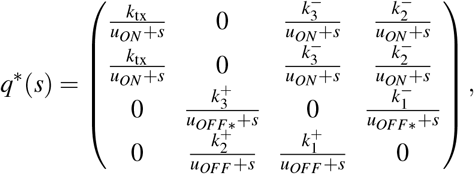

where 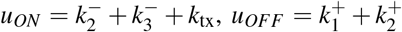 We filter *E* to the reduced state space {J} which yields the interarrival time distribution *f*_*τ*_ of the output. Since the geometric series of a 3 × 3-matrix is hard to evaluate, we do the filtering state by state, from right to left. In the first step we identify

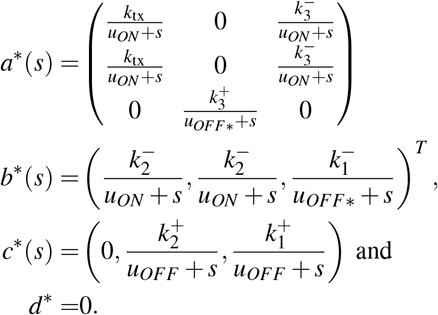

Thus

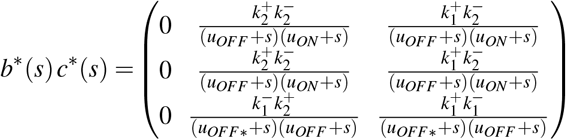

and

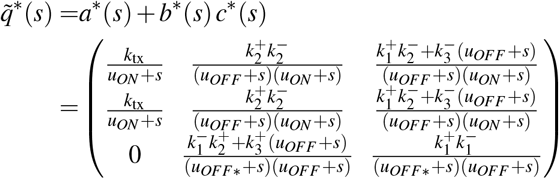

In the second step we identify

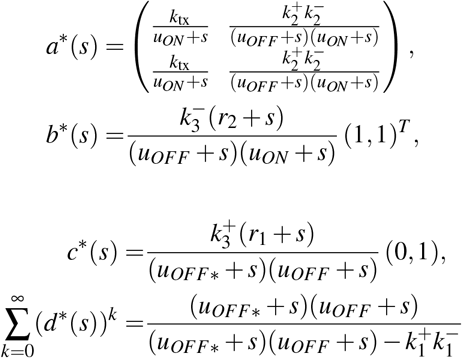

with 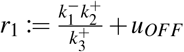 and 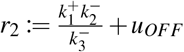. We denote the roots of 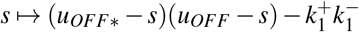 by *w*_1_, *w*_2_. Then

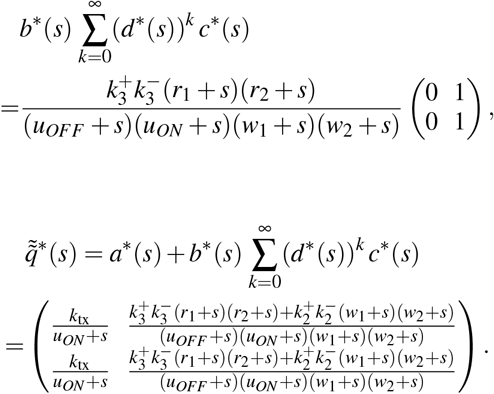

Denoting the roots of 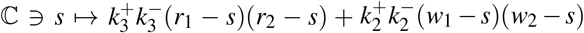 by *q*_1_, *q*_2_ we rewrite

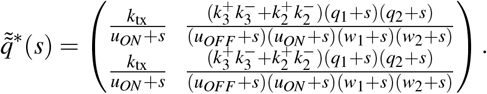

For the last filtering step we use that 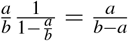 and 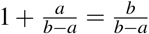 for any *a*≠ *b*. We obtain

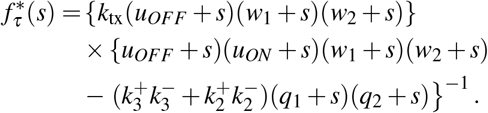

Denote *R*(*s*) := *k*_tx_(*u*_*OFF*_ + *s*)(*w*_1_ + *s*)(*w*_2_ + *s*) and let *v*_1_, …, *v*_4_ be the roots of 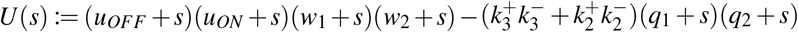. Then

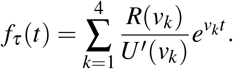

