## Supplementary Material S1 for "Biochemical communication with noisy feedback under energy constraints"

(Dated: July 7, 2026)

### CONTENTS

S1.1. Review of measure theoretic notions 1

S1.2. Mutual information between subnetworks of Markovian CRNs 3
- A. Marginalization and equivalent representations 3
- B. Marked point processes on the state space $\mathscr{X} \times \mathscr{Y}$ 4
- C. Proof of Theorem 1 5

S1.3. Stochastic filtering equations for the intensity processes 7

S1.4. Integral version of directed information 8

S1.5. Proof of directed information between subnetworks of Markovian CRNs 12
- A. Newton directed information 12
- B. Instantaneous information exchange 13
- C. Massey directed information 14

S1.6. Proofs for CRNs as communication channels 14
- A. Equivalence of message and source code conditioning 14
- B. Fano-type converse theorem 17
- C. Energy-per-bit 18

References 19

### S1.1. REVIEW OF MEASURE THEORETIC NOTIONS

We briefly review the main measure-theoretic notions used throughout this work; see [1, 2] for standard references.

A probability space is a triple $(\Omega, \mathscr{F}, \mathbb{P})$ consisting of an abstract set $\Omega$, a $\sigma$-algebra $\mathscr{F}$ on $\Omega$, and a probability measure $\mathbb{P}\colon \mathscr{F} \to [0,1]$ on $(\Omega, \mathscr{F})$, where $\mathbb{P}(\Omega) = 1$. The $\sigma$-algebra specifies which subsets of $\Omega$ are measurable and can therefore be assigned probabilities. It also provides the basic structure needed to define Lebesgue integration on abstract spaces. This extends familiar notions such as Riemann or Stieltjes integration and allows one to integrate random variables on very general state spaces. In particular, if $X\colon \Omega \to \mathbb{R}$ is integrable, then

$$\mathbb{E}[X] = \int_\Omega X(\omega)\, \mathbb{P}(\mathrm{d}\omega).$$

The power set, denoted with $\mathfrak{P}$ contains all subsets of a particular set. Power sets can however usually not be used together with any probability measure, in particular if the space is uncountable. The underlying problem is that there are subsets with very irregular structure that can lead to irregularities in the assignment of probabilities to other sets as well. The typically considered, smaller $\sigma$-algebras do not contain such problematic sets, such that a designated probability measure is well-defined on all sets it contains.

**Definition 1** (Measurable space)**.** *A* measurable space *is a pair* $(\mathscr{X}, \Sigma_X)$ *consisting of a set* $\mathscr{X}$ *and a* $\sigma$*-algebra* $\Sigma_X \subseteq \mathfrak{P}(\mathscr{X})$*. The sets in* $\Sigma_X$ *are called* measurable sets*.*

Thus, a measurable space is the basic object on which one can define measures and measurable maps. A probability space is simply a measurable space together with a probability measure.

**Definition 2** (Generated $\sigma$-algebra)**.** *Let* $\mathscr{C} \subseteq \mathfrak{P}(\mathscr{X})$ *be a collection of subsets of* $\mathscr{X}$*. The* $\sigma$-algebra generated by $\mathscr{C}$*, denoted by* $\sigma(\mathscr{C})$*, is the smallest* $\sigma$*-algebra on* $\mathscr{X}$ *that contains* $\mathscr{C}$*.*

Here "smallest" means that $\sigma(\mathscr{C})$ is contained in every $\sigma$-algebra on $\mathscr{X}$ that contains $\mathscr{C}$. In this sense, $\sigma(\mathscr{C})$ is the minimal measurable structure generated by the collection $\mathscr{C}$.

**Definition 3** (Measurable map and $\sigma$-algebra generated by a random variable)**.** *Let* $(\mathscr{X}, \Sigma_X)$ *and* $(\mathscr{Y}, \Sigma_Y)$ *be measurable spaces. A map*

$$f\colon \mathscr{X} \to \mathscr{Y}$$

*is called* $(\Sigma_X, \Sigma_Y)$-measurable*, or simply* measurable*, if*

$$f^{-1}(B) \in \Sigma_X \qquad \text{for all } B \in \Sigma_Y.$$

*If* $X\colon \Omega \to \mathscr{X}$ *is measurable, then the* $\sigma$*-algebra generated by* $X$ *is*

$$\sigma(X) := \sigma\big(\{X^{-1}(B) \mid B \in \Sigma_X\}\big).$$

Thus, $\sigma(X)$ is the smallest sub-$\sigma$-algebra of $\mathscr{F}$ with respect to which $X$ is measurable. Intuitively, it captures exactly the information contained in the random variable $X$.

**Definition 4** (Borel $\sigma$-algebra)**.** *Let* $\mathscr{X}$ *be a topological or metric space. The* Borel $\sigma$-algebra *on* $\mathscr{X}$*, denoted by* $\mathfrak{B}(\mathscr{X})$*, is the* $\sigma$*-algebra generated by the open subsets of* $\mathscr{X}$*.*

For example, $\mathfrak{B}(\mathbb{R})$ is generated by the open intervals. In many applications, especially on path spaces equipped with a topology or a metric, the Borel $\sigma$-algebra is the natural measurable structure.

**Definition 5** (Join of $\sigma$-algebras). *Let $\Sigma_1$ and $\Sigma_2$ be $\sigma$-algebras on the same set $\mathscr{X}$. Their* join *is*

$$\Sigma_1 \vee \Sigma_2 := \sigma(\Sigma_1 \cup \Sigma_2),$$

*that is, the smallest $\sigma$-algebra on $\mathscr{X}$ containing both $\Sigma_1$ and $\Sigma_2$.*

More generally, if $(\Sigma_i)_{i\in I}$ is a family of $\sigma$-algebras on $\mathscr{X}$, then $\bigvee_{i\in I}\Sigma_i$ denotes the $\sigma$-algebra generated by their union.

**Definition 6** (Almost sure property). *A property $P(\omega)$ is said to hold* $\mathbb{P}$-almost surely, *abbreviated* $\mathbb{P}$-a.s., *if there exists a set $N \in \mathscr{F}$ with $\mathbb{P}(N) = 0$ such that $P(\omega)$ holds for all $\omega \in \Omega \setminus N$.*

Thus, a $\mathbb{P}$-a.s. statement may fail on a null set $N$, i.e., a set with probability zero, but only on such a set.

**Definition 7** (Standard Borel space). *A measurable space $(\mathscr{X}, \Sigma_X)$ is called a* standard Borel space *if it is measurably equivalent to a Borel subset of a complete separable metric space.*

Informally, standard Borel spaces are measurable spaces that are general enough for most applications and still regular enough for probability theory to work smoothly on them. Many basic constructions and existence results are especially clean on such spaces. Important examples are countable sets with their power set, Euclidean spaces with their Borel $\sigma$-algebras, and many path spaces from stochastic-process theory.

**Definition 8** (Filtration). *Let $(\Omega, \mathscr{F})$ be a measurable space and let $\mathbb{T} \subseteq \mathbb{R}$ be a time index set. A family $(\mathscr{F}_t)_{t\in\mathbb{T}}$ of sub-$\sigma$-algebras of $\mathscr{F}$ is called a* filtration *if*

$$\mathscr{F}_s \subseteq \mathscr{F}_t \qquad \text{whenever } s \le t.$$

A filtration models the information available over time: the inclusion $\mathscr{F}_s \subseteq \mathscr{F}_t$ means that information can accumulate as time progresses. If $X = (X_t)_{t\in\mathbb{T}}$ is a stochastic process with values in $(\mathscr{X}, \Sigma_X)$, then its *natural filtration* is

$$\mathscr{F}_t^X := \sigma\big(X_s^{-1}(B) \colon s \le t,\, B \in \Sigma_X\big).$$

**Definition 9** (Conditional expectation with respect to a sub-$\sigma$-algebra). *Let $Z\colon \Omega \to \mathbb{R}$ be an integrable random variable and let $\mathscr{G} \subseteq \mathscr{F}$ be a sub-$\sigma$-algebra. A random variable $U\colon \Omega \to \mathbb{R}$ is called a* conditional expectation of $Z$ given $\mathscr{G}$ *if*

1. *$U$ is $\mathscr{G}$-measurable,*
2. *for all $A \in \mathscr{G}$,*

$$\int_A U(\omega)\,\mathbb{P}(\mathrm{d}\omega) = \int_A Z(\omega)\,\mathbb{P}(\mathrm{d}\omega).$$

*Any such random variable is denoted by*

$$\mathbb{E}[Z \mid \mathscr{G}].$$

The conditional expectation $\mathbb{E}[Z \mid \mathscr{G}]$ is unique $\mathbb{P}$-a.s. It is the $\mathscr{G}$-measurable random variable that reproduces the averages of $Z$ on all events in $\mathscr{G}$. In this sense, it is the canonical summary of $Z$ based only on the information encoded by $\mathscr{G}$.

**Definition 10** (Conditional expectation given a random variable). *Let $Z\colon \Omega \to \mathbb{R}$ be an integrable random variable and let $X\colon \Omega \to \mathscr{X}$ be a random variable into a measurable space $(\mathscr{X}, \Sigma_X)$. The* conditional expectation of $Z$ given $X$ *is defined by*

$$\mathbb{E}[Z \mid X] := \mathbb{E}[Z \mid \sigma(X)].$$

Thus, $\mathbb{E}[Z \mid X]$ means conditioning on the information contained in the random variable $X$. It should not be confused with conditioning on a specific value $X = x$, which is a different notion and typically requires a regular conditional distribution.

**Definition 11** (Conditional probability). *Let $A \in \mathscr{F}$ and let $\mathscr{G} \subseteq \mathscr{F}$ be a sub-$\sigma$-algebra. The* conditional probability of $A$ given $\mathscr{G}$ *is defined by*

$$\mathbb{P}(A \mid \mathscr{G}) := \mathbb{E}[\mathbb{1}_A \mid \mathscr{G}].$$

*If $X\colon \Omega \to \mathscr{X}$ is a random variable into a measurable space $(\mathscr{X}, \Sigma_X)$, then*

$$\mathbb{P}(A \mid X) := \mathbb{P}(A \mid \sigma(X)) = \mathbb{E}[\mathbb{1}_A \mid X].$$

*More generally, if $Y\colon \Omega \to \mathscr{Y}$ is a random variable into a measurable space $(\mathscr{Y}, \Sigma_Y)$, then for each $B \in \Sigma_Y$,*

$$\mathbb{P}(Y \in B \mid X) := \mathbb{E}[\mathbb{1}_{\{Y\in B\}} \mid X].$$

For fixed $B \in \Sigma_Y$, the quantity $\mathbb{P}(Y \in B \mid X)$ is a $\sigma(X)$-measurable random variable on $\Omega$. At this stage, conditional probability is still an object on the underlying probability space. A stochastic kernel provides a stronger representation in which the conditional law is indexed directly by values of $X$.

**Definition 12** (Stochastic kernel). *Let $(\mathscr{X}, \Sigma_X)$ and $(\mathscr{Y}, \Sigma_Y)$ be measurable spaces. A map*

$$\kappa\colon \Sigma_Y \times \mathscr{X} \to [0,1]$$

*is called a* stochastic kernel *from $(\mathscr{X}, \Sigma_X)$ to $(\mathscr{Y}, \Sigma_Y)$ if*

1. *for every $x \in \mathscr{X}$, the map*

   $$B \mapsto \kappa(B, x)$$

   *is a probability measure on $(\mathscr{Y}, \Sigma_Y)$;*
2. *for every $B \in \Sigma_Y$, the map*

   $$x \mapsto \kappa(B, x)$$

   *is $\Sigma_X$-measurable.*

A stochastic kernel describes a probability distribution on the target space that depends measurably on a point of the source space. In probability theory and information theory, stochastic kernels are the natural measure-theoretic formulation of channels and conditional laws.

**Definition 13** ($\mathbb{P}$-regular conditional distribution). *Let* $(\Omega,\mathscr{F},\mathbb{P})$ *be a probability space, and let*

$$X:\Omega\to\mathscr{X},\qquad Y:\Omega\to\mathscr{Y}$$

*be random variables into measurable spaces* $(\mathscr{X},\Sigma_X)$ *and* $(\mathscr{Y},\Sigma_Y)$. *A stochastic kernel*

$$\kappa\colon \Sigma_Y\times\mathscr{X}\to[0,1]$$

*is called a* $\mathbb{P}$-regular conditional distribution of $Y$ given $X$ *if, for every* $B\in\Sigma_Y$,

$$\kappa(B,X(\omega))=\mathbb{P}(Y\in B\mid X)(\omega),\qquad \mathbb{P}\text{-}a.s.$$

Thus, a $\mathbb{P}$-regular conditional distribution is a stochastic kernel that realizes the conditional probabilities of $Y$ given $X$ as a measurable family of probability measures indexed by $x\in\mathscr{X}$. We use the slightly nonstandard terminology $\mathbb{P}$-*regular conditional distribution* to emphasize that such a kernel is always defined relative to a fixed ambient probability measure $\mathbb{P}$ on the underlying probability space.

### S1.2. MUTUAL INFORMATION BETWEEN SUBNETWORKS OF MARKOVIAN CRNS

In the following we provide the details for the proof of Theorem 1. We first discuss the marginalization and equivalent counting process representations of the relevant processes, which we subsequently use for the proof.

#### A. Marginalization and equivalent representations

The process $(X,Y)$ – even if it is not bipartite – is a fundamental process of the kind in [3, §2.13 ], since $\mathscr{R}$ is finite. The coordinate map $\pi_{XY}:\mathscr{V}\to\mathbb{Z}_{\geq 0}^{|\mathscr{S}_X\cup\mathscr{S}_Y|}$ yields the nonzero change vectors $\mathscr{V}_{XY}:=\{\pi_{XY}(v)|v\in\mathscr{V}\}\setminus\{0\}$. Analogously to (104), we define the reaction counting processes $R_s^{XY}(v_{XY})$ for all $v_{XY}\in\mathscr{V}_{XY}$. In turn, they have predictable $\mathscr{F}^{XY}$-intensity processes $\lambda^{XY}(v_{XY})$, such that

$$\lambda_s^{XY}(v_{XY})=\mathbb{E}\left[\lambda^{XY}(v_{XY},U(s^-))\mid\mathscr{F}_{s-}^{XY}\right]\qquad\text{(S1.1)}$$

with $\lambda^{XY}(v_X,u):=\sum_{v\in\pi_{XY}^{-1}(v_{XY})}\lambda_{V^{-1}(v)}(u)$.

Consequently, the joint subsystems obey the following stochastic equation

$$\begin{pmatrix}X(t)\\Y(t)\end{pmatrix}=\begin{pmatrix}X(0)\\Y(0)\end{pmatrix}+\sum_{v_{XY}\in\mathscr{V}_{XY}}v_{XY}\,R_t^{XY}(v_{XY})\qquad\text{(S1.2)}$$

and are hence equivalently represented by

$$A_{[0,t]}:=\left\{(X(0),Y(0)),R_s^{XY}(v_{XY})\colon v_{XY}\in\mathscr{V}_{XY}\right\}_{s\in[0,t]}.$$

That is, there exist maps that uniquely map between realizations of $A_{[0,t]}$ and $(X,Y)_{[0,t]}$ for all $t\geq 0$ and, in particular, it holds that $\mathscr{F}^A=\mathscr{F}^{XY}$ (for details see, e.g., the Appendix of [4]). Formally, the identity (S1.1) follows by the "Change of History for Intensities" Theorem [5, cf. p.32 - 33] applied to $A$, the process equivalence and the predictability of the r.h.s. of (S1.1). The path-distribution of the process (S1.2) on the interval $[0,t]$ is $\mathbb{P}_{[0,t]}^{XY}$.

Similarly, the marginal processes, following $\mathbb{P}_{[0,t]}^X$ and $\mathbb{P}_{[0,t]}^Y$, respectively, are given by (106) with predictable $\mathscr{F}^X$- and $\mathscr{F}^Y$-intensity processes $\hat{\lambda}^X(v_X)$, $\hat{\lambda}^Y(v_Y)$ as in (108) and (110). The process $X$ is equivalently represented by

$$A_{[0,t]}^X:=\left\{X(0),R_s^X(v_X)\colon v_X\in\mathscr{V}_X\right\}_{s\in[0,t]}$$

and analogously for $Y$.

If $X$ and $Y$ are strongly bipartite and $\mathscr{V}_X,\mathscr{V}_Y\neq\emptyset$, then exists a bijection between $\mathscr{V}_{XY}$ and $(\mathscr{V}_X\times\{0\})\cup(\{0\}\times\mathscr{V}_Y)$, such that

$$\lambda_s^{XY}(v_{XY})=\lambda_s^X(v_X)\mathbb{1}\{v_Y=0\}+\lambda_s^Y(v_Y)\mathbb{1}\{v_X=0\}$$

for all $(v_X,v_Y)^\mathsf{T}=v_{XY}\in\mathscr{V}_{XY}$. Then (S1.2) can be rewritten as

$$\begin{aligned}\begin{pmatrix}X(t)\\Y(t)\end{pmatrix}=\begin{pmatrix}X(0)\\Y(0)\end{pmatrix}&+\sum_{v_X\in\mathscr{V}_X}\begin{pmatrix}v_X\\0\end{pmatrix}R_t^X(v_X)\\&+\sum_{v_X\in\mathscr{V}_X}\begin{pmatrix}0\\v_Y\end{pmatrix}R_t^Y(v_Y)\end{aligned}$$

with reaction counters as defined in (104). This equation implies the identity $A_{[0,t]}=B_{[0,t]}$ with

$$B_{[0,t]}:=\left\{(X(0),Y(0)),R_s^X(v_X),R_s^Y(v_Y)\colon v_X\in\mathscr{V}_X,v_Y\in\mathscr{V}_Y\right\}_{s\in[0,t]}.$$

Now consider the case that $X$ and $Y$ are not strongly bipartite. Interestingly, when reducing to marginal change vectors on the subsystems, reactions that simultaneously affect both $X$ and $Y$ can merge with those that affect only $X$ or $Y$ individually. For example, consider the species X, Y, A with the reactions $\emptyset\to\mathsf{X}$, $\mathsf{Y}\to\mathsf{X}$, $\mathsf{A}+\mathsf{X}\to 2\mathsf{X}$, $\mathsf{Y}\to\emptyset$, $\emptyset\to\mathsf{Y}$. We identify the process $X$ with the dynamics of the X copy number, and analogously for $Y$. The reactions in the set $\{\emptyset\to\mathsf{X},\mathsf{Y}\to\mathsf{X},\mathsf{A}+\mathsf{X}\to 2\mathsf{X}\}$ are $X$-indistinguishable with $v_X=1$ and those in $\{\mathsf{Y}\to\mathsf{X},\mathsf{Y}\to\emptyset\}$ are $Y$-indistinguishable with $v_Y=-1$. Consequently, $R^X(1)$ and $R^Y(-1)$ exhibit both simultaneous and individual jumps.

While without the bipartite assumption we still have

$$\begin{pmatrix}X(t)\\Y(t)\end{pmatrix}=\begin{pmatrix}X(0)\\Y(0)\end{pmatrix}+\begin{pmatrix}\sum_{v_X\in\mathscr{V}_X}v_X\,R_t^X(v_X)\\\sum_{v_Y\in\mathscr{V}_Y}v_Y\,R_t^Y(v_Y)\end{pmatrix},$$

with definition (104) and we have $\mathscr{F}^B=\mathscr{F}^A=\mathscr{F}^{XY}$, the process $B$ no longer classifies as a multivariate point process [5, 6], since these must always correspond to "orthogonally thinned" univariate point processes. That is, no two coordinates are allowed to have common jump times in a multivariate point process. Similarly, the process $B$ is also incompatible with the fundamental jump process definition [3].

Note that, while $R^X(v_X)$ and $R^Y(v_Y)$ still have $\mathscr{F}^{XY}$-intensity processes $\lambda^X(v_X)$ and $\lambda^Y(v_Y)$ (cf. Eqs. (107),

(109)), the respective martingales are no longer orthogonal. Hence, stochastic simulation of $B$ with these intensities requires an additional decision-step that chooses, whether a given change in, e.g., $R^X(\nu_X)$ at time $s>0$, is accompanied with a simultaneous change in $R^Y(\nu_Y)$ for some $\nu_Y \in \mathscr{V}_Y$. The decision is made upon sampling

$$\nu_Y \sim \rho_s^{\nu_X}(\nu_Y) := \frac{\lambda_s^{XY}\left(\begin{pmatrix}\nu_X\\ \nu_Y\end{pmatrix}\right)}{\lambda_s^X(\nu_X)},$$

where $\nu_Y = 0$ means "no common jump" and $\nu_Y \neq 0$ implies a common jump, i.e., raising the counter $\Delta R_s^Y(\nu_Y) = 1$. Omitting these update steps would result in a path probability, where common jumps have probability zero.

In conclusion, $B$ is an equivalent counting process description of $(X,Y)$ if and only if $X$ and $Y$ are weakly bipartite (cf. Def. 14).

#### B. Marked point processes on the state space $\mathscr{X} \times \mathscr{Y}$

Additionally to the equivalent multivariate point process descriptions $A$, $A^X$ and $A^Y$ with coordinates for each reaction type, we now introduce equivalent marked point processes [5, 6] via counting measures on the measurable state spaces $(\mathscr{X}\times\mathscr{Y}, \Sigma_X \otimes \Sigma_Y)$, $(\mathscr{X}, \Sigma_X)$ and $(\mathscr{Y}, \Sigma_Y)$. Let

$$N^{XY}(s) := \sum_{\nu_{XY}\in\mathscr{V}_{XY}} R_s^{XY}(\nu_{XY})$$
$$N^X(s) := \sum_{\nu_X\in\mathscr{V}_X} R_s^X(\nu_X), \quad N^Y(s) := \sum_{\nu_Y\in\mathscr{V}_Y} R_s^Y(\nu_Y)$$

denote the counting process of all jumps of $(X,Y)$, $X$ and $Y$, respectively.

In accordance with the notation in [3, §2.9] we define the random counting measure associated with $(X,Y)$ as

$$P_1(C,s) := \delta_{(X(0),Y(0))}(C) + \int_0^t \mathbb{1}\{(X(s),Y(s)) \in C\}\, \mathrm{d}N_s^{XY}$$

for all $C \in \Sigma_X \otimes \Sigma_Y$ and $s \geq 0$ with Dirac measure $\delta_{(x,y)}(C) = \mathbb{1}\{(x,y)\in C\}$ for all $(x,y) \in \mathscr{X}\times\mathscr{Y}$. Similarly define random counting measures for $X$ and $Y$:

$$P^X(C_X,s) := \delta_{X(0)}(C_X) + \int_0^t \mathbb{1}\{X(s)\in C_X\}\,\mathrm{d}N^X(s), \quad C_X \in \Sigma_X$$

$$P^Y(C_Y,s) := \delta_{Y(0)}(C_Y) + \int_0^t \mathbb{1}\{Y(s)\in C_Y\}\,\mathrm{d}N^Y(s), \quad C_Y \in \Sigma_Y.$$

These random measures count the number of visits of the respective set of states in the interval $[0,s]$.

The integrator $\mu$ specified in Theorem 4.1, is the product measure

$$\mu = \Big(\sum_{x\in\mathbb{Z}_{\geq 0}^{|\mathscr{S}^X|}} \delta_x \quad \otimes \sum_{y\in\mathbb{Z}_{\geq 0}^{|\mathscr{S}^Y|}} \delta_y\Big) \otimes (\delta_0 + \tilde{\mu})$$

where $\delta_x$ denotes a Dirac measure at $x \in \mathbb{Z}_{\geq 0}^{|\mathscr{S}^X|}$, $\delta_0$ a Dirac measure at $0 \in \mathbb{R}_{\geq 0}$ and $\tilde{\mu}$ is the Lebesgue measure on $\mathbb{R}_{\geq 0}$. For any function $f\colon \mathscr{X}\times\mathscr{Y}\times\mathbb{R}_{\geq 0} \to \mathbb{R}_{\geq 0}$ we can then express integrals

$$\int_{\mathscr{X}_\circ\times\mathscr{Y}_\circ\times[0,t]} f(x,y,s)\,\mathrm{d}\mu(x,y,s) = \sum_{x\in\mathscr{X}_\circ}\sum_{y\in\mathscr{Y}_\circ}\left(f(x,y,0) + \int_0^t f(x,y,s)\,\mathrm{d}s\right)$$

for any $\mathscr{X}_\circ \subseteq \mathscr{X}$, $\mathscr{Y}_\circ \subseteq \mathscr{Y}$.

With this integrator we can express the compensating random measures

$$\tilde{P}_1(C,t) := \int_{C\times[0,t]} g_1(x,y,s)\,\mathrm{d}\mu(x,y,s)$$

where $\tilde{P}_1 \ll \mu$ and the density has the form

$$\begin{aligned} g_1(x,y,s) &:= \sum_{r=1}^{|\mathscr{R}|} \lambda_r\left(U(s^-),s\right) \\ &\qquad \times \mathbb{1}\{\pi_{XY}(\nu_r) = \left(x - X(s^-), y - Y(s^-)\right)^\mathsf{T}\} \\ &= \sum_{\nu_{XY}\in\mathscr{V}_{XY}} \lambda^{XY}\left(\nu_{XY}, U(s^-), s\right) \\ &\qquad \times \mathbb{1}\{\nu_{XY} = \left(x - X(s^-), y - Y(s^-)\right)^\mathsf{T}\}, \end{aligned}$$

for $s>0$, where we even allow for time-inhomogeneities in the propensity functions, and $g_1(x,y,0) := \mathbb{P}(X(0)=x, Y(0)=y)$. With $\tilde{P}$ defined in this way,

$$P_1(C,\cdot) - \tilde{P}_1(C,\cdot)$$

is an $\mathscr{F}$-local martingale for all $C \in \Sigma_X \otimes \Sigma_Y$. By virtue of (S1.1) and [3, Prop. 4.1] we obtain the unique compensating random measure

$$\begin{aligned} \tilde{P}_1^{XY}(C,t) &:= \int_{C\times[0,t]} \hat{g}_1(x,y,s)\,\mathrm{d}\mu(x,y,s) \\ \hat{g}_1(x,y,s) &:= \sum_{\nu_{XY}\in\mathscr{V}_{XY}} \lambda_s^{XY}(\nu_{XY}) \\ &\qquad \times \mathbb{1}\{\nu_{XY} = \left(x - X(s^-), y - Y(s^-)\right)^\mathsf{T}\} \end{aligned}$$

for $s>0$ and $\hat{g}_1(x,y,0) := \mathbb{P}(X(0)=x, Y(0)=y) =: p_0^{XY}(x,y)$ with respect to the filtration $\mathscr{F}^{XY}$. In the same way we derive compensating measures for $X$ and $Y$.

Let $\mu_X := \left(\sum_{x\in\mathbb{Z}_{\geq 0}^{|\mathscr{S}^X|}} \delta_x\right) \otimes (\delta_0 + \tilde{\mu})$ and $\mu_Y := \left(\sum_{y\in\mathbb{Z}_{\geq 0}^{|\mathscr{S}^Y|}} \delta_y\right) \otimes (\delta_0 + \tilde{\mu})$. Then

$$\tilde{P}^X(C_X,t) := \int_{C_X\times[0,t]} \hat{g}^X(x,s)\,\mathrm{d}\mu_X(x,s),$$
$$\tilde{P}^Y(C_Y,t) := \int_{C_Y\times[0,t]} \hat{g}^Y(y,s)\,\mathrm{d}\mu_Y(y,s)$$

with the respective densities

$$\begin{aligned}\hat{g}^X(x,s) :=&\mathbb{P}(X(0)=x)\mathbb{1}\{s=0\}\\ &+\mathbb{1}\{s>0\}\sum_{v_X\in\mathscr{V}_X}\hat{\lambda}_s^X(v_X)\mathbb{1}\{v_X=x-X(s^-)\},\\ \hat{g}^Y(y,s) :=&\mathbb{P}(Y(0)=y)\mathbb{1}\{s=0\}\\ &+\mathbb{1}\{s>0\}\sum_{v_Y\in\mathscr{V}_Y}\hat{\lambda}_s^Y(v_Y)\mathbb{1}\{v_Y=y-Y(s^-)\}\end{aligned}$$

satisfy that $P^X(C_X,\cdot)-\tilde{P}^X(C_X,\cdot)$ is an $\mathscr{F}^X$-local martingale and $P^Y(C_Y,\cdot)-\tilde{P}^Y(C_Y,\cdot)$ is an $\mathscr{F}^Y$-local martingale for all $C_X\in\Sigma_X, C_Y\in\Sigma_Y$. Given all these definitions, we finally conduct the proof of Theorem 1.

### C. Proof of Theorem 1

To formally distinguish between the joint and the product path probability measures we introduce a second auxiliary probability measure $\mathbb{Q}$, such that $(\Omega,\mathscr{F}^{XY},\mathbb{Q})$ is a probability space and $\mathbb{Q}^{XY}_{[0,t]}:=\mathbb{Q}\circ(X,Y)^{-1}_{[0,t]}=\mathbb{P}^X_{[0,t]}\otimes\mathbb{P}^Y_{[0,t]}$. Our goal is to obtain the Radon-Nikodym derivative

$$L_t(\omega):=\mathbb{E}\left[\frac{\mathrm{d}\mathbb{P}}{\mathrm{d}\mathbb{Q}}\middle|\mathscr{F}^{XY}_t\right](\omega)=\frac{\mathrm{d}\mathbb{P}^{XY}_{[0,t]}}{\mathrm{d}\mathbb{Q}^{XY}_{[0,t]}}\left(X_{[0,t]}(\omega),Y_{[0,t]}(\omega)\right)$$

in order to evaluate the MI

$$\mathbb{I}(X_{[0,t]};Y_{[0,t]})=\mathbb{E}\left[\log(L_t)\right].$$

We do the proof in four steps. First we show some basic properties of $\mathbb{Q}$, which will be used later on. Secondly, we show that $(X,Y)$ is weakly bipartite under $\mathbb{Q}$, even if it is not under $\mathbb{P}$. This property is used to construct the counting measure and compensator of $(X,Y)$ under $\mathbb{Q}$. Thirdly, we apply Theorem 4.1 of [3] to the strongly bipartite case and obtain the MI for this case. Lastly, we show how the non-existence of the RN-derivative almost immediately follows for the given non-bipartite conditions.

*Step 1:*
Denote the expectations w.r.t. $\mathbb{Q}$ with $\mathbb{E}_\mathbb{Q}$ and consider a factorizing function $f(x_{[0,t]},y_{[0,t]},t)=f_X(x_{[0,t]},t)f_Y(y_{[0,t]},t)$ with $\mathbb{E}_\mathbb{Q}[|f(X_{[0,t]},Y_{[0,t]},t)|]<\infty$. Then, by the definition of $\mathbb{Q}$ it holds for all $t\geq 0$ that

$$\begin{aligned}&\mathbb{E}_\mathbb{Q}[f(X_{[0,t]},Y_{[0,t]},t)]\\ =&\mathbb{E}[f_X(X_{[0,t]},t)]\,\mathbb{E}[f_Y(Y_{[0,t]},t)]\\ =&\mathbb{E}_\mathbb{Q}[f_X(X_{[0,t]},t)]\,\mathbb{E}_\mathbb{Q}[f_Y(Y_{[0,t]},t)],\end{aligned}$$

and for any $s\in[0,t]$ that

$$\begin{aligned}&\mathbb{E}_\mathbb{Q}[f(X_{[0,t]},Y_{[0,t]},t)\mid\mathscr{F}^{XY}_s]\\ =&\mathbb{E}[f_X(X_{[0,t]},t)\mid\mathscr{F}^X_s]\,\mathbb{E}[f_Y(Y_{[0,t]},t)\mid\mathscr{F}^Y_s]\\ =&\mathbb{E}_\mathbb{Q}[f_X(X_{[0,t]},t)\mid\mathscr{F}^X_s]\,\mathbb{E}_\mathbb{Q}[f_Y(Y_{[0,t]},t)\mid\mathscr{F}^Y_s].\end{aligned}\tag{S1.3}$$

*Step 2:*
We now turn to the proof of weak bipartiteness under $\mathbb{Q}$. First note that

$$\begin{aligned}\mathbb{Q}(\Delta X(s)\neq 0)=\mathbb{P}(\Delta X(s)\neq 0)=0\\ \mathbb{Q}(\Delta Y(s)\neq 0)=\mathbb{P}(\Delta Y(s)\neq 0)=0\end{aligned}$$

for any $s>0$, which follows generally by continuous waiting time distributions. Further, the set of jump times

$$\begin{aligned}\mathscr{T}^X(t)&:=\{s\in(0,t]:=\Delta X(s)\neq 0\},\\ \mathscr{T}^Y(t)&:=\{s\in(0,t]:=\Delta Y(s)\neq 0\}\end{aligned}$$

are both countable $\mathbb{Q}$-a.s. by the non-explosiveness assumption. We denote the counting process of common jumps in $(0,t]$ as

$$N(t):=\sum_{s\in\mathscr{T}^X(t)}\mathbb{1}\{\Delta Y(s)\neq 0\}.$$

Hence we have that

$$\mathbb{Q}\left(\exists s\in(0,t]\colon\Delta X(s)\neq 0,\Delta Y(s)\neq 0\right)=\mathbb{Q}\left(N(t)>0\right).$$

By $N(t)\geq 0$ it suffices to show $\mathbb{E}_\mathbb{Q}[N(t)]=0$ in order to show weak bipartiteness, restricted to the interval $[0,t]$.

$$\begin{aligned}\mathbb{E}_\mathbb{Q}[N(t)]=&\mathbb{E}_\mathbb{Q}\left[\mathbb{E}_\mathbb{Q}\left[N(t)\middle|\mathscr{F}^X_t\right]\right]\\ =&\mathbb{E}_\mathbb{Q}\left[\sum_{s\in\mathscr{T}^X(t)}\mathbb{E}_\mathbb{Q}\left[\mathbb{1}\{\Delta Y(s)\neq 0\}\middle|\mathscr{F}^X_t\right]\right]\\ =&\mathbb{E}_\mathbb{Q}\left[\sum_{s\in\mathscr{T}^X(t)}\mathbb{P}\left(\Delta Y(s)\neq 0\right)\right]=0.\end{aligned}$$

Since the result holds for all $t>0$, let $(t_n)_{n\in\mathbb{N}}$ be a strictly increasing sequence on $(0,\infty)$ with $\lim_{n\to\infty}t_n=\infty$ and note that $\{N(t_n)>0\}\subset\{N(t_{n+1})>0\}$ as well as

$$\{\exists t>0\colon\Delta X(s)\neq 0,\Delta Y(s)\neq 0\}=\bigcup_{n\in\mathbb{N}}\{N(t_n)>0\}.$$

By continuity from below of measures, we then have

$$\mathbb{Q}\left(\exists t>0\colon\Delta X(s)\neq 0,\Delta Y(s)\neq 0\right)=\lim_{n\to\infty}\mathbb{Q}\left(N(t_n)>0\right)=0.$$

Hence, both, under $\mathbb{P}$ in the strongly bipartite case and under $\mathbb{Q}$ in every case the random counting measure associated with $(X,Y)$ almost surely has the "strongly bipartite" form

$$\begin{aligned}P(C,s):=&\delta_{(X(0),Y(0))}(C)+\int_0^t\mathbb{1}\{(X(s),Y(s))\in C\}\,\mathrm{d}N^X_s\\ &+\int_0^t\mathbb{1}\{(X(s),Y(s))\in C\}\,\mathrm{d}N^Y_s\end{aligned}$$

for all $C\in\Sigma_X\otimes\Sigma_Y$ and $s\geq 0$. That is, in the strongly bipartite case it holds that $P_1=P$, $\mathbb{P}$-a.s., and the $(\mathscr{F}^{XY},\mathbb{P})$-intensity kernel can be rewritten as

$$\begin{aligned}\hat{g}_1(x,y,s):=&\mathbb{1}\{Y(s^-)=y\}\sum_{v_X\in\mathscr{V}_X}\lambda^X_s(v_X)\mathbb{1}\{v_X=x-X(s^-)\}\\ &+\mathbb{1}\{X(s^-)=x\}\sum_{v_Y\in\mathscr{V}_Y}\lambda^Y_s(v_Y)\mathbb{1}\{v_Y=y-Y(s^-)\}\end{aligned}$$

for all $s > 0$. As the compensating measure under $\mathbb{Q}$ we propose

$$\tilde{P}(C,t) := \int_{C\times[0,t]} \hat{g}(x,y,s)\,\mathrm{d}\mu(x,y,s)$$

for all $C \in \Sigma_X \otimes \Sigma_Y$ with the $(\mathscr{F}^{XY},\mathbb{Q})$-intensity kernel

$$\begin{aligned}\hat{g}(x,y,s) :=\ & \mathbb{1}\{Y(s^-)=y\}\textstyle\sum_{v_X\in\mathscr{V}_X} \hat{\lambda}_s^X(v_X)\\ &\qquad\times\mathbb{1}\{v_X = x - X(s^-)\}\\ &+\mathbb{1}\{X(s^-)=x\}\textstyle\sum_{v_Y\in\mathscr{V}_Y} \hat{\lambda}_s^Y(v_Y)\\ &\qquad\times\mathbb{1}\{v_Y = y - Y(s^-)\}\\ =\ & \mathbb{1}\{Y(s^-)=y\}\hat{g}^X(x,s) + \mathbb{1}\{X(s^-)=x\}\hat{g}^Y(y,s)\end{aligned}$$

for all $s > 0$ and $\hat{g}(x,y,0) := \mathbb{P}(X(0)=x)\mathbb{P}(Y(0)=y) =: p_0^X(x)p_0^Y(y) = \hat{g}^X(x,0)\hat{g}^Y(y,0)$, such that $Q(C,t) := P(C,\cdot) - \tilde{P}(C,\cdot)$ is a $(\mathscr{F}^{XY},\mathbb{Q})$-local martingale. For brevity, we only provide some hints on how to prove this local martingale property.

Considering the jump times $(\tau_n^{XY})_{n\in\mathbb{Z}_{\geq 0}}$ of $(X,Y)$ with $\tau_0^{XY} = 0$, $\mathbb{Q}$-a.s. First, note that

$$\mathbb{E}[Q(C,0)] = \mathbb{E}_{\mathbb{Q}}\left[\delta_{(X(0),Y(0))}(C) - \sum_{(x,y)\in C} p_0^X(x)p_0^Y(y)\right] = 0.$$

Secondly, note that for $t > 0$

$$\begin{aligned}&\mathbb{E}_{\mathbb{Q}}\left[\int_0^t \mathbb{1}\{(X(s),Y(s))\in C\}\,\mathrm{d}N_s^X\right] = \sum_{v_X\in\mathscr{V}_X}\mathbb{E}_{\mathbb{Q}}\left[\int_0^t \mathbb{1}\{(X(s),Y(s))\in C\}\,\mathrm{d}R_s^X(v_X)\right]\\ &= \sum_{v_X\in\mathscr{V}_X}\mathbb{E}_{\mathbb{Q}}\left[\int_0^t \sum_{(x,y)\in C}\mathbb{1}\{Y(s^-)=y\}\mathbb{1}\{v_X = x - X(s^-)\}\,\mathrm{d}R_s^X(v_X)\right]\\ &= \sum_{v_X\in\mathscr{V}_X}\mathbb{E}_{\mathbb{Q}}\left[\int_0^t \sum_{(x,y)\in C}\mathbb{1}\{Y(s^-)=y\}\mathbb{1}\{v_X = x - X(s^-)\}\hat{\lambda}_s^X(v_X)\,\mathrm{d}s\right]\\ &=\mathbb{E}_{\mathbb{Q}}\left[\int_0^t \sum_{(x,y)\in C}\mathbb{1}\{Y(s^-)=y\}\sum_{v_X\in\mathscr{V}_X}\mathbb{1}\{v_X = x - X(s^-)\}\hat{\lambda}_s^X(v_X)\,\mathrm{d}s\right]\\ &=\mathbb{E}_{\mathbb{Q}}\left[\int_{C\times(0,t]}\mathbb{1}\{Y(s^-)=y\}\hat{g}^X(x,s)\,\mathrm{d}\mu(x,y,s)\right],\end{aligned} \tag{S1.9}$$

where the second and third equality follow by virtue of [6, Thm. 18.6] and the fact that $\hat{\lambda}_s^X(v_X)$ is the $(\mathscr{F}^{XY},\mathbb{Q})$-intensity process of $R^X(v_X)$ since by (S1.3) we have that

$$\begin{aligned}&\mathbb{E}_{\mathbb{Q}}\left[R^X_{t\wedge\tau_n^X}(v_X) - \int_0^{t\wedge\tau_n^X}\hat{\lambda}_{s'}^X(v_X)\,\mathrm{d}s' \,\middle|\, \mathscr{F}_s^{XY}\right]\\ =&\mathbb{E}\left[R^X_{t\wedge\tau_n^X}(v_X) - \int_0^{t\wedge\tau_n^X}\hat{\lambda}_{s'}^X(v_X)\,\mathrm{d}s' \,\middle|\, \mathscr{F}_s^{X}\right]\\ =&R^X_{s\wedge\tau_n^X}(v_X) - \int_0^{s\wedge\tau_n^X}\hat{\lambda}_{s'}^X(v_X)\,\mathrm{d}s'\end{aligned}$$

with $(\tau_n^X)_{n\in\mathbb{N}}$ being the $\mathscr{F}^{XY}$-jump times of $X$. With the analogous equation for the $Y$-related terms, this implies $\mathbb{E}_{\mathbb{Q}}[Q(C,t)] = 0$ for all $t \geq 0$. The local martingale property then follows along similar lines via

$$\begin{aligned}&\mathbb{E}_{\mathbb{Q}}\left[Q(C,t\wedge\tau_n^{XY})\middle|\mathscr{F}_s^{XY}\right]\\ =&\mathbb{E}_{\mathbb{Q}}\left[Q(C,t\wedge\tau_n^{XY}) - Q(C,s\wedge\tau_n^{XY})\middle|\mathscr{F}_s^{XY}\right]\\ &+Q(C,s\wedge\tau_n^{XY}).\end{aligned}$$

and noting that the Theorem used for (S1.9) also implies

$$\mathbb{E}_{\mathbb{Q}}\left[Q(C,t\wedge\tau_n^{XY}) - Q(C,s\wedge\tau_n^{XY})\middle|\mathscr{F}_s^{XY}\right] = 0, \quad \mathbb{Q}\text{-a.s.}$$

*Step 3:*
We now show that the $\mathbb{P} \ll \mathbb{Q}$ on $\mathscr{F}^{XY}$ by virtue of [3, Thm. 3.3]. The $\mathscr{F}^{XY}$-compensating measure under $\mathbb{P}$ can be expressed as

$$\begin{aligned}\tilde{P}_1^{XY}(C,t) &= \int_{C\times[0,t]} \hat{g}_1(x,y,s)\,\mathrm{d}\mu(x,y,s)\\ &= \int_{C\times[0,t]} \frac{\hat{g}_1(x,y,s)}{\hat{g}(x,y,s)}\frac{\hat{g}(x,y,s)}{\sum_{x',y'}\hat{g}(x,y,s)}\sum_{\check{x},\check{y}}\hat{g}(\check{x},\check{y},s)\,\mathrm{d}\mu(x,y,s)\\ &= \int_{[0,t]}\sum_{(x,y)\in C}\frac{\hat{g}_1(x,y,s)}{\hat{g}(x,y,s)}\frac{\hat{g}(x,y,s)}{\sum_{x',y'}\hat{g}(x,y,s)}\,\tilde{P}(\mathscr{X}\times\mathscr{Y},\mathrm{d}s)\\ &= \int_{C\times[0,t]}\frac{\hat{g}_1(x,y,s)}{\hat{g}(x,y,s)}\,\tilde{P}((\mathrm{d}x,\mathrm{d}y),\mathrm{d}s).\end{aligned}$$

Hence, $(X,Y)$ has the intrinsic local description [3, §2.12] $\big(\hat{n}_1, \tilde{P}(\mathscr{X}\times\mathscr{Y},\cdot)\big)$ under $\mathbb{P}$ with

$$\hat{n}_1(C,s) := \int_C \frac{\hat{g}_1(x,y,s)}{\hat{g}(x,y,s)}\,\hat{n}(\mathrm{d}x,\mathrm{d}y,s),$$

$$\hat{n}(C,s) := \sum_{(x,y)\in C}\frac{\hat{g}(x,y,s)}{\sum_{x',y'}\hat{g}(x,y,s)}$$

for all $C \in \Sigma_X \otimes \Sigma_Y$ and $\left(\hat{n}, \tilde{P}(\mathscr{X} \times \mathscr{Y}, \cdot)\right)$ being the intrinsic local description of $(X,Y)$ under $\mathbb{Q}$. Then, the absolute continuity $\mathbb{P} \ll \mathbb{Q}$ follows by [3, Thm. 3.3], where $\mathbb{E}_{\mathbb{Q}}[L_t] = 1$ can be shown by expressing the expectation as an integral on the space of marked point process sequences and using the likelihood of $P$ under $\mathbb{Q}$ (cf. [7, Prop. 7.3.III.]) for the change of measure.

Now all conditions for Theorem 4.1 in [3] are satisfied, such that

$$
\begin{aligned}
&\ln(L_t)\\
=&\ln\left(\frac{p_0^{XY}(X(0),Y(0))}{p_0^X(X(0))p_0^Y(Y(0))}\right) + \int_0^t \ln\left(\frac{\hat{g}_1(X(s),Y(s),s)}{\hat{g}(X(s),Y(s),s)}\right)\mathrm{d}N^{XY}(s) - \int_{\mathscr{X}\times\mathscr{Y}\times[0,t]} \left(\hat{g}_1(x,y,s) - \hat{g}(x,y,s)\right)\mathrm{d}\mu(x,y,s)\\
=&\ln\left(\frac{p_0^{XY}(X(0),Y(0))}{p_0^X(X(0))p_0^Y(Y(0))}\right) - \left(p_0^{XY}(X(0),Y(0)) - p_0^X(X(0)p_0^Y(Y(0)))\right)\\
&+\int_0^t \ln\left(\frac{\sum_{\nu_X\in\mathscr{V}_X}\lambda_s^X(\nu_X)\,\mathbb{1}\{\nu_X = X(s)-X(s^-)\}}{\sum_{\nu_X\in\mathscr{V}_X}\hat{\lambda}_s^X(\nu_X)\,\mathbb{1}\{\nu_X = X(s)-X(s^-)\}}\right)\mathrm{d}N^X(s) + \int_0^t \ln\left(\frac{\sum_{\nu_Y\in\mathscr{V}_Y}\lambda_s^Y(\nu_Y)\,\mathbb{1}\{\nu_Y = Y(s)-Y(s^-)\}}{\sum_{\nu_Y\in\mathscr{V}_Y}\hat{\lambda}_s^Y(\nu_Y)\,\mathbb{1}\{\nu_Y = Y(s)-Y(s^-)\}}\right)\mathrm{d}N^Y(s)\\
&-\sum_{x\in\mathscr{X}}\int_0^t \sum_{\nu_X\in\mathscr{V}_X} \mathbb{1}\{\nu_X = x - X(s^-)\}\left(\lambda_s^X(\nu_X) - \hat{\lambda}_s^X(\nu_X)\right)\mathrm{d}s - \sum_{y\in\mathscr{Y}}\int_0^t \sum_{\nu_Y\in\mathscr{V}_Y} \mathbb{1}\{\nu_Y = y - Y(s^-)\}\left(\lambda_s^Y(\nu_Y) - \hat{\lambda}_s^Y(\nu_Y)\right)\mathrm{d}s\\
=&\ln\left(\frac{p_0^{XY}(X(0),Y(0))}{p_0^X(X(0))p_0^Y(Y(0))}\right) - \left(p_0^{XY}(X(0),Y(0)) - p_0^X(X(0))p_0^Y(Y(0))\right)\\
&+\sum_{\nu_X\in\mathscr{V}_X}\int_0^t \ln\left(\frac{\lambda_s^X(\nu_X)}{\hat{\lambda}_s^X(\nu_X)}\right)\mathrm{d}N^X(s) - \int_0^t \sum_{\nu_X\in\mathscr{V}_X}\left(\lambda_s^X(\nu_X) - \hat{\lambda}_s^X(\nu_X)\right)\mathrm{d}s\\
&+\sum_{\nu_Y\in\mathscr{V}_Y}\int_0^t \ln\left(\frac{\lambda_s^Y(\nu_Y)}{\hat{\lambda}_s^Y(\nu_Y)}\right)\mathrm{d}N^Y(s) - \int_0^t \sum_{\nu_Y\in\mathscr{V}_Y}\left(\lambda_s^Y(\nu_Y) - \hat{\lambda}_s^Y(\nu_Y)\right)\mathrm{d}s
\end{aligned}
$$

Taking the expectation $\mathbb{E}$ yields the desired expression for the MI if the integrability condition (iii) is satisfied (cf. Proof of Theorem 4.2 in [3]).

*Step 4:*
Lastly, assume that $X$ and $Y$ are not weakly bipartite under $\mathbb{P}$, restricted to $\mathscr{F}^{XY}$. This directly implies $\mathbb{P}$ is not absolutely continuous with respect to $\mathbb{Q}$. The additional condition implies that $\mathbb{P}^{XY}_{[0,t]} \not\ll \mathbb{Q}^{XY}_{[0,t]}$ for all $t > 0$. Since $X$ and $Y$ are not weakly bipartite, they are also not strongly bipartite. Hence, exists $r \in |\mathscr{R}|$, such that $\pi_X(\nu_r), \pi_Y(\nu_r) \neq 0$. Then

$$\{\exists s \in (0,t] \colon \Delta X(s) \neq 0, \Delta Y(s) \neq 0\} \supseteq \{R_r(t) > 0\}$$

and since $\mathbb{E}[R_r(t)] > 0$ for all $t > 0$ we have

$$0 < \mathbb{P}(R_r(t)) \leq \mathbb{P}\left(\exists s \in (0,t] \colon \Delta X(s) \neq 0, \Delta Y(s) \neq 0\right)$$

for all $t > 0$, but $\mathbb{Q}\left(\exists s \in (0,t] \colon \Delta X(s) \neq 0, \Delta Y(s) \neq 0\right) = 0$ for all $t > 0$.

### S1.3. STOCHASTIC FILTERING EQUATIONS FOR THE INTENSITY PROCESSES

In order to compute the MI or DI from the expressions in the Theorems 1 and 2, we need to evaluate the intensity processes $\lambda^X$, $\hat{\lambda}^X$, $\lambda^Y$ and $\hat{\lambda}^Y$. Computing the MI from similar expressions and approximations thereof has been discussed extensively in [8]. If, however, there are $(X,Y)$-, $X$- or $Y$-indistinguishable reactions, then this needs to be accounted for [9] (see also the discussion in Sec. VI). To improve accessibility, we provide the relevant expressions (Eq. (21) and (22) in [9]) in our notation.

We maintain all assumptions from Sec. VI. Let $\mathscr{S}_A \subset \mathscr{S}$ be an arbitrary subset of species indices of the core system U, and its complement $\mathscr{S}_B := \mathscr{S} \setminus \mathscr{S}_A$. The set $\mathscr{S}_A$ represents the set of species with respect to whom's history we want to filter the propensity functions in Eq (105). Hence, we choose $\mathscr{S}_A \in \{\mathscr{S}_X, \mathscr{S}_Y, \mathscr{S}_X \cup \mathscr{S}_Y\}$ in this context. Define $A := \{U_d(s) \colon d \in \mathscr{S}_A\}_{s\geq 0}$ and $B := \{U_d(s) \colon d \in \mathscr{S}_B\}_{s\geq 0}$, together with the natural filtration $\mathscr{F}^A$, as well as the coordinate maps $\pi_A$, $\pi_B$ and the non-zero subnetwork change vectors $\mathscr{V}_A$, $\mathscr{V}_B$. In the following we substitute $U(s) = (A(s), B(s))$.

Exemplary, we want to evaluate the $\mathscr{F}^A$-intensity process of the reaction counter $R^X(\nu_X)$, $\nu_X \in \mathscr{V}_X$:

$$
\begin{aligned}
&\mathbb{E}\left[\lambda^X(\nu_X, U(s^-)) \middle| \mathscr{F}^A_{s-}\right]\\
=&\mathbb{E}\left[\lambda^X(\nu_X, A(s^-), B(s^-)) \middle| \mathscr{F}^A_{s-}\right]\\
=&\sum_b \Pi_b(s^-)\ \lambda^X(\nu_X, A(s^-), b)
\end{aligned}
$$

for all $s \in [0,t]$, where $\Pi_b(s)$ denotes the conditional probability mass function $s \mapsto \Pi_b(s) := \mathbb{P}\left(B(s) = b \middle| \mathscr{F}^A_s\right)$ as a function of time, which is actually a piecewise-deterministic stochastic process on the state space $[0,1]$ for all $b \in \mathbb{Z}_{\geq 0}^{|\mathscr{S}_B|}$. We refer to $\{\Pi_b(s),\, b \in \mathbb{Z}_{\geq 0}^{|\mathscr{S}_B|}\}$ as the filtering distribution at time $s \geq 0$. It follows the stochastic differential equation

$$\begin{aligned}
\mathrm{d}\Pi_b(s) = &\left( -\lambda^A(0, A(s^-), b)\, \Pi_b(s^-) + \sum_{v \in \pi_A^{-1}(0) \cap \pi_B^{-1}(\mathscr{V}_B)} \lambda_{V^{-1}(v)}(A(s^-), b - \pi_B(v))\, \Pi_{b-\pi_B(v)}(s^-) \right) \mathrm{d}s \\
&- \sum_{v_A \in \mathscr{V}_A} \left( \lambda^A(v_A, A(s^-), b) - \hat{\lambda}_s^A(v_A) \right) \Pi_b(s^-)\, \mathrm{d}s \\
&+ \sum_{v_A \in \mathscr{V}_A} \left( \frac{\sum_{v \in \pi_A^{-1}(v_A)} \lambda_{V^{-1}(v)}(A(s^-), b - \pi_B(v))\, \Pi_{b-\pi_B(v)}(s^-)}{\hat{\lambda}_s^A(v_A)} - \Pi_b(s^-) \right) \mathrm{d}R_s^A(v_A)
\end{aligned}$$

such that $\Pi_b(s) = \int_0^s \mathrm{d}\Pi_b(u)$, where

$$\begin{aligned}
\lambda^A(v_A, a, b) &:= \sum_{v \in \pi_A^{-1}(v_A)} \lambda_{V^{-1}(v)}(a, b) \\
\hat{\lambda}_s^A(v_A) &:= \mathbb{E}\left[ \lambda^A(v_A, A(s^-), B(s^-)) \middle| \mathscr{F}_{s-}^A \right] \\
&= \sum_b \lambda^A(v_A, A(s^-), b)\, \Pi_b(s^-)
\end{aligned}$$

Importantly, the stochastic updates at jumps of the counters $R^A(v_A), v_A \in \mathscr{V}_A$ depend on the correct conditioning. Using the reaction counters $R_r, r \in \mathscr{R}^X$ (cf. Eq. (VI)) instead would add the surplus information, which of some $X$-indistinguishable reaction has happened. In particular, the enumerator and denominator of the update equation satisfy

$$\sum_{v \in \pi_A^{-1}(v_A)} \lambda_{V^{-1}(v)}(A(s^-), b - \pi_B(v))\, \Pi_{b-\pi_B(v)}(s^-)$$

$$= \sum_{r \in V^{-1}(\pi_A^{-1}(v_A))} \lambda_r(A(s^-), b - \pi_B(v_r))\, \Pi_{b-\pi_B(v_m)}(s^-)$$

$$\hat{\lambda}_s^A(v_A) = \sum_{r \in V^{-1}(\pi_A^{-1}(v_A))} \sum_b \lambda_r(A(s^-), b) \Pi_b(s^-)$$

with $V^{-1}(\pi_A^{-1}(v_A)) \subseteq \mathscr{R}^X$ being a set of $X$-indistinguishable reaction indices. The continuous part of the evolution equation does not depend on the choice of conditioning, discussed here [9].

### S1.4. INTEGRAL VERSION OF DIRECTED INFORMATION

Weissman *et al.*, in Proposition 2 of [10], provided conditions for the existence of the right derivative of the Massey directed information (DI) for paths on $[0,t)$, along with an explicit expression. They informally stated an "integral version" of this Proposition, implying that $s \mapsto \mathbb{I}_M(X_{[0,s)} \to Y_{[0,s)})$ be absolutely continuous. However, even if the conditions for the right derivative are satisfied for all $s \in [0,t)$, discontinuities in the Massey DI may still occur. In such cases, the Massey DI is not absolutely continuous if the left derivative fails to exist somewhere in the interval. For example, consider a piecewise constant process with càdlàg paths that exhibit deterministic jump times, but random values between jumps: the right derivative exists and is zero everywhere, yet the left derivative does not exist at the jump times, violating the uniform convergence condition in an analogous left-sided version of Proposition 2 in [10]. Additionally, if the random paths lack a fixed deterministic initial value, the Massey DI may have a positive value at $s = 0$, in accordance with the fundamental theorem of calculus. In what follows, we build on the ideas in the original proofs to establish rigorous conditions for the existence of the common derivatives of $s \mapsto \mathbb{I}_M(X_{[0,s)} \to Y_{[0,s)})$. Alongside, we establish analogous results for the generalization of Newton's DI (95) via a similar structural, supremum-based approach and also for the continuous-time instantaneous information exchange (98).

The following lemma is analogous to Proposition 1 in [10], but also extends it to include the relevant results for Newton's DI and instantaneous information exchange. It motivates the respective infimum- or supremum-based definitions. For brevity we denote

$$\begin{aligned}
I_{X\to Y}(\boldsymbol{t}) &:= \sum_{i=2}^{n(\boldsymbol{t})} \mathbb{I}(Y_{[t_{i-1},t_i)}; X_{[0,t_{i-1})} \mid Y_{[0,t_{i-1})}) \\
I_{Y\to X}(\boldsymbol{t}) &:= \sum_{i=2}^{n(\boldsymbol{t})} \mathbb{I}(X_{[t_{i-1},t_i)}; Y_{[0,t_{i-1})} \mid X_{[0,t_{i-1})}) \\
I_{X\leftrightarrow Y}(\boldsymbol{t}) &:= \sum_{i=1}^{n(\boldsymbol{t})} \mathbb{I}(Y_{[t_{i-1},t_i)}; X_{[t_{i-1},t_i)} \mid Y_{[0,t_{i-1})}, X_{[0,t_{i-1})})
\end{aligned}$$

**Lemma S1.1.** *Let $\boldsymbol{t}, \boldsymbol{t}' \in \mathscr{T}(0,t)$ be partitions of $[0,t]$, such that $\boldsymbol{t}'$ is a refinement of $\boldsymbol{t}$, i.e., exists a subsequence $(n_m)_{m\in\{1,\dots,n(\boldsymbol{t})\}}$ of $(1,\dots,n(\boldsymbol{t}'))$, such that $t_m = t'_{n_m}$ for all m. Then*

$$\sum_{i=1}^{n(\boldsymbol{t}')} \mathbb{I}(Y_{[t'_{i-1},t'_i)}; X_{[0,t'_i)} \mid Y_{[0,t'_{i-1})}) \leq \sum_{i=1}^{n(\boldsymbol{t})} \mathbb{I}(Y_{[t_{i-1},t_i)}; X_{[0,t_i)} \mid Y_{[0,t_{i-1})}),$$

$$I_{X\to Y}(\boldsymbol{t}') \geq I_{X\to Y}(\boldsymbol{t}) \quad \textit{and} \quad I_{X\leftrightarrow Y}(\boldsymbol{t}') \leq I_{X\leftrightarrow Y}(\boldsymbol{t}).$$

□

*Proof.* Let $\boldsymbol{t} \in \mathscr{T}(0,t)$. It suffices to prove the claims for $\boldsymbol{t}' \in \mathscr{T}(0,t)$, such that $t'_i = t_j$ for $j \le i-1$ and $t'_{i+i} = t_j$ for $j \ge i$, with $t'_i \in (t_{i-1}, t_i)$. For brevity, denote $s_1 := t_{i-1}$, $s_2 := t'_i$ and $s_3 := t_i$.

Further let $A_1 := X_{[0,s_1)}$, $A_2 := X_{[s_1,s_2)}$, $A_3 := X_{[s_2,s_3)}$, and analogously $B_1 := Y_{[0,s_1)}$, $B_2 := Y_{[s_1,s_2)}$, $B_3 := Y_{[s_2,s_3)}$. In the following we make use of the chain rule of information and the non-negativity of the (conditional) mutual information.

(A)

$$\sum_{i=1}^{n(\boldsymbol{t})} \mathbb{I}(Y_{[t_{i-1},t_i)}; X_{[0,t_i)} \mid Y_{[0,t_{i-1})}) - \sum_{i=1}^{n(\boldsymbol{t}')} \mathbb{I}(Y_{[t'_{i-1},t'_i)}; X_{[0,t'_i)} \mid Y_{[0,t'_{i-1})})$$

$$\begin{aligned}
=&\mathbb{I}(A_1,A_2,A_3;B_2,B_3 \mid B_1) \\
&- [\mathbb{I}(A_1,A_2;B_2 \mid B_1) + \mathbb{I}(A_1,A_2,A_3;B_3 \mid B_1,B_2)] \\
=&\mathbb{I}(A_1,A_2,A_3;B_3 \mid B_1,B_2) + \mathbb{I}(A_1,A_2,A_3;B_2 \mid B_1) \\
&- [\mathbb{I}(A_1,A_2;B_2 \mid B_1) + \mathbb{I}(A_1,A_2,A_3;B_3 \mid B_1,B_2)] \\
=&\mathbb{I}(A_1,A_2,A_3;B_2 \mid B_1) - \mathbb{I}(A_1,A_2;B_2 \mid B_1) \\
=&\mathbb{I}(A_3;B_2 \mid B_1,A_1,A_2) + \mathbb{I}(A_1,A_2;B_2 \mid B_1) \\
&- \mathbb{I}(A_1,A_2;B_2 \mid B_1) \\
=&\mathbb{I}(A_3;B_2 \mid B_1,A_1,A_2) \ge 0
\end{aligned}$$

(B)

$$\begin{aligned}
&\sum_{i=2}^{n(\boldsymbol{t})} \mathbb{I}(Y_{[t_{i-1},t_i)}; X_{[0,t_{i-1})} \mid Y_{[0,t_{i-1})}) \\
&- \sum_{i=2}^{n(\boldsymbol{t}')} \mathbb{I}(Y_{[t'_{i-1},t'_i)}; X_{[0,t'_{i-1})} \mid Y_{[0,t'_{i-1})}) \\
=&\mathbb{I}(A_1;B_2,B_3 \mid B_1) \\
&- [\mathbb{I}(A_1;B_2 \mid B_1) + \mathbb{I}(A_1,A_2;B_3 \mid B_1,B_2)] \\
=&\mathbb{I}(A_1;B_2 \mid B_1) + \mathbb{I}(A_1;B_3 \mid B_1,B_2) \\
&- [\mathbb{I}(A_1;B_2 \mid B_1) + \mathbb{I}(A_1,A_2;B_3 \mid B_1,B_2)] \\
=&\mathbb{I}(A_1;B_3 \mid B_1,B_2) - \mathbb{I}(A_1,A_2;B_3 \mid B_1,B_2) \\
=&\mathbb{I}(A_1;B_3 \mid B_1,B_2) \\
&- [\mathbb{I}(A_1;B_3 \mid B_1,B_2) + \mathbb{I}(A_2;B_3 \mid B_1,B_2,A_1)] \\
\le& 0
\end{aligned}$$

(C)

$$\begin{aligned}
&\sum_{i=1}^{n(\boldsymbol{t})} \mathbb{I}(Y_{[t_{i-1},t_i)}; X_{[t_{i-1},t_i)} \mid Y_{[0,t_{i-1})}, X_{[0,t_{i-1})}) \\
&- \sum_{i=1}^{n(\boldsymbol{t}')} \mathbb{I}(Y_{[t'_{i-1},t'_i)}; X_{[t'_{i-1},t'_i)} \mid Y_{[0,t'_{i-1})}, X_{[0,t'_{i-1})}) \\
=&\mathbb{I}(A_2,A_3;B_2,B_3 \mid A_1,B_1) \\
&- [\mathbb{I}(A_2;B_2 \mid A_1,B_1) + \mathbb{I}(A_3;B_3 \mid A_1,A_2,B_1,B_2)] \\
=&\mathbb{I}(A_3;B_3 \mid A_1,A_2,B_1,B_2) + \mathbb{I}(A_3;B_2 \mid A_1,A_2,B_1) \\
&+ \mathbb{I}(A_2;B_3 \mid A_1,B_1,B_2) + \mathbb{I}(A_2;B_2 \mid A_1,B_1) \\
&- [\mathbb{I}(A_2;B_2 \mid A_1,B_1) + \mathbb{I}(A_3;B_3 \mid A_1,A_2,B_1,B_2)] \\
=&\mathbb{I}(A_2;B_3 \mid A_1,B_1,B_2) + \mathbb{I}(A_3;B_2 \mid A_1,A_2,B_1) \ge 0
\end{aligned}$$

□

**Lemma S1.2.** *For every $\delta > 0$ there exists a partition $\boldsymbol{t} \in \mathscr{T}(0,t)$ such that simultaneously*

$$\left| \mathbb{I}(X_{[0,t)} \to Y_{[0,t)}) - I_{X\to Y}(\boldsymbol{t}) \right| < \delta,$$
$$\left| \mathbb{I}(Y_{[0,t)} \to X_{[0,t)}) - I_{Y\to X}(\boldsymbol{t}) \right| < \delta,$$
$$\left| \mathbb{I}(X_{[0,t)} \leftrightarrow Y_{[0,t)}) - I_{X\leftrightarrow Y}(\boldsymbol{t}) \right| < \delta.$$

*Proof.* Let $\delta > 0$. Now consider $\boldsymbol{t}^{(1)}, \boldsymbol{t}^{(2)}, \boldsymbol{t}^{(3)} \in \mathscr{T}(0,t)$, such that

$$|\mathbb{I}(X_{[0,t)} \to Y_{[0,t)}) - I_{X\to Y}(\boldsymbol{t}^{(1)})| < \delta$$
$$|\mathbb{I}(Y_{[0,t)} \to X_{[0,t)}) - I_{Y\to X}(\boldsymbol{t}^{(2)})| < \delta$$
$$|\mathbb{I}(X_{[0,t)} \leftrightarrow Y_{[0,t)}) - I_{X\leftrightarrow Y}(\boldsymbol{t}^{(3)})| < \delta$$

By Lemma S1.1, the joined sequence $\boldsymbol{t} := \boldsymbol{t}^{(1)} \vee \boldsymbol{t}^{(2)} \vee \boldsymbol{t}^{(3)}$ simultaneously satisfies the inequalities. □

Note that Lemma S1.2 implies

$$\begin{aligned}
&\sup_{t\in\mathscr{T}(0,t)} \; (I_{X\to Y}(\boldsymbol{t}) + I_{Y\to X}(\boldsymbol{t})) \\
=&\mathbb{I}(X_{[0,t)} \to Y_{[0,t)}) + \mathbb{I}(Y_{[0,t)} \to X_{[0,t)}).
\end{aligned} \tag{S1.10}$$

The information conservation law for Newton's DI [11] can be generalized to its continuous-time version (97) with the definitions (95) and (98).

**Proposition S1.1.** *If $\mathbb{I}(X_{[0,t)};Y_{[0,t)}) < \infty$, then*

$$\begin{aligned}
\mathbb{I}(X_{[0,t)};Y_{[0,t)}) =&\mathbb{I}(X_{[0,t)} \to Y_{[0,t)}) + \mathbb{I}(Y_{[0,t)} \to X_{[0,t)}) \\
&+ \mathbb{I}\left(X_{[0,t)} \leftrightarrow Y_{[0,t)}\right).
\end{aligned}$$

*Proof.* Let $\boldsymbol{t} \in \mathscr{T}(0,t)$ be arbitrary, apply the discrete-sequence information conservation law [11] and shift the terms of the instantaneous information exchange to the l.h.s. of the equation, i.e.,

$$\mathbb{I}(X_{[0,t)};Y_{[0,t)}) - I_{X\leftrightarrow Y}(\boldsymbol{t}) = I_{X\to Y}(\boldsymbol{t}) + I_{Y\to X}(\boldsymbol{t}).$$

Applying the supremum over $\boldsymbol{t} \in \mathscr{T}(0,t)$ on both sides of the equality, using Lemma S1.2 (cf. (S1.10)) on the r.h.s. and $\sup(-C) = -\inf C$ for some set $C \subseteq \mathbb{R}$ on the l.h.s. yields the desired relation. □

The following lemma resembles a chain rule for directed information. It has also been provided in [10] with a slightly less detailed proof.

**Lemma S1.3.** *Let $0 < s < t$. Then*

$$\begin{aligned}
\mathbb{I}_M(X_{[0,t)} \to Y_{[0,t)}) =&\mathbb{I}_M(X_{[0,s)} \to Y_{[0,s)}) \\
&+ \mathbb{I}_M(X_{[0,t)} \to Y_{[s,t)} \mid Y_{[0,s)}), \\
\mathbb{I}(X_{[0,t)} \to Y_{[0,t)}) =&\mathbb{I}(X_{[0,s)} \to Y_{[0,s)}) \\
&+ \mathbb{I}(X_{[0,t)} \to Y_{[s,t)} \mid Y_{[0,s)})
\end{aligned}$$

*where we denote*

$$\begin{aligned}&\mathbb{I}_M(X_{[0,t)} \to Y_{[s,t)} \mid Y_{[0,s)})\\ &:= \inf_{\boldsymbol{t}\in\mathscr{T}(s,t)} \sum_{i=1}^{n(\boldsymbol{t})} \mathbb{I}(Y_{[t_{i-1},t_i)};X_{[0,t_i)} \mid Y_{[0,t_{i-1})})\\ &\mathbb{I}(X_{[0,t)} \to Y_{[s,t)} \mid Y_{[0,s)})\\ &:= \sup_{\boldsymbol{t}\in\mathscr{T}(s,t)} \sum_{i=2}^{n(\boldsymbol{t})} \mathbb{I}(Y_{[t_{i-1},t_i)};X_{[0,t_{i-1})} \mid Y_{[0,t_{i-1})}).\end{aligned}$$

*Proof.* Consider the function $\Phi\colon \mathscr{T}(0,s)\times\mathscr{T}(s,t)\to\mathscr{T}(0,t)$, $(\boldsymbol{a},\boldsymbol{b})\mapsto(a_1,\ldots,a_{n(\boldsymbol{a})},b_1,\ldots,b_{n(\boldsymbol{b})})$ with $a_{n(\boldsymbol{a})}=s$ and image $\Phi(\mathscr{T}(0,s)\times\mathscr{T}(s,t))$. Then

$$\begin{aligned}&\mathbb{I}_M(X_{[0,t)} \to Y_{[0,t)})\\ &= \inf_{\boldsymbol{t}\in\mathscr{T}(0,t)} \sum_{i=1}^{n(\boldsymbol{t})} \mathbb{I}(Y_{[t_{i-1},t_i)};X_{[0,t_i)} \mid Y_{[0,t_{i-1})})\\ &\le \inf_{\boldsymbol{t}\in\Phi(\mathscr{T}(0,s)\times\mathscr{T}(s,t))} \sum_{i=1}^{n(\boldsymbol{t})} \mathbb{I}(Y_{[t_{i-1},t_i)};X_{[0,t_i)} \mid Y_{[0,t_{i-1})})\\ &=\mathbb{I}_M(X_{[0,s)} \to Y_{[0,s)}) + \mathbb{I}_M(X_{[0,t)} \to Y_{[s,t)} \mid Y_{[0,s)}),\end{aligned}$$

where the inequality follows from $\Phi(\mathscr{T}(0,s)\times\mathscr{T}(s,t))\subseteq\mathscr{T}(0,t)$ and the last equality follows by the additive property of the supremum/infimum [12, Thm. 1.15]. Similarly, we obtain

$$\begin{aligned}&\mathbb{I}(X_{[0,t)} \to Y_{[0,t)})\\ &\ge\mathbb{I}(X_{[0,s)} \to Y_{[0,s)}) + \mathbb{I}(X_{[0,t)} \to Y_{[s,t)} \mid Y_{[0,s)}).\end{aligned}$$

For the reverse inequality consider the partitions $\boldsymbol{t}',\boldsymbol{t}\in\mathscr{T}(0,t)$, such that $\boldsymbol{t}'=\boldsymbol{t}$ if there is a $j$ such that $t_j=s$ and otherwise $n(\boldsymbol{t}')=n(\boldsymbol{t})+1$ and there is a $j$ such that $t'_j=s$, $t'_i=t_i$ for all $i<j$ and $t'_{i+1}=t_i$ for all $i\ge j$. Now define $\Psi\colon\mathscr{T}(0,t)\to\mathscr{T}(0,s)\times\mathscr{T}(s,t)$ such that $(\Phi\circ\Psi)(\boldsymbol{t})=\boldsymbol{t}'$ and note that

$$\begin{aligned}&\mathbb{I}_M(X_{[0,s)} \to Y_{[0,s)}) = \inf_{\boldsymbol{s}\in\mathscr{T}(0,s)} \sum_{i=1}^{n(\boldsymbol{s})} \mathbb{I}(Y_{[s_{i-1},s_i)};X_{[0,s_i)} \mid Y_{[0,s_{i-1})})\\ &= \inf_{\boldsymbol{t}\in\mathscr{T}(0,t)} \sum_{i=1}^{n(\boldsymbol{t}')} \mathbb{I}(Y_{[t'_{i-1},t'_i)};X_{[0,t'_i)} \mid Y_{[0,t'_{i-1})})\mathbb{1}\{t'_i\le s\} \text{ and}\\ &\mathbb{I}_M(X_{[0,t)} \to Y_{[s,t)} \mid Y_{[0,s)})\\ &= \inf_{\boldsymbol{s}\in\mathscr{T}(s,t)} \sum_{i=1}^{n(\boldsymbol{s})} \mathbb{I}(Y_{[s_{i-1},s_i)};X_{[0,s_i)} \mid Y_{[0,s_{i-1})})\\ &= \inf_{\boldsymbol{t}\in\mathscr{T}(0,t)} \sum_{i=1}^{n(\boldsymbol{t}')} \mathbb{I}(Y_{[t'_{i-1},t'_i)};X_{[0,t'_i)} \mid Y_{[0,t'_{i-1})})\mathbb{1}\{t'_i> s\},\end{aligned}$$

where we use $\boldsymbol{t}'$ instead of explicitly writing $(\Phi\circ\Psi)(\boldsymbol{t})$ for brevity. Thus we obtain

$$\begin{aligned}&\mathbb{I}_M(X_{[0,s)} \to Y_{[0,s)}) + \mathbb{I}_M(X_{[0,t)} \to Y_{[s,t)} \mid Y_{[0,s)})\\ &\le \inf_{\boldsymbol{t}\in\mathscr{T}(0,t)} \sum_{i=1}^{n(\boldsymbol{t}')} \mathbb{I}(Y_{[t'_{i-1},t'_i)};X_{[0,t'_i)} \mid Y_{[0,t'_{i-1})})\\ &\le \inf_{\boldsymbol{t}\in\mathscr{T}(0,t)} \sum_{i=1}^{n(\boldsymbol{t})} \mathbb{I}(Y_{[t_{i-1},t_i)};X_{[0,t_i)} \mid Y_{[0,t_{i-1})})\\ &=\mathbb{I}_M(X_{[0,t)} \to Y_{[0,t)}),\end{aligned}$$

where for the first inequality we applied the well-known inequality for the supremum/infimum of the sum of real-valued bounded functions, and for the second we applied Lemma S1.1. The inequality

$$\begin{aligned}&\mathbb{I}(X_{[0,t)} \to Y_{[0,t)})\\ &\le\mathbb{I}(X_{[0,s)} \to Y_{[0,s)}) + \mathbb{I}(X_{[0,t)} \to Y_{[s,t)} \mid Y_{[0,s)}).\end{aligned}$$

follows analogously. □

The central functions for deriving a density of the DI are $\xi_M\colon\mathscr{G}_0\to\mathbb{R}_{\ge0}$ and $\xi\colon\mathscr{G}_0\to\mathbb{R}_{\ge0}$ with $\mathscr{G}_0:=\{(s,h)\in(0,t)\times\mathbb{R}\setminus\{0\}\colon 0<s+h<t\}$ and

$$\xi_M(s,h) := \begin{cases}\frac{1}{h}\mathbb{I}(Y_{[s,s+h)};X_{[0,s+h)} \mid Y_{[0,s)}) & \text{for } h>0\\ -\frac{1}{h}\mathbb{I}(Y_{[s+h,s)};X_{[0,s)} \mid Y_{[0,s+h)}) & \text{for } h<0,\end{cases}$$

$$\xi(s,h) := \begin{cases}\frac{1}{h}\mathbb{I}(Y_{[s,s+h)};X_{[0,s)} \mid Y_{[0,s)}) & \text{for } h>0\\ -\frac{1}{h}\mathbb{I}(Y_{[s+h,s)};X_{[0,s+h)} \mid Y_{[0,s+h)}) & \text{for } h<0.\end{cases}$$

The condition in $\mathscr{G}_0$ serves as a technical restriction on the range of $h$ for different $s$, resulting in a set that has a trapezoidal shape. The definition further implies $\xi_M(s,h)=\xi_M(s+h,-h)$ for all $(s,h)\in\mathscr{G}_0$ and analogously for $\xi$. Let

$$\begin{aligned}\mathscr{D}_M &:= \{s\in(0,t)\colon \lim_{h\to0}\xi_M(s,h) \text{ exists}\},\\ \mathscr{D} &:= \{s\in(0,t)\colon \lim_{h\to0}\xi(s,h) \text{ exists}\}\end{aligned}$$

be the sets, where the $h\mapsto\xi_M(s,h)$ and $h\mapsto\xi(s,h)$ have a continuous extension at $h=0$. With these sets define

$$\begin{aligned}\mathscr{G}_M &:= \{(u,h)\in\mathscr{D}_M\times\mathbb{R}\colon 0<u+h<t\},\\ \mathscr{G} &:= \{(u,h)\in\mathscr{D}\times\mathbb{R}\colon 0<u+h<t\}\end{aligned}$$

and the functions $\overline{\xi}_M\colon\mathscr{G}_M\to\mathbb{R}_{\ge0}$, $\overline{\xi}\colon\mathscr{G}\to\mathbb{R}_{\ge0}$ with

$$\overline{\xi}_M(s,h) = \begin{cases}\xi_M(s,h) & \text{for } h\ne0\\ \lim_{h\to0}\xi_M(s,h) & \text{for } h=0.\end{cases}$$

$$\overline{\xi}(s,h) = \begin{cases}\xi(s,h) & \text{for } h\ne0\\ \lim_{h\to0}\xi(s,h) & \text{for } h=0.\end{cases}$$

We denote the open ball of radius $\varepsilon>0$ around $(s,0)\in\mathscr{G}_M$ or $(s,0)\in\mathscr{G}$, respectively, by

$$U_\varepsilon(s) := \{(u,h)\in(0,t)\times\mathbb{R}\colon \sqrt{h^2+(u-s)^2}<\varepsilon\}.$$

Note that if $(s-\varepsilon', s+\varepsilon') \subseteq \mathscr{D}_M$ for some $\varepsilon' > 0$, then always exists $0 < \varepsilon < \varepsilon'$ with $U_\varepsilon(s) \subseteq \mathscr{G}_M$, and analogously for $\mathscr{D}$ and $\mathscr{G}$.

In simple words, the following theorem states that the derivative of the DI exists at $s \in (0,t)$ if the bivariate function $\overline{\xi}$ exists and is continuous in the neighborhood of $(s,0)$.

**Theorem S1.1.** *(a) Let $s \in \mathscr{D}_M$. If $\varepsilon > 0$ exists, such that $(s-\varepsilon, s+\varepsilon) \subseteq \mathscr{D}_M$, $U_\varepsilon(s) \subseteq \mathscr{G}_M$ and $\overline{\xi}_M$ is continuous on $U_\varepsilon(s)$, then*

$$\frac{\mathrm{d}}{\mathrm{d}s}\mathbb{I}_M(X_{[0,s)} \to Y_{[0,s)}) = \overline{\xi}_M(s,0).$$

*(b) Let $s \in \mathscr{D}$. If $\varepsilon > 0$ exists, such that $(s-\varepsilon, s+\varepsilon) \subseteq \mathscr{D}$, $U_\varepsilon(s) \subseteq \mathscr{G}$ and $\overline{\xi}$ is continuous on $U_\varepsilon(s)$, then*

$$\frac{\mathrm{d}}{\mathrm{d}s}\mathbb{I}(X_{[0,s)} \to Y_{[0,s)}) = \overline{\xi}(s,0).$$

*Proof.* For brevity, we will only prove part (b), as part (a) is analogous. Let $h \in (-\varepsilon, \varepsilon) \setminus \{0\}$. Then $K_{|h|}(s) := \{(u,\delta) \in (0,t) \times \mathbb{R} \colon \sqrt{\delta^2 + (u-s)^2} \leq |h|\} \subset U_\varepsilon(s)$ is a compact subset. By Lemma S1.3 the difference quotient of the DI at $s \in \mathscr{D}$, for any $h$, is given by

$$\frac{1}{h}(\mathbb{I}(X_{[0,s+h)} \to Y_{[0,s+h)}) - \mathbb{I}(X_{[0,s)} \to Y_{[0,s)}))$$
$$= \begin{cases} \frac{1}{|h|}\mathbb{I}(X_{[0,s+h)} \to Y_{[s,s+h)} \mid Y_{[0,s)}) & \text{for } h > 0 \\ \frac{1}{|h|}\mathbb{I}(X_{[0,s)} \to Y_{[s+h,s)} \mid Y_{[0,s+h)}) & \text{for } h < 0. \end{cases}$$

Then we have for $h \in (0, \varepsilon)$

$$\frac{1}{|h|}\mathbb{I}(X_{[0,s+h)} \to Y_{[s,s+h)} \mid Y_{[0,s)})$$
$$= \frac{1}{|h|} \sup_{\boldsymbol{t} \in \mathscr{T}(s,s+h)} \sum_{i=2}^{n(\boldsymbol{t})} \mathbb{I}(Y_{[t_{i-1},t_i)}; X_{[0,t_{i-1})} \mid Y_{[0,t_{i-1})}) \quad \text{(S1.11)}$$
$$= \frac{1}{|h|} \sup_{\boldsymbol{t} \in \mathscr{T}(s,s+h)} \sum_{i=2}^{n(\boldsymbol{t})} (t_i - t_{i-1})\overline{\xi}(t_{i-1}, t_i - t_{i-1})$$

and for $h \in (-\varepsilon, 0)$

$$\frac{1}{|h|}\mathbb{I}(X_{[0,s)} \to Y_{[s+h,s)} \mid Y_{[0,s+h)})$$
$$= \frac{1}{|h|} \sup_{\boldsymbol{t} \in \mathscr{T}(s+h,s)} \sum_{i=2}^{n(\boldsymbol{t})} \mathbb{I}(Y_{[t_{i-1},t_i)}; X_{[0,t_{i-1})} \mid Y_{[0,t_{i-1})}) \quad \text{(S1.12)}$$
$$= \frac{1}{|h|} \sup_{\boldsymbol{t} \in \mathscr{T}(s+h,s)} \sum_{i=2}^{n(\boldsymbol{t})} (t_i - t_{i-1})\overline{\xi}(t_{i-1}, t_i - t_{i-1})$$

By the continuity of $\overline{\xi}$ on all compact sets $K(|h|)$, both $\max_{(\tau,\kappa)\in K(|h|)} \overline{\xi}(\tau,\kappa)$ and $\min_{(\tau,\kappa)\in K(|h|)} \overline{\xi}(\tau,\kappa)$ exist [12] (cf. Theorem 4.28 therein). Since supremum and infimum are taken on the set $K(|h|)$ it follows that $\lim_{h\to 0} \max_{(\tau,\kappa)\in K(|h|)} \overline{\xi}(\tau,\kappa) = \lim_{h\to 0} \min_{(\tau,\kappa)\in K(|h|)} \overline{\xi}(\tau,\kappa) = \overline{\xi}(s,0)$. Noting that $\mathscr{T}(s+\delta, s) = \mathscr{T}(s, s-\delta) = \emptyset$ for any $\delta \in (0,\varepsilon)$ and using (S1.11) and (S1.12) we obtain for any $h \in (-\varepsilon,\varepsilon) \setminus \{0\}$

$$\min_{(\tau,\kappa)\in K(|h|)} \overline{\xi}(\tau,\kappa)$$
$$= \frac{1}{|h|} \sup_{\boldsymbol{t} \in \mathscr{T}(s+h,s) \cup \mathscr{T}(s,s+h)} \sum_{i=2}^{n(\boldsymbol{t})} (t_i - t_{i-1}) \min_{(\tau,\kappa)\in K(|h|)} \overline{\xi}(\tau,\kappa)$$
$$\leq \frac{1}{|h|} \sup_{\boldsymbol{t} \in \mathscr{T}(s+h,s) \cup \mathscr{T}(s,s+h)} \sum_{i=2}^{n(\boldsymbol{t})} (t_i - t_{i-1})\overline{\xi}(t_{i-1}, t_i - t_{i-1})$$
$$= \frac{1}{h}\left(\mathbb{I}(X_{[0,s+h)} \to Y_{[0,s+h)}) - \mathbb{I}(X_{[0,s)} \to Y_{[0,s)})\right)$$
$$\leq \frac{1}{|h|} \sup_{\boldsymbol{t} \in \mathscr{T}(s+h,s) \cup \mathscr{T}(s,s+h)} \sum_{i=2}^{n(\boldsymbol{t})} (t_i - t_{i-1}) \max_{(\tau,\kappa)\in K(|h|)} \overline{\xi}(\tau,\kappa)$$
$$= \max_{(\tau,\kappa)\in K(|h|)} \overline{\xi}(\tau,\kappa).$$

Taking the limit $h \to 0$ on both sides of the inequality finishes the proof. □

We complete the section by providing the desired "integral version".

**Corollary S1.1.** *(a) If $\mathscr{D}_M = (0,t)$ and $\overline{\xi}_M$ is continuous on $\mathscr{G}_M$, then*

$$\mathbb{I}_M(X_{[0,t)} \to Y_{[0,t)})$$
$$= \mathbb{I}(Y_0; X_0) + \int_0^t \lim_{h\searrow 0} \frac{1}{h}\mathbb{I}(Y_{[s,s+h)}; X_{[0,s+h)} \mid Y_{[0,s)})\,\mathrm{d}s \quad \text{(S1.13)}$$
$$+ \lim_{h\searrow 0} \mathbb{I}(Y_{[t-h,t)}; X_{[0,t)} \mid Y_{[0,t-h)}).$$

*(b) If $\mathscr{D} = (0,t)$ and $\overline{\xi}$ is continuous on $\mathscr{G}$, then*

$$\mathbb{I}(X_{[0,t)} \to Y_{[0,t)}) = \int_0^t \lim_{h\searrow 0} \frac{1}{h}\mathbb{I}(Y_{[s,s+h)}; X_{[0,s)} \mid Y_{[0,s)})\,\mathrm{d}s \quad \text{(S1.14)}$$
$$+ \lim_{h\searrow 0} \mathbb{I}(Y_{[t-h,t)}; X_{[0,t-h)} \mid Y_{[0,t-h)}).$$

*Proof.* By Theorem S1.1 and the fundamental theorem of calculus, we directly obtain

$$\lim_{h\searrow 0}\left(\mathbb{I}_M(X_{[0,t-h)} \to Y_{[0,t-h)}) - \mathbb{I}_M(X_{[0,h)} \to Y_{[0,h)})\right)$$
$$= \int_0^t \lim_{h\searrow 0} \frac{1}{h}\mathbb{I}(Y_{[s,s+h)}; X_{[0,s+h)} \mid Y_{[0,s)})\,\mathrm{d}s.$$

Analogously, we have

$$\lim_{h\searrow 0}\left(\mathbb{I}(X_{[0,t-h)} \to Y_{[0,t-h)}) - \mathbb{I}(X_{[0,h)} \to Y_{[0,h)})\right)$$
$$= \int_0^t \lim_{h\searrow 0} \frac{1}{h}\mathbb{I}(Y_{[s,s+h)}; X_{[0,s)} \mid Y_{[0,s)})\,\mathrm{d}s.$$

We start proving the boundary values for part (a). At the right boundary

$$\begin{aligned}
&\lim_{h\searrow 0}\left(\mathbb{I}_M(X_{[0,t)}\to Y_{[0,t)})-\mathbb{I}_M(X_{[0,t-h)}\to Y_{[0,t-h)})\right)\\
=&\lim_{h\searrow 0}\mathbb{I}_M(X_{[0,t)}\to Y_{[t-h,t)}\mid Y_{[0,t-h)})\\
=&\lim_{h\searrow 0}\inf_{\boldsymbol{t}\in\mathscr{T}(t-h,t)}\sum_{i=1}^{n(\boldsymbol{t})}\mathbb{I}(Y_{[t_{i-1},t_i)};X_{[0,t_i)}\mid Y_{[0,t_{i-1})})\\
\geq&\lim_{h\searrow 0}\inf_{\boldsymbol{t}\in\mathscr{T}(t-h,t)}\mathbb{I}(Y_{[t_{n(\boldsymbol{t})-1},t)};X_{[0,t)}\mid Y_{[0,t_{n(\boldsymbol{t})-1})})\\
=&\lim_{h\searrow 0}\lim_{\delta\searrow 0}\mathbb{I}(Y_{[t-\delta,t)};X_{[0,t)}\mid Y_{[0,t-\delta)}),
\end{aligned}$$

where the first equality follows with Lemma S1.3 and the last equality follows with the fact that $\delta\mapsto\mathbb{I}(Y_{[t-\delta,t)};X_{[0,t)}\mid Y_{[0,t-\delta)})$ is a monotone decreasing function, as can be shown with the chain rule for mutual information. On the other hand

$$\begin{aligned}
&\lim_{h\searrow 0}\inf_{\boldsymbol{t}\in\mathscr{T}(t-h,t)}\sum_{i=1}^{n(\boldsymbol{t})}\mathbb{I}(Y_{[t_{i-1},t_i)};X_{[0,t_i)}\mid Y_{[0,t_{i-1})})\\
\leq&\lim_{h\searrow 0}\inf_{\boldsymbol{t}\in\mathscr{T}(t-h,t)}\mathbb{I}(Y_{[t-h,t)};X_{[0,t)}\mid Y_{[0,t-h)})\\
=&\lim_{h\searrow 0}\mathbb{I}(Y_{[t-h,t)};X_{[0,t)}\mid Y_{[0,t-h)})
\end{aligned}$$

by Lemma S1.1, and setting $\boldsymbol{t}=t_1=t$. At the left boundary we have

$$\begin{aligned}
&\lim_{h\searrow 0}\mathbb{I}_M(X_{[0,h)}\to Y_{[0,h)})\\
=&\lim_{h\searrow 0}\inf_{\boldsymbol{t}\in\mathscr{T}(0,h)}\sum_{i=1}^{n(\boldsymbol{t})}\mathbb{I}(Y_{[t_{i-1},t_i)};X_{[0,t_i)}\mid Y_{[0,t_{i-1})})\\
\geq&\lim_{h\searrow 0}\inf_{\boldsymbol{t}\in\mathscr{T}(0,h)}\mathbb{I}(Y_{[0,t_1)};X_{[0,t_1)})\\
=&\mathbb{I}(X_0;Y_0),
\end{aligned}$$

where we used that $\delta\mapsto\mathbb{I}(X_{[0,\delta)};Y_{[0,\delta)})$ is monotone increasing by the chain rule for mutual information. By Lemma S1.1 and setting $\boldsymbol{t}=t_1=h$ we obtain

$$\begin{aligned}
&\lim_{h\searrow 0}\inf_{\boldsymbol{t}\in\mathscr{T}(0,h)}\sum_{i=1}^{n(\boldsymbol{t})}\mathbb{I}(Y_{[t_{i-1},t_i)};X_{[0,t_i)}\mid Y_{[0,t_{i-1})})\\
\leq&\lim_{h\searrow 0}\inf_{\boldsymbol{t}\in\mathscr{T}(0,h)}\mathbb{I}(Y_{[0,h)};X_{[0,h)})\\
=&\mathbb{I}(X_0;Y_0).
\end{aligned}$$

Now we turn to the boundaries for part (b). For the left boundary we have

$$\begin{aligned}
0\leq&\lim_{h\searrow 0}\mathbb{I}(X_{[0,h)}\to Y_{[0,h)})\\
\leq&\lim_{h\searrow 0}\left(\mathbb{I}(X_{[0,h)};Y_{[0,h)})-\mathbb{I}(X_0;Y_0)\right)\\
=&0,
\end{aligned}$$

where we used non-negativity of the directed information for the first inequality and Proposition S1.1 with $\mathbb{I}(Y_{[0,t)}\to X_{[0,t)})\geq 0$ and $\mathbb{I}\left(X_{[0,t)}\leftrightarrow Y_{[0,t)}\right)\geq\mathbb{I}(X_0;Y_0)$ for the second inequality.

At the right boundary we have that

$$\begin{aligned}
&\lim_{h\searrow 0}\left(\mathbb{I}(X_{[0,t)}\to Y_{[0,t)})-\mathbb{I}(X_{[0,t-h)}\to Y_{[0,t-h)})\right)\\
=&\lim_{h\searrow 0}\mathbb{I}(X_{[0,t)}\to Y_{[t-h,t)}\mid Y_{[0,t-h)})\\
=&\lim_{h\searrow 0}\sup_{\boldsymbol{t}\in\mathscr{T}(t-h,t)}\sum_{i=2}^{n(\boldsymbol{t})}\mathbb{I}(Y_{[t_{i-1},t_i)};X_{[0,t_{i-1})}\mid Y_{[0,t_{i-1})})\\
\geq&\lim_{h\searrow 0}\mathbb{I}(Y_{[t_{n(\boldsymbol{t})-1},t)};X_{[0,t_{n(\boldsymbol{t})-1})}\mid Y_{[0,t_{n(\boldsymbol{t})-1})})\\
=&\lim_{h\searrow 0}\mathbb{I}(Y_{[t-h,t)};X_{[0,t-h)}\mid Y_{[0,t-h)}),
\end{aligned}$$

as well as

$$\begin{aligned}
&\lim_{h\searrow 0}\sup_{\boldsymbol{t}\in\mathscr{T}(t-h,t)}\sum_{i=2}^{n(\boldsymbol{t})}\mathbb{I}(Y_{[t_{i-1},t_i)};X_{[0,t_{i-1})}\mid Y_{[0,t_{i-1})})\\
\leq&\lim_{h\searrow 0}\sup_{\boldsymbol{t}\in\mathscr{T}(t-h,t)}\sum_{i=2}^{n(\boldsymbol{t})}\mathbb{I}(Y_{[t_{i-1},t_i)};X_{[0,t-h)}\mid Y_{[0,t_{i-1})})\\
\leq&\lim_{h\searrow 0}\sup_{\boldsymbol{t}\in\mathscr{T}(t-h,t)}\mathbb{I}(Y_{[t-h,t)};X_{[0,t-h)}\mid Y_{[0,t-h)})\\
=&\lim_{h\searrow 0}\mathbb{I}(Y_{[t-h,t)};X_{[0,t-h)}\mid Y_{[0,t-h)}).
\end{aligned}$$

□

### S1.5. PROOF OF DIRECTED INFORMATION BETWEEN SUBNETWORKS OF MARKOVIAN CRNS

#### A. Newton directed information

In the following we constructively show that the conditions for Equation (S1.14) are satisfied and thereby verify the expressions (111), (112) and (96) for the DI between subnetworks. We start by defining the relevant filtrations $\mathscr{F}^1$, $\mathscr{F}^{X,2}$, $\mathscr{F}^{Y,2}$ such that for any fixed $s\geq 0$ and all $\tau\in[0,s+h)$ they satisfy

$$\mathscr{F}^{X,2}_\tau=\begin{cases}\mathscr{F}^X_\tau\vee\mathscr{F}^Y_{s-} & \text{for } \tau<s\\ \mathscr{F}^{XY}_{s-} & \text{for } \tau\geq s\end{cases}$$

$$\mathscr{F}^{Y,2}_\tau=\begin{cases}\mathscr{F}^Y_{s-} & \text{for } \tau<s\\ \mathscr{F}^Y_\tau & \text{for } \tau\geq s,\end{cases}$$

$$\mathscr{F}^1_\tau=\begin{cases}\mathscr{F}^X_\tau\vee\mathscr{F}^Y_{s-} & \text{for } \tau<s\\ \mathscr{F}^X_{s-}\vee\mathscr{F}^Y_\tau & \text{for } \tau\geq s.\end{cases}$$

The measurable space of trajectories of $(X,Y)_{[0,t]}$ is given by the product space $(\mathscr{X}^{[0,t]}\times\mathscr{Y}^{[0,t]},\Sigma_X^{\otimes[0,t]}\otimes\Sigma_Y^{\otimes[0,t]})$. We define the conditional probability measures $\mathbb{P}^1_{\tau,s}$, $\mathbb{P}^2_{\tau,s}$ such that for all $\tau>s$, $A\in\Sigma_X^{\otimes[0,s)}$ and $B\in\Sigma_Y^{\otimes[s,\tau)}$ it holds that

$$\begin{aligned}
\mathbb{P}^1_{\tau,s}(A\times B;\omega)=&\mathbb{E}[\mathbb{1}\{X_{[0,s)}\in A,\ Y_{[s,\tau)}\in B\}\mid\mathscr{F}^Y_{s-}](\omega)\text{ and}\\
\mathbb{P}^2_{\tau,s}(A\times B;\omega)=&\mathbb{E}[\mathbb{1}\{X_{[0,s)}\in A\}\mid\mathscr{F}^Y_{s-}](\omega)\\
&\otimes\mathbb{E}[\mathbb{1}\{Y_{[s,\tau)}\in B\}\mid\mathscr{F}^Y_{s-}](\omega).
\end{aligned}$$

Again, we assume that $X$ and $Y$ are strongly bipartite. While the r.h.s. of (111) and (112) are well-defined, even if $X$ and $Y$ are not weakly bipartite, i.e., the intensity processes exist for all times, the compensators required in the following proof are not absolutely continuous for all times. This issue may be avoided by defining the DI directly via a Radon-Nikodym derivative of causally conditioned probability measures, similar to the transfer entropy (B2).

In the strongly bipartite case and using the regular version of these conditional probabilities, we obtain for $h > 0$

$$\begin{aligned}
&\mathbb{I}(Y_{[s,s+h)};X_{[0,s)} \mid Y_{[0,s)})\\
=&\mathbb{E}\left[\mathbb{E}^1_{s+h,s}\left[\ln\left(\frac{\mathrm{d}\mathbb{P}^1_{s+h,s}}{\mathrm{d}\mathbb{P}^2_{s+h,s}}(X_{[0,s)},Y_{[s,s+h)};Y_{[0,s)})\right)\right]\right]\\
=&\mathbb{E}\left[\mathbb{E}\left[\ln\left(\frac{\mathrm{d}\mathbb{P}^1_{s+h,s}}{\mathrm{d}\mathbb{P}^2_{s+h,s}}(X_{[0,s)},Y_{[s,s+h)};Y_{[0,s)})\right)\middle|\mathscr{F}^Y_{s-}\right]\right] \qquad \text{(S1.15)}\\
=&\mathbb{E}\left[\ln\left(\frac{\mathrm{d}\mathbb{P}^1_{s+h,s}}{\mathrm{d}\mathbb{P}^2_{s+h,s}}(X_{[0,s)},Y_{[s,s+h)};Y_{[0,s)})\right)\right]\\
=&\int_{\mathscr{Y}^{[0,s)}}\mathbb{E}\left[\ln\left(\frac{\mathrm{d}\mathbb{P}^1_{s+h,s}}{\mathrm{d}\mathbb{P}^2_{s+h,s}}\right)\middle|Y_{[0,s)}=y_{[0,s)}\right]\mathrm{d}\mathbb{P}^Y_{[0,s)}(y_{[0,s)}).
\end{aligned}$$

With Theorem 4.1 in [3], the conditional expectation in (S1.15) obeys

$$\begin{aligned}
&\mathbb{E}\left[\ln\left(\frac{\mathrm{d}\mathbb{P}^1_{s+h,s}}{\mathrm{d}\mathbb{P}^2_{s+h,s}}\right)\middle|\mathscr{F}^Y_{s-}\right]\\
=&\sum_{\nu_X\in\mathscr{V}_X}\int_0^s\mathbb{E}\left[\phi(\lambda^{X,1}_\tau(\nu_X))-\phi(\lambda^{X,2}_\tau(\nu_X))\mid\mathscr{F}^Y_{s-}\right]\mathrm{d}\tau\\
&+\sum_{\nu_Y\in\mathscr{V}_Y}\int_s^{s+h}\mathbb{E}\left[\phi(\lambda^{Y,1}_\tau(\nu_Y))-\phi(\lambda^{Y,2}_\tau(\nu_Y))\mid\mathscr{F}^Y_{s-}\right]\mathrm{d}\tau,
\end{aligned}$$

with newly defined intensity processes

$$\begin{aligned}
\lambda^{X,1}_\tau(\nu_X)=&\mathbb{E}\left[\lambda^X(\nu_X,U(\tau^-))\mid\mathscr{F}^1_{\tau-}\right]\\
\lambda^{X,2}_\tau(\nu_X)=&\mathbb{E}\left[\lambda^X(\nu_X,U(\tau^-))\mid\mathscr{F}^{X,2}_{\tau-}\right]
\end{aligned}$$

for all $\tau \geq 0$ and

$$\begin{aligned}
\lambda^{Y,1}_\tau(\nu_Y)=&\mathbb{E}\left[\lambda^Y(\nu_Y,U(\tau^-))\mid\mathscr{F}^1_{\tau-}\right]\\
\lambda^{Y,2}_\tau(\nu_Y)=&\mathbb{E}\left[\lambda^Y(\nu_Y,U(\tau^-))\mid\mathscr{F}^{Y,2}_{\tau-}\right]
\end{aligned}$$

for all $\tau \geq s$. Now note that $\mathscr{F}^1_\tau = \mathscr{F}^{X,2}_\tau$ for all $\tau < s$, such that for all $\nu_X$ the integrals vanish. Lastly, $\mathscr{F}^1_{s-} = \mathscr{F}^{XY}_{s-}$ and $\mathscr{F}^{Y,2}_\tau = \mathscr{F}^Y_\tau$ for all $\tau \geq s$, which implies $\lambda^{Y,1}_s(\nu_Y) = \lambda^Y_s(\nu_Y)$ and $\lambda^{Y,2}_\tau(\nu_Y) = \hat{\lambda}^Y_\tau(\nu_Y)$ for $\tau \geq s$. Concluding, we have

$$\begin{aligned}
&\lim_{h\searrow 0}\frac{1}{h}\mathbb{I}(Y_{[s,s+h)};X_{[0,s+)]}\mid Y_{[0,s)})\\
=&\sum_{\nu_Y\in\mathscr{V}_Y}\mathbb{E}\left[\phi(\lambda^Y_s(\nu_Y))-\phi(\hat{\lambda}^Y_s(\nu_Y))\right],
\end{aligned}$$

for all $s > 0$. The left-limit follows analogously and coincides with the right limit. Hence, $\overline{\xi}$ is continuous on its whole domain and Eq. (111) follows directly by Corollary S1.1.

#### B. Instantaneous information exchange

Although we did not explicitly prove the "integral version" (cf. Corollary S1.1) for the instantaneous information exchange, such an expression can be derived using the same steps as for the Massey DI. Clearly, it holds that

$$\begin{aligned}
&\mathbb{I}(X_0;Y_0)\\
=&\lim_{h\searrow 0}\inf_{\boldsymbol{t}\in\mathscr{T}(0,h)}\mathbb{I}(X_{[0,t_1)};Y_{[0,t_1)})\\
\leq&\lim_{h\searrow 0}\inf_{\boldsymbol{t}\in\mathscr{T}(0,h)}\sum_{i=1}^{n(\boldsymbol{t})}\mathbb{I}(X_{[t_{i-1},t_i)};Y_{[t_{i-1},t_i)}\mid X_{[0,t_{i-1})},Y_{[0,t_{i-1})})\\
\leq&\lim_{h\searrow 0}\mathbb{I}(X_{[0,h)};Y_{[0,h)})\\
=&\mathbb{I}(X_0;Y_0).
\end{aligned}$$

As for the Newton DI, we now define relevant filtrations $\mathscr{F}^1$, $\mathscr{F}^{X,2}$, $\mathscr{F}^{Y,2}$ such that for any fixed $s \geq 0$ and all $\tau \in [s,s+h)$ they satisfy

$$\mathscr{F}^{X,2}_\tau=\mathscr{F}^X_\tau\vee\mathscr{F}^Y_{s-},\quad \mathscr{F}^{Y,2}_\tau=\mathscr{F}^X_{s-}\vee\mathscr{F}^Y_\tau,\quad \mathscr{F}^1_\tau=\mathscr{F}^{XY}_\tau.$$

Again, we assume that $X$ and $Y$ are strongly bipartite. Define the conditional probability measures $\mathbb{P}^1_{\tau,s}$, $\mathbb{P}^2_{\tau,s}$ such that for all $\tau > s$, $A \in \Sigma_X^{\otimes[s,\tau)}$ and $B \in \Sigma_Y^{\otimes[s,\tau)}$ it holds that

$$\begin{aligned}
\mathbb{P}^1_{\tau,s}(A\times B;\omega)=&\mathbb{E}[\mathbb{1}\{X_{[s,\tau)}\in A,\ Y_{[s,\tau)}\in B\}\mid\mathscr{F}^{XY}_{s-}](\omega)\text{ and}\\
\mathbb{P}^2_{\tau,s}(A\times B;\omega)=&\mathbb{E}[\mathbb{1}\{X_{[s,\tau)}\in A\}\mid\mathscr{F}^{XY}_{s-}](\omega)\\
&\otimes\mathbb{E}[\mathbb{1}\{Y_{[s,\tau)}\in B\}\mid\mathscr{F}^{XY}_{s-}](\omega).
\end{aligned}$$

In short notation, we then have that

$$\mathbb{I}(Y_{[s,s+h)};X_{[s,s+h)}\mid(X,Y)_{[0,s)})=\mathbb{E}\left[\mathbb{E}\left[\ln\left(\frac{\mathrm{d}\mathbb{P}^1_{s+h,s}}{\mathrm{d}\mathbb{P}^2_{s+h,s}}\right)\middle|\mathscr{F}^{XY}_{s-}\right]\right]$$

and with Theorem 4.1 in [3]

$$\begin{aligned}
&\mathbb{E}\left[\ln\left(\frac{\mathrm{d}\mathbb{P}^1_{s+h,s}}{\mathrm{d}\mathbb{P}^2_{s+h,s}}\right)\middle|\mathscr{F}^{XY}_{s-}\right]\\
=&\sum_{\nu_X\in\mathscr{V}_X}\int_s^{s+h}\mathbb{E}\left[\phi(\lambda^{X,1}_\tau(\nu_X))-\phi(\lambda^{X,2}_\tau(\nu_X))\mid\mathscr{F}^{XY}_{s-}\right]\mathrm{d}\tau\\
&+\sum_{\nu_Y\in\mathscr{V}_Y}\int_s^{s+h}\mathbb{E}\left[\phi(\lambda^{Y,1}_\tau(\nu_Y))-\phi(\lambda^{Y,2}_\tau(\nu_Y))\mid\mathscr{F}^{XY}_{s-}\right]\mathrm{d}\tau,
\end{aligned}$$

with newly defined intensity processes

$$\begin{aligned}
\lambda^{X,1}_\tau(\nu_X)=&\mathbb{E}\left[\lambda^X(\nu_X,U(\tau^-))\mid\mathscr{F}^1_{\tau-}\right]\\
\lambda^{X,2}_\tau(\nu_X)=&\mathbb{E}\left[\lambda^X(\nu_X,U(\tau^-))\mid\mathscr{F}^{X,2}_{\tau-}\right]\\
\lambda^{Y,1}_\tau(\nu_Y)=&\mathbb{E}\left[\lambda^Y(\nu_Y,U(\tau^-))\mid\mathscr{F}^1_{\tau-}\right]\\
\lambda^{Y,2}_\tau(\nu_Y)=&\mathbb{E}\left[\lambda^Y(\nu_Y,U(\tau^-))\mid\mathscr{F}^{Y,2}_{\tau-}\right]
\end{aligned}$$

for all $\tau \geq s$. Now its holds that $\lambda_s^{X,1}(\nu_X) = \lambda_s^{X,2}(\nu_X)$ and $\lambda_s^{Y,1}(\nu_Y) = \lambda_s^{Y,2}(\nu_Y)$, which implies

$$\lim_{h \searrow 0} \frac{1}{h} \mathbb{I}(Y_{[s,s+h)}; X_{[s,s+h)]} \mid (X,Y)_{[0,s)}) = 0$$

and hence

$$\mathbb{I}(X_{[0,t)} \leftrightarrow Y_{[0,t)}) = \mathbb{I}(X_0; Y_0)$$

for all $t > 0$.

#### C. Massey directed information

We show that the conditions for Eq. (S1.13) are satisfied and thereby verify expression (V) and the relations presented in Theorem 2 for the Massey DI between subnetworks $X$ and $Y$ that are strongly bipartite. We define the relevant filtrations $\mathscr{F}^1$, $\mathscr{F}^{X,2}$, $\mathscr{F}^{Y,2}$ such that for any fixed $s \geq 0$ and all $\tau \geq 0$ they satisfy

$$\begin{aligned} \mathscr{F}_\tau^{X,2} =& \mathscr{F}_\tau^X \vee \mathscr{F}_{s-}^Y \\ \mathscr{F}_\tau^{Y,2} =& \begin{cases} \mathscr{F}_{s-}^Y & \text{for } \tau < s \\ \mathscr{F}_\tau^Y & \text{for } \tau \geq s, \end{cases} \\ \mathscr{F}_\tau^1 =& \begin{cases} \mathscr{F}_\tau^X \vee \mathscr{F}_{s-}^Y & \text{for } \tau < s \\ \mathscr{F}_\tau^{XY} & \text{for } \tau \geq s. \end{cases} \end{aligned}$$

Now let $\mathbb{P}_{\tau,s}^1$, $\mathbb{P}_{\tau,s}^2$ be the conditional probability measures such that for all $\tau \geq s$, $A \in \Sigma_X^{\otimes[0,\tau)}$ and $B \in \Sigma_Y^{\otimes[s,\tau)}$ it holds that

$\mathbb{P}_{\tau,s}^1(A \times B) = \mathbb{E}[\mathbb{1}\{X_{[0,\tau)} \in A,\, Y_{[s,\tau)} \in B\} \mid \mathscr{F}_{s-}^Y]$ and
$\mathbb{P}_{\tau,s}^2(A \times B) = \mathbb{E}[\mathbb{1}\{X_{[0,\tau)} \in A\} \mid \mathscr{F}_{s-}^Y] \otimes \mathbb{E}[\mathbb{1}\{Y_{[s,\tau)} \in B\} \mid \mathscr{F}_{s-}^Y]$.

Then, for $h > 0$ we obtain

$$\begin{aligned} &\mathbb{I}(Y_{[s,s+h]}; X_{[0,s+h]} \mid Y_{[0,s)}) \\ =& \mathbb{E}\left[\mathbb{E}\left[\ln\left(\frac{\mathrm{d}\mathbb{P}_{s+h,s}^1}{\mathrm{d}\mathbb{P}_{s+h,s}^2}\right) \middle| \mathscr{F}_{s-}^Y\right]\right]. \end{aligned} \tag{S1.16}$$

With Theorem 4.1 in [3], the conditional expectation in (S1.16) obeys

$$\begin{aligned} &\mathbb{E}\left[\ln\left(\frac{\mathrm{d}\mathbb{P}_{s+h,s}^1}{\mathrm{d}\mathbb{P}_{s+h,s}^2}\right) \middle| \mathscr{F}_{s-}^Y\right] \\ =& \sum_{\nu_X \in \mathscr{V}_X} \int_0^{s+h} \mathbb{E}\left[\phi(\lambda_u^{X,1}(\nu_X)) - \phi(\lambda_u^{X,2}(\nu_X)) \mid \mathscr{F}_{s-}^Y\right] \mathrm{d}u \\ &+ \sum_{\nu_Y \in \mathscr{V}_Y} \int_s^{s+h} \mathbb{E}\left[\phi(\lambda_u^{Y,1}(\nu_Y)) - \phi(\lambda_u^{Y,2}(\nu_Y)) \mid \mathscr{F}_{s-}^Y\right] \mathrm{d}u, \end{aligned}$$

with intensity processes

$$\begin{aligned} \lambda_\tau^{X,1}(\nu_X) =& \mathbb{E}\left[\lambda^X(\nu_X, U(\tau^-)) \mid \mathscr{F}_{\tau-}^1\right] \\ \lambda_\tau^{X,2}(\nu_X) =& \mathbb{E}\left[\lambda^X(\nu_X, U(\tau^-)) \mid \mathscr{F}_{\tau-}^{X,2}\right] \end{aligned}$$

for all $\tau \geq 0$ and

$$\begin{aligned} \lambda_\tau^{Y,1}(\nu_Y) =& \mathbb{E}\left[\lambda^Y(\nu_Y, U(\tau^-)) \mid \mathscr{F}_{\tau-}^1\right] \\ \lambda_\tau^{Y,2}(\nu_Y) =& \mathbb{E}\left[\lambda^Y(\nu_Y, U(\tau^-)) \mid \mathscr{F}_{\tau-}^{Y,2}\right] \end{aligned}$$

for all $\tau \geq s$. Now note that $\mathscr{F}_\tau^1 = \mathscr{F}_\tau^{X,2}$ for all $\tau < s$, such that for all $\nu_X$ the integrals vanish on $[0,s]$. Further, $\lim_{h \searrow 0} \mathscr{F}_{s+h}^1 = \mathscr{F}_s^{X,2}$, such that $\mathbb{E}\left[\phi(\lambda_s^{X,1}(\nu_X)) - \phi(\lambda_s^{X,2}(\nu_X)) \mid \mathscr{F}_{s-}^Y\right] = 0$. Hence,

$$\lim_{h \searrow 0} \frac{1}{h} \int_s^{s+h} \mathbb{E}\left[\phi(\lambda_\tau^{X,1}(\nu_X)) - \phi(\lambda_\tau^{X,2}(\nu_X)) \mid \mathscr{F}_{s-}^Y\right] \mathrm{d}\tau = 0$$

Lastly, $\mathscr{F}_\tau^1 = \mathscr{F}_\tau^{XY}$ and $\mathscr{F}_\tau^{Y,2} = \mathscr{F}_\tau^Y$ for all $\tau \geq s$, which implies $\lambda_\tau^{Y,1}(\nu_Y) = \lambda_\tau^Y(\nu_Y)$ and $\lambda_\tau^{Y,2}(\nu_Y) = \hat{\lambda}_\tau^Y(\nu_Y)$. Concluding, we have

$$\begin{aligned} &\lim_{h \searrow 0} \frac{1}{h} \mathbb{I}(Y_{[s,s+h)}; X_{[0,s+h)} \mid Y_{[0,s)}) \\ =& \sum_{\nu_Y \in \mathscr{V}_Y} \mathbb{E}\left[\phi(\lambda_s^Y(\nu_Y)) - \phi(\hat{\lambda}_s^Y(\nu_Y))\right]. \end{aligned}$$

The left-limit follows analogously and coincides with the right limit. Hence, $\overline{\xi}_M$ is continuous on its whole domain and the relations in Theorem 2 follow directly by Corollary S1.1.

### S1.6. PROOFS FOR CRNS AS COMMUNICATION CHANNELS

#### A. Equivalence of message and source code conditioning

For the proof of the equivalence of intensity processes that are conditioned either on the message or the noiseless source encoding, we first show a more general equality of conditional probabilities in Proposition S1.2, which will serve as a lemma for the equality of intensity processes. Therefore, we introduce a short notation for natural filtrations of a subset of reaction processes and a subset of initial values $U_d(0)$, $d \in \mathscr{S}$. We represent the initial values of certain species just by their species index in $\mathscr{S}$. In contrast to the main document, let $\mathscr{R}$ be the set of all reactions and not just the reversible reaction channels. Now we can choose subsets of $\mathscr{R}$ and $\mathscr{S}$ to select exactly those reactions and initial values that we want to keep track of in a natural filtration.

For any set $G \subseteq \mathscr{S} \cup \mathscr{R}$ we assign the subfiltration $\mathscr{F}^G$ such that for all $s \in [0,t]$

$$\mathscr{F}_s^G := \left(\bigvee_{r \in G \cap \mathscr{R}} \sigma(R_r(\tau) \colon \tau \in [0,s])\right) \vee \left(\bigvee_{i \in G \cap \mathscr{S}} \sigma(U_d(0))\right).$$

We further denote $\mathscr{F}_s^{\{\theta\} \cup G} := \sigma(\theta(\tau), \tau \in [0,s)) \vee \mathscr{F}_s^G$ and $\mathscr{F}_s^{\{\mathsf{M}\} \cup G} := \sigma(M) \vee \mathscr{F}_s^G$ for all $G \subseteq \mathscr{S} \cup \mathscr{R}$, $s \geq 0$.

**Proposition S1.2.** *If $\theta$ is a noiseless source encoding, i.e., it is defined as in Sec. VII, then*

$$\mathbb{P}(F \mid \mathscr{F}_0^{\{\mathsf{M}\} \cup J}) = \mathbb{P}(F \mid \mathscr{F}_s^{\{\theta\} \cup J})$$

*for all $s \in [0,t]$, $F \in \mathscr{F}_s^{\mathscr{R}}$ and $J \subseteq \mathscr{S}$.*

In other words, for all times $s \geq 0$ the conditional distribution of $R_{[0,s]}$, as a short notation for all reaction counters in $\mathscr{R}$, given any subset of species initial values and $M$ equals the conditional distribution given the same subset of initial values and the history of the protocol up to time $s^-$. The equality of intensity processes can then be obtained by specifying the initial value set to $\mathscr{S}_X$ (or $\mathscr{S}_Y$) and the all-reactions process to only the effective reaction processes $R^X$ accounting for changes of $X$ (or $R^Y$ accounting for changes of $Y$).

*Proof.* Let $s \in [0,t]$. By definition, the channel encoding propensities (119) (and actually the propensities of all reaction channels $r \in \mathscr{R}_\circ^X$) depend on $m$ only via $\Gamma_m^{\mathsf{X}}(s)$ for all $s \in [0,t]$. Since all other reaction channels have no $m$-dependence, the propensity functions for all $r \in \mathscr{R}$ depend on $m$ only via $\Gamma_m^{\mathsf{X}}(s)$ for all $s \in [0,t]$.

*Step 1: Introduction of two probability kernels.* Let $\mathbb{Q}_s, \tilde{\mathbb{Q}}_s \colon \mathscr{F}_s^{\mathscr{R}} \times \Omega \to [0,1]$ be regular probability kernels on $(\Omega, \mathscr{F}_s^{\mathscr{R}})$ such that almost surely

$$(F,\omega) \mapsto \mathbb{Q}_s(F,\omega) := \mathbb{P}\left(F \,\middle|\, \mathscr{F}_0^{\{\mathsf{M}\}\cup\mathscr{S}}\right)(\omega),$$
$$(F,\omega) \mapsto \tilde{\mathbb{Q}}_s(F,\omega) := \mathbb{P}\left(F \,\middle|\, \mathscr{F}_s^{\{\theta\}\cup\mathscr{S}}\right)(\omega).$$

Note that these probability kernels are conditioned on the species initial value set $\mathscr{S}$.

*Step 2: Define a common conditional intensity process on $\mathscr{F}$.* For every $r \in \mathscr{R}$ let $\psi_\tau^r \colon \mathscr{U} \times \mathscr{U} \times \theta([0,t]) \to \mathbb{R}_{\geq 0}$, where $\theta([0,t]) \subseteq \mathbb{R}_{\geq 0}$ denotes the state space (not the trajectory space) of $\theta$ on $[0,t]$, such that

$$\psi_\tau^r(u,u',\vartheta) := \begin{cases} \lambda_r(u,\vartheta) & \text{for } \tau > 0 \\ \lambda_r(u',\vartheta) & \text{for } \tau = 0 \end{cases}$$

Let $\omega' \in \Omega$. Then

$$(\omega,\tau) \mapsto \psi_\tau^r(U(\tau^-,\omega), U(0,\omega'), \theta(\tau^-,\omega')) \tag{S1.17}$$

is an $\mathscr{F}^{\mathscr{R}}$-predictable process, $\omega'$-$\mathbb{P}$-a.s. Predictability follows since is is an $\mathscr{F}^{\mathscr{R}}$-adapted process, i.e., given $\omega'$, the value of $U(\tau^-,\omega)$ is completely determined by $U(0,\omega')$, $\theta(\tau^-,\omega')$ and $\boldsymbol{R}_{[0,\tau)}(\omega)$ for all $\tau \in [0,s)$, $\omega'$-$\mathbb{P}$-a.s., and it has left-continuous trajectories. For fixed $\omega \in \Omega$ and $\tau \in [0,s)$ the random variable

$$\omega' \mapsto \psi_\tau^r(U(\tau^-,\omega), U(0,\omega'), \theta(\tau^-,\omega'))$$

is $\mathscr{F}_{\tau-}^{\{\theta\}\cup\mathscr{S}}$-measurable. By $\mathscr{F}_{\tau-}^{\{\theta\}\cup\mathscr{S}} \subseteq \mathscr{F}_0^{\{\mathsf{M}\}\cup\mathscr{S}}$ it is also $\mathscr{F}_0^{\{\mathsf{M}\}\cup\mathscr{S}}$-measurable.

Since $\psi_\cdot^r(U(\cdot^-), U(0), \theta(\cdot^-))$ is a $\mathscr{F}^{\{\theta\}\cup\mathscr{S}\cup\mathscr{R}}$-intensity (and a $\mathscr{F}^{\{\mathsf{M}\}\cup\mathscr{S}\cup\mathscr{R}}$-intensity) of $R_r$, it holds by definition [5, p. 27] that for all $r \in \mathscr{R}$

$$\mathbb{E}\left[\int_0^\infty \phi_\tau \, \mathrm{d}R_r(\tau)\right] = \mathbb{E}\left[\int_0^\infty \phi_\tau \, \psi_\tau^r(U(\tau^-), U(0), \theta(\tau^-)) \, \mathrm{d}\tau\right] \tag{S1.18}$$

for all $\mathscr{F}^{\{\theta\}\cup\mathscr{S}\cup\mathscr{R}}$-predictable (and all $\mathscr{F}^{\{\mathsf{M}\}\cup\mathscr{S}\cup\mathscr{R}}$-predictable) processes $\phi$. This equation is equivalent to saying that each $R_r$ has the above intensity process on both probability spaces $(\Omega, \mathscr{F}^{\{\theta\}\cup\mathscr{S}\cup\mathscr{R}}, \mathbb{P})$ and $(\Omega, \mathscr{F}^{\{\mathsf{M}\}\cup\mathscr{S}\cup\mathscr{R}}, \mathbb{P})$.

*Step 3: Reduce the intensity process to $\mathscr{F}_{[0,s]}^{\mathscr{R}}$.* What we actually want are conditional intensity processes on $(\Omega, \mathscr{F}^{\mathscr{R}}, \mathbb{P})$, which does no longer measure explicitly the information about initial values. This requires two essential steps, which are both a bit technical: (i) a conditioning step, and (ii) a disintegration step.

*Step 3 (i): Conditioning.* What we would like to obtain is the equality

$$\begin{aligned} &\mathbb{E}\left[\int_0^s \phi'_\tau \, \mathrm{d}R_r(\tau) \,\middle|\, \mathscr{F}_s^{\{\theta\}\cup\mathscr{S}}\right] \\ =&\mathbb{E}\left[\int_0^s \phi'_\tau \, \psi_\tau^r(U(\tau^-), U(0), \theta(\tau^-)) \, \mathrm{d}\tau \,\middle|\, \mathscr{F}_s^{\{\theta\}\cup\mathscr{S}}\right] \end{aligned} \tag{S1.19}$$

for all $\mathscr{F}_{[0,s]}^{\mathscr{R}}$-predictable non-negative processes $\phi'$. This is almost a conditional intensity definition. The final reduction of the filtration does then follow via disintegration. Eq. (S1.18) can be written equivalently as

$$\begin{aligned} &\mathbb{E}\left[\mathbb{1}_G \int_0^\infty \mathbb{1}_{[0,s]} \phi'_\tau \, \mathrm{d}R_r(\tau)\right] \\ =&\mathbb{E}\left[\mathbb{1}_G \int_0^\infty \mathbb{1}_{[0,s]} \phi'_\tau \, \psi_\tau^r(U(\tau^-), U(0), \theta(\tau^-)) \, \mathrm{d}\tau\right] \end{aligned} \tag{S1.20}$$

for all $G \in \mathscr{F}_s^{\{\theta\}\cup\mathscr{S}}$. To get this equality we will show a slightly more general one.

Let $H$ be a bounded, non-negative $\mathscr{F}_\infty^{\{\theta\}\cup\mathscr{S}\cup\mathscr{R}}$-measurable random variable. We define its $\mathscr{F}^{\{\theta\}\cup\mathscr{S}\cup\mathscr{R}}$-predictable projection as $h_s := \mathbb{E}[H \mid \mathscr{F}_{s-}^{\{\theta\}\cup\mathscr{S}\cup\mathscr{R}}]$ for all $s \geq 0$ and we denote by $(\tau_n(r))_{n\in\mathbb{N}}$ the jump times of $R_r$. For brevity we omit the functional arguments of $\psi_s^r$ in the following. Let $\phi$ be an arbitrary non-negative $\mathscr{F}^{\{\theta\}\cup\mathscr{S}\cup\mathscr{R}}$-predictable process.Then

$$\begin{aligned} \mathbb{E}\left[H \int_0^\infty \phi_s \, \mathrm{d}R_r(s)\right] &= \sum_{n=1}^\infty \mathbb{E}\left[H \, \phi_{\tau_n(r)}\right] \\ &= \sum_{n=1}^\infty \mathbb{E}\left[\mathbb{E}\left[H \, \phi_{\tau_n(r)} \mid \mathscr{F}_{\tau_n(r)-}^{\{\theta\}\cup\mathscr{S}\cup\mathscr{R}}\right]\right] \\ &= \sum_{n=1}^\infty \mathbb{E}\left[\mathbb{E}\left[H \mid \mathscr{F}_{\tau_n(r)-}^{\{\theta\}\cup\mathscr{S}\cup\mathscr{R}}\right] \phi_{\tau_n(r)}\right] \\ &= \sum_{n=1}^\infty \mathbb{E}\left[h_{\tau_n(r)} \, \phi_{\tau_n(r)}\right] \\ &= \mathbb{E}\left[\int_0^\infty h_s \, \phi_s \, \mathrm{d}R_r(s)\right] \\ &= \mathbb{E}\left[\int_0^\infty h_s \, \phi_s \, \psi_s^r \, \mathrm{d}s\right] \\ &= \int_0^\infty \mathbb{E}\left[h_s \, \phi_s \, \psi_s^r\right] \mathrm{d}s \\ &= \int_0^\infty \mathbb{E}\left[\mathbb{E}\left[H \, \phi_s \, \psi_s^r \mid \mathscr{F}_{s-}^{\{\theta\}\cup\mathscr{S}\cup\mathscr{R}}\right]\right] \mathrm{d}s \\ &= \int_0^\infty \mathbb{E}\left[H \, \phi_s \, \psi_s^r\right] \mathrm{d}s \\ &= \mathbb{E}\left[H \int_0^\infty \phi_s \, \psi_s^r \, \mathrm{d}s\right]. \end{aligned}$$

Here, we used the sum-representation of the counting integral, Fubini/Tonelli [2, Thm. 1.27] to exchange the order of summation and expectation, the definition (S1.18) of intensities to switch from the counting process integral to an ordinary integral, standard properties of the conditional expectation together with the fact that $\phi_{\tau_n(r)}$ is $\mathscr{F}^{\{\theta\}\cup\mathscr{S}\cup\mathscr{R}}_{\tau_n(r)-}$-measurable since it is predictable, and lastly, that $s \mapsto h_s\,\phi_s$ is a non-negative $\mathscr{F}^{\{\theta\}\cup\mathscr{S}\cup\mathscr{R}}$-predictable process.

Now, substituting $H = \mathbb{1}_G$ and $\phi_\tau = \mathbb{1}_{[0,s]}\phi'_\tau$ for all $\tau \geq 0$ yields (S1.20).

*Step 3 (ii): Disintegration.* To finish the step 3 we need to obtain the disintegration of (S1.17) within the conditional intensity "definition" (S1.19) to turn the equality into a proper intensity definition.

Using Fubini/Tonelli, we rewrite (S1.19) as

$$\begin{aligned}&\mathbb{E}\left[\int_0^s \phi'_\tau \,\mathrm{d}R_r(\tau)\middle|\mathscr{F}_s^{\{\theta\}\cup\mathscr{S}}\right]\\ &=\int_0^s \mathbb{E}\left[\phi'_\tau\,\psi^r_\tau(U(\tau^-),U(0),\theta(\tau^-))\middle|\mathscr{F}_s^{\{\theta\}\cup\mathscr{S}}\right]\mathrm{d}\tau\end{aligned}$$

Since $\phi'$ is not required to be integrable, we approximate argument of the conditional expectation from below with

$$\begin{aligned}&\lim_{n\to\infty}\min\{\phi'_\tau\,\psi^r_\tau(U(\tau^-),U(0),\theta(\tau^-)),n\}\\ =&\phi'_\tau\,\psi^r_\tau(U(\tau^-),U(0),\theta(\tau^-))\end{aligned}$$

Then, by monotone convergence and the disintegration theorem

$$\begin{aligned}&\mathbb{E}\left[\phi'_\tau\,\psi^r_\tau(U(\tau^-),U(0),\theta(\tau^-))\;\middle|\;\mathscr{F}_s^{\{\theta\}\cup\mathscr{S}\cup\mathscr{R}}\right](\omega)\\ =&\mathbb{E}\left[\lim_{n\to\infty}\min\{\phi'_\tau\,\psi^r_\tau(U(\tau^-),U(0),\theta(\tau^-)),n\}\;\middle|\;\mathscr{F}_s^{\{\theta\}\cup\mathscr{S}\cup\mathscr{R}}\right](\omega)\\ =&\lim_{n\to\infty}\mathbb{E}\left[\min\{\phi'_\tau\,\psi^r_\tau(U(\tau^-),U(0),\theta(\tau^-)),n\}\;\middle|\;\mathscr{F}_s^{\{\theta\}\cup\mathscr{S}\cup\mathscr{R}}\right](\omega)\\ =&\lim_{n\to\infty}\mathbb{E}\left[\min\{\phi'_\tau\,\psi^r_\tau(U(\tau^-),U(0,\omega),\theta(\tau^-,\omega)),n\}\;\middle|\;\mathscr{F}_s^{\{\theta\}\cup\mathscr{S}\cup\mathscr{R}}\right](\omega)\\ =&\mathbb{E}\left[\lim_{n\to\infty}\min\{\phi'_\tau\,\psi^r_\tau(U(\tau^-),U(0,\omega),\theta(\tau^-,\omega)),n\}\;\middle|\;\mathscr{F}_s^{\{\theta\}\cup\mathscr{S}\cup\mathscr{R}}\right](\omega)\\ =&\mathbb{E}\left[\phi'_\tau\,\psi^r_\tau(U(\tau^-),U(0,\omega),\theta(\tau^-,\omega))\;\middle|\;\mathscr{F}_s^{\{\theta\}\cup\mathscr{S}\cup\mathscr{R}}\right](\omega)\end{aligned}$$

In conclusion, we have

$$\begin{aligned}&\mathbb{E}\left[\int_0^s \phi'_\tau\,\mathrm{d}R_r(\tau)\;\middle|\;\mathscr{F}_s^{\{\theta\}\cup\mathscr{S}\cup\mathscr{R}}\right](\omega)\\ =&\mathbb{E}\left[\int_0^s \phi'_\tau\,\psi^r_\tau(U(\tau^-),U(0,\omega),\theta(\tau^-,\omega))\,\mathrm{d}\tau\;\middle|\;\mathscr{F}_s^{\{\theta\}\cup\mathscr{S}\cup\mathscr{R}}\right](\omega),\end{aligned}$$

which proves that $\psi^r_\tau(U(\tau^-),U(0,\omega),\theta(\tau^-,\omega))$, $s\in[0,t]$, is the $(\mathbb{Q}_s(\cdot,\omega),\mathscr{F}_s^{\mathscr{R}})$-intensity of $R_r$. The proof to establish that these processes are also $(\tilde{\mathbb{Q}}_s,\mathscr{F}_s^{\mathscr{R}})$-intensities of $R_r$ is identical and directly follows by substitution of $\mathscr{F}_s^{\{\theta\}\cup\mathscr{S}\cup\mathscr{R}}$ with $\mathscr{F}_0^{\{\mathsf{M}\}\cup\mathscr{S}\cup\mathscr{R}}$.

*Step 4: Equality of the probability kernels.* Since the two kernels have the same intensity process, the uniqueness theorem [5, p. 64] asserts $\mathbb{Q}(\cdot,\omega)=\tilde{\mathbb{Q}}(\cdot,\omega)$, $\mathbb{P}$-a.s.

*Step 5: Conditioning on subsets.* The generalization to arbitrary $J\subseteq\mathscr{S}$ follows by marginalization, i.e., integration of the kernels with

$$\mathbb{P}(G\,|\,\mathscr{F}_0^J)=\mathbb{P}(G\,|\,\mathscr{F}_0^{\{\mathsf{M}\}\cup J})=\mathbb{P}(G\,|\,\mathscr{F}_s^{\{\theta\}\cup J}),\quad G\in\mathscr{F}^{J^C},$$

where the equality $\mathbb{P}$-a.s follows by the independence of $M$ (and $\theta$) from $U(0)$. □

The following lemma provides us with an abstraction of the marginalization to aggregated reaction counters. Instead of looking at the aggregated processes, we may also just look at the original counters, but with respect to a smaller $\sigma$-algebra on the mark space. This reformulation facilitates the application of traditional results from filtering theory for multivariate point processes [5].

For all $G\subseteq\mathscr{R}$ denote $R_G(s):=\sum_{r\in G}R_r(s)$. Let $F\subseteq\mathscr{R}$ and $\Sigma_F$ be an arbitrary $\sigma$-algebra on $F$. For example, if $\Sigma_F=\{\emptyset,F\}$, then $\sigma(R_G(s)\colon s\in[0,t],G\in\Sigma_F)=\sigma(\sum_{r\in F}R_r(s)\colon s\in[0,t])$. Hence, we can group different reactions by choosing $\Sigma_F$ smaller than the power set.

**Lemma S1.4.** *Let $F\subseteq\mathscr{R}$, $N\leq|F|$ and $F_1,\ldots,F_N\subseteq F$ be a partition of $F$. If $\Sigma_F=\sigma(\{F_1,\ldots,F_N\})$, then for all $t\geq0$*

$$\sigma(R_G(s)\colon s\in[0,t],G\in\Sigma_F)=\sigma(\sum_{r\in F_k}R_r(s)\colon s\in[0,t],1\leq k\leq N).$$

*Proof.* Let $k\in\{1,\ldots,N\}$ and define the process $\{S_k(t)\}_{t\geq0}$

such that $S_k(t) := \sum_{r\in F_k} R_r(t) = R_{F_k}(t)$ for all $t \geq 0$. Let

$$\begin{aligned}\mathfrak{A} &:= \sigma(R_G(s)\colon s\in[0,t], G\in\Sigma_F),\\ \mathfrak{B} &:= \sigma(S_k(s)\colon s\in[0,t], 1\leq k\leq N).\end{aligned}$$

We prove $\mathfrak{A} = \mathfrak{B}$ by double inclusion.

$\mathfrak{B} \subseteq \mathfrak{A}$ is follows directly from $B_k \in \Sigma_F$ for all $k$, such that all $S_k$ are among the generating processes of $\mathfrak{A}$.

For the reverse inclusion note that for a $\sigma$-algebra, generated by a finite partition, every $G \in \Sigma_F$ can be represented as a (disjoint) union $G = \cup_{k\in I(G)} F_k$ for some $I(G) \subseteq \{1,\ldots,N\}$. Therefore, for all $s \in [0,t]$,

$$R_G(s) = \sum_{r\in G} R_r(s) = \sum_{k\in I(G)} \sum_{r\in F_k} R_r(s) = \sum_{k\in I(G)} S_k(s).$$

Hence $R_G(s)$ is a finite sum of $S_k(s)$ and thus $\mathfrak{B}$-measurable for all $G \in \Sigma_F$ and all $s \in [0,t]$, which implies $\mathfrak{A} \subseteq \mathfrak{B}$. □

For the formal statement of this equivalence of intensity processes we need the notion of the $\mathscr{F}$-predictable $\sigma$-algebra on $(0,\infty)\times\Omega$ [5, p. 8]:

$$\mathscr{P}(\mathscr{F}) := \sigma\left((s,\infty)\times A\colon A\in\mathscr{F}_s, s\geq 0\right)$$

This sigma-algebra is generated by all left-continuous processes. In turn, $((0,\infty)\times\Omega, \mathscr{P}(\mathscr{F}), \mu\otimes\mathbb{P})$ is a measure space, where $\mu$ denotes the Lebesgue measure. For our purpose we only look at the restricted space $(0,t]\times\Omega$ and denote restricted filtrations as $\mathscr{F}_{[0,t]}$.

The following proposition is a mathematically rigorous version of Proposition 2 in the main document. In contrast to Prop. S1.2, the following proposition is written under the reversible reaction notation of the main document, where $\mathscr{R}$ denotes reversible reaction channels with a forward and a backward direction. Its proof will, however, again follow the same conventions as in Prop. S1.2.

**Proposition S1.3.** *If $\theta$ is a noiseless source encoding, i.e., it is defined as specified in Sec. VII, then the channel encoding intensities are equivalent to the source-channel encoding intensities. That is, for all $\varepsilon \in \{+,-\}$ and $\nu_X \in \mathscr{V}_X$ it holds that*

$$\lambda^X(\varepsilon\nu_X) = \lambda^{M\to X}(\varepsilon\nu_X) \quad (\mu\otimes\mathbb{P})\text{-a.e. on } \mathscr{P}(\mathscr{F}^{XY}_{[0,t]}\vee\sigma(M)).$$

*Further, the source-output intensities satisfy*

$$\lambda^{M\to Y}_s(\varepsilon\nu_Y) = \mathbb{E}\left[\lambda^Y(\varepsilon\nu_Y, U(s^-))\middle|\mathscr{F}^{\theta Y}_{s-}\right],$$

*$(\mu\otimes\mathbb{P})$-a.e. on $\mathscr{P}(\mathscr{F}^Y_{[0,t]}\vee\sigma(M))$, for all $\varepsilon\in\{+,-\}$ and $\nu_Y \in \mathscr{V}_Y$.*

*Proof of Proposition S1.3.* Note that the equality of conditional probabilities in Proposition S1.2 holds for each $F \in \mathscr{H}_s := \sigma(R^X_\tau(\nu_X), R^Y_\tau(\nu_Y), \tau\in[0,s], \nu_X\in\mathscr{V}_X, \nu_Y\in\mathscr{V}_Y) \subseteq \mathscr{F}^{\mathscr{R}}_s$ (cf. Lemma S1.4) and $J = \mathscr{S}_X \cup \mathscr{S}_Y$. By the uniqueness theorem of predictable intensities [5, p. 31] the equality of probability kernels implies $\omega'$-$\mathbb{P}$-a.s.

$$\tilde{\lambda}^{M\to X}(\nu_X,\omega') = \tilde{\lambda}^X(\nu_X,\omega'), \quad (\mu\otimes\mathbb{P})\text{-a.e. on } \mathscr{P}(\mathscr{H}_t).$$

The tilde-notation indicates the disintegration of the original intensity processes $\lambda^{M\to X}(\nu_X)$, $\lambda^X(\nu_X)$ with respect to (i) the $\mathscr{H}_{s-}$-measurable degrees of freedom "$\omega$" (not explicitly visible in the given notation for brevity), and (ii) the disintegrated degrees of freedom "$\omega'$", which are $\mathscr{F}^{\{\theta\}\cup J}_s$-measurable . Such a disintegrated version exists by the factorization lemma [1]; for comparison see the disintegration in the case of Eq. (S1.17). What is left to do to obtain the desired equality is to reverse the disintegration.

Let $\phi\colon \Omega\times[0,t]\to\mathbb{R}_{\geq 0}$ be an arbitrary non-negative $\mathscr{F}^{XY}\vee\mathscr{F}^{\{\theta\}}$-predictable process. Consider the predictable projection $\tilde{\phi}_s = \mathbb{E}[\phi_s \mid \mathscr{H}_{s-}]$. Then for all $\nu_X \in \mathscr{V}_X$ it holds

$$\begin{aligned}\mathbb{E}\left[\int_0^t \phi_s\,\mathrm{d}R^X_s(\nu_X)\right] &= \sum_{n=1}^\infty \mathbb{E}\left[\phi_{\tau^X_n(\nu_X)-}\right]\\ &= \sum_{n=1}^\infty \mathbb{E}\left[\tilde{\phi}_{\tau^X_n(\nu_X)-}\right] = \mathbb{E}\left[\int_0^t \tilde{\phi}_s\,\mathrm{d}R^X_s(\nu_X)\right]\\ &= \mathbb{E}\left[\mathbb{E}\left[\int_0^t \tilde{\phi}_s\,\mathrm{d}R^X_s(\nu_X)\mid\mathscr{H}_t\right]\right]\\ &= \int \mathbb{E}\left[\int_0^t \tilde{\phi}_s\tilde{\lambda}^X_s(\nu_X,\omega')\,\mathrm{d}s\mid\mathscr{H}_t\right](\omega')\,\mathbb{P}(\mathrm{d}\omega)\\ &= \int_0^t\int \mathbb{E}\left[\phi_s\tilde{\lambda}^X_s(\nu_X,\omega')\mid\mathscr{H}_{s-}\right](\omega')\,\mathbb{P}(\mathrm{d}\omega')\,\mathrm{d}s\\ &= \int_0^t\int \mathbb{E}\left[\phi_s\lambda^X_s(\nu_X)\mid\mathscr{H}_{s-}\right](\omega')\,\mathbb{P}(\mathrm{d}\omega')\,\mathrm{d}s\\ &= \int_0^t \mathbb{E}\left[\phi_s\lambda^X_s(\nu_X)\right]\,\mathrm{d}s = \mathbb{E}\left[\int_0^t \phi_s\lambda^X_s(\nu_X)\,\mathrm{d}s\right]\end{aligned}$$

Here, we used the conditional intensity definition in line 4, Fubini/Tonelli [2, Thm. 1.27], the tower property and $\mathscr{H}_{s-}$-measurability of $\tilde{\lambda}^X(\nu_X,\omega')$ in line 5, the disintegration theorem [2, Thm. 6.4] in line 6 and again Fubini/Tonelli in the last equality. The equality follows analogously with $\lambda^{M\to X}(\nu_X)$.

Applying the same steps with $\mathscr{H}_s := \sigma(R^Y_\tau(\nu_Y), \tau\in[0,s], \nu_Y\in\mathscr{V}_Y) \subseteq \mathscr{F}^{\mathscr{R}}_s$ and $J := \mathscr{S}_Y$ proves the equivalence of channel output intensities and source-output intensities. □

#### B. Fano-type converse theorem

*Proof of Lemma 2.* Assume that $M \overset{d}{=} \theta_{[0,t]}$. Then

$$\begin{aligned}R &= \frac{1}{t}H(M) = \frac{1}{t}\left[\mathbb{I}(\theta_{[0,t]};D(Y_{[0,t]})) + H(M\mid D(Y_{(0,t]}))\right]\\ &\leq \frac{1}{t}\left[\mathbb{I}(\theta_{[0,t]};D(Y_{(0,t]})) + H_{\mathrm{B}}\left(\mathbb{P}(M\neq D(Y_{(0,t]}))\right)\right.\\ &\qquad \left.+\mathbb{P}(M\neq D(Y_{[0,t]}))\log(|\mathscr{M}|)\right]\\ &= \frac{1}{t}\mathbb{I}(\theta_{(0,t]};D(Y_{(0,t]})) + \frac{1}{t}H_{\mathrm{B}}\left(\mathbb{P}(M\neq D(Y_{(0,t]}))\right)\\ &\qquad +\mathbb{P}(M\neq D(Y_{(0,t]}))R_0,\end{aligned}$$

where the inequality follows from the following version of Fano's inequality [13]

$$\begin{aligned}H(M\mid D(Y_{(0,t]})) \leq& H_{\mathrm{B}}\left(\mathbb{P}(M\neq D(Y_{(0,t]}))\right)\\ &+\mathbb{P}(M\neq D(Y_{(0,t]}))\log(|\mathscr{M}|).\end{aligned}$$

By the data processing inequality [13] it holds that

$$\mathbb{I}(\theta_{[0,t]};D(Y_{(0,t]})) \leq \mathbb{I}(\theta_{[0,t]};Y_{(0,t]}),$$

which finishes the proof. □

*Proof of Thm. 3.* Let $\varepsilon \in (0,\frac{1}{2})$. By assumption $R$ is an achievable rate and hence also an $\varepsilon$-achievable rate. So, for all $\delta > 0$ exists a $t_0(\delta) > 0$ such that for all $t \geq t_0$ exists a message $M_t\colon \Omega \to \mathscr{M}_t$ and an $(|\mathscr{M}_t|,,t,\varepsilon)$-code for $M_t$, and it holds that $R-\delta \leq \inf_{t\in[t_0,\infty)} \frac{1}{t}H(M_t)$. Lemma 2 now asserts

$$\begin{aligned}
R-\delta &\leq \inf_{t\in[t_0,\infty)} \frac{1}{t}H(M_t)\\
&\leq \inf_{t\in[t_0,\infty)} \left\{\frac{1}{t}\mathbb{I}(\theta_{[0,t]};Y_{(0,t]}) + \frac{1}{t}H_{\mathrm{B}}(\varepsilon) + \varepsilon R_0\right\}\\
&= \varepsilon R_0 + \inf_{t\in[t_0,\infty)} \frac{1}{t}\mathbb{I}(\theta_{[0,t]};Y_{(0,t]})\\
&\leq \varepsilon R_0 + \inf_{t\in[t_0,\infty)} c_{\mathbb{I}}(t,Q_0)
\end{aligned}$$

Since $\inf_{t\in[t_0,\infty)} c_{\mathbb{I}}(t,Q_0)$ is monotonously increasing in $t_0$ and $t_0(\delta)$ is non-decreasing as $\delta \searrow 0$, we obtain

$$R \leq \varepsilon R_0 + C_{\mathbb{I}}(Q_0)$$

for all $\varepsilon \in (0,\frac{1}{2})$, i.e., $R \leq C_{\mathbb{I}}(Q_0)$ for all achievable rates. □

#### C. Energy-per-bit

*Proof of Lemma 3.* Let $R > 0$. Define for $t > 0$

$$\begin{aligned}
e_t &:= e_{\mathrm{b}}^{\min}(t,R)\\
E_* &:= E_{\mathrm{b}}^{\min}(R)
\end{aligned}$$

We start showing $\limsup_{t\to\infty} e_{\mathrm{b}}^{\min}(t,R) \leq E_{\mathrm{b}}^{\min}(R)$. Let $\delta > 0$. By (141) we then have

$$\begin{aligned}
&\liminf_{t\to\infty} c_{\mathbb{I}}(t,(E_*+\delta)R)\\
=&C_{\mathbb{I}}((E_*+\delta)R) \geq R > R-\delta
\end{aligned}$$

Hence, exists $t_0 > 0$ such that for all $t \geq t_0$

$$c_{\mathbb{I}}(t,(E_*+\delta)R) > R-\delta.$$

By (142) this implies

$$e_{\mathrm{b}}^{\min}(t,R-\delta) \leq \frac{R}{R-\delta}(E_*+\delta),$$

which through $\delta \searrow 0$ and the left-continuity of $e_{\mathrm{b}}^{\min}(t,R)$ in $R$ establishes

$$e_{\mathrm{b}}^{\min}(t,R) \leq E_*.$$

for all $t \geq t_0$. Consequently,

$$\limsup_{t\to\infty} e_{\mathrm{b}}^{\min}(t,R) \leq E_*.$$

So we turn to proving $E_{\mathrm{b}}^{\min}(R) \leq \limsup_{t\to\infty} e_{\mathrm{b}}^{\min}(t,R)$. Define $e_* := \limsup_{t\to\infty} e_t$. Then, for $\delta > 0$ exists $t_0 > 0$ such that such $e_t \leq e_* + \delta$ for all $t \geq t_0$. With

$$c_{\mathbb{I}}(t,(e_t+\delta)R) \geq R.$$

this implies

$$\begin{aligned}
C_{\mathbb{I}}((e_*+2\delta)R) &= \liminf_{t\to\infty} c_{\mathbb{I}}(t,(e_*+2\delta)R)\\
&\geq \liminf_{t\to\infty} c_{\mathbb{I}}(t,(e_t+\delta)R) > R-2\delta
\end{aligned}$$

and then

$$E_{\mathrm{b}}^{\min}(R-2\delta) \leq \frac{R}{R-2\delta}(e_*+2\delta)$$

which converges to the desired result as $\delta \searrow 0$ by the left-continuity of $E_{\mathrm{b}}^{\min}$. □

*Proof of Proposition 4.* Let $t,R > 0$. We start showing the inequality

$$e_{\mathrm{b}}^{\min}(t,R) \leq \frac{1}{R} q_{\mathbb{I}}^{\min}(t,R) \tag{S1.21}$$

Let $Q_* = q_{\mathbb{I}}^{\min}(t,R)$. By (143) exists a sequence of protocol-input pairs $(\theta_{[0,t]}^{(n)},X_{[0,t]}^{(n)})$ such that $\frac{1}{t}\mathbb{E}\left[q^{(n)}(t)\right]$ is a monotonously decreasing sequence with $\lim_{n\to\infty} \frac{1}{t}\mathbb{E}\left[q^{(n)}(t)\right] = Q_*$. For all $n \in \mathbb{N}$ it holds that $\mathbb{I}(\theta_{[0,t]}^{(n)};Y_{[0,t]}^{(n)}) \geq Rt$. Set $\varepsilon > 0$ and choose $n$ large enough such that $\frac{1}{t}\mathbb{E}\left[q^{(n)}(t)\right] \leq Q_* + \varepsilon$. Then, by (139) it holds that

$$c_{\mathbb{I}}(t,Q_*+\varepsilon) \geq \frac{1}{t}\mathbb{I}(\theta_{[0,t]}^{(n)};Y_{[0,t]}^{(n)}) \geq R$$

for $n$ large enough. Thus

$$e_{\mathrm{b}}^{\min}(t,R) \leq \frac{Q_*+\varepsilon}{R}$$

and (S1.21) follows by taking the limit $\varepsilon \searrow 0$.

Turning to the proof of the reverse inequality

$$\frac{1}{R} q_{\mathbb{I}}^{\min}(t,R) \leq e_{\mathrm{b}}^{\min}(t,R), \tag{S1.22}$$

let $\varepsilon > 0$ and set $E_* := e_{\mathrm{b}}^{\min}(t,R)$. By (142) it holds that $c_{\mathbb{I}}(t,(E_*+\varepsilon)R) \geq R > R-\varepsilon$. So there exist protocol-input pairs $(\theta_{[0,t]},X_{[0,t]})$ that satisfy

$$\begin{aligned}
\frac{1}{t}\mathbb{I}\left(\theta_{[0,t]},X_{[0,t]}\right) &\geq R-\varepsilon\\
\frac{1}{t}\mathbb{E}\left[q(t)\right] &\leq (E_*+\varepsilon)R
\end{aligned}$$

So taking the infimum as in (143) yields

$$q_{\mathbb{I}}^{\min}(t,R-\varepsilon) \leq (E_*+\varepsilon)R.$$

As $q_{\mathbb{I}}^{\min}(t,R)$ is left-continuous in $R$, taking the limit $\varepsilon \searrow 0$ on both sides yields (S1.22).

The equality for the asymptotic quantities follows directly by Lemma 3. □

[1] A. Klenke, *Probability Theory : A Comprehensive Course*, 3rd ed. (Springer International Publishing, Cham, 2020).
[2] O. Kallenberg, *Foundations of modern probability* (Springer, 1997).
[3] R. Boel, P. Varaiya, and E. Wong, Martingales on jump processes. II: Applications, SIAM Journal on Control **13**, 1022 (1975).
[4] M. Gehri, N. Engelmann, and H. Koeppl, Mutual information of a class of Poisson-type channels using Markov renewal theory, in *2024 IEEE International Symposium on Information Theory (ISIT)* (2024) pp. 1931–1936.
[5] P. M. Brémaud, *Point Processes and Queues: Martingale Dynamics*, Vol. 50 (Springer, 1981).
[6] R. S. Liptser and A. N. Shiryaev, *Statistics of random processes: I. General theory*, Vol. 5 (Springer Science & Business Media, 2013).
[7] D. J. Daley and D. Vere-Jones, *An Introduction to the Theory of Point Processes: Volume I: Elementary Theory and Methods* (Springer Science & Business Media, 2006).
[8] A.-L. Moor and C. Zechner, Dynamic information transfer in stochastic biochemical networks, Physical Review Research **5**, 013032 (2023).
[9] L. Bronstein and H. Koeppl, Marginal process framework: A model reduction tool for Markov jump processes, Physical Review E **97**, 062147 (2018).
[10] T. Weissman, Y.-H. Kim, and H. H. Permuter, Directed information, causal estimation, and communication in continuous time, IEEE Transactions on Information Theory **59**, 1271 (2012).
[11] N. J. Newton, Transfer entropy and directed information in Gaussian diffusion processes, arXiv preprint arXiv:1604.01969 (2016).
[12] T. M. Apostol, *Mathematical Analysis*, 2nd ed. (Addison-Wesley, 1974).
[13] J. A. T. Thomas M. Cover, Entropy, relative entropy, and mutual information, in *Elements of Information Theory* (John Wiley & Sons, Ltd, 2005) Chap. 2, pp. 13–55.
