## Supplementary Material S2 for "Biochemical communication with noisy feedback under energy constraints"

(Dated: July 7, 2026)

### CONTENTS

S2.1. Coarse-graining and derivation of mesoscopic energies 1

S2.2. Local detailed balance of open chemical reaction networks and the multi-reservoir picture 4

S2.3. Entropy Production Rate of driven conversion networks and general CTMCs with multiple reservoirs 4

S2.4. Macroscopic consistency of local detailed balance for open chemical reaction networks 5

S2.5. Adiabatic and non-adiabatic entropy production 6

S2.6. Stochastic consistency and statistical time reversal 7

S2.7. Proofs for stochastic thermodynamics 10

References 13

In this Supplementary Materials, we offer additional background on stochastic thermodynamics and include technical proofs that were too extensive for the Primer Section III. This includes, in particular, a review of a coarse-graining procedure that connects the microscopic dynamics of a chemical system with both the coarse-grained mesoscopic dynamics of an open chemical reaction network and the related macroscopic thermodynamic quantities, ensuring consistency across all levels.

Additionally, we review stochastic consistency, which relates thermodynamic reversibility to the distributional equality of a forward process with the time-reverse of the backward process, that is, statistical reversibility. The concept of stochastic consistency is important for trajectory-level second law-like relations, known as fluctuation theorems. In this context, the entropy production rate was shown to coincide with the KL-divergence between forward and backward trajectories under the assumption of stationarity.

### S2.1. COARSE-GRAINING AND DERIVATION OF MESOSCOPIC ENERGIES

In this section, we review the coarse-graining construction underlying the mesoscopic thermodynamic potentials used in the main text, following Peliti and Pigolotti [1, Ch. 3.12]. Starting from microscopic ensemble definitions, with potentially strong coupling between the core species U and the solvent [2, 3], we derive mesoscopic thermodynamic potentials for solvated systems. The construction is guided by macroscopic thermodynamic laws and relies on a separation of time scales between fast microscopic equilibration and slow mesoscopic state changes. Finally, we use this time-scale separation to motivate the trajectory-level calorimetric heat, including both the reaction-induced contribution associated with jumps of the core system U and, under external manipulation, the continuous contribution generated by protocol-induced re-equilibration of the microscopic states.

Let $\xi$ be a state of the stochastic process $\{\Xi(t)\}_{t\geq 0}$, representing the microscopic ensemble including chemical composition, and dimensionless particle positions and moments of both core system species U and solvent species S. The solvent species act as the environment of the core species, but the solution is itself typically embedded in (weakly coupled to) an external macroscopic environment. The energy landscape of the microstates in a closed system be dictated by the Hamiltonian $\xi \mapsto \mathfrak{H}(\xi)$, which includes both the kinetic energy and the potential energy for any chemical composition. Its potential energy part may include non-reactive particle interactions like Coulomb forces or van-der-Waals interactions. Changes in chemical composition, like covalent bond formation, cannot be accounted for in a classical Hamiltonian, which is why the energy landscape is given for every chemical composition.

Each microstate consist of the degrees of freedom of the core system and the solvent, i.e., $\xi = (\xi_{\mathsf{U}}, \xi_{\mathsf{S}})$. A dilute solution $\mathsf{U} \cup \mathsf{S}$ is called weakly coupled, if interaction energies between the core system and the solvent are negligible, such that the Hamiltonian approximately decomposes as

$$\mathfrak{H}(\xi) \approx \mathfrak{H}_{\mathsf{U}}(\xi_{\mathsf{U}}) + \mathfrak{H}_{\mathsf{S}}(\xi_{\mathsf{S}}).$$

In biological cells, biomolecules U are typically strongly coupled to the solvent, resulting in an interaction term:

$$\mathfrak{H}(\xi) = \mathfrak{H}_{\mathsf{Z}}(\xi_{\mathsf{Z}}) + \mathfrak{H}_{\mathsf{S}}(\xi_{\mathsf{S}}) + \mathfrak{H}_{\mathsf{Z}\leftrightarrow\mathsf{S}}(\xi_{\mathsf{Z}}, \xi_{\mathsf{S}})$$

Previous works have discussed consistent definitions of stochastic thermodynamic quantities for the marginal degrees of freedom $\xi_{\mathsf{U}}$ of the core system, both in classical [2, 3] and quantum [4] settings. Here we adopt a complementary viewpoint; we treat the combined solution $\mathsf{U} \cup \mathsf{S}$ as the thermodynamic system and assume only that this full solution is weakly coupled to an external macroscopic environment. In this way, strong coupling between solute and solvent within the solution is fully compatible with the microscopic starting point. The mesoscopic thermodynamic potentials derived below therefore pertain to the entire solution $\mathsf{U} \cup \mathsf{S}$, rather than to the solute $\mathsf{U}$ alone. Deriving effective potentials for the core system would require an additional marginalization over solvent degrees of freedom, as discussed by Jarzynski [3] and, for CRNs, by Schmiedl and Seifert [5, App. A]. We omit this further step here in order to keep the derivation focused on the mesoscopic structure used in the main text.

The macroscopic internal energy of the solvated system $\mathsf{U} \cup \mathsf{S}$ at time $t$ is the mean of the microscopic Hamiltonian:

$$E_{\mathrm{i}}[\Xi(t)] := \mathbb{E}[\mathfrak{H}(\Xi(t))].$$

The macroscopic Boltzmann entropy is given by $k_{\mathrm{B}} H[\Xi(t)]$, where $H[\cdot]$ denotes the Shannon entropy. The macroscopic nonequilibrium Helmholtz free energy is then defined as

$$F[\Xi(t)] := E_{\mathrm{i}}[\Xi(t)] - k_{\mathrm{B}} T\, H[\Xi(t)]. \tag{S2.1}$$

According to the laws of thermodynamics, a closed system of fixed volume in contact with a thermal reservoir at temperature $T$ will relax toward a state that minimizes its Helmholtz free energy. The zeroth law implies convergence in distribution to an equilibrium ensemble $\Xi_{\mathrm{eq}} = \lim_{t\to\infty} \Xi(t)$, for which

$$F^{\mathrm{eq}} = \min_{\Xi} F[\Xi] = F[\Xi^{\mathrm{eq}}]. \tag{S2.2}$$

Equilibrium statistical mechanics identifies this equilibrium distribution with the Gibbs–Boltzmann distribution of a canonical ensemble:

$$p(\Xi^{\mathrm{eq}} = \xi) = \frac{1}{P_\Xi} e^{-\beta \mathfrak{H}(\xi)} \tag{S2.3}$$

with $\beta = (k_{\mathrm{B}} T)^{-1}$ and the partition function

$$P_\Xi := \int e^{-\beta \mathfrak{H}(\xi)}\, \mathrm{d}\mu_\Xi(\xi).$$

The reference measure $\mu_\Xi$ is a hybrid product measure combining a Lebesgue measure over the continuous degrees of freedom (e.g., particle positions and momenta) with a counting measure over discrete degrees of freedom, such as chemical compositions. The continuous phase space is rendered dimensionless via normalization by Planck's constant $h$, i.e., by dividing each momentum-position phase space volume element by $h^3$. To account for particle indistinguishability, the discrete measure includes a combinatorial prefactor. For instance, for a composition $u \in \mathscr{U}$ of the core species, the discrete normalization of $\xi_{\mathsf{U}}$ involves the factor $1/u! = \prod_{n=1}^{N} 1/u_n!$. For simplicity, we keep all these normalizations implicit.

Inserting (S2.3) into (S2.2) yields

$$F^{\mathrm{eq}} = -k_{\mathrm{B}} T \ln P_\Xi, \tag{S2.4}$$

which relates the macroscopic free energy at equilibrium directly to the microscopic partition function.

We now introduce the coarse-graining procedure by identifying the states $\mathscr{U}$ of the closed CRN over the species $\mathsf{U}$ as the mesostates of the full microscopic ensemble process $\Xi$. The state space of $\Xi$ be partitioned into disjoint subsets $\mathscr{A}_u$, such that $\{U(t) = u\} = \{\Xi(t) \in \mathscr{A}_u\}$ for all $u \in \mathscr{U}$. We assume a separation of time scales whereby transitions between mesostates $u$ occur on a much slower time scale than the equilibration of the microstates within each $\mathscr{A}_u$. This assumption allows us to impose a special case of local equilibrium [6] condition: the conditional distribution of microstates, given the mesostate, at time $t$ coincides with the conditional equilibrium distribution,

$$\begin{aligned}\mathbb{P}(\Xi(t) = \xi \mid U(t) = u) &= \mathbb{P}(\Xi^{\mathrm{eq}} = \xi \mid U^{\mathrm{eq}} = u) \\ &= \mathbb{1}\{\xi \in \mathscr{A}_u\} \frac{e^{-\beta \mathfrak{H}(\xi)}}{P_u},\end{aligned} \tag{S2.5}$$

where $P_u := \int_{\mathscr{A}_u} e^{-\beta \mathfrak{H}(\xi)}\, \mathrm{d}\mu_\Xi(\xi)$ is the conditional partition function given the mesostate $u$. For simplicity, we neglected the reactions among the solvent species $\mathsf{S}$. While the coarse-graining scheme is not unique, the combination of time-scale separation and local equilibrium is a central modeling assumption in stochastic thermodynamics [1].

The marginal distribution over mesostates at equilibrium is

$$\begin{aligned}\pi(u) :&= \mathbb{P}(U^{\mathrm{eq}} = u) = \mathbb{P}(\Xi^{\mathrm{eq}} \in \mathscr{A}_u) \\ &= \frac{1}{P_\Xi} \int_{\mathscr{A}_u} e^{-\beta \mathfrak{H}(\xi)}\, \mathrm{d}\mu_\Xi(\xi) = \frac{P_u}{P_\Xi}.\end{aligned}$$

We now define the mesoscopic Helmholtz free energy as the function

$$f(u) := -k_{\mathrm{B}} T \ln P_u,$$

which allows us to express the equilibrium distribution of mesostates as the Gibbs-Boltzmann distribution

$$\mathbb{P}(U^{\mathrm{eq}} = u) = \frac{1}{P_\Xi} e^{-\beta f(u)}.$$

The function $f(u)$ has the same functional form as the macroscopic Helmholtz free energy in Eq. (S2.4), but evaluated over the conditional ensemble restricted to mesostate $u$. Indeed, by (S2.5) it satisfies the identity

$$f(u) = \mathbb{E}[\mathfrak{H}(\Xi^{\mathrm{eq}}) \mid U^{\mathrm{eq}} = u] - k_B T\, H[\Xi^{\mathrm{eq}} \mid U^{\mathrm{eq}} = u].$$

Consequently, we define the mesoscopic internal energy $e_{\mathrm{i}}$ and the internal (Shannon) entropy $h_{\mathrm{i}}$ of the state $u$ as

$$\begin{aligned}e_{\mathrm{i}}(u) &:= \mathbb{E}[\mathfrak{H}(\Xi^{\mathrm{eq}}) \mid U^{\mathrm{eq}} = u] \\ h_{\mathrm{i}}(u) &:= H[\Xi^{\mathrm{eq}} \mid U^{\mathrm{eq}} = u],\end{aligned}$$

which yields the mesoscopic identity

$$f(u) = e_{\mathrm{i}}(u) - k_B T h_{\mathrm{i}}(u).$$

Furthermore, the macroscopic nonequilibrium Helmholtz free energy from Eq. (S2.1) can be decomposed as

$$\begin{aligned}F[\Xi(t)] =& \mathbb{E}[\mathbb{E}[\mathfrak{H}(\Xi(t)) \mid U(t)]] - k_{\mathrm{B}} T H[\Xi(t) \mid U(t)] \\ &- k_{\mathrm{B}} T H[U(t)] \\ =& \sum_u (\mathbb{E}[\mathfrak{H}(\Xi(t)) \mid U(t) = u] \\ &\quad - k_{\mathrm{B}} T H[\Xi(t) \mid U(t) = u]) \mathbb{P}(U(t) = u) \\ &- k_{\mathrm{B}} T H[U(t)],\end{aligned}$$

At equilibrium, this expression reduces to

$$F^{\mathrm{eq}} = \sum_u f(u)\,\pi(u) - k_{\mathrm{B}} T\, H[U^{\mathrm{eq}}],$$

even without invoking the time-scale separation assumption. Under the local equilibrium assumption (S2.5), the nonequilibrium free energy admits the simplified representation

$$F[\Xi(t)] = \mathbb{E}\left[f(U(t))\right] - k_{\mathrm{B}} T\, H[U(t)],$$

demonstrating that the mesoscopic free energy function $f(u)$ plays a role on the level of mesostates analogous to that of the microscopic Hamiltonian $\mathfrak{H}$ on the level of microstates.

However, it is important to emphasize that $\mathbb{E}\left[f(U(t))\right] \neq E_{\mathrm{i}}[\Xi(t)]$ in general. This distinction must be kept in mind when interpreting the first and second laws of stochastic thermodynamics in the mesoscopic setting discussed in the main text in Sec. III B and III C.

Similar derivations of mesoscopic thermodynamic potentials, represented by mesoscopic free energy functions, can be carried out for any generalized canonical ensemble [7] of the solvated system. The formal structure of the laws of stochastic thermodynamics remains consistent across such ensembles, with the mesoscopic free energy selected in accordance with the constraints imposed by the ensemble.

In particular, biological cells typically operate under constant-pressure conditions. The flexibility and selective permeability of the cell membrane support the regulation of intracellular pressure through osmotic exchange with the environment. To incorporate solvent exchange through the semipermeable membrane, we consider the solvent as partitioned into molecules $\xi_{\mathsf{S}}$ within the cellular volume and molecules $\xi_{\mathsf{E}}$ in the external environment. While the extracellular compartment contains only solvent, the intracellular region includes both solvent and solute species $\mathsf{U}$.

The isothermal-isobaric ensemble inside the cell corresponds to a generalized canonical ensemble, where the total enthalpy of the solvated system is given by

$$\mathfrak{H}_{\mathrm{tot}}(\xi) = \mathfrak{H}(\xi_{\mathsf{U}}, \xi_{\mathsf{S}}) + \mathsf{p}\, v(\xi_{\mathsf{U}}, \xi_{\mathsf{S}}),$$

with $\mathsf{p}$ denoting the external pressure and $v(\xi)$ the volume occupied by the solvated system. Here, $\mathsf{p}$ and $v$ represent a conjugate pair of thermodynamic variables: a generalized force and its corresponding coordinate.

The embedding of the cell into a larger external solvent reservoir at the same pressure can be modeled by weakly coupling the system to the external solvent degrees of freedom $\xi_{\mathsf{E}}$. The total enthalpy of the combined system $\mathsf{U} \cup \mathsf{S} \cup \mathsf{E}$ is then

$$\mathfrak{H}_{\mathsf{U}\cup\mathsf{S}\cup\mathsf{E}}(\xi, \xi_{\mathsf{E}}) = \mathfrak{H}_{\mathrm{tot}}(\xi) + \mathfrak{H}_{\mathsf{E}}(\xi_{\mathsf{E}}) + \mathsf{p}\, v_{\mathsf{E}}(\xi_{\mathsf{E}})$$

where $v_{\mathsf{E}}$ denotes the volume occupied by the external compartment. Assuming such weak coupling, the equilibrium distribution of the total system is proportional to $e^{-\beta \mathfrak{H}_{\mathsf{U}\cup\mathsf{S}\cup\mathsf{E}}(\xi, \xi_{\mathsf{E}})}$, and the marginal equilibrium distribution of the intracellular solvated system takes the generalized canonical form

$$p(\Xi^{\mathrm{eq}} = \xi) = \frac{1}{\mathscr{P}_{\Xi}}\, e^{-\beta \mathfrak{H}_{\mathrm{tot}}(\xi)} = \frac{1}{\mathscr{P}_{\Xi}}\, e^{-\beta \mathfrak{H}(\xi) - \beta \mathsf{p} v(\xi)},$$

with partition function

$$\mathscr{P}_{\Xi} := \int e^{-\beta \mathfrak{H}(\xi) - \beta \mathsf{p} v(\xi)}\, \mathrm{d}\mu_{\Xi}(\xi).$$

The macroscopic nonequilibrium Gibbs free energy is defined as

$$\begin{aligned} G[\Xi(t)] :=& \mathbb{E}\left[\mathfrak{H}(\Xi(t))\right] + \mathsf{p}\, \mathbb{E}\left[v(\Xi(t))\right] \\ & - k_{\mathrm{B}} T\, H[\Xi(t)], \end{aligned} \tag{S2.6}$$

where the second term represents the mean mechanical work associated with the system volume. For a closed system maintained at constant temperature and pressure, the Gibbs free energy reaches its minimum at equilibrium. Hence, the equilibrium free energy satisfies

$$G^{\mathrm{eq}} = G[\Xi^{\mathrm{eq}}] = -k_{\mathrm{B}} T \ln(\mathscr{P}_{\Xi}),$$

where $\mathscr{P}_{\Xi}$ denotes the isothermal–isobaric partition function.

To avoid redundancy, we do not repeat the full coarse-graining derivation here. The steps proceed analogously to the isothermal case, with $\mathfrak{H}$ replaced by the total enthalpic Hamiltonian $\mathfrak{H}_{\mathrm{tot}}$. The mesoscopic Gibbs free energy associated with mesostate $u$ is then given by

$$g(u) = \mathbb{E}\left[\mathfrak{H}_{\mathrm{tot}}(\Xi^{\mathrm{eq}}) \mid U_{\mathrm{eq}} = u\right] - k_{\mathrm{B}} T\, H\left[\Xi^{\mathrm{eq}} \mid U_{\mathrm{eq}} = u\right],$$

and under the local equilibrium assumption, the nonequilibrium Gibbs free energy becomes

$$G[\Xi(t)] = \mathbb{E}\left[g(U(t))\right] - k_{\mathrm{B}} T\, H[U(t)].$$

Defining the mesoscopic enthalpy $\mathfrak{h}$ and solution volume $\mathfrak{v}$ as

$$\begin{aligned} \mathfrak{h}(u) :=& \mathbb{E}\left[\mathfrak{H}_{\mathrm{tot}}(\Xi^{\mathrm{eq}}) \mid U_{\mathrm{eq}} = u\right] = e_{\mathrm{i}}(u) + \mathsf{p}\mathfrak{v}(u), \\ \mathfrak{v}(u) :=& \mathbb{E}\left[v(\Xi^{\mathrm{eq}}) \mid U_{\mathrm{eq}} = u\right] \end{aligned}$$

yields a mesocopic identity, analogous to the macroscopic (S2.6):

$$g(u) = \mathfrak{h}(u) - k_{\mathrm{B}} T\, h_{\mathrm{i}}(u).$$

It is worth noting that fluctuations in the mesoscopic volume function $\mathfrak{v}(u)$ across different mesostates $u$ are typically considered negligible and are therefore omitted in most kinetic models of biochemical reaction networks, as discussed in Sec. II.

In an analogous manner, we now motivate the calorimetric heat exchanged between the solution and its environment. Under the local equilibrium assumption, the process $U(t)$ can be treated as a mesoscopic parameter of a macroscopic state transformation, that is, of a change in macroscopic energies and entropies through heat exchange and work. If the system is externally manipulated, the microscopic energy landscape itself may vary in time through a determined protocol, so that both the microscopic Hamiltonian and the volume function become time dependent, $\mathfrak{H}(\xi, t)$ and $v(\xi, t)$. In the isothermal–isobaric setting, this amounts to a time-dependent total enthalpic Hamiltonian

$$\mathfrak{H}_{\mathrm{tot}}(\xi, t) := \mathfrak{H}(\xi, t) + \mathsf{p}\, v(\xi, t),$$

and hence to a time-dependent mesoscopic enthalpy

$$\mathfrak{h}(u,t) := \mathbb{E}[\mathfrak{H}_{\text{tot}}(\Xi^{\text{eq}},t) \mid U^{\text{eq}} = u].$$

The local equilibrium assumption for this mesoscopic potential now also includes that the re-equilibration of microscopic states is fast compared to the change of the Hamiltonian. The differential first law of macroscopic thermodynamics reads [6]

$$\mathrm{d}E_{\mathrm{i}} = \delta W - \delta Q,$$

where $\delta W$ denotes the differential work performed on the system and $\delta Q$ the heat released to the environment, with the sign convention of stochastic thermodynamics. At a reaction jump time, however, the protocol is instantaneously frozen, so that its explicit time dependence contributes only continuously between jumps. The jump contribution is therefore determined solely by the state change $U(t) \to U(t+)$. Since the solvated system $\mathsf{U} \cup \mathsf{S}$ is treated here as the thermodynamic system, the only jump-associated work contribution is the pressure-volume term; chemical work appears explicitly only later, in the open-system description of the main text, where the chemostats are moved outside the system boundary. Accordingly, the mesoscopic first-law balance at a jump reads

$$\mathrm{d}e_{\mathrm{i}}(U(t)) = \mathrm{d}w^{\mathrm{r}}(t) - \mathrm{d}q^{\mathrm{r}}(t),$$

where $\mathrm{d}q^{\mathrm{r}}(t)$ denotes the reaction-induced increment of the calorimetric heat and $\mathrm{d}w^{\mathrm{r}}(t)$ the work increment associated with the jump. Hence, for a closed isothermal-isobaric system, the jump-associated work contribution is the pressure-volume term,

$$\begin{aligned}\mathrm{d}w^{\mathrm{r}}(t) &= -\,\mathsf{p}\,(\mathfrak{v}(U(t+),t) - \mathfrak{v}(U(t),t)),\\ \mathrm{d}q^{\mathrm{r}}(t) &= \mathfrak{h}(U(t),t) - \mathfrak{h}(U(t+),t),\end{aligned}$$

in agreement with the macroscopic enthalpy balance. The full trajectory-level calorimetric heat is then obtained by supplementing this jump contribution with the continuous manipulation-induced part generated by the explicit time dependence of the protocol.

### S2.2. LOCAL DETAILED BALANCE OF OPEN CHEMICAL REACTION NETWORKS AND THE MULTI-RESERVOIR PICTURE

Transitions between states of $\mathscr{U}$ can be interpreted as interactions with multiple reservoirs $r$, each associated with a reaction channel $\mathfrak{R}_r$. These reservoirs are characterized by macroscopic equilibrium chemical potentials $\mu_l^{(r)}$ and temperatures $T_r$. If the chemostat species $\mathsf{Z}$ were explicitly modeled via their mesoscopic copy numbers $z' = z\mathsf{V}$ and their exchanges with the environment treated as reactions, then the mesoscopic open system state $(u,z')$ would admit a well-defined grand potential relative to reservoir $r$:

$$\begin{aligned}\Phi^{(r)}(u,z') :=& \mathfrak{h}(u,z') - k_{\mathrm{B}} T_r h_{\mathrm{i}}(u,z')\\ &- \sum_{l \in \{l' :\, r_{l'}^{(r)} \neq 0 \text{ or } s_{l'}^{(r)} \neq 0\}} \mu_l^{(r)} z_l',\end{aligned}$$

where only the species $\mathsf{Z}$ are coupled to the reservoirs [8, Eq. (1)]. Assuming all reservoirs share a common temperature $T$ and identical chemical potentials $\mu_l$, the grand potential for the full system, coupled to $|\mathscr{R}|$ reservoirs, becomes

$$\Phi(u,z') := \mathfrak{h}(u,z') - k_{\mathrm{B}} T\, h_{\mathrm{i}}(u,z') - \sum_{l=1}^{L} \mu_l z_l'.$$

The corresponding grand canonical equilibrium distributions satisfy

$$\begin{aligned}\pi_r(u,z') &\propto e^{-\beta \Phi^{(r)}(u,z')},\\ \pi(u,z') &\propto e^{-\beta \Phi(u,z')}.\end{aligned} \tag{S2.7}$$

Note that for these distributions to be kinetically consistent, all exchange reactions with the reservoirs must be included.

A system coupled to a single reservoir satisfies the detailed balance condition

$$\lambda_r^+(u,z')\,\pi_r(u,z') = \lambda_r^-(u+\nu_r,z')\,\pi_r(u+\nu_r,z'), \tag{S2.8}$$

which holds even when the reservoirs differ in temperature or chemical potentials. Note that, by construction, the reaction $\mathfrak{R}_r$ models only changes in the core species and not the explicit exchange of particles with the reservoir, although detailed balance would also apply to such exchange reactions if they were included in the set of reaction channels. In the special case where all reservoirs share the same temperature and chemical potentials, the condition (S2.8) also holds with the multi-reservoir equilibrium distribution $\pi(u,z')$.

Under the chemostat assumption, which can be understood as a coarse-graining where $\mathsf{Z}$ equilibrates much faster than $U$, both the single-reservoir and multi-reservoir potential changes satisfy

$$-\beta \Delta_r^+ \Phi(u,z') = -\beta \Delta_r^+ \Phi^{(r)}(u,z') = \beta\, \Delta_r^+ Q(u) + \Delta_r^+ h_{\mathrm{i}}(u),$$

recovering the right-hand side of (12), since enthalpy and entropy changes in $z'$ are immediately compensated. Reservoir exchanges of $\mathsf{Z}$ are not tracked explicitly in the marginal thermodynamic description of $U$.

Combining (S2.7) and (S2.8) (either with single-reservoir coupling or multi-reservoir coupling) and subsequently applying the chemostat assumption yields the local detailed balance relation (12). Importantly, under the chemostat assumption, detailed balance no longer holds for the marginal distribution $\pi(u)$, where the chemostat degrees of freedom have been integrated out.

In particular, this derivation shows that the central local detailed balance condition can be recovered by considering each $\mathfrak{R}_r$ as a separate reservoir coupling and neglecting the other reactions.

### S2.3. ENTROPY PRODUCTION RATE OF DRIVEN CONVERSION NETWORKS AND GENERAL CTMCS WITH MULTIPLE RESERVOIRS

Much of the literature on entropy production rate employs a formulation based on the infinitesimal generator (transition rate matrix) $\Lambda$ of a CTMC, rather than a reaction network perspective [1, 9, 10]. A bridge between these two pictures can be constructed by introducing an index set $\mathscr{J} := \{1,\ldots,|\mathscr{U}|\}$, where each mesoscopic state $u \in \mathscr{U}$ is uniquely assigned an index $j \in \mathscr{J}$ (with $|\mathscr{U}| = \infty$ allowed). This indexing defines a one-hot encoding of the state space, $\mathscr{U}' = \{e_j \colon j \in \mathscr{J}\}$, where $e_j$ denotes the unit vector with a single non-zero entry at position $j$.

Viewed as a population representation, this encoding interprets each state $e_j$ as indicating the presence of a single effective species $\mathsf{U}'_j$, with all other species absent. This reformulation recasts the original reaction network into a conversion network: a collection of microscopically reversible transitions between discrete one-hot states. Each conversion $i \to j$ (with change vector $v_{(i,j)} = e_j - e_i$) may be coupled to one or more reservoirs $r$, corresponding to the reaction channels $\mathfrak{R}_r$, as discussed in Sec. S2.2.

For each reservoir $r$, we denote the set of reversible transitions it mediates as

$$\mathscr{R}^{(r)} := \{(i,j) \in \mathscr{J}^2 \mid i<j,\ \lambda^{(+r)}_{(i,j)}(e_i) > 0\}.$$

with stoichiometric balance equations

$$\mathfrak{R}^{(r)}_{(i,j)}: \quad \mathsf{U}_i \underset{\lambda^{(-r)}_{(i,j)}}{\overset{\lambda^{(+r)}_{(i,j)}}{\rightleftarrows}} \mathsf{U}_j, \quad \text{with } (i,j) \in \mathscr{R}^{(r)}$$

and reservoir-specific propensity functions $\lambda^{(\pm r)}_{(i,j)}(u')$.

The effective Markov generator $\Lambda$ of such a network, coupled to multiple reservoirs $r$ is then given as the sum of reservoir-wise generators $\Lambda^{(r)}$, such that the transition rates for $i \to j$ (and also diagonal elements) satisfy [8]

$$\Lambda_{ij} = \sum_{r\in\mathscr{R}} \Lambda^{(r)}_{ij}.$$

For each $(i,j) \in \mathscr{R}^{(r)}$ the reservoir-wise propensity functions and generators are related as

$$\begin{aligned} \lambda^{(+r)}_{(i,j)}(u') &= \Lambda^{(r)}_{ij}\, \mathbb{1}\{u' = e_i\}, \\ \lambda^{(-r)}_{(i,j)}(u') &= \Lambda^{(r)}_{ji}\, \mathbb{1}\{u' = e_j\}. \end{aligned} \tag{S2.9}$$

The reservoir-wise generators then satisfy the local detailed balance relation for $(i,j) \in \mathscr{R}^{(r)}$ (cf. Sec. S2.2 and [8, Eq. (4)])

$$\ln\left(\frac{\Lambda^{(r)}_{ij}}{\Lambda^{(r)}_{ji}}\right) = \ln\left(\frac{\lambda^{(+r)}_{(i,j)}(e_i)}{\lambda^{(-r)}_{(i,j)}(e_j)}\right) = \beta\, \Delta^+_r Q(e_i) + \Delta^+_r h_{\mathrm{i}}(e_i).$$

By (36) we obtain

$$\frac{\mathrm{d}}{\mathrm{d}t} H^{\mathrm{tot}}(t) = \sum_{i\in\mathscr{J}} \sum_{r\in\mathscr{R}} \sum_{(k,j)\in\mathscr{R}^{(r)}} \ln\left(\frac{p(e_i,t)\,\lambda^{(+r)}_{(k,j)}(e_k)}{p(e_i+e_j-e_k,t)\,\lambda^{(-r)}_{(k,j)}(e_j)}\right) \times (\lambda^{(+r)}_{(k,j)}(e_k) p(e_i,t) - \lambda^{(-r)}_{(k,j)}(e_j) p(e_i+e_j-e_k,t))$$

and hence substituting (S2.9) yields the multi-reservoir EPR of a consistently modeled CTMC [8]

$$\frac{\mathrm{d}}{\mathrm{d}t} H^{\mathrm{tot}}(t) = \sum_{i\in\mathscr{J}} \sum_{r\in\mathscr{R}} \sum_{j>i} \ln\left(\frac{p(e_i,t)\,\Lambda^{(r)}_{ij}}{p(e_j,t)\,\Lambda^{(r)}_{ji}}\right) \times (\Lambda^{(r)}_{ij}\, p(e_i,t) - \Lambda^{(r)}_{ji}\, p(e_j,t)).$$

Importantly, neglecting the contribution of individual reservoirs can result in an underestimation of the EPR; cf. [1, Eq. (3.81)].

### S2.4. MACROSCOPIC CONSISTENCY OF LOCAL DETAILED BALANCE FOR OPEN CHEMICAL REACTION NETWORKS

We review the consistency of (12) in the deterministic limit of an ideal dilute solution under the mass action assumption. Hence, we take the macroscopic limit of the Gibbs free energy of the state and the propensity functions for a large volume $\mathsf{V}$, where all molecules are accounted for in real valued concentrations and fluctuations are negligible.

For positive concentrations $\tilde{u}_d > 0$ under the large volume assumption, we typically have $a^{(r)}_d \ll u_d$, which implies $\frac{u_d!}{(u_d - a^{(r)}_d)!} \approx u_d^{a^{(r)}_d}$. Let $g$ be the potential of an ideal dilute solution, where the total Gibbs free energy decomposes into a sum of Gibbs free energies for each species $\mathsf{U}_d$. Then

$$\begin{aligned} g(\mathsf{V}\tilde{u}) &\approx \mathsf{V} \sum_{d\in\mathscr{S}} \int_0^{\tilde{u}_d} (g^\circ_d + \beta^{-1}\ln(y_d))\,\mathrm{d}y_d \\ \Delta^+_r g(\mathsf{V}\tilde{u}) &\approx v_r(\mathsf{V}^{-1}\nabla_{\tilde{u}}) g(\mathsf{V}\tilde{u}) \\ &= \sum_{d\in\mathscr{S}} (b^{(r)}_d - a^{(r)}_d)(g^\circ_d + \beta^{-1}\ln(\tilde{u}_d)) \\ \lambda^+_r(\mathsf{V}\tilde{u}) &\approx \kappa^+_r \mathsf{V} \left(\prod_{l=1}^{L} z_l^{s^{(r)}_l}\right)\left(\prod_{d\in\mathscr{S}} \tilde{u}_d^{a^{(r)}_d}\right), \\ \lambda^-_r(\mathsf{V}\tilde{u}) &\approx \kappa^-_r \mathsf{V} \left(\prod_{l=1}^{L} z_l^{t^{(r)}_l}\right)\left(\prod_{d\in\mathscr{S}} \tilde{u}_d^{b^{(r)}_d}\right), \end{aligned}$$

where $g^\circ_d$ is the standard Gibbs free energy of species $\mathsf{U}_d$, $\tilde{u} = \mathsf{V}^{-1}u$, and the difference operator $\Delta^+_r$ be comes a differential operator with $\nabla_{\tilde{u}}$ denoting the gradient operator w.r.t. the components of $\tilde{u}$. Further we identify $\mu_l = \mu^\circ_l + \beta^{-1}\ln(z_l)$, with standard potential energy $\mu^\circ_l$, and observe that $-W^{(\pm r)}_r$ precisely represents the change in Gibbs free energy of the chemostat species in the macroscopic limit. Thus, if the species $\mathsf{Z}_1,\ldots,\mathsf{Z}_L$ and the core system $\mathsf{U}$ were to form a closed system, then the r.h.s. of the thermodynamic consistency relation for open CRNs (12), i.e., $\Delta^+_r Q(u) + k_{\mathrm{B}}T\,\Delta^+_r h_{\mathrm{i}}(u) = W^{(r)}_r - \Delta^+_r g(u)$, would converge in the macroscopic limit to the negative change in macroscopic Gibbs free energy of the total system upon the occurrence of reaction $\mathfrak{R}_r$ in forward direction. So, in the macroscopic limit, (12), i.e.,

$$\ln\left(\frac{\lambda_r^+(\mathsf{V}\tilde{u})}{\lambda_r^-(\mathsf{V}\tilde{u})}\right) = \beta\left(-\sum_{l=1}^{L}(t_l^{(r)} - s_l^{(r)})\mu_l - \Delta_r^+ g(\mathsf{V}\tilde{u})\right), \tag{S2.10}$$

is consistent with (5) under the assumption that the chemostat is an ideal dilute solution.

Substituting the macroscopic mass action approximation and the approximate definitions of $\Delta_r^+ g(\mathsf{V}\tilde{u})$ and $\mu_l$ for ideal dilute solution in (S2.10) yields the well known relation for the macroscopic rate constants [11]

$$\begin{aligned}\ln(K_{\text{eq}}^{(r)}) &= \ln\left(\frac{\kappa_r^+}{\kappa_r^-}\right)\\ &= -\sum_{l=1}^{L}(t_l^{(r)} - s_l^{(r)})\beta\mu_l^\circ\\ &\quad -\sum_{d\in\mathscr{S}}(b_d^{(r)} - a_d^{(r)})\beta g_d^\circ,\end{aligned}$$

where $K_{\text{eq}}^{(r)}$ is the equilibrium constant of reaction $\mathfrak{R}_r$.

Consistency in the mesoscopic limit can be established by dropping the chemostat assumptions on the chemostat species Z and noting that the sum of the potential of the chemostat species and $g$ represents the mesoscopic Gibbs free energy of the total system. This total Gibbs free energy and the respective mesoscopic rates then satisfy Equation (5).

#### S2.5. ADIABATIC AND NON-ADIABATIC ENTROPY PRODUCTION

The central equality (38) motivates the definition of the so-called housekeeping heat $q^{\text{hk}}(t)$ [5, 12, 13] at trajectory level

$$\begin{aligned}&\beta q^{\text{hk}}(t)\\ &:= \sum_{r\in\mathscr{R}}\int_0^t \ln\left(\frac{\lambda_r^+(U(s^-),s)\pi(U(s^-),s)}{\lambda_r^-(U(s^-)+\nu_r,s)\pi(U(s^-)+\nu_r,s)}\right)\mathrm{d}R_r^+(s)\\ &\quad+\sum_{r\in\mathscr{R}}\int_0^t \ln\left(\frac{\lambda_r^-(U(s^-),s)\pi(U(s^-),s)}{\lambda_r^+(U(s^-)-\nu_r,s)\pi(U(s^-)-\nu_r,s)}\right)\mathrm{d}R_r^-(s),\end{aligned} \tag{S2.11}$$

where we again generalized to time-dependent protocols $\Gamma$, using the instantaneous stationary distribution $\pi(u,s)$. The housekeeping heat captures the portion of the total heat dissipation $q(t)$ along a trajectory $U_{[0,t]}$ that is required to maintain the instantaneous NESS of the system at any time $s\in[0,t]$. In other words, $q^{\text{hk}}(t)$ quantifies the dissipated heat that can be attributed to broken detailed balance. Consequently, $q^{\text{hk}}(t)$ vanishes in the case that the instantaneous stationary distribution $\pi(\cdot,s)$ is an equilibrium distribution for all $s\in[0,t]$, which is the case for any manipulated closed CRN – that is without driving. Since the stationary distribution $\pi(u,s)$ is only an auxiliary quantity, which is fully determined by the propensity functions $\lambda_r^\varepsilon(u,s)$ for all $s\in[0,t]$ (cf. Eq. (15)), the definition is valid without assuming stationarity. In turn, this allows to distinguish it from the excess heat, defined as

$$q^{\text{ex}}(t) := q(t) - q^{\text{hk}}(t),$$

which quantifies the additional heat dissipated during relaxation to the (non-equilibrium) steady state.

Similarly, there is the notion of adiabatic entropy production [13]

$$h^{\text{a}}(t) = \beta q^{\text{hk}}(t),$$

which is indeed also defined by the r.h.s. of (S2.11). In the tradition of stochastic thermodynamics, a manipulation protocol that satisfies the quasistatic ad-hoc assumption of instantaneous stationarity $p(\cdot,s)=\pi(\cdot,s)$ for all $s\in[0,t]$ is called an adiabatic protocol [13]. This notion should not be confused with notion of an adiabatic process in classical equilibrium thermodynamics, which refers to vanishing heat exchange. For general time-dependent protocols, the quasistatic assumption is an ideal limiting notion associated with an infinitely slow manipulation, such that relaxation to steady state becomes much faster compared to variations in the manipulation. The quasistatic limit can be approximated arbitrarily well by any manipulation protocol, but it can be realized exactly only by constant protocols at stationarity [1]. In this exact case, it holds $h^{\text{tot}}(t) = h^{\text{a}}(t)$.

The adiabatic entropy production allows for the decomposition of the total entropy production into an adiabatic term and the non-adiabatic entropy production

$$h^{\text{na}}(t) := h^{\text{tot}}(t) - h^{\text{a}}(t).$$

By (28) and (31) the non-adiabatic entropy production satisfies

$$h^{\text{na}}(t) = h^{\text{sys}}(t) + \beta q^{\text{ex}}(t).$$

On the other hand, (32) implies that

$$h^{\text{a}}(t) = \beta q^{\text{mes}}(t) + \sum_{\varepsilon}\sum_{r\in\mathscr{R}}\int_0^t \ln\left(\frac{\pi(U(s^-),s)}{\pi(U(s^-)+\varepsilon\nu_r,s)}\right)\mathrm{d}R_r^\varepsilon(s),$$

which – consistently with the houskeeping interpretation – contains only reactive changes. In turn, by (34) we obtain

$$\begin{aligned}&h^{\text{na}}(t)\\ &= h^{\mathsf{U}}(t) - \sum_{\varepsilon}\sum_{r\in\mathscr{R}}\int_0^t \ln\left(\frac{\pi(U(s^-),s)}{\pi(U(s^-)+\varepsilon\nu_r,s)}\right)\mathrm{d}R_r^\varepsilon(s)\\ &= \sum_{\varepsilon}\sum_{r\in\mathscr{R}}\int_0^t \ln\left(\frac{p(U(s^-),s)/\pi(U(s^-),s)}{p(U(s^-)+\varepsilon\nu_r,s)/\pi(U(s^-)+\varepsilon\nu_r,s)}\right)\mathrm{d}R_r^\varepsilon(s)\\ &\quad -\int_0^t \partial_s \ln(p(U(s),s))\,\mathrm{d}s.\end{aligned}$$

For further intuition, consider a manipulated closed system (i.e., without driving). Then $\pi(\cdot,s)$ is always an equilibrium distribution since a closed system always converges to equilibrium. Hence, $h^{\text{a}}(t)=0$ irrespective of whether the manipulation is adiabatic or not. In this case $h^{\text{tot}}(t)=h^{\text{na}}(t)$. Now additionally assume an ideally adiabatic, time-varying protocol $\Gamma$ such that the process satisfies the quasistatic ad-hoc assumption $p(\cdot,s)=\pi(\cdot,s)$ for all $s\in[0,t]$. Then the non-adiabatic entropy production reduces to [1, Eq. (4.69)]

$$h^{\text{na}}(t) = -\int_0^t \partial_s \ln(\pi(U(s),s))\,\mathrm{d}s,$$

where $\pi(\cdot,s)$ does no follow the chemical master equation. Nevertheless, for a sufficiently well-behaved protocol, it is straightforward that in this case

$$\mathbb{E}[h^{\mathrm{na}}(t)] = -\int_0^t \sum_u \pi(u,s)\,\partial_s \ln(\pi(u,s))\,\mathrm{d}s = 0.$$

According to [1], the expected non-adiabatic entropy production does generally not vanish in the quasistatic limit of slow manipulation if the intitial and the final stationary distributions differ.

This example shows that the "housekeeping" interpretation is sound even at the trajectory level, while the interpretation of adiabatic entropy production as the fraction of entropy production that remains under an ideally adiabatic protocol is not always true. Therefore, adiabatic entropy production is better understood as housekeeping entropy production, and non-adiabatic entropy production as excess entropy production.

Finally, the two quantities $h^{\mathrm{a}}(t)$ and $h^{\mathrm{na}}(t)$ exhibit important statistical properties such as non-negative expectations [13]

$$\begin{aligned}\mathbb{E}[h^{\mathrm{a}}(t)] &\geq 0,\\ \mathbb{E}[h^{\mathrm{na}}(t)] &\geq 0.\end{aligned}$$

### S2.6. STOCHASTIC CONSISTENCY AND STATISTICAL TIME REVERSAL

In Sections III A and III B, we reviewed thermodynamic consistency relations derived from equilibrium statistical mechanics under coarse-graining (cf. Supplement Sec. S2.1). These relations ensure that the mesoscopic CRN description remains compatible with macroscopic thermodynamics, the underlying microscopic ensemble picture, and the detailed-balance condition (4). In particular, they provide a consistent link between kinetic rates and mesoscopic thermodynamic quantities that underlies the trajectory-level functionals introduced in Sec. III C.

We now turn to a complementary notion of consistency, namely *stochastic consistency*. Here the basic question is whether the path statistics of the CRN reproduce the fluctuation relations of stochastic thermodynamics, i.e., whether the ratio of forward and suitably time-reversed trajectory likelihoods agrees with the trajectory-level total entropy production [3, 14]. Unless stated otherwise, we allow for deterministic external manipulation.

We begin by defining the time-reversed process on the interval $[0,t]$. The time-reversed reaction counters are

$$\tilde{R}_r^+(\tau) := R_r^-(t) - R_r^-(t-\tau), \qquad \tilde{R}_r^-(\tau) := R_r^+(t) - R_r^+(t-\tau),$$

for all $\tau \in [0,t]$. The corresponding time-reversed state trajectory $\tilde{U}(\tau) := U(t-\tau)$ therefore satisfies

$$\tilde{U}(\tau) = \tilde{U}(0) + \sum_{r\in\mathscr{R}} \nu_r \big(\tilde{R}_r^+(\tau) - \tilde{R}_r^-(\tau)\big)$$

for all $\tau \in [0,t]$. In the following, we use $\sim$ as the time-reversal operator.

To derive the propensities of the time-reversed counters, we make use of mesoscopic consistency; see Sec. S2.2. Consider the full closed CRN obtained by reintroducing the chemostat species $\mathsf{Z}$ with mesoscopic copy numbers $z' = z\mathsf{V}$, instead of treating them as fixed. The augmented process $(U,Z')$ is then a Markov jump process without $(\mathsf{U},\mathsf{Z})$-parallel reaction channels. If the full system is driven by a protocol $\Gamma(s)$ on $[0,t]$ and we define the reversed protocol by

$$\tilde{\Gamma}(\tau) := \Gamma(t-\tau), \qquad \tau \in [0,t],$$

then the propensities of the exact time-reversed augmented process are given by the standard reverse-time formula for time-inhomogeneous Markov chains [15, p. 58]:

$$\begin{aligned}
&\tilde{\lambda}_r^{\pm}(u,z',\tilde{\Gamma}(\tau),\tau)\\
&= \lim_{h\searrow 0} \frac{1}{h}\mathbb{P}\big(\tilde{U}(\tau+h) = u\pm\nu_r, \tilde{Z}'(\tau+h) = z'\pm\gamma_r \,\big|\, \dots\big)\\
&= \lim_{h\searrow 0} \frac{1}{h}\mathbb{P}\big(U(t-\tau) = u, Z'(t-\tau) = z' \,\big|\, \dots\big)\\
&\quad\times \frac{\mathbb{P}\big(U(t-\tau-h) = u\pm\nu_r, Z'(t-\tau-h) = z'\pm\gamma_r\big)}{\mathbb{P}\big(U(t-\tau) = u, Z'(t-\tau) = z'\big)}\\
&= \lambda_r^{\mp}(u\pm\nu_r, z'\pm\gamma_r, \Gamma(t-\tau))\\
&\quad\times \frac{\mathbb{P}(\tilde{U}(\tau) = u\pm\nu_r, \tilde{Z}'(\tau) = z'\pm\gamma_r)}{\mathbb{P}(\tilde{U}(\tau) = u, \tilde{Z}'(\tau) = z')}.
\end{aligned}$$

Thus, the time-reversed Markov process does generally *not* satisfy the statistical time reversibility property when the original dynamics is time-inhomogeneous [15]. The general external manipulation $\Gamma$ under consideration includes the ability to alter the mean value of $Z'(s)$ for all times $s \geq 0$ by adjusting the propensities of reservoir exchange reactions, which were not explicitly modeled.

Reapplying the chemostat approximation means assuming $\gamma_r/z' \ll 1$, so that

$$\mathbb{P}(\tilde{Z}'(\tau) = z'\pm\gamma_r) \approx \mathbb{P}(\tilde{Z}'(\tau) = z').$$

The resulting propensities of the conditional time-reversed core process $\tilde{U}(\tau)$ are therefore

$$\begin{aligned}
\tilde{\lambda}_r^{\pm}(u,\tau) &= \lambda_r^{\mp}(u\pm\nu_r, t-\tau)\,\frac{\tilde{p}(u\pm\nu_r,\tau)}{\tilde{p}(u,\tau)}\\
&= \lambda_r^{\mp}(u\pm\nu_r, t-\tau)\,\frac{p(u\pm\nu_r,t-\tau)}{p(u,t-\tau)},
\end{aligned} \tag{S2.12}$$

where $\tilde{p}(u,\tau) := \mathbb{P}(\tilde{U}(\tau) = u) = p(u,t-\tau)$.

Using (3) together with (S2.12), the probability evolution of the exact time-reversed process follows as

$$\begin{aligned}
\partial_\tau \tilde{p}(u,\tau) = -\sum_{r\in\mathscr{R}} &\lambda_r^+(u-\nu_r, t-\tau)\tilde{p}(u-\nu_r,\tau)\\
&+ \lambda_r^-(u+\nu_r, t-\tau)\tilde{p}(u+\nu_r,\tau)\\
&- (\lambda_r^+(u,t-\tau) + \lambda_r^-(u,t-\tau))\tilde{p}(u,\tau).
\end{aligned}$$

We now distinguish between *statistical* and *thermodynamic* reversibility.

**Definition S2.1.** *A stationary Markov process U is called statistically reversible if*

$$U_{[0,t]} \overset{d}{=} \tilde{U}_{[0,t]}$$

*for all* $t \geq 0$.

At the level of the state process alone, this notion does not distinguish between U-parallel reaction channels. Indeed, $U_{[0,t]}$ is equivalently represented by

$$\left\{U(0), (\overline{R}^{+}_{\alpha}(s), \overline{R}^{-}_{\alpha}(s)) : \alpha \in \mathscr{C}\right\}_{s\in[0,t]},$$

where the effective counters $\overline{R}^{\pm}_{\alpha}$ were introduced in Sec. III E. As similarly observed in [16, 17], this representation forgets which particular U-parallel channel was used. In a thermodynamic setting this loss of channel information is problematic, because different channels typically correspond to different chemostat exchanges. We therefore need a channel-resolved notion of reversibility.

**Definition S2.2.** *A stationary Markov CRN U is called statistically reversible if*

$$C_{[0,t]} \overset{d}{=} \tilde{C}_{[0,t]}$$

*for all* $t \geq 0$*, where*

$$C_{[0,t]} := \left\{U(0), (R^{+}_{r}(s), R^{-}_{r}(s)) : r \in \mathscr{R}\right\}_{s\in[0,t]},$$
$$\tilde{C}_{[0,t]} := \left\{\tilde{U}(0), (\tilde{R}^{+}_{r}(s), \tilde{R}^{-}_{r}(s)) : r \in \mathscr{R}\right\}_{s\in[0,t]}.$$

In words, Definition S2.2 states that, based on the full channel-resolved trajectory, one cannot tell whether the process was run forward or backward in time solely based on the trajectory statistics. Each trajectory is equally likely in both temporal directions.

For notational convenience we introduce the multivariate counting process

$$R_{[0,t]} := \left\{(R^{+}_{r}(s), R^{-}_{r}(s)) : r \in \mathscr{R}\right\}_{s\in[0,t]},$$

and similarly its time reverse $\tilde{R}_{[0,t]}$, so that $C_{[0,t]} = (U(0), R_{[0,t]})$.

The Kullback–Leibler divergence between the path measures of $C_{[0,t]}$ and $\tilde{C}_{[0,t]}$ quantifies the degree of statistical irreversibility. In particular,

$$\mathbb{D}(\mathbb{P}^{C}_{[0,t]} \| \mathbb{P}^{\tilde{C}}_{[0,t]}) = 0$$

for all $t \geq 0$ if and only if the CRN is statistically reversible.

This still does not capture the notion of *thermodynamic* reversibility relevant to fluctuation relations. To see why, consider an extended protocol

$$\Gamma^{\mathrm{fb}}(s) := \Gamma(s)\mathbb{1}\{s \in [0,t)\} + \tilde{\Gamma}(s)\mathbb{1}\{s \in [t,2t]\}$$

on $[0,2t]$, which first runs the forward protocol and then the reversed protocol. The first half of the corresponding extended process is just the original forward process. We denote the second half by $C'$ and refer to it as the backward process:

$$C'_{[0,t]} := \left\{U'(0), (R'^{+}_{r}(s), R'^{-}_{r}(s)) : r \in \mathscr{R}\right\}_{s\in[0,t]}.$$

By construction, the backward process is initialized with $U'(0) \overset{d}{=} U(t)$, i.e., with the final distribution of the forward process,

$$p'(u,0) = p(u,t),$$

and the counters $R'^{\pm}_{r}$ evolve under the reversed protocol, i.e., with propensities $(u,s) \mapsto \lambda^{\pm}_{r}(u,t-s)$. Thermodynamic reversibility means that a forward realization is as likely as the time-reversed realization generated by this backward experiment.

**Definition S2.3.** *A Markov CRN U following the protocol* $\Gamma$ *on* $[0,t]$ *is called thermodynamically reversible if*

$$C_{[0,t]} \overset{d}{=} \tilde{C}'_{[0,t]},$$

*where* $\tilde{C}'_{[0,t]} = (\tilde{U}'(0), \tilde{R}'_{[0,t]})$ *is the time-reversed backward process.*

The propensities of $\tilde{R}'$ are

$$\tilde{\lambda}'^{\pm}_{r}(u,\tau) = \lambda^{\mp}_{r}(u \pm \nu_r, \tau)\,\frac{p'(u \pm \nu_r, t-\tau)}{p'(u, t-\tau)}, \tag{S2.13}$$

where $p'(u,s) := \mathbb{P}(U'(s) = u)$ and $p'(u,0) = p(u,t)$. In the absence of external manipulation, $\tilde{C}_{[0,t]}$ and $\tilde{C}'_{[0,t]}$ trivially coincide in distribution, as both the initial distribution and propensities are identical.

Let $\zeta\colon \Omega \times [0,t] \to \mathbb{Z}^{|\mathscr{S}|}_{\geq 0}$ denote a sample path of either $U_{[0,t]}$, $\tilde{U}_{[0,t]}$, or $\tilde{U}'_{[0,t]}$, and let $\rho\colon \Omega \times [0,t] \to \mathbb{Z}^{2|\mathscr{R}|}_{\geq 0}$ denote a sample path of the channel-resolved counting processes $R_{[0,t]}$, $\tilde{R}_{[0,t]}$, or $\tilde{R}'_{[0,t]}$. Thus, for all such sample paths it holds

$$\zeta(\omega,s) = \zeta(\omega,0) + \sum_{r\in\mathscr{R}} \nu_r\big(\rho^{+}_{r}(\omega,s) - \rho^{-}_{r}(\omega,s)\big)$$

for all $\omega \in \Omega$ and all $s \in [0,t]$. The path measures $\mathbb{P}^{C}_{[0,t]}$ and $\mathbb{P}^{\tilde{C}'}_{[0,t]}$ assign a positive likelihood only to trajectories satisfying this constraint. Their Radon-Nikodym derivative therefore takes the standard form for multivariate counting processes

[18, 19]:

$$
\begin{aligned}
&\frac{\mathrm{d}\mathbb{P}^{C}_{[0,t]}}{\mathrm{d}\mathbb{P}^{\tilde{C}'}_{[0,t]}}(\zeta_{[0,t]},\rho_{[0,t]})\\
=&\frac{\mathrm{d}\mathbb{P}^{U(0)}}{\mathrm{d}\mathbb{P}^{\tilde{U}'(0)}}(\zeta(0))\frac{\mathrm{d}\mathbb{P}^{R|U(0)}_{[0,t]}}{\mathrm{d}\mathbb{P}^{\tilde{R}'|\tilde{U}'(0)}_{[0,t]}}(\zeta_{[0,t]},\rho_{[0,t]})\\
=&\frac{p(\zeta(0),0)}{\tilde{p}'(\zeta(0),0)}\\
&\times\exp\Bigg(-\sum_{r\in\mathscr{R}}\int_0^t\Big[\lambda_r^{+}(\zeta(s),s)+\lambda_r^{-}(\zeta(s),s)\\
&\qquad\qquad-\tilde{\lambda}_r^{\prime +}(\zeta(s),s)-\tilde{\lambda}_r^{\prime -}(\zeta(s),s)\Big]\,\mathrm{d}s\Bigg)\\
&\times\exp\left(\sum_{r\in\mathscr{R}}\int_0^t\ln\left(\frac{\lambda_r^{+}(\zeta(s^-),s)}{\tilde{\lambda}_r^{\prime +}(\zeta(s^-),s)}\right)\mathrm{d}\rho_r^{+}(s)\right)\\
&\times\exp\left(\sum_{r\in\mathscr{R}}\int_0^t\ln\left(\frac{\lambda_r^{-}(\zeta(s^-),s)}{\tilde{\lambda}_r^{\prime -}(\zeta(s^-),s)}\right)\mathrm{d}\rho_r^{-}(s)\right).
\end{aligned}
$$

This expression compares the likelihood of the same forward trajectory under the physical forward process and the time-reversed backward reference process.

To connect this Radon-Nikodym derivative with the likelihood-ratio expressions commonly used in stochastic thermodynamics, it is useful to make the underlying path densities explicit. A convenient reference density for counting processes is the local Janossy density [20, Ch. 7.3], often simply called the path likelihood in the stochastic thermodynamics literature. Including the initial distribution, the local Janossy density of $C$ is

$$
\begin{aligned}
&L^{C}_{[0,t]}(\zeta_{[0,t]},\rho_{[0,t]})\\
=&p(\zeta(0),0)\\
&\times\exp\left(-\sum_{r\in\mathscr{R}}\int_0^t\lambda_r^{+}(\zeta(s),s)+\lambda_r^{-}(\zeta(s),s)\,\mathrm{d}s\right)\\
&\times\exp\left(\sum_{r\in\mathscr{R}}\int_0^t\ln\big(\lambda_r^{+}(\zeta(s^-),s)\big)\,\mathrm{d}\rho_r^{+}(s)\right)\\
&\times\exp\left(\sum_{r\in\mathscr{R}}\int_0^t\ln\big(\lambda_r^{-}(\zeta(s^-),s)\big)\,\mathrm{d}\rho_r^{-}(s)\right).
\end{aligned}
\tag{S2.14}
$$

Analogous expressions for $L^{C'}_{[0,t]}$, $L^{\tilde{C}'}_{[0,t]}$, and $L^{\tilde{C}}_{[0,t]}$ are obtained by replacing the propensity functions with those of the respective processes. In particular,

$$
\frac{\mathrm{d}\mathbb{P}^{C}_{[0,t]}}{\mathrm{d}\mathbb{P}^{\tilde{C}'}_{[0,t]}}(\zeta_{[0,t]},\rho_{[0,t]})=\frac{L^{C}_{[0,t]}(\zeta_{[0,t]},\rho_{[0,t]})}{L^{\tilde{C}'}_{[0,t]}(\zeta_{[0,t]},\rho_{[0,t]})}.
$$

The next step is the identity

$$
L^{\tilde{C}'}_{[0,t]}(\zeta_{[0,t]},\rho_{[0,t]})=L^{C'}_{[0,t]}(\tilde{\zeta}_{[0,t]},\tilde{\rho}_{[0,t]}),\tag{S2.15}
$$

proved below in Sec. S2.7. We state this identity explicitly because it provides the bridge between the Radon-Nikodym derivative written entirely on the forward path space and the more familiar likelihood ratio involving a forward trajectory and the reversed backward trajectory. Combining (S2.15) with the previous display yields

$$
\frac{\mathrm{d}\mathbb{P}^{C}_{[0,t]}}{\mathrm{d}\mathbb{P}^{\tilde{C}'}_{[0,t]}}(\zeta_{[0,t]},\rho_{[0,t]})=\frac{L^{C}_{[0,t]}(\zeta_{[0,t]},\rho_{[0,t]})}{L^{C'}_{[0,t]}(\tilde{\zeta}_{[0,t]},\tilde{\rho}_{[0,t]})},\tag{S2.16}
$$

which is the form commonly used in stochastic thermodynamics; see, e.g., [5, Eq. (49)], [1, Eq. (4.9)], [21, Eqs. (32)–(33)], and [22, Eq. (81)].

Moreover, this likelihood ratio can be written entirely in terms of the forward trajectory:

$$
\begin{aligned}
&\frac{L^{C}_{[0,t]}(\zeta_{[0,t]},\rho_{[0,t]})}{L^{C'}_{[0,t]}(\tilde{\zeta}_{[0,t]},\tilde{\rho}_{[0,t]})}\\
=&\frac{p(\zeta(0),0)}{p(\zeta(t),t)}\\
&\times\exp\left(\sum_{r\in\mathscr{R}}\int_0^t\ln\left(\frac{\lambda_r^{+}(\zeta(s^-),s)}{\lambda_r^{-}(\zeta(s^-)+\nu_r,s)}\right)\mathrm{d}\rho_r^{+}(s)\right)\\
&\times\exp\left(\sum_{r\in\mathscr{R}}\int_0^t\ln\left(\frac{\lambda_r^{-}(\zeta(s^-),s)}{\lambda_r^{+}(\zeta(s^-)-\nu_r,s)}\right)\mathrm{d}\rho_r^{-}(s)\right).
\end{aligned}
$$

We emphasize this intermediate identity because it makes transparent how the path-space Radon-Nikodym derivative reduces to a purely forward, causal likelihood ratio.

The right-hand side of (S2.16) is directly identified with trajectory-level thermodynamic quantities. More precisely,

$$
\begin{aligned}
\frac{\mathrm{d}\mathbb{P}^{C}_{[0,t]}}{\mathrm{d}\mathbb{P}^{\tilde{C}'}_{[0,t]}}(\zeta_{[0,t]},\rho_{[0,t]})&=\exp\big(h^{\mathrm{U}}(t)+\beta q^{\mathrm{mes}}(t)\big)\\
&=\exp\big(h^{\mathrm{tot}}(t)\big),
\end{aligned}
\tag{S2.17}
$$

where a rigorous derivation is given in Sec. S2.7.

Following Horowitz [14], one may therefore call a stochastic process description *stochastically consistent* with thermodynamics if,

$$
\beta q(t)=\ln\left(\frac{\mathrm{d}\mathbb{P}^{C}_{[0,t]}}{\mathrm{d}\mathbb{P}^{\tilde{C}'}_{[0,t]}}(\zeta_{[0,t]},\rho_{[0,t]})\right)-h^{\mathrm{sys}}(t)
$$

holds pathwise, even under external manipulation. This means that the trajectory statistics are compatible with the thermodynamic pathwise identification of heat exchange, system entropy change, and total entropy production in Eqs. (28) and (31). In open CRNs, stochastic consistency enforces local detailed balance (12). This concept of stochastic consistency is not limited to discrete-state stochastic processes; it also extends to continuous-state processes, such as diffusion processes.

The KL divergence between the two path measures is

$$
\mathbb{D}(\mathbb{P}^{C}_{[0,t]}\|\mathbb{P}^{\tilde{C}'}_{[0,t]}):=\mathbb{E}\left[\ln\left(\frac{\mathrm{d}\mathbb{P}^{C}_{[0,t]}}{\mathrm{d}\mathbb{P}^{\tilde{C}'}_{[0,t]}}(U_{[0,t]},R_{[0,t]})\right)\right].
$$

Equation (S2.17) implies

$$\mathbb{D}(\mathbb{P}^{C}_{[0,t]}\|\mathbb{P}^{\tilde{C}'}_{[0,t]}) = \mathbb{E}[h^{\text{tot}}(t)] = H^{\text{tot}}(t). \tag{S2.18}$$

This motivates the following measure-theoretic definition of entropy production rate [23].

**Definition S2.4.** *Let U be a potentially time-dependent open CRN. Then*

$$e_p(t) := \lim_{h\searrow 0}\frac{1}{h}\mathbb{D}(\mathbb{P}^{C}_{[t,t+h]}\|\mathbb{P}^{\tilde{C}'}_{[t,t+h]})$$

*is called the instantaneous entropy production rate, and in the time-independent case*

$$e_p := \lim_{t\to\infty}\frac{1}{t}\mathbb{D}(\mathbb{P}^{C}_{[0,t]}\|\mathbb{P}^{\tilde{C}'}_{[0,t]})$$

*is called the limiting entropy production rate.*

Under stochastic consistency, this measure-theoretic definition agrees with the thermodynamic notions introduced earlier. Moreover, since

$$\mathbb{D}(\mathbb{P}^{C}_{[t,t+h]}\|\mathbb{P}^{\tilde{C}'}_{[t,t+h]}) \geq 0$$

for all $t,h\geq 0$, it immediately yields the second law for open CRNs,

$$e_p(t), e_p \geq 0,$$

in agreement with (III C).

We conclude by reviewing equivalent characterizations of thermodynamic equilibrium for non-explosive open CRNs, similar to [24, Thm. 2.5].

**Theorem S2.1.** *Let U be stationary with stationary distribution $\pi(u)$. Then the following are equivalent:*

*(a) U is thermodynamically reversible. In particular,*

$$\mathbb{D}(\mathbb{P}^{C}_{[0,t]}\|\mathbb{P}^{\tilde{C}}_{[0,t]}) = 0$$

*for all $t\geq 0$, and the time-reversed propensities satisfy*

$$\tilde{\lambda}^{\pm}_r(u,t) = \lambda^{\pm}_r(u).$$

*(b) $e_p = 0$.*

*(c) Stoichiometric Kolmogorov (Wegscheider) cycle condition: Any admissible closed cycle of states*

$$u_0 \xrightarrow{\varepsilon_1 r_1} u_1 \xrightarrow{\varepsilon_2 r_2} \dots \xrightarrow{\varepsilon_j r_j} u_n \xrightarrow{\varepsilon_{n+1} r_{n+1}} u_0,$$

*originating from a sequence of reaction channel uses $r_j$ in the directions $\varepsilon_j\in\{+,-\}$, such that $\varepsilon_j \nu_{r_j} = (u_j - u_{j-1})$, satisfies*

$$\frac{\lambda^{\varepsilon_1}_{r_1}(u_0)\lambda^{\varepsilon_2}_{r_2}(u_1)\cdots\lambda^{\varepsilon_{n+1}}_{r_{n+1}}(u_n)}{\lambda^{-\varepsilon_{n+1}}_{r_{n+1}}(u_0)\lambda^{-\varepsilon_n}_{r_n}(u_n)\cdots\lambda^{-\varepsilon_1}_{r_1}(u_1)} = 1.$$

*(d) U satisfies the detailed balance condition*

$$\lambda^{+}_r(u)\pi(u) = \lambda^{-}_r(u+\nu_r)\pi(u+\nu_r)$$

*for all u and all $r\in\mathscr{R}$.*

The stoichiometric cycle condition, together with thermodynamic consistency, is the discrete analogue of path-independence for conservative force fields. It characterizes whether the effective forces driving the CRN derive from a potential or contain genuinely nonconservative components.

A proof of Theorem S2.1 is given in Sec. S2.7. The limiting distribution need not be unique; the equivalent equilibrium conditions remain valid when the stationary distribution $\pi$ is a mixture of equilibrium distributions on individual stoichiometric compatibility classes.

### S2.7. PROOFS FOR STOCHASTIC THERMODYNAMICS

*Proof of* (20). For brevity we denote with $q(t)$ and $Q(t)$ only the reaction-related contribution in the dissipated heat. Utilizing the Poisson-type process definition in [25, p. 7] the expectations of the stochastic integrals can be rewritten as expectations of Lebesgue/Riemann integrals:

$$\begin{aligned}\mathbb{E}[q(t)] &= \sum_{r\in\mathscr{R}}\mathbb{E}\left[\int_0^t \Delta^{+}_r Q(U(s^-))\,\mathrm{d}R^{+}_r(s)\right.\\ &\qquad\left.+\int_0^t \Delta^{-}_r Q(U(s^-))\,\mathrm{d}R^{-}_r(s)\right]\\ &= \sum_{r\in\mathscr{R}}\mathbb{E}\left[\int_0^t \Delta^{+}_r Q(U(s))\lambda^{+}_r(U(s))\,\mathrm{d}s\right.\\ &\qquad\left.+\int_0^t \Delta^{-}_r Q(U(s))\lambda^{-}_r(U(s))\,\mathrm{d}s\right],\end{aligned}$$

where the expectation of $q$ is taken over the distribution of all trajectories $U_{[0,t]}$. Exchanging expectation and integration by Fubini's theorem and differentiating with respect to time gives

$$\begin{aligned}\dot{Q}(t) = \sum_{r\in\mathscr{R}}\big\{&\mathbb{E}[\Delta^{+}_r Q(U(t))\lambda^{+}_r(U(t))]\\ &+\mathbb{E}[\Delta^{-}_r Q(U(t))\lambda^{-}_r(U(t))]\big\}.\end{aligned}$$

Evaluating the expectations yields

$$\dot{Q}(t) = \sum_u\sum_{r\in\mathscr{R}}(\Delta^{+}_r Q(u)\lambda^{+}_r(u) + \Delta^{-}_r Q(u)\lambda^{-}_r(u))p(u,t).$$

Equation (20) then follows by rearranging the summation of the last term and using the property of Equation (11). □

Before proving (S2.15), we make explicit that the trajectory-level thermodynamic quantities $h^{\text{tot}}(\zeta,t)$, $h^{\text{U}}(\zeta,t)$, etc. are evaluated along a fixed path $(\zeta_{[0,t]},\rho_{[0,t]})$; for brevity, we suppress the explicit dependence on $\rho$. The proof relies on two simple time-reversal identities, one for the continuous-time derivative of the state probability and one for counting-process integrals.

For $\tau = t - s$, the chain rule gives

$$\begin{aligned}\partial_s \ln p(\zeta(s),s) &= -\partial_\tau \ln p(\tilde{\zeta}(\tau), t-\tau) \\ &= -\partial_\tau \ln \tilde{p}(\tilde{\zeta}(\tau),\tau),\end{aligned} \tag{S2.19}$$

and therefore

$$\int_0^t \partial_s \ln p(\zeta(s),s)\,\mathrm{d}s = -\int_0^t \partial_s \ln \tilde{p}(\tilde{\zeta}(s),s)\,\mathrm{d}s.$$

Next, let $\tau_i^{(\pm r)}$, $i \in \mathbb{N}$, denote the jump times of the forward counter $\rho_r^{\pm}$, and define

$$\tilde{\tau}_i^{(\pm r)} := t - \tau_i^{(\mp r)}.$$

At such times,

$$\zeta(\tau_i^{(\pm r)}-) = \zeta((t-\tilde{\tau}_i^{(\mp r)})^-) = \tilde{\zeta}(\tilde{\tau}_i^{(\mp r)}+) = \tilde{\zeta}(\tilde{\tau}_i^{(\mp r)}-) \mp \nu_r.$$

Hence, for any real-valued function $\chi(u,s)$,

$$\int_0^t \chi(\zeta(s^-),s)\,\mathrm{d}\rho_r^{\pm}(s) = \int_0^t \chi(\tilde{\zeta}(s^-)\mp\nu_r, t-s)\,\mathrm{d}\tilde{\rho}_r^{\mp}(s), \tag{S2.20}$$

which is the required change-of-variables formula for counting-process integrals under time reversal.

These two identities already imply the involution property of the population entropy. Using (29), (S2.19) and (S2.20), we obtain

$$\begin{aligned}h^{\mathsf{U}}(\zeta,t) =& -\int_0^t \partial_s \ln p(\zeta(s),s)\,\mathrm{d}s \\ &+ \sum_{r\in\mathscr{R}} \int_0^t \ln\left(\frac{p(\zeta(s^-),s)}{p(\zeta(s^-)+\nu_r,s)}\right)\mathrm{d}\rho_r^+(s) \\ &+ \sum_{r\in\mathscr{R}} \int_0^t \ln\left(\frac{p(\zeta(s^-),s)}{p(\zeta(s^-)-\nu_r,s)}\right)\mathrm{d}\rho_r^-(s) \\ =& \int_0^t \partial_s \ln p(\tilde{\zeta}(s),t-s)\,\mathrm{d}s \\ &- \sum_{r\in\mathscr{R}} \int_0^t \ln\left(\frac{p(\tilde{\zeta}(s^-),t-s)}{p(\tilde{\zeta}(s^-)+\nu_r,t-s)}\right)\mathrm{d}\tilde{\rho}_r^+(s) \\ &- \sum_{r\in\mathscr{R}} \int_0^t \ln\left(\frac{p(\tilde{\zeta}(s^-),t-s)}{p(\tilde{\zeta}(s^-)-\nu_r,t-s)}\right)\mathrm{d}\tilde{\rho}_r^-(s) \\ =& -h^{\mathsf{U}}(\tilde{\zeta},t).\end{aligned}$$

This "odd-parity" property under time-reversal of the argument is called involution [1], and it similarly holds for all other trajectory quantities, even the internal ones. For example,

$$\Delta_r^{\pm} Q(u,s) = -\Delta_r^{\mp} Q(u\pm\nu_r,s), \qquad W_r^{(r)}(s) = -W_r^{(-r)}(s).$$

In what follows, we use this involution property whenever it is convenient.

*Proof of* (S2.15). The proof proceeds in three steps. First, we rewrite the backward Janossy density evaluated on the time-reversed path as a functional of the forward path. Second, we rewrite the initial factor by introducing the population entropy change of the backward process. Third, we use (S2.13) to show that all remaining terms combine into the Janossy density of the time-reversed backward process.

By definition of the local Janossy density,

$$\begin{aligned}&L^{C'}_{[0,t]}(\tilde{\zeta}_{[0,t]},\tilde{\rho}_{[0,t]}) \\ =& p'(\tilde{\zeta}(0),0) \\ &\times \exp\left(-\sum_{r\in\mathscr{R}}\int_0^t \lambda_r^+(\tilde{\zeta}(s),t-s)+\lambda_r^-(\tilde{\zeta}(s),t-s)\,\mathrm{d}s\right) \\ &\times \exp\left(\sum_{r\in\mathscr{R}}\int_0^t \ln\big(\lambda_r^+(\tilde{\zeta}(s^-),t-s)\big)\,\mathrm{d}\tilde{\rho}_r^+(s)\right) \\ &\times \exp\left(\sum_{r\in\mathscr{R}}\int_0^t \ln\big(\lambda_r^-(\tilde{\zeta}(s^-),t-s)\big)\,\mathrm{d}\tilde{\rho}_r^-(s)\right).\end{aligned}$$

Using (S2.20), this becomes

$$\begin{aligned}&L^{C'}_{[0,t]}(\tilde{\zeta}_{[0,t]},\tilde{\rho}_{[0,t]}) \\ =& p'(\tilde{\zeta}(0),0) \\ &\times \exp\left(-\sum_{r\in\mathscr{R}}\int_0^t \lambda_r^+(\zeta(s),s)+\lambda_r^-(\zeta(s),s)\,\mathrm{d}s\right) \\ &\times \exp\left(\sum_{r\in\mathscr{R}}\int_0^t \ln\big(\lambda_r^-(\zeta(s^-)+\nu_r,s)\big)\,\mathrm{d}\rho_r^+(s)\right) \\ &\times \exp\left(\sum_{r\in\mathscr{R}}\int_0^t \ln\big(\lambda_r^+(\zeta(s^-)-\nu_r,s)\big)\,\mathrm{d}\rho_r^-(s)\right).\end{aligned} \tag{S2.21}$$

We next rewrite the initial factor. Let

$$h^{\mathsf{U}}_{\mathrm{B}}(\tilde{\zeta},t) = \ln\frac{p'(\tilde{\zeta}(0),0)}{p'(\tilde{\zeta}(t),t)}$$

denote the population entropy change of the backward process along the reversed path. Then

$$\begin{aligned}p'(\tilde{\zeta}(0),0) &= p'(\tilde{\zeta}(t),t)\exp\big(h^{\mathsf{U}}_{\mathrm{B}}(\tilde{\zeta},t)\big) \\ &= \tilde{p}'(\zeta(0),0)\exp\big(-h^{\mathsf{U}}_{\mathrm{B}}(\zeta,t)\big),\end{aligned} \tag{S2.22}$$

where the second line uses the involution property.

Finally, by first using (30), and subsequently (S2.13), (S2.19), and (S2.20), the backward population entropy along the forward path is rewritten as

$$\begin{aligned}
&h_{\mathrm{B}}^{\mathrm{U}}(\zeta,t)\\
&=\sum_{r\in\mathscr{R}}\int_0^t\Bigg\{\lambda_r^+(\zeta(s)-\nu_r,s)\frac{p'(\zeta(s)-\nu_r,t-s)}{p'(\zeta(s),t-s)}\\
&\qquad-(\lambda_r^+(\zeta(s),s)+\lambda_r^-(\zeta(s),s))\\
&\qquad+\lambda_r^-(\zeta(s)+\nu_r,s)\frac{p'(\zeta(s)+\nu_r,t-s)}{p'(\zeta(s),t-s)}\Bigg\}\mathrm{d}s\\
&\quad+\sum_{r\in\mathscr{R}}\int_0^t\ln\left(\frac{p'(\zeta(s^-),t-s)}{p'(\zeta(s^-)+\nu_r,t-s)}\right)\mathrm{d}\rho_r^+(s)\\
&\quad+\sum_{r\in\mathscr{R}}\int_0^t\ln\left(\frac{p'(\zeta(s^-),t-s)}{p'(\zeta(s^-)-\nu_r,t-s)}\right)\mathrm{d}\rho_r^-(s)\\
&=\sum_{r\in\mathscr{R}}\int_0^t\Big[\tilde{\lambda}_r'^+(\zeta(s),s)+\tilde{\lambda}_r'^-(\zeta(s),s)\\
&\qquad-(\lambda_r^+(\zeta(s),s)+\lambda_r^-(\zeta(s),s))\Big]\mathrm{d}s\\
&\quad+\sum_{r\in\mathscr{R}}\int_0^t\ln\left(\frac{\lambda_r^-(\zeta(s^-)+\nu_r,s)}{\tilde{\lambda}_r'^+(\zeta(s^-),s)}\right)\mathrm{d}\rho_r^+(s)\\
&\quad+\sum_{r\in\mathscr{R}}\int_0^t\ln\left(\frac{\lambda_r^+(\zeta(s^-)-\nu_r,s)}{\tilde{\lambda}_r'^-(\zeta(s^-),s)}\right)\mathrm{d}\rho_r^-(s).
\end{aligned}\tag{S2.23}$$

Substituting (S2.23) into (S2.22), and then (S2.22) into (S2.21), all terms reorganize into the Janossy density of the time-reversed backward process evaluated on the forward path. This yields (S2.15). ☐

*Proof of* (S2.17). We now use (S2.16) together with the explicit Janossy densities (S2.14) and (S2.21). This gives

$$\begin{aligned}
&\frac{L^C_{[0,t]}(\zeta_{[0,t]},\rho_{[0,t]})}{L^{C'}_{[0,t]}(\tilde{\zeta}_{[0,t]},\tilde{\rho}_{[0,t]})}\\
&=\frac{p(\zeta(0),0)}{p'(\tilde{\zeta}(0),0)}\\
&\quad\times\exp\left(-\sum_{r\in\mathscr{R}}\int_0^t\lambda_r^+(\zeta(s),s)+\lambda_r^-(\zeta(s),s)\,\mathrm{d}s\right)\\
&\quad\times\exp\left(\sum_{r\in\mathscr{R}}\int_0^t\ln\big(\lambda_r^+(\zeta(s^-),s)\big)\,\mathrm{d}\rho_r^+(s)\right)\\
&\quad\times\exp\left(\sum_{r\in\mathscr{R}}\int_0^t\ln\big(\lambda_r^-(\zeta(s^-),s)\big)\,\mathrm{d}\rho_r^-(s)\right)\\
&\quad\times\exp\left(\sum_{r\in\mathscr{R}}\int_0^t\lambda_r^+(\zeta(s),s)+\lambda_r^-(\zeta(s),s)\,\mathrm{d}s\right)\\
&\quad\times\exp\left(-\sum_{r\in\mathscr{R}}\int_0^t\ln\big(\lambda_r^-(\zeta(s^-)+\nu_r,s)\big)\,\mathrm{d}\rho_r^+(s)\right)\\
&\quad\times\exp\left(-\sum_{r\in\mathscr{R}}\int_0^t\ln\big(\lambda_r^+(\zeta(s^-)-\nu_r,s)\big)\,\mathrm{d}\rho_r^-(s)\right).
\end{aligned}$$

The continuous integral factors cancel. Since $\tilde{\zeta}(0)=\zeta(t)$ and the backward process is initialized from the final forward law, $p'(\tilde{\zeta}(0),0)=p(\zeta(t),t)$. Hence,

$$\begin{aligned}
&\frac{L^C_{[0,t]}(\zeta_{[0,t]},\rho_{[0,t]})}{L^{C'}_{[0,t]}(\tilde{\zeta}_{[0,t]},\tilde{\rho}_{[0,t]})}\\
&=\frac{p(\zeta(0),0)}{p(\zeta(t),t)}\\
&\quad\times\exp\left(\sum_{r\in\mathscr{R}}\int_0^t\ln\left(\frac{\lambda_r^+(\zeta(s^-),s)}{\lambda_r^-(\zeta(s^-)+\nu_r,s)}\right)\mathrm{d}\rho_r^+(s)\right)\\
&\quad\times\exp\left(\sum_{r\in\mathscr{R}}\int_0^t\ln\left(\frac{\lambda_r^-(\zeta(s^-),s)}{\lambda_r^+(\zeta(s^-)-\nu_r,s)}\right)\mathrm{d}\rho_r^-(s)\right).
\end{aligned}$$

The first factor is exactly $\exp(h^{\mathrm{U}}(\zeta,t))$, while the remaining two factors give $\exp(\beta q^{\mathrm{mes}}(\zeta,t))$. This proves the first line of (S2.17). The second line then follows from (34). ☐

*Proof of Thm. S2.1.* We prove the implications

$$(d)\Leftrightarrow(c),\qquad(d)\Leftrightarrow(b),\qquad(a)\Rightarrow(b),\qquad(d)\Rightarrow(a),$$

which together yield the equivalence of (a)–(d).

$(d)\implies(c)$. Let

$$u_0\xrightarrow{\varepsilon_1 r_1}u_1\xrightarrow{\varepsilon_2 r_2}\dots\xrightarrow{\varepsilon_{n+1}r_{n+1}}u_{n+1}=u_0$$

be an admissible closed cycle. For each $j\in\{1,\dots,n+1\}$, the detailed-balance condition gives

$$\lambda_{r_j}^{\varepsilon_j}(u_{j-1})\,\pi(u_{j-1})=\lambda_{r_j}^{-\varepsilon_j}(u_j)\,\pi(u_j).$$

Hence

$$\frac{\lambda_{r_j}^{\varepsilon_j}(u_{j-1})}{\lambda_{r_j}^{-\varepsilon_j}(u_j)}=\frac{\pi(u_j)}{\pi(u_{j-1})}.$$

Multiplying these identities along the cycle and using $u_{n+1}=u_0$ yields

$$\prod_{j=1}^{n+1}\frac{\lambda_{r_j}^{\varepsilon_j}(u_{j-1})}{\lambda_{r_j}^{-\varepsilon_j}(u_j)}=\prod_{j=1}^{n+1}\frac{\pi(u_j)}{\pi(u_{j-1})}=\frac{\pi(u_{n+1})}{\pi(u_0)}=1,$$

which is exactly the stoichiometric Kolmogorov cycle condition.

$(c)\implies(d)$. Since $\pi$ is stationary, its support is a union of closed communicating classes. It is therefore enough to verify detailed balance on each closed communicating class $\mathscr{K}$ contained in a stoichiometric compatibility class.

Fix such a class $\mathscr{K}$ and choose a reference state $u_*\in\mathscr{K}$. For any $u\in\mathscr{K}$, choose an admissible path

$$u_*=u_0\xrightarrow{\varepsilon_1 r_1}u_1\xrightarrow{\varepsilon_2 r_2}\dots\xrightarrow{\varepsilon_m r_m}u_m=u$$

and define

$$\Phi_{\mathscr{K}}(u):=\prod_{j=1}^{m}\frac{\lambda_{r_j}^{\varepsilon_j}(u_{j-1})}{\lambda_{r_j}^{-\varepsilon_j}(u_j)},\qquad\Phi_{\mathscr{K}}(u_*):=1.$$

By the cycle condition in (c), $\Phi_{\mathscr{K}}(u)$ is independent of the chosen path: if two admissible paths connect $u_*$ to $u$, then traversing one path forward and the other backward gives a closed cycle, and the corresponding ratio equals 1.

Consequently, for every admissible edge

$$u \xrightarrow{+r} u+\nu_r$$

inside $\mathscr{K}$, we have

$$\frac{\Phi_{\mathscr{K}}(u+\nu_r)}{\Phi_{\mathscr{K}}(u)} = \frac{\lambda_r^+(u)}{\lambda_r^-(u+\nu_r)}.$$

Equivalently,

$$\Phi_{\mathscr{K}}(u)\,\lambda_r^+(u) = \Phi_{\mathscr{K}}(u+\nu_r)\,\lambda_r^-(u+\nu_r) \qquad \text{(S2.24)}$$

for all admissible reaction edges in $\mathscr{K}$.

Equation (S2.24) shows that $\Phi_{\mathscr{K}}$ is a reversible, and therefore invariant, measure on $\mathscr{K}$; see Ref. [26, Remark 19.19]. Since $\mathscr{K}$ is irreducible and $\pi(\cdot \mid \mathscr{K})$ is a stationary probability measure on $\mathscr{K}$, invariant measures on $\mathscr{K}$ are unique up to a multiplicative constant [26, Remark 19.19]. Hence there exists $c_{\mathscr{K}} > 0$ such that

$$\pi(u \mid \mathscr{K}) = c_{\mathscr{K}}\,\Phi_{\mathscr{K}}(u), \qquad u \in \mathscr{K}.$$

Substituting this into (S2.24) yields

$$\lambda_r^+(u)\pi(u \mid \mathscr{K}) = \lambda_r^-(u+\nu_r)\pi(u+\nu_r \mid \mathscr{K})$$

for all admissible edges in $\mathscr{K}$.

Since the support of $\pi$ is a union of such closed communicating classes, detailed balance holds on the whole support of $\pi$, and therefore on the whole state space.

$(d) \implies (b)$. At stationarity, the entropy production rate admits the edgewise representation

$$e_p = \sum_u \sum_{r\in\mathscr{R}} \Psi\big(\lambda_r^+(u)\pi(u),\, \lambda_r^-(u+\nu_r)\pi(u+\nu_r)\big), \quad \text{(S2.25)}$$

where

$$\Psi(a,b) := (a-b)\ln\frac{a}{b} \geq 0$$

for all $a,b > 0$, with equality if and only if $a = b$. Under detailed balance, every term in (S2.25) vanishes, hence $e_p = 0$.

$(b) \implies (d)$. Again by (S2.25), all summands are nonnegative. Therefore, if $e_p = 0$, every term must vanish individually. Hence

$$\lambda_r^+(u)\pi(u) = \lambda_r^-(u+\nu_r)\pi(u+\nu_r)$$

for all $u$ and all $r \in \mathscr{R}$, i.e., detailed balance holds.

$(a) \implies (b)$. By thermodynamic reversibility,

$$\mathbb{P}^C_{[0,t]} = \mathbb{P}^{\tilde{C}}_{[0,t]}$$

for all $t \geq 0$. Hence

$$\mathbb{D}\Big(\mathbb{P}^C_{[0,t]} \,\Big\|\, \mathbb{P}^{\tilde{C}}_{[0,t]}\Big) = 0$$

for all $t \geq 0$. Applying Definition S2.4 gives $e_p = 0$.

$(d) \implies (a)$. Assume detailed balance. Since the process is stationary, the initial law is invariant under time reversal:

$$\mathbb{P}^{U(0)} = \mathbb{P}^{\tilde{U}(0)}.$$

Moreover, by (S2.12),

$$\tilde{\lambda}_r^{\pm}(u,\tau) = \lambda_r^{\mp}(u \pm \nu_r)\frac{\pi(u\pm\nu_r)}{\pi(u)}.$$

Using detailed balance, this becomes

$$\tilde{\lambda}_r^{\pm}(u,\tau) = \lambda_r^{\pm}(u)$$

for all $\tau \geq 0$, all $u$, and all $r \in \mathscr{R}$. Thus the forward process and its time reverse have the same initial law and the same jump intensities. Equivalently,

$$\mathbb{D}\Big(\mathbb{P}^C_{[0,t]} \,\Big\|\, \mathbb{P}^{\tilde{C}}_{[0,t]}\Big) = 0$$

for all $t \geq 0$, so $C_{[0,t]} \overset{d}{=} \tilde{C}_{[0,t]}$. Hence the CRN is thermodynamically reversible. □


[1] L. Peliti and S. Pigolotti, *Stochastic thermodynamics: an introduction* (Princeton University Press, 2021).

[2] U. Seifert, First and second law of thermodynamics at strong coupling, Physical review letters **116**, 020601 (2016).

[3] C. Jarzynski, Stochastic and macroscopic thermodynamics of strongly coupled systems, Physical Review X **7**, 011008 (2017).

[4] H.-P. Breuer and F. Petruccione, *The theory of open quantum systems* (OUP Oxford, 2002).

[5] T. Schmiedl and U. Seifert, Stochastic thermodynamics of chemical reaction networks, The Journal of Chemical Physics **126** (2007).

[6] D. Kondepudi and I. Prigogine, *Modern thermodynamics: from heat engines to dissipative structures* (John Wiley & Sons, 2014) Chap. 15.

[7] M. Kardar, Classical statistical mechanics, in *Statistical Physics of Particles* (Cambridge University Press, 2007) p. 98–125.

[8] M. Esposito, Stochastic thermodynamics under coarse graining, Physical Review E—Statistical, Nonlinear, and Soft Matter Physics **85**, 041125 (2012).

[9] H. Qian and H. Ge, *Stochastic Chemical Reaction Systems in Biology* (Springer International Publishing, 2021).

[10] J. Schnakenberg, Network theory of microscopic and macroscopic behavior of master equation systems, Rev. Mod. Phys. **48**, 571 (1976).

[11] T. Renner, E. R. Cohen, T. Cvitas, J. G. Frey, B. Holström, K. Kuchitsu, R. Marquardt, I. Mills, F. Pavese, M. Quack,

J. Stohner, H. L. Strauss, M. Takami, and A. J. Thor, *Quantities, units and symbols in physical chemistry* (The Royal Society of Chemistry, 2007).

[12] Y. Oono and M. Paniconi, Steady state thermodynamics, Progress of Theoretical Physics Supplement **130**, 29 (1998).

[13] M. Esposito and C. Van den Broeck, Three detailed fluctuation theorems, Physical review letters **104**, 090601 (2010).

[14] J. M. Horowitz, Diffusion approximations to the chemical master equation only have a consistent stochastic thermodynamics at chemical equilibrium, The Journal of chemical physics **143** (2015).

[15] S. Asmussen, S. Asmussen, and S. Asmussen, *Applied probability and queues*, Vol. 2 (Springer, 2003).

[16] A detailed balanced reaction network is sufficient but not necessary for its Markov chain to be detailed balanced, Discrete and Continuous Dynamical Systems - B **20**, 1077.

[17] C. Jia, D.-Q. Jiang, and Y. Li, Detailed balance, local detailed balance, and global potential for stochastic chemical reaction networks, Advances in Applied Probability **53**, 886 (2021).

[18] J. Jacod, Multivariate Point Processes: Predictable Projection, Radon-Nikodym Derivatives, Representation of Martingales, Zeitschrift für Wahrscheinlichkeitstheorie und verwandte Gebiete **31**, 235 (1975).

[19] R. Boel, P. Varaiya, and E. Wong, Martingales on jump processes. II: Applications, SIAM Journal on Control **13**, 1022 (1975).

[20] D. J. Daley and D. Vere-Jones, *An Introduction to the Theory of Point Processes: Volume I: Elementary Theory and Methods* (Springer Science & Business Media, 2006).

[21] U. Seifert, Stochastic thermodynamics: From principles to the cost of precision, Physica A: Statistical Mechanics and its Applications **504**, 176 (2018).

[22] C. Van den Broeck and M. Esposito, Ensemble and trajectory thermodynamics: A brief introduction, Physica A: Statistical Mechanics and its Applications **418**, 6 (2015).

[23] D.-Q. Jiang and D. Jiang, *Mathematical Theory of Nonequilibrium Steady States: On the Frontier of Probability and Dynamical Systems* (Springer Science & Business Media, 2004).

[24] X.-J. Zhang, H. Qian, and M. Qian, Stochastic theory of nonequilibrium steady states and its applications. part i, Physics Reports **510**, 1 (2012).

[25] P. M. Brémaud, *Point Processes and Queues: Martingale Dynamics*, Vol. 50 (Springer, 1981).

[26] A. Klenke, *Probability Theory : A Comprehensive Course*, 3rd ed. (Springer International Publishing, Cham, 2020).
